# Native Hydrogen/Deuterium Exchange Ion Mobility Mass Spectrometry of Structured DNA Oligonucleotides

**DOI:** 10.64898/2026.09.03.749223

**Authors:** Matthieu Ranz, Romane Guisiano, Eric Largy, Valérie Gabelica

## Abstract

Hydrogen/deuterium exchange coupled to mass spectrometry (HDX/MS) is a powerful technique to probe nucleic acid secondary structures and dynamics, but its ability to resolve conformers with identical masses remains limited. To overcome this challenge, we integrated ion mobility spectrometry (IMS) into our native HDX/MS workflow, and tested the approach on a variety of model DNA G-quadruplex structures. We show several examples of human telomeric sequence oligonucleotides where complexes of the same mass differ in their collision cross section, and each gas-phase conformational ensembles corresponds to unique solution exchange behaviors, allowing kinetic analysis beyond what is possible with HDX/native MS alone. But we also found examples where several gas-phase populations separated in ion mobility have exactly the same solution exchange behavior, suggesting that conformational rearrangements occur either during electrospray or at later stages in the gas phase. Finally, we show how IMS filtering can be leveraged to distinguish groups of non-specific cation binding on a given conformational ensemble, as indicated by populations with different masses and same exchange rates. These findings establish IMS as an essential tool for complementing HDX/MS in the characterization of structural polymorphism and conformational ensembles in DNA oligonucleotides.

## 1. Introduction

Hydrogen-Deuterium eXchange (HDX) is a powerful method that measures the kinetics of H/D exchange reactions in deuterated solvents. HDX can be monitored by^1^H-NMR,^1,2^ or mass spectrometry (HDX/MS).^3–6^ In macromolecules, the rate of exchange depends on both hydrogen bonding involvement and solvent accessibility, thus providing detailed information about their structure and dynamics. For these reasons, HDX/MS has become a widely adopted technique for studying protein structures and interactions.^7–13^ A typical experiment involves a bottom-up workflow: the reaction is quenched, followed by analyte denaturation and enzymatic digestion prior to mass spectrometry analysis. But while this workflow facilitates robust comparative studies between samples, it inherently limits the ability to resolve and analyze complex conformational ensembles in solution.

Our group has demonstrated that HDX/MS is an excellent tool for characterizing DNA oligonucleotides, whose exchange kinetics inform both their structure and solution stability.^14–16^ Furthermore, we have evidenced that lower-stability conformers undergo cooperative unfolding across multiple sites, a feature that can be leveraged to detect and characterize transient (un)folding intermediates.^15,16^ A key difference in our approach is the use of native MS, a powerful method for studying noncovalent complexes formed by oligonucleotides.^17^ Native MS relies on the conservation of noncovalent interactions from solution to the gas phase, allowing for the accurate determination of binding stoichiometries, affinities, and kinetics.

To illustrate the potential resolving power of integrating HDX with native MS, we focused on G-quadruplexes (G4s). G4s constitute a major class of nucleic acid secondary structures formed by the stacking of at least two guanine tetrads, further stabilized by coordination with a cation (typically K^+^) positioned between the tetrad planes.^18^ Each planar tetrad consists of four guanines linked by eight hydrogen bonds (Figure 1A), making G4s particularly suited for HDX/MS studies. Crucially, G4s can co-exist in complex ensembles of conformers in dynamic equilibrium, differing by their topology and number of tetrads.^16,19,20^ Topologies are typically defined by the relative 5’ → 3’ orientation of the four guanine tracts: parallel (all four tracks oriented similarly), antiparallel (two tracks opposite to the other two), and hybrid (three tracks opposite the fourth one) (Figure 1B). The ability to easily shift these conformational equilibria using simple solution manipulations, such as changes in cation type and concentration, temperature, co-solvents, or small molecule binders,^18,20–22^ makes G4s *ad hoc* systems for testing the capabilities of HDX/native MS.

**Figure 1.**
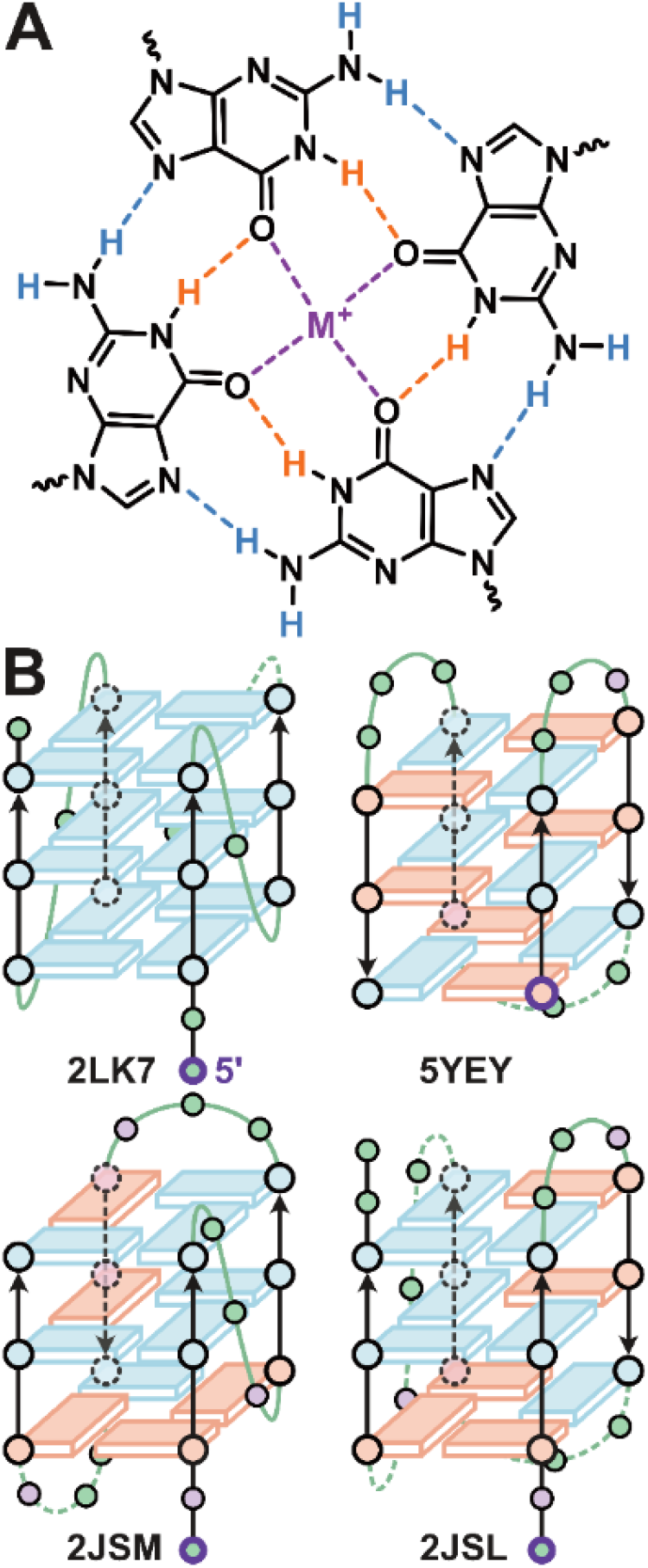
A. Structure of a G-tetrad with exchangeable amino protons in blue and imino protons in orange. The potassium cation (purple) is coordinated by the oxygen atoms from two consecutive G-tetrads. B. Examples of G4 topologies as defined by the relative 5’-3’ orientation of G-tracts (black) linked by loops (green): parallel (2LK7), antiparallel (5YEY), hybrid-1 (2JSM) and hybrid-2 (2JSL). The guanines are shown in blue (anti glycosidic bond orientation) and salmon (syn), the thymines in green and adenines in purple. Potassium cations are not shown for the sake of clarity.

We previously demonstrated the capability of HDX/native MS on complex conformational mixtures of G4s. Under native MS conditions, the exchange kinetics of two- and three-tetrad G4 ensembles were selectively measured because they are mass-resolved thanks to differences in their K^+^ coordination stoichiometry (1 and 2, respectively).^23–25^ However, HDX/native MS alone fails to measure selectively the kinetics of same-mass conformers; instead, the measured HDX rates represent an average weighted by the respective abundances of these species.

To overcome this limitation, we explored the possibility of separating analytes of same mass using ion-mobility spectrometry (IMS). IMS is an effective tool for studying noncovalent structures and interactions in nucleic acids,^26–29^ including G-quadruplexes.^30–32^ We will show how IMS deconvolution can be leveraged to resolve conformation-specific HDX kinetics, overcoming the limitations of native MS alone. Moreover, while examining various G-quadruplexes (Table 1), we also found two other interesting case scenarios and alternative uses of HDX/native IMS-MS, to filter out the contribution of non-specific adducts, and to distinguish if different gas-phase conformers have been issued from different solution-phase conformations. The interpretation of exchange rates in terms of G-quadruplex biophysics, as carried out for HDX/native MS alone,^16^ will be reported elsewhere.

**Table 1.**
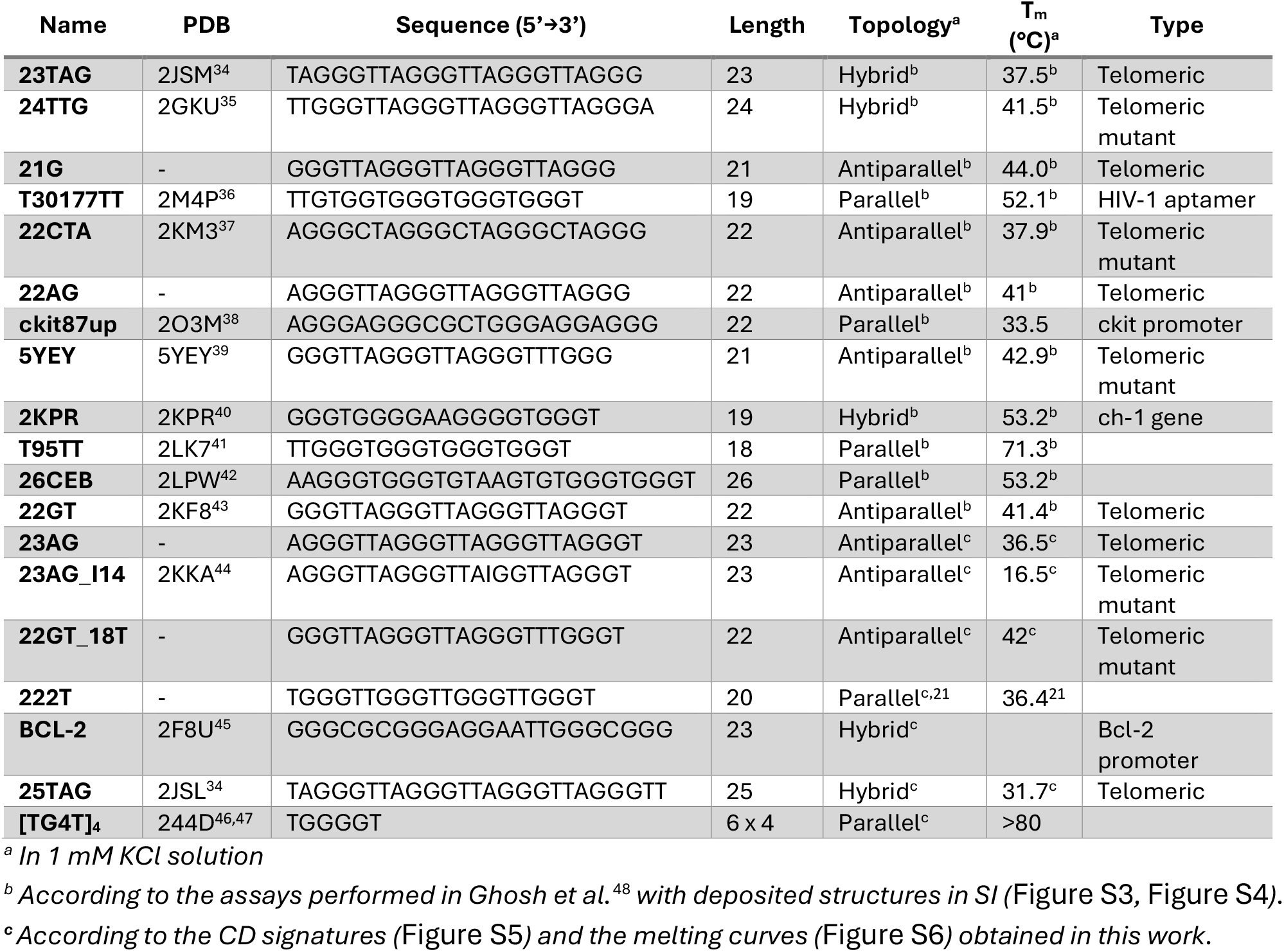
Oligonucleotides used in this assay, with their PDB codes and their topology in 100 mM TMAA + 1 mM KCl.

## 2. Materials and methods

### 2.1. Sample preparation

Oligodeoxynucleotides sequences (Table 1) were purchased lyophilized from Eurogentec (Seraing, Belgium) and resuspended in D2O (99.9% D atom Sigma-Aldrich, Saint-Quentin Fallavier, France) to reach approximately 1 mM. Desalting on 1.5-mL Amicon Ultra-0.5 centrifugal filters (3 kDa) was carried out four times with 150 mM NH_4_OAc/D_2_O buffer then four times with D_2_O, for 10 minutes at 14,000 g. The accurate concentration was measured with a UV spectrophotometer (Uvikon XS, Secomam, France) using the nearest-neighbors approach to determine extinction coefficients._33_

HDX/MS samples were prepared at 90% deuterium content by mixing 100 µM of the DNA solutions prepared as above with 100 mM trimethylammonium acetate (TMAA, 1M in H_2_O, SantaCruz Biotechnology, Heidelberg, Germany), 1 mM KCl (99.999% trace metal basis, Sigma-Aldrich, Saint-Quentin Fallavier, France) and D_2_O. The sequences T95TT, T30177TT were folded in 0.5 mM KCl due to their high stability even at this concentration. 24TTG and 222T were investigated at 1 mM KCl and 0.1 mM KCl. Samples were left at least one night at 4°C to ensure complete folding. The exchanging buffer was prepared extemporaneously by mixing 1 mM KCl and 100 mM TMAA in ULC/MS – CC/SFC water (Biosolve Chimie, Dieuze, France).

### 2.2. IMS-MS

The HDX/IMS experiments were carried out on an Agilent DT-IMS Q-TOF 6560. In drift-tube IMS (Figure S1), ions travel through a gas-filled (here, helium) drift tube under a uniform electric field. The ion mobility depends on the species’ collision cross section (CCS) and charge, allowing for separation.^49–51^ The resulting arrival time distributions (ATD) can be converted into CCS values that directly relate to the gas-phase structures of the analytes (Figure S2).

The instrument was tuned according to the recommendations set in our lab in negative mode using “Optimized parameters” for drift and “Compromised parameters” for the post-IMS region, as defined in reference (Figure S1).^52^ The impact of key instrument parameters on the deuteration and signal-to-noise ratio was assessed by systematic variation of their values; methods and results are summarized in Supporting Information (Figure S7-Figure S17, Table S1).

The acquisition time of IMS-MS experiments was 2 minutes per timepoint. The fragmentor voltage was set by default at 320 V to carry experiments in native conditions and increased to improve the signal-to-noise ratio where relevant. Similarly, the nebulizer pressure default value was 12 psi, the drying gas flow rate 2 L/min and the drift voltage −650 V. An internal CCS calibration was systematically performed to adjust the helium pressure in the drift tube before each experiment session using the G-quadruplex [(dTG4T)4·(NH4^+^)3] for which the experimental CCS values should match the theoretical values (780−788 Å² for the 5- charge state and 730−734 Å² for the 4- charge state).^24^ The 2D IMS figures were generated for each oligonucleotide using the last time point recorded and were exported in .csv format from Agilent MassHunter IM-MS Browser (version B.08.00). ATD to CCS conversion method and results are available in Supporting Information, including ATDs at different drift voltages, corresponding fitting results and CCS distributions for each oligonucleotide (Figure S18-Figure S54). Unless otherwise mentioned, the 4- charge states were analyzed, as they generally had much higher abundances and improved IMS separation.

Collision-Induced Unfolding (CIU) experiments were conducted with the parameters above and varying the fragmentor voltage between 290 V and 470 V with a 20 V increment per segment (2 min of acquisition per voltage). The data was collected with CIU Suite 2,^53^ then plotted in R. All CIU experiments are available in SI (Figure S55-Figure S98).

### 2.3. HDX/IMS-MS

We used our previously described continuous flow setup:^14^ a syringe containing the sample and another containing the exchanging buffer are set in parallel in a two-channels Chemyx Fusion 4000 syringe pump. Both syringes were connected to a mixing tee (IDEX, ref U-466) with its output connected to the mass spectrometer by a removable PEEK tube. The flow rates of the syringe pump were set to 30, 35 or 40 µL/min with a 1:9 sample:diluent ratio, and 8 output tubes of different volumes were used (Figure S99) to generate different time points. The final DNA concentration is therefore 10 µM and the D2O content drops to 9%.

To access individual exchange rates, the data analysis workflow combines three nested separation steps:

1. For all analytes: *m/z* separation of different cations stoichiometries
2. For each *m/z*-separated ensemble: IMS separation by arrival time of different CCS
3. For each *m/z*- and IMS-separated analyte: deconvolution of bimodal isotopic distribution resulting from transient unfolding, where necessary

Data extraction was done using the IMSBrowser software. After selecting the m/z region of interest for the most abundant charge states (step 1), arrival time distribution (ATD) deconvolution was performed in R (4.3.2) using a minimum number of *n* Gaussian peaks, following Equation 1 where counts represent the signal intensity, *t_D_* is the drift time in ms, and *A_i_*, *x_c_*_,*i*_ and *w_i_* are the height, center and standard deviation of peak *i*, respectively.

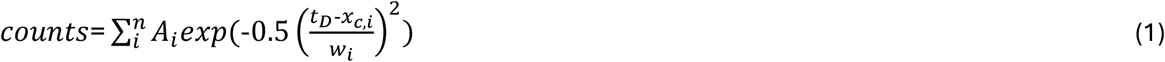

The selection of the ATD zones (step 2) was done by selecting parts of the Gaussian populations large enough to obtain a sufficient signal-to-noise ratio, while avoiding contamination by other populations. The raw mass data was converted into numbers of unexchanged sites (NUS) for each m/z and ATD-resolved species using Equation 2 in the OligoR software.^14,15^

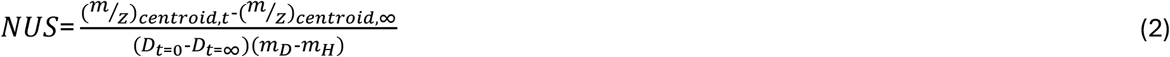

with *D_t_*_=0_=90%, *D_t_*_=∞_=9%, *m_H_* (resp. *m_D_*) the mass of hydrogen (resp. deuterium), (*^m^*⁄*_z_*)*_centroid_*_,∞_ standing for the centroid mass when the exchange is complete, and (*^m^*⁄*_z_*)*_centroid_*_,*t*_ the centroid at time *t*. Exchange curves for mass and IMS-resolved conformer populations are given in Supporting Information, alongside the density maps showing the selected m/z and ATD regions and the CCS distribution of the corresponding m/z-selected species (Figure S100-Figure S137). Deconvolutions of bimodal isotopic distributions was achieved with OligoR^15^ (Figure S138-Figure S173), and are given in Supporting information together with the corresponding exchange kinetics in Figure S174-Figure S187.

## 3. Results

### 3.1. Case 1: Conformation-specific HDX/MS of complex mixtures

We assess here the ability of IMS to separate conformers of same *m/z*, but different CCS, to access their specific HDX rates and exchange behaviors. To that end, we used model oligonucleotides with four repeats of the human telomeric GGGTTA motif (23TAG, 5YEY), which often form dynamic mixtures of conformers.^48,54^ Discrete variations of the oligonucleotide sequence impacts the structural equilibrium,^55^ which is valuable for method development. HDX/native MS provides information on the dynamics of these conformational ensembles, but cannot resolve conformers of same mass.^16^

#### 3.1.1. Accessing exchange rates and mechanisms for closely related conformers

In 1 mM KCl, the telomeric sequence 23TAG forms a mixture of 3-tetrad hybrid-1 and hybrid-2 conformers,^34^ as well as an ensemble of 2-tetrad antiparallel conformers.^56^ The 2- and 3-tetrad ensembles can be separated by mass under native conditions because they specifically bind one and two potassium cations, respectively (detected as MK and MK_2_ in Figure 2). However, the two 3- tetrad conformers are not separated by mass, within MK_2_. The ATD of MK_2_ exhibits two distinct peaks, corresponding to the hybrid-1 (higher CCS) and hybrid-2 (lower CCS) populations.^56^ This bimodal ATD can be deconvoluted into two Gaussian distributions, which serve as a basis for selecting two arrival time windows (Figure 2), to maximize the signal of each conformer while minimizing cross-contamination.

**Figure 2.**
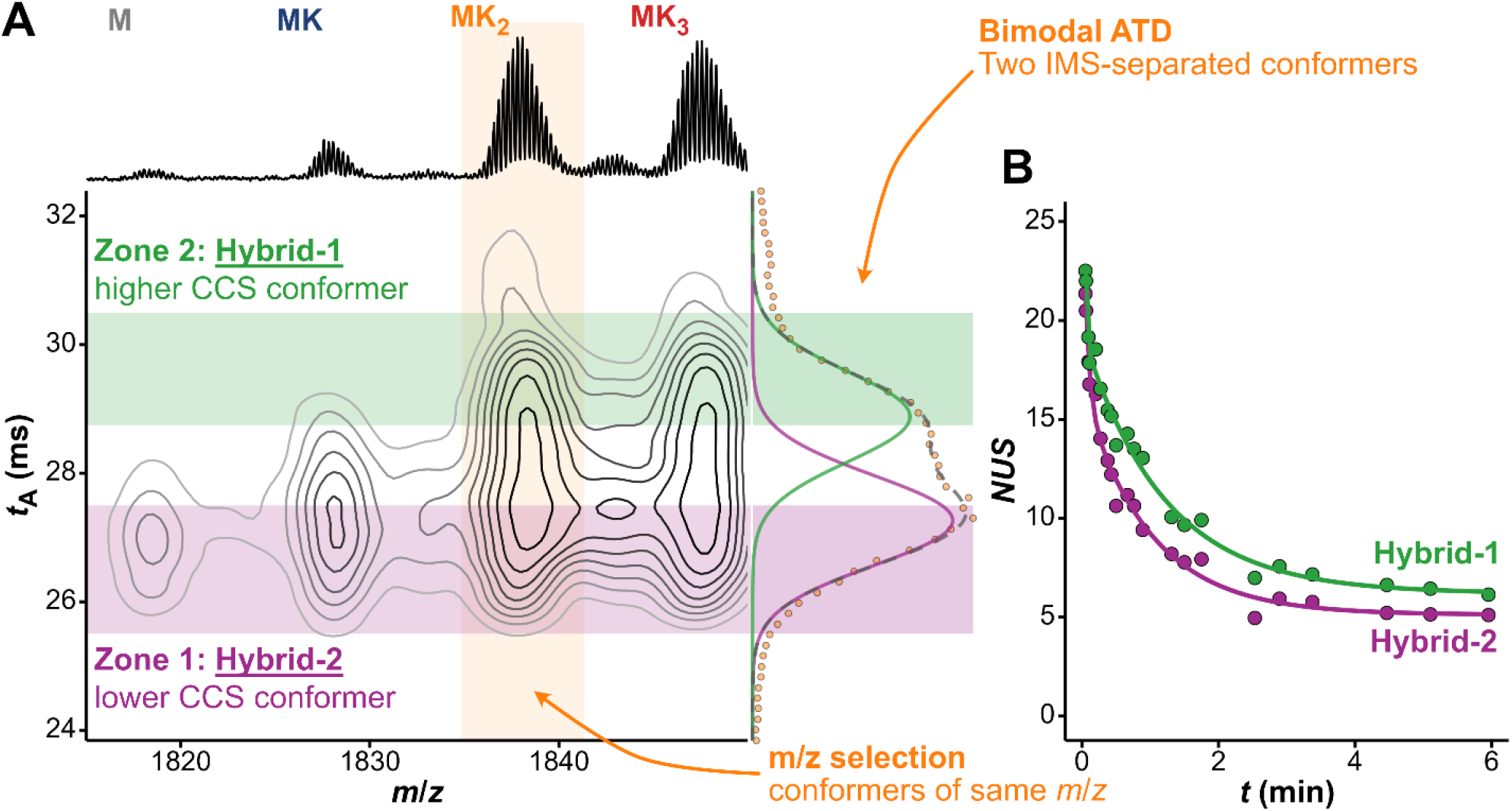
A. IMS-MS density map of the oligonucleotide 23TAG in TMAA/KCl buffer, zoomed on the 4- charge state (fragmentor voltage: 370 V): mass separation on the x-axis and shape separation of the y-axis. The corresponding mass spectrum is shown on top, and the arrival time (t_A_) distribution for the selected MK_2_ species (selection zone in orange) on the right (orange points). Deconvolution with two Gaussians (purple and green lines) allows selecting appropriate t_A_ zones (purple and green areas) of distinct conformations. B. Exchange kinetics of the two conformers of 23TAG·2K^+^, derived from the data at the intersections of the m/z and t_A_ zones of panel A.

The fragmentor voltage was increased from 320 to 370 V, *i.e.*, just below the onset of gas-phase unfolding (Figure S188A), to offset the relatively low signal-to-noise ratio (S/N) and limited resolution of the IMS populations. At higher voltage, the selected *m/z* and ATD regions are more clearly discriminated (Figure S188B), enabling better-resolved HDX kinetics (Figure 2B, Figure S188C). The hybrid-2 conformer displays a faster apparent exchange rate than its hybrid-1 counterpart. Different ATD zone widths were tested to assess the method robustness, with no significant changes observed in the kinetics (Figure S189). The increase in fragmentor voltage also enables the discrimination of two IMS populations within MK, revealing the presence of at least two 2-tetrad ensembles with distinct exchange rates (Figure S118).

The isotopic distributions of both IMS-separated conformers are bimodal (Figure 3A); their deconvolution can inform on their stability and tendency to unfold dynamically. Both conformers undergo local exchange of individual sites at similar rates, characterized by a gradual shift of the isotopic distribution (Figure 3B; EX2 exchange), suggesting that they have a similar Δ*G*^0^.^16^ However, these conformers have distinct EX1 exchange contributions, wherein the highly-exchanged isotopic population (at low m/z) is the result from species that have visited a partially or fully unfolded state (at least once), at the unfolding rate *k_op_*.^8,15,57,58^ Thus, the hybrid-2 conformer undergoes partial unfolding at a faster rate than the hybrid-1 (Figure 3C; *k_HDX_*_,*EX*1_ ≈ *k_op_* ≈ 0.57 min^-1^ vs. 1.76 min^-1^, respectively). In both cases, the isotopic populations undergoing EX1 are not fully exchanged at short time points, consistent with the EX1-competent transient species being only partially unfolded, such as into a G-triplex structure.^59–61^

**Figure 3.**
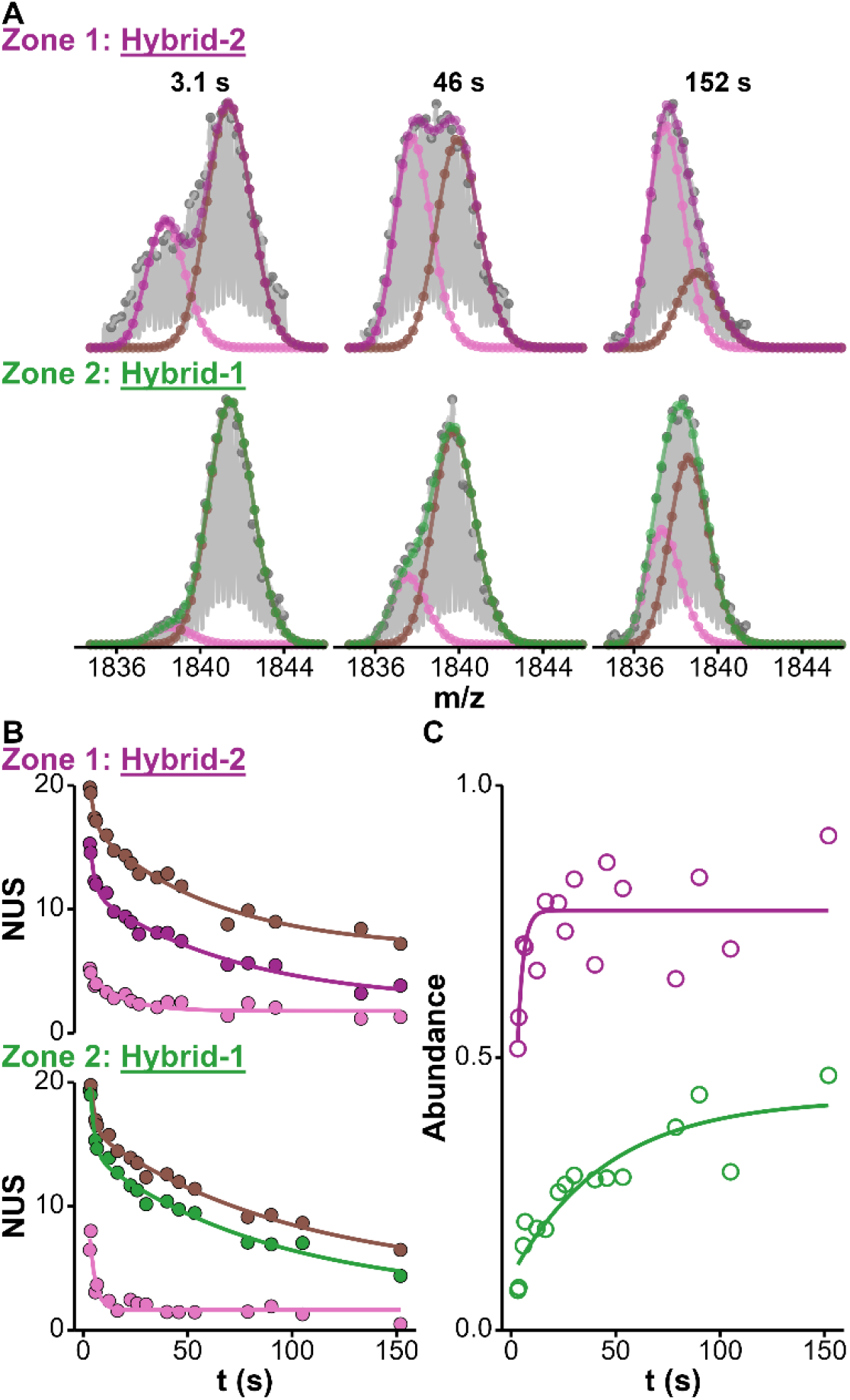
A. Deconvoluted mass spectra at selected exchange timepoints for the two conformers of 23TAG·2K^+^ separated by IMS (Figure 2). High- and low-exchange isotopic populations are fitted in pink and brown, respectively. B. Exchange kinetics for both conformers: the apparent exchange is shown in purple (zone 1) or green (zone 2) and deconvoluted EX2 contributions in pink and brown, as above. C. Abundance of the low- exchanged isotopic populations against the exchange time, highlighting the faster unfolding of the conformer from zone 1 compared to zone 2 (0.57 min^-1^ vs. 1.7c min^-1^, respectively).

In summary, the difference in apparent exchange rates between these two conformers, despite their similar structure and stability, arises not from local fluctuations, but from differences in their unfolding dynamics. This is consistent with a more indirect measurement of their unfolding rates by dynamic fitting of native IMS-MS measurements (*k_op_* ≈ 0.05 min^-1^ for the hybrid-2 vs. 0.73 min^-1^ for the hybrid-1).^56^

When this approach was applied to the closely related sequence 24TTG, which also adopts a mixture of hybrid-1 and hybrid-2 conformers under our experimental conditions,^56^ no difference in apparent exchange was observed (Figure S122). These conformers likely exchange with similar EX2 rates and are too stable to undergo EX1 exchange, rendering them indistinguishable by HDX/IMS-MS. To overcome this limitation, EX1 exchange was induced by reducing the KCl concentration to 0.5 mM, which destabilized the G4 structure sufficiently to promote transient unfolding (Figure S123), without producing a completely unfolded population as observed in 0.1 mM solutions (Figure S124).

#### 3.1.2. Applications and limitations

5YEY is a 21-mer human telomeric motif where a thymine replaces adenine 18 (compared to 21G) (Table 1), whose main conformer adopts a chair-type topology (Figure 1B). The NMR data, which we replicated under our conditions,^48^ suggest the presence of several other conformers,^39^ providing a particularly challenging case study. The results below are similar for 22GT_T18, a variant of 22GT with the same A18-to-T mutation (Figures S108, S109, S151, S152, S177, S184).

Two populations of 5YEY are separated by mass, binding either 1 or 2 K⁺ ions. The minor 5YEY·1K⁺ population is protected from exchange at low exchange times only, consistent with the formation of 2-tetrad G4s, possibly involving additional stacked base pairs or triads, as suggested by their relatively slow exchange compared to typical 2-tetrad G4s.^16^ Two IMS populations, exchanging at distinct rates, were resolved (Figure S102), and both unfold at different rates as indicated by their EX1 exchange profiles (Figure S149 and Figure S150).

The 5YEY·2K⁺ ensemble can be separated into three main IMS populations (Figure 4A). These populations have protection levels reflecting the formation of 3-tetrad G4s, but with significantly different rates (Figure 4B). Like 23TAG, the key difference between these species lies in their unfolding propensity rather than their local dynamics. Hence, population 1 (CCS = 686 Å^2^) exhibits a significant EX1 contribution, indicating partial unfolding (the exchange is incomplete at early timepoints). On the other hand, population 3 (CCS = 762 Å^2^) exchanges exclusively via EX2, indicating that it does not transiently unfold (Figure 4C—E). A limitation of this approach lies in population 2 (CCS = 721 Å^2^), which appears to transiently unfold like population 1, albeit slower. However, because it is not entirely IMS-resolved from populations 1 and 3, it is possible that the data is not perfectly deconvoluted despite our efforts in selecting a pure arrival time distribution zone.

**Figure 4.**
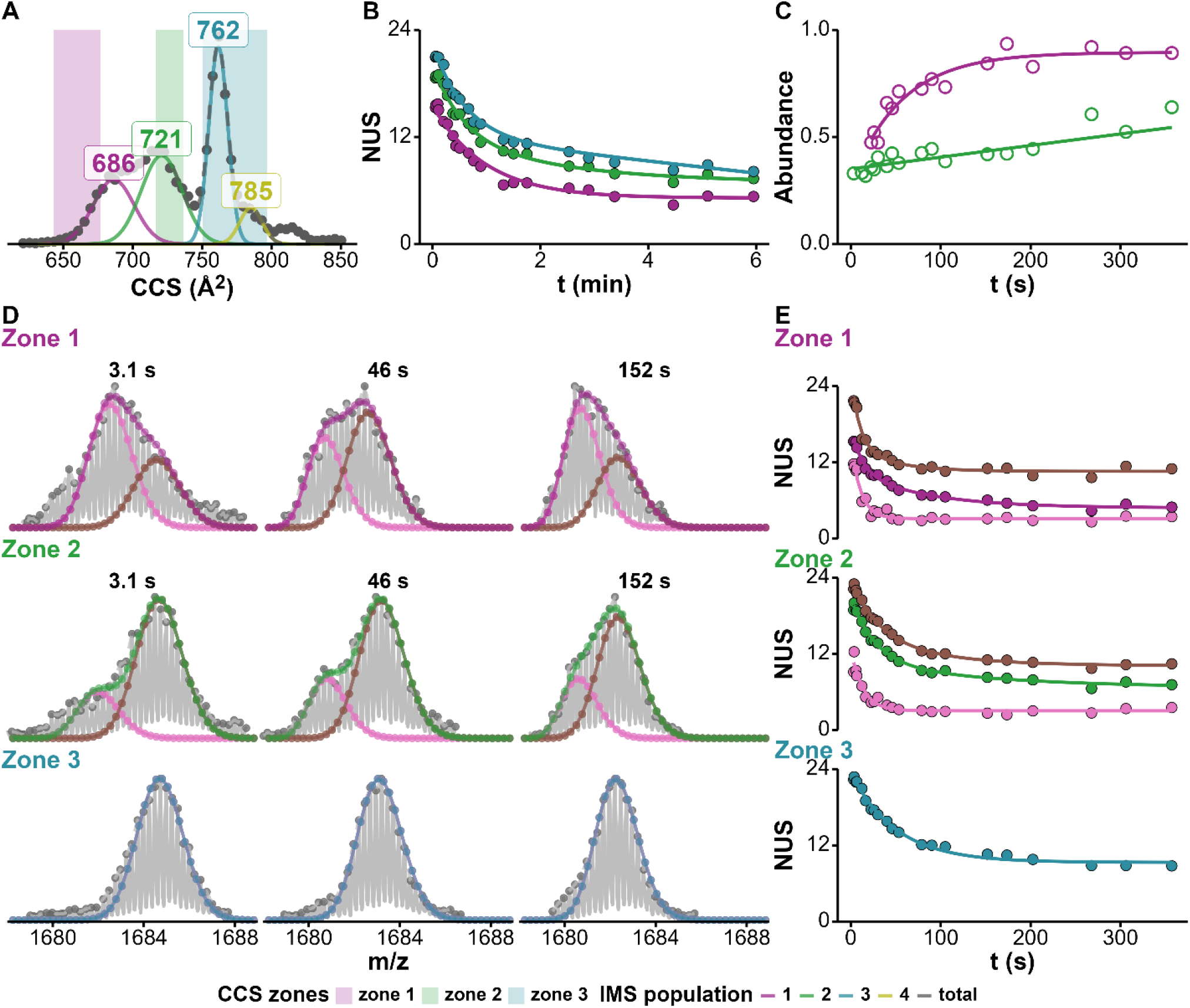
5YEY·2K^+^ has a complex CCS distribution from which three IMS populations are defined (A), each with distinct apparent exchange kinetics (B). Deconvolution of the isotopic populations (D; selected time points) gives access to the pure EX2 (E; high- and low-exchange in pink and brown, respectively) and EX1 (C) exchange kinetics. The 762Å² conformer (blue) exchanges via EX2 exclusively. Both the 686 Å² (purple) and 721 Å² (green) conformers dynamically unfold, with the latter unfolding more slowly.

Lack of IMS peak separation is an issue we encountered with other oligonucleotides from the panel, sometimes to a much larger extent. For example, the wild-type counterparts of 5YEY and 22GT_18T, 21G·2K⁺ (Figure S101) and 22GT·2K^+^ (Figure S109), do not display well-resolved IMS populations, and consequently, no distinct exchange dynamics could be characterized.

### 3.2. Case 2: IMS-filtering of non-specific adducts

We focus here on species of same HDX rate and CCS but different m/z, resulting from the non-specific binding of cations to oligonucleotides. Non-specific binding is defined here as cations binding to the negatively charged phosphate backbone without displacement of their outer-sphere hydration shell, without changes of secondary structure,^18,24^ and therefore with virtually unaltered CCS. This type of electrostatic interactions is prominent in native ESI-MS, and complicates data analysis and interpretation. For instance, the determination of the specific coordination stoichiometry of cations between tetrads by native MS alone can be used as a proxy for estimating the number of G-tetrads, but this can be ambiguous when specific and non-specific adduct distributions are superimposed.^24,62^ We have previously shown that coupling in-solution HDX allows revealing non-ambiguously if different cation adduct stoichiometries are linked to different conformers in solution, as these differences lead to distinct exchange rates.^16^ However, this approach remains insufficient to filter-out non-specific from specific binding and provide cleaned-up HDX data. We therefore investigate here whether IMS can be leveraged to filter-out non-specific cation adduction.

#### 3.2.1. Filtering non-specific adduct for a polymorphic system

To do so, we examined the 222T oligonucleotide (M; Figure 5) in 100-µM K^+^ solutions, where it coexists as 3-tetrad (3T), 2-tetrad (2T) and unfolded (U) conformers.^21^ Accounting for non-specific binding, MK_2_ contains a mixture of 3T specifically binding two K⁺ (3T·2K⁺), 2T binding one K⁺ specifically and one non-specifically (2T·(1+1)K⁺) and U binding non-specifically two K⁺ (U·(0+2)K^+^), all sharing the same m/z.^14,21^ Similarly, MK may contain mixtures of 2T·1K⁺ and U·(0+1)K⁺.

**Figure 5.**
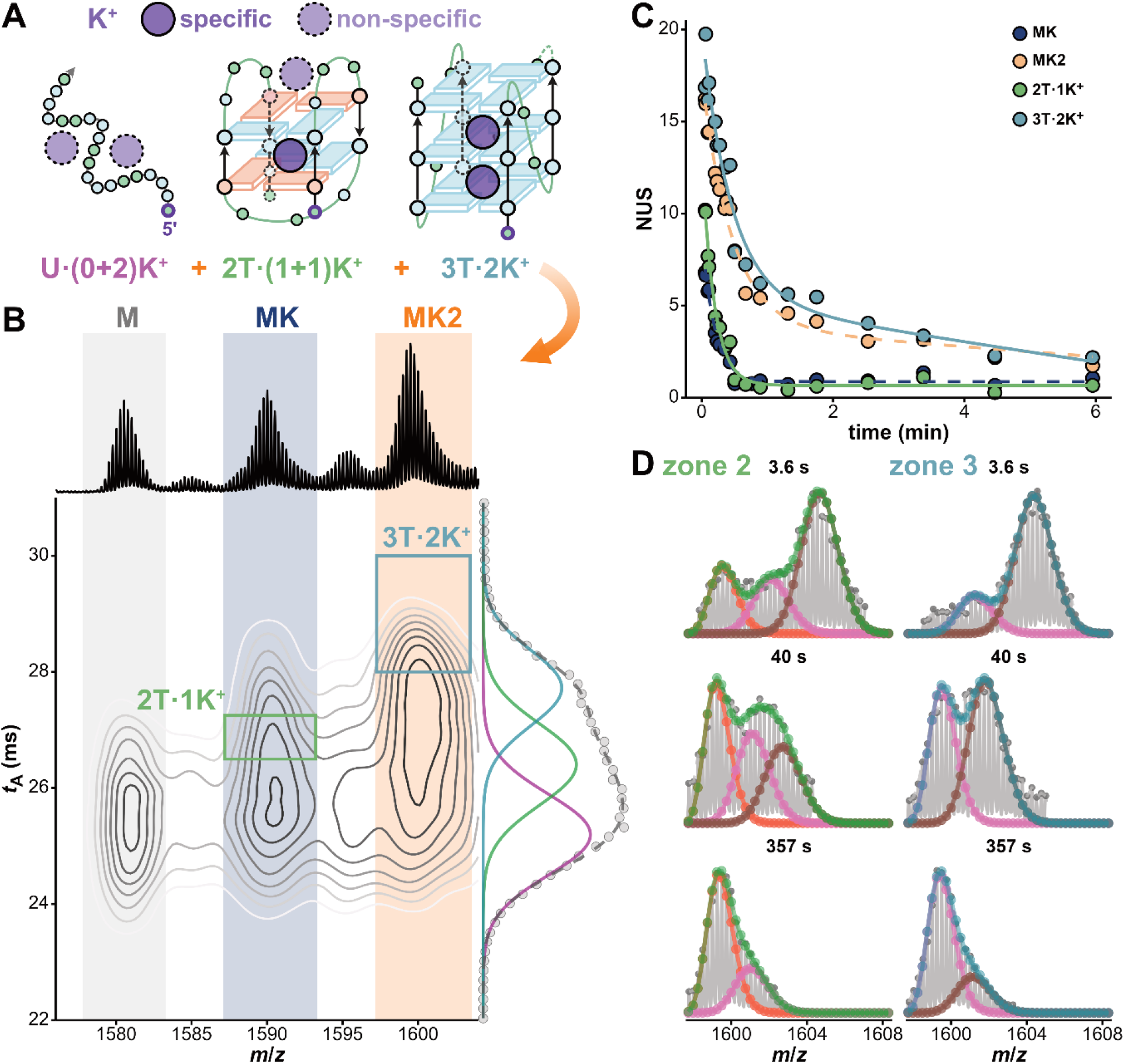
Filtering out non-specific cation binding by IMS: A. The presence of nonspecific K^+^ adducts gives rise to different conformations of same m/z: here 2T binding one specific and one non-specific cation (2T·(1+1)K^+^) and U binding two non-specific cations (U·(0+2)K^+^) have the same m/z has 3T·2K^+^ and are convoluted in MK_2_. The same occurs for MK that contains 2T·1K^+^ and U·(0+1)K^+^. Note that the structure shown here for 2T is not known and likely a mixture of conformers. B. IMS density map of 222T in 0.1 mM KCl (top: native mass spectrum; right: arrival time distribution. C. Extraction of specific IMS-MS regions (rectangles in panel B) provides HDX kinetics of species 3T·2K^+^ and 2T·1K^+^ without the non-specific contributions initially present in MK_2_ and MK, respectively. D. Deconvolution of multimodal isotopic distribution for selected time points of highlight the presence of a contaminant in zone 2 (green) of MK_2_ that is filtered out in zone 3 (light blue).

We processed the HDX/IMS-MS data in a stepwise fashion. First, the ATD of M was fitted with a single broad Gaussian, consistent with an ensemble of unstructured conformations in solution that *i*) undergo significant compaction in the gas phase (CCS = 690 Å²; Figure 5B) and *ii*) exchange within the experimental dead time (Figure 5C). Next, fitting the ATD of MK requires a second broad Gaussian (CCS = 737 Å²), whose resolved 2T·1K⁺ population show marked protection at short exchange times, a characteristic feature of 2-tetrad G4s (Figure S125). Finally, the ATD of MK_2_ includes a third, narrower population at a higher CCS (753 Å²), corresponding to the formation of the well-exchange-protected 3T·2K⁺. Thus, the specific HDX kinetics of 2T·1K⁺ and 3T·2K⁺ are accessed by selection of the intersecting area between their specific m/z and arrival times (green and blue rectangles, respectively; Figure 5B). The “pure” kinetics of 2T·1K⁺ and 3T·2K⁺ are slower than those of their “contaminated” ensembles (MK and MK_2_, respectively), owing to the filtering-out of non-specific contributions from faster-exchanging species (Figure 5C).

IMS filtering of non-specific adducts is also useful to determine exchange mechanisms unambiguously. Both MK and MK_2_ feature bimodal isotopic distributions upon exchange, potentially indicating EX1 behavior, but this could also arise from contaminating species.^16^ The IMS-resolved 3T·2K⁺ population undergoes both EX2 and, unambiguously, slow EX1 exchange, consistent with the high stability of a 3-tetrad G4 (Figure 5D, Figure S163). In contrast, the low-CCS population of MK_2_ exhibits a more complex isotopic pattern comprising three coexisting populations. Deconvolution was carried out with an updated version of the OligoR software that can handle trimodal distributions :^15^ the two more protected populations correspond to the slow, mixed EX1/EX2 exchange of 3T·2K⁺, while the most exchanged population likely originates from a mixture of 2T·(1+1)K⁺ and U·(0+2)K⁺ (Figure 5D, Figure S162).

#### 3.2.2. Applications and limitations

A limitation of IMS-based filtering is its dependence on sufficient separation between conformer populations, a condition that is not always met with inherently broad, gas-phase conformer populations. In previous work, we demonstrated that the mass-resolved ckit87up·2K⁺ and BCL- 2·2K⁺ analytes are both contaminated with low-protection species.^16^ IMS filtering proved only partially effective for ckit87up: while IMS filtering improved the separation, it did not fully eliminate contaminants, resulting in a trimodal isotopic distribution that required deconvolution for accurate interpretation (Figure S169-Figure S171), and access to the EX1 exchange of its 3-tetrad conformation (Figure S128, Figure S187). We advise to systematically evaluate isotopic patterns to detect residual non-specific complexes, which will be reflected by additional isotopic distributions.^16^ IMS-filtering proved ineffective for BCL-2, for which the same apparent exchange and EX1 rates were observed across different IMS zones, indicating that the resolution was insufficient to distinguish between the conformers formed (Figure S54,Figure S129,Figure S186).

The study of conformational mixtures from several human telomeric oligonucleotides (21G, 22AG, 22GT, 22CTA, and 23AG), which have been shown by NMR to form complex conformational mixtures in both ∼100 mM KCl and native MS buffer,^48^ was equally limited. HDX/IMS-MS results confirm the presence of both 2- and 3-tetrad species, as detailed in the Supporting Information (Figure S190). Unfortunately, oligonucleotides containing 12 guanines can theoretically form numerous distinct 2- tetrad conformers of similar stability, resulting in broad IMS distributions that limit further structural resolution.

### 3.3. Case 3: Distinguishing if multiple IMS peaks come from one or several solution ensembles, as defined from HDX

Intuitively, we may think that two well defined IMS peaks always come from two well defined solution ensembles, as leveraged in case 1. Here we show that, sometimes, two IMS peaks can come from the same solution ensemble, as defined by their HDX exchange profile.

Several well-characterized, monomorphic G4-forming oligonucleotides display multiple IMS peaks but identical HDX rates for IMS-resolved populations. This is the case of *i*) T95TT and T30177TT, both forming 3-tetrad parallel G4s with single thymines in the loops, as shown by in-solution NMR and under native MS conditions (Figure 6, Figure S133, Figure S134),^36,41,48^ *ii*) 2KPR and 26CEB, two G4s featuring longer loops with increasing dynamics, (Figure S130-Figure S132),^40,42,48^ and *iii*) [TG4T]4, which form a loop-less tetramolecular G4 (similar observations for both 4– and, to a lesser extent, 5– charge states and in K^+^ and NH4^+^ solutions; Figure S135-Figure S137).

**Figure 6.**
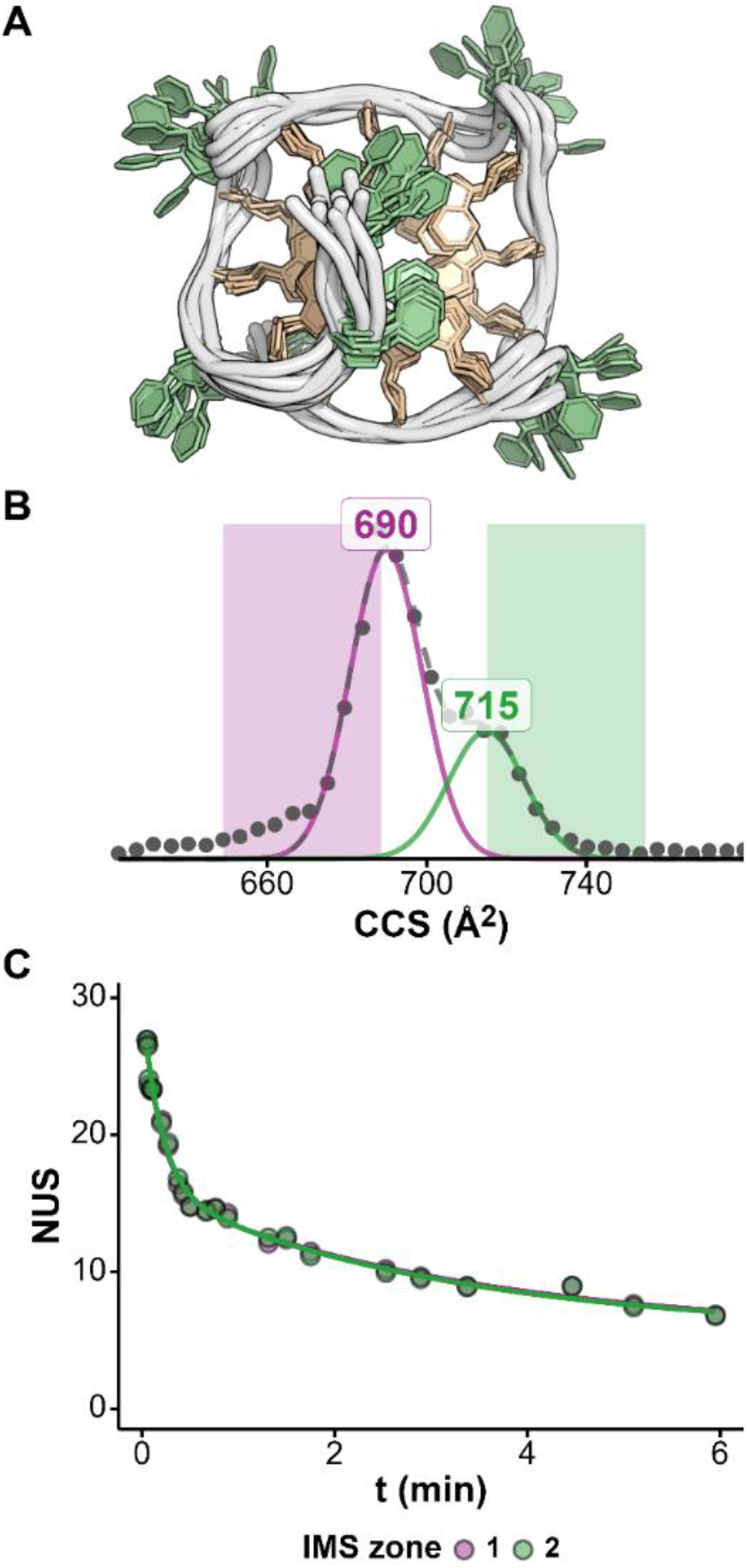
Detection of gas-phase rearrangements of T30177TT by HDX/IM-MS. A. top view from the 5’ end of the 10 lowest-energy conformers deposited in the PDB (2M4P).^3c^ B. The ATD of [T30177TT·2K^+^]^4-^ in 100 mM TMAA + 0.5 mM KCl presents a bimodal distribution indicating the presence of at least two conformational ensembles in the gas-phase. C. These two ensembles exchange at the same rate, suggesting that they are produced from a single structure in solution.

One explanation for these observations is that homogeneous populations in solutions may produce heterogeneous gas-phase ion populations due to different electrospray droplet evolution, including differences in droplet ionic strength, size and fission pathways, and charge-residue (CRM) versus chain-ejection-like (CEM) release mechanisms.^23,63,64^ Alternatively, these observations may also be ascribed to the presence of structural microheterogeneities in solution, specifically fast dynamics of loop and/or flanking nucleotides. These structural features do not affect apparent HDX rates since *i*) unpaired, solvent-exposed residues do not contribute significantly to exchange and *ii*) any potential influence on the exchange rate (transient pairing, stacking) would be too fast (< 1 s) to be sampled by our HDX/MS setup.^14^ However, different conformations of these structural features may lead to distinct post-ESI gas-phase rearrangements, producing multimodal ATDs. This is consistent with the increase of the phenomenon at higher fragmentor voltage, which we previously reported for 2KPR.^48^

On the one hand, coupling IMS to HDX may therefore be useful to suggest the presence of fast dynamics that are invisible to HDX/alone. On the other hand, adding HDX to native IMS/MS allows determining whether IMS populations arise from different conformers in solution, or are only the result of gas-phase rearrangements.^65^ Finally, it is important to note that identical HDX rates for two IMS populations do not necessarily equate to a single solution-phase structure. Two distinct conformers can exchange at similar rates, as illustrated by the hybrid-1 and hybrid-2 G4s of 24TTG in section 3.1.1 (Figure S122). However, this is dependent on solution conditions and differences were revealed at lower [KCl] (Figure S124).

## 4. Conclusions

In summary, coupling IMS with native HDX/MS significantly enhances our ability to probe the structural dynamics of DNA oligonucleotides. Our approach resolves three key scenarios:

1. IMS-HDX/MS overcomes a major limitation of native HDX/MS alone, by allowing specific HDX rate measurement for coexisting conformers of identical *m/z* but different CCS. We exemplified this capability on complex conformational mixtures formed by 23TAG, in agreement with a previous report,^56^ and 5YEY. However, this approach remains limited to sufficiently-well IMS resolved conformers. More generally, IMS-enabled native HDX/MS should prove useful to study the structural dynamics of oligonucleotides whose folding does not necessarily triggers an *m/z* shift, which is often the case for non-G4 species.
2. A common occurrence in native ESI-MS of nucleic acids is the non-specific binding of cations to analytes, which complicates data analysis and interpretation. Non-specific binding translates into several species of different *m/z* but sharing the same CCS and HDX rates. When several stoichiometries of specific and non-specific adducts coexist, mass- separation alone becomes insufficient to resolve different conformers. Combining IMS is effective to filter non-specific complexes out, yielding more accurate HDX rates and mechanisms. This approach relies on the ability to effectively separate the conformers by IMS, which is not always possible. Careful analysis of HDX isotopic patterns provides an internal control for evaluating the effectiveness of IMS filtering, since non-specific complexes generate additional isotopic populations distinct from those produced by EX1 exchange ^16^.
3. Several species detected with distinct CCS despite having the same *m/z* and HDX rates indicate distinct gas-phase conformational ensembles issued from the same solution conformational ensemble. In some cases, distinct conformers may exchange at similar rates, but this is solution-condition dependent. HDX is therefore a precious aid in the interpretation of multimodal collision cross distributions.

Together, these findings establish HDX/IMS-MS as a powerful tool for studying the structural polymorphism and folding of structured nucleic acids. Because the method can now be applied to mixtures of analytes of same mass, we anticipate that it will also be applicable to other classes of nucleic acid secondary structures beyond G4s.

## Supporting Information

The Supporting Information file contains:

- Additional schematics (Figure S1-Figure S4, Figure S99)
- Additional methods and results for circular dichroism (Figure S5, Figure S6), mass spectrometry parameter optimization (Figure S7-Figure S17), CCS conversion (Figure S18- Figure S54), collision-induced unfolding (Figure S55-Figure S98), HDX/IMS (Figure S100- Figure S137), isotopic distribution deconvolution and corresponding EX2 and EX1 exchange kinetics (Figure S138-Figure S173), cation adduction specificity (Figure S174-Figure S190)

## Acknowledgements

We thank Dr. Frédéric Rosu and Dr. Corinne Buré at the Institut Européen de Chimie et Biologie (IECB/CNRS UMR3033) for access to the mass spectrometry platform and their invaluable help.

