## Supporting Information for "Native Hydrogen/Deuterium Exchange Ion Mobility Mass Spectrometry of Structured DNA Oligonucleotides"

### Ion Mobility schematics

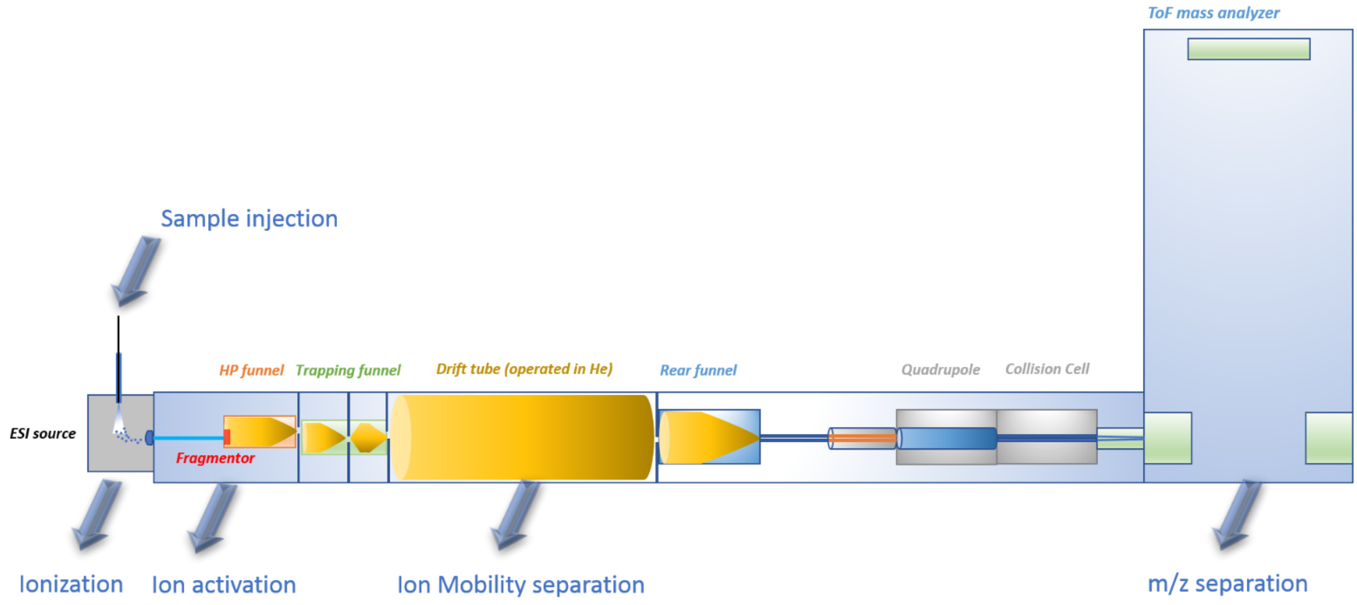


*Figure S1. Scheme of the Agilent ESI-IMS-QTOF 6560 with the position of the fragmentor in red, located at the entrance of the high-pressure funnel after the resistive-glass capillary. The drift tube is filled with helium.*

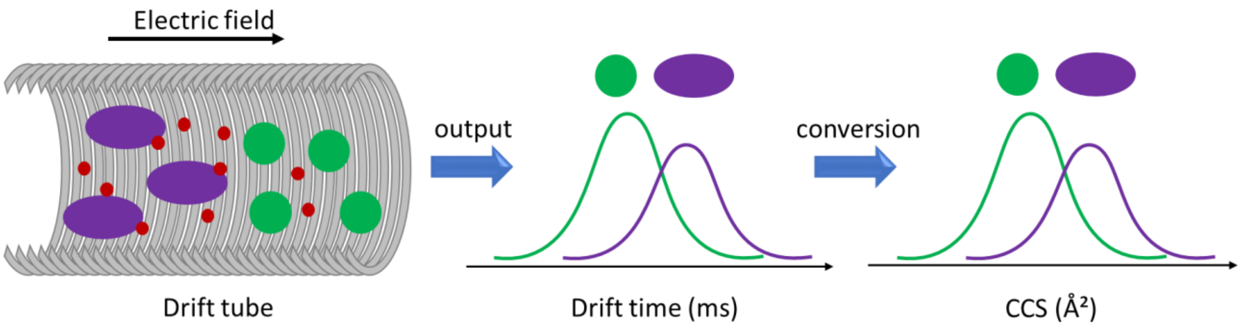


*Figure S2. Simplified IMS drift tube mechanism. In the drift time for ions of same charges, the more compact (in orange) undergo fewer collisions with the drift gas (in red) compared to more elongated structures (in blue) which leads to a faster passage in the drift tube. The output is a distribution of drift times that is multimodal when species of different shapes travel through the tube. Finally the arrival time distribution can be converted into CCS using intrinsic parameters (mass, charge, pressure, temperature)*

### G4 structures

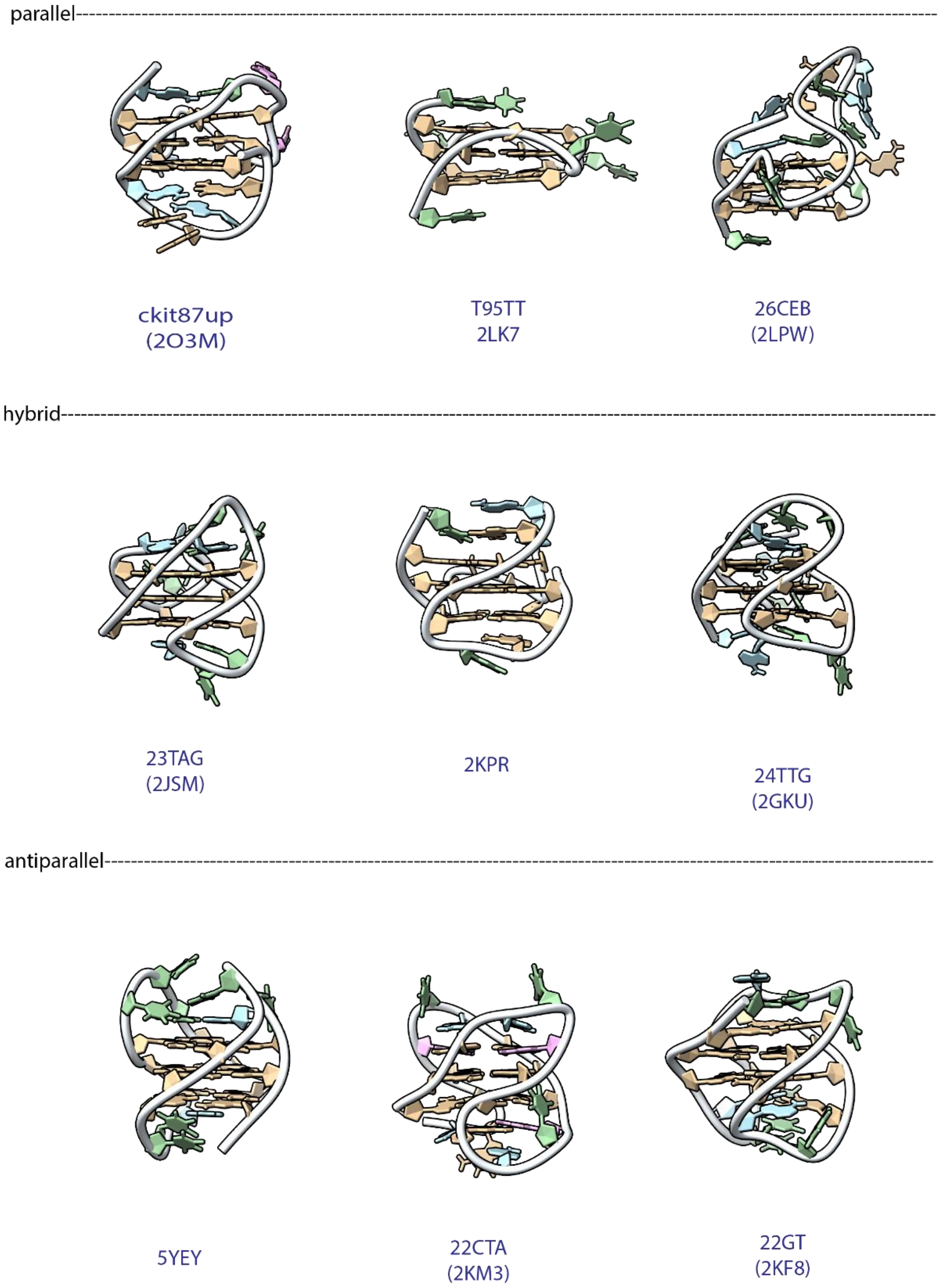


*Figure S3. PDB-deposited structures of different G4 structures used in this assay grouped by topology in 1 mM KCl. The phosphate backbone is represented by a white line, the guanines are colored in tan, thymines in green, adenines in blue and cytosines in pink.*

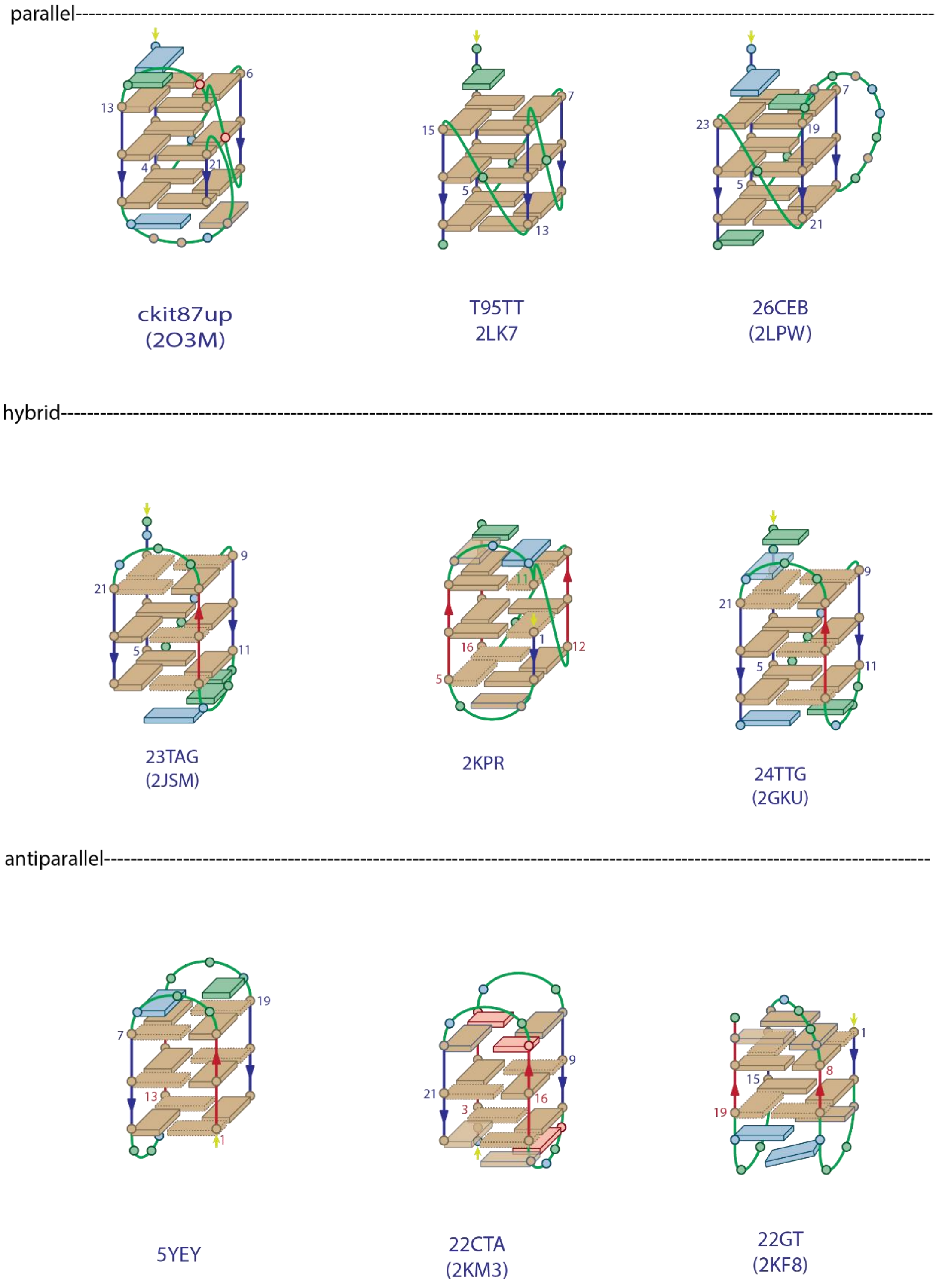


*Figure S4. Schematic version of different G4 structures used in this assay grouped by topology in 1 mM KCl. The guanines are represented by tan rectangles with straight (resp. dotted) lines for the anti (resp. syn) conformation, thymines in green, adenines in blue and cytosines in pink. The strand direction is indicated by blue and red lines. 5YEY is classified as an antiparallel G4 in respect of the strand orientation however the G-tetrads bases stacking follows a hybrid pattern which makes the topology assignment of this sequence ambiguous.*

### CD experiments

Circular Dichroism spectroscopy is a frequently used spectroscopic technique to get insights of the topology adopted by G4 structures.[1–4] The CD spectra of the oligonucleotides were recorded with samples at 10 µM DNA in the same buffer as the HDX/IMS experiment on a JASCO J-1500 spectrophotometer in 1 cm pathlength quartz cuvettes. Blank samples with adequate buffer and %D were also measured. The wavelength scanning was set from 220 to 335 nm at 50 nm/min with three accumulations. The CD signal obtained was blank-subtracted, baseline-corrected and the ellipticity obtained in mdeg *(θ)* was converted to molar absorptivity *(Δε)*, accounting for the pathlength *(l)* and the concentration of the analyte *(c)* using Equation 3.

$$\Delta\varepsilon = \frac{\theta}{32980 \times l \times c} \quad (3)$$

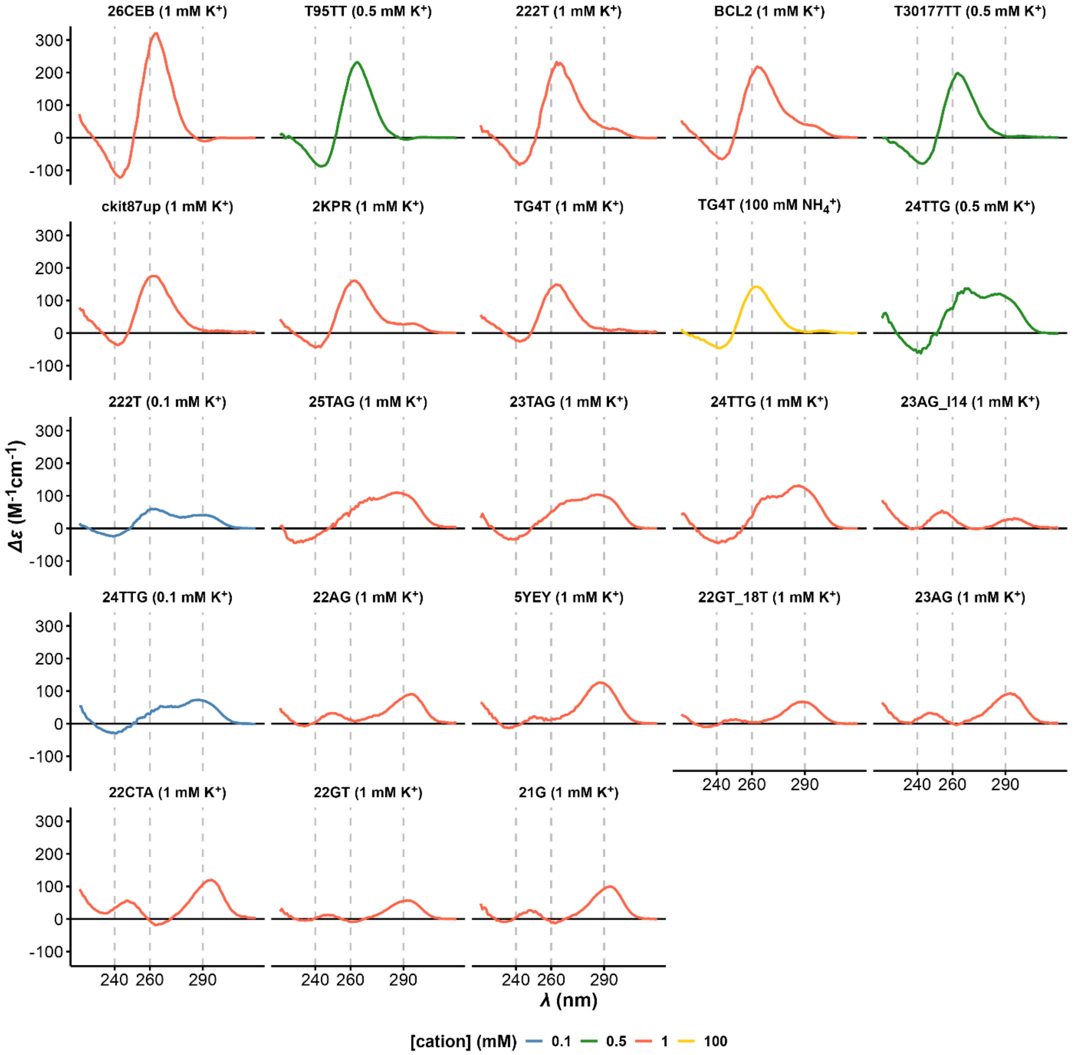


*Figure S5. CD spectra of the oligonucleotides in the panel at 10 µM G4 ranked by decreasing CD value at 260 nm. A sole positive band at 260nm indicates a parallel topology, Positive bands at 250 nm and 290 nm and a negative band at 260 nm are characteristic of antiparallel topologies. Two consecutive positive peaks in the 260-290 range is indicative of a hybrid topology. The curves color corresponds to the buffer.*

### CD-melting experiments

Melting curves of 22GT_18T, 23AG and 23AG_I14 were obtained from CD-melting analysis on the same JASCO instrument used for CD spectra. Temperature range: 4-90°C, 1 spectrum/°C, 1°C/min range. Scanning range: 220-350nm, 100 nm/min in quartz cuvettes, 1 cm pathlength.

The wavelength selected for $T_m$ determination corresponds to the maximum of CD absorbance (specified on the melting curves for each oligonucleotide).

Blank subtraction, baselines correction and conversion from CD units to folded fraction was in the same way as in previous works,[5,6] according to standard protocols in literature.[7,8]

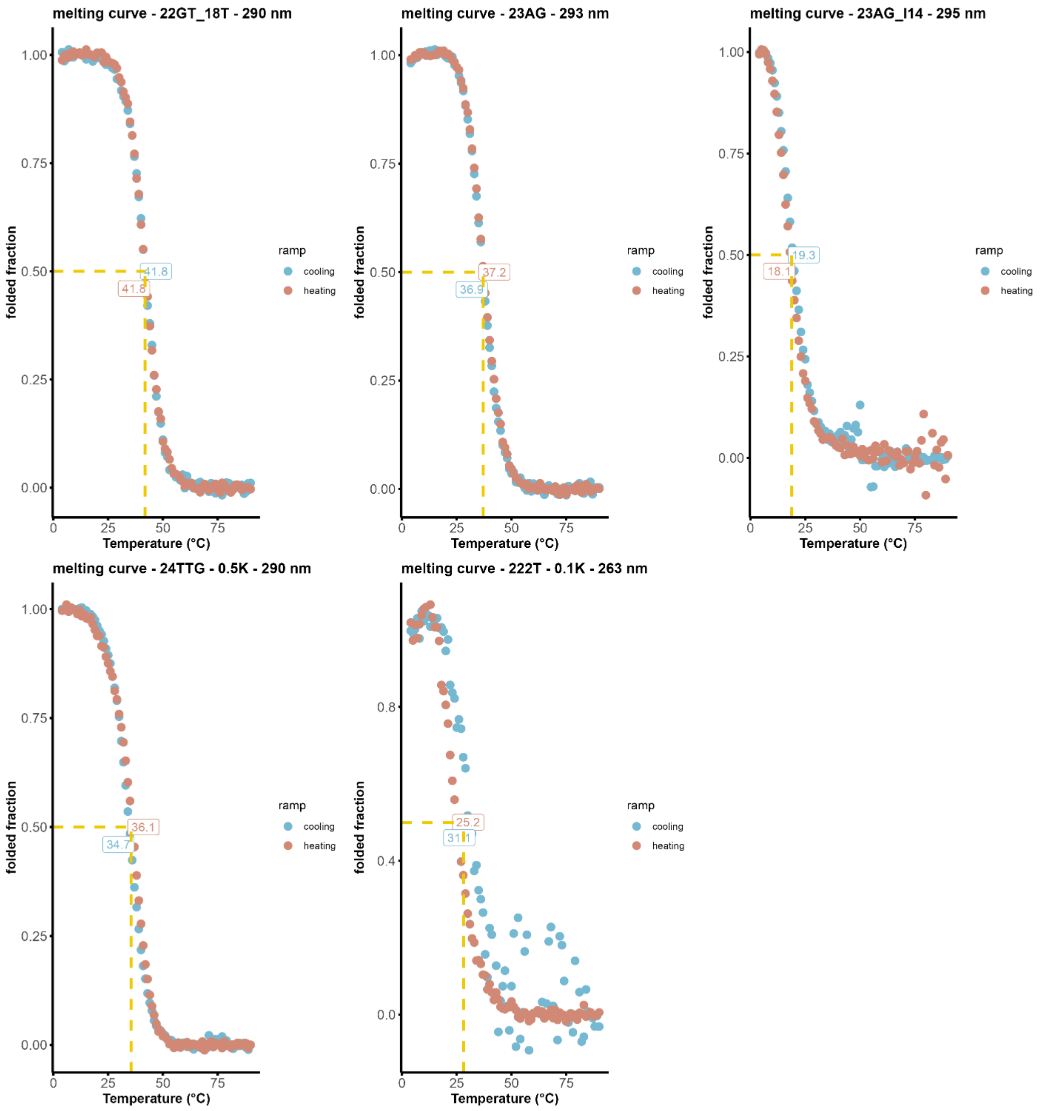


*Figure S6. Melting curve of the oligonucleotides 22GT_18T, 23AG and 23AG_I14 in 100 mM TMAA+1mMKCl, melting curve of 24TTG+0.5 mM KCl and melting curve of 222T in 0.1 mM KCl.*

### Instrument parameter influence

The HDX/MS measurements accuracy and precision have already been determined for the study of several structured and unstructured oligonucleotides.[9] ESI source atmosphere can affect the deuteration of oligonucleotides, in particular in too harsh conditions or in presence of $D_2O$ vapors. Potential gas-phase or in-droplet exchange must therefore be evaluated for coupling of IMS to native HDX/MS, where the ions spend more time in the gas phase. Additionally, instrumental settings must also be optimized to work in native conditions while ensuring sufficient signal-to-noise ratio (S/N) and resolution of conformer populations by IMS. We therefore evaluated the impact of the source nebulizer pressure and drying gas flow (impacting the spray formation and desolvation process), the fragmentor voltage (influencing the softness and efficacy of desolvation), and drift voltage, which needs to be varied for CCS measurements (Figure S1).

To avoid any experimental bias, the experiments were performed with both increasing and decreasing parameter values, interspersed by an evacuation of the source, and at two exchange times (4.2 and 53.4 s). We used two intramolecular G4s, 23TAG·$2K^+$ (lower stability, polymorphic, in 1 mM KCl) and T30177TT·$2K^+$ (higher stability, monomorphic) and a tetramolecular G4 $[TG4T]_4$·$3NH_4^+$ (in 100 ammonium acetate) (Figure S7-Figure S17).

### Methods

The impact of key instrument parameters on the deuteration and signal-to-noise ratio was assessed by systematic variation of their values: nebulizer pressure (4—48 psi) dry gas flow (1—5 L/min), fragmentor voltage (300—440 V), and drift voltage (650—1000 V). Duplicated, 1-min acquisitions were performed for 4.19 s and 53.42 s mixing times, in both increasing and decreasing values. The source was evacuated for 2.0 min between each timepoint and change of parameter.

The impact of the instrumental parameters on deuteration was quantified for all populated charge states as the relative variation of exchange $\frac{\Delta NUS}{NUS}$ using Equation 4, where $NUS^0$ is the $NUS$ value at the default parameter value.

$$\frac{\Delta NUS}{NUS} = \frac{NUS\text{-}NUS^0}{NUS} \quad (4)$$

The signal to noise ratio (S/N) was defined for 0.9 min acquisitions by Equation 5, where $I_{peak}$ is the intensity of the peak of interest divided and $\sigma_{base}$ the standard deviation of the baseline intensity. The baseline is defined as an 80 *m/z*-wide region devoid of signal, positioned immediately before or after the peak of interest.

$$\frac{S}{N} = \frac{I_{peak}}{3\sigma_{base}} \quad (5)$$

### Results

#### *Influence of IMS source parameters on the H/D exchange*

The in-solution exchange is performed in the D-to-H direction, therefore additional exchange in droplets or in the gas-phase with water vapor should increase the apparent exchange, yielding lower NUS values. Overall, there is no significant impact on deuteration when varying the source nebulizer pressure, drying gas flow, and drift tube voltage (Table S1, Figure S11). For 23TAG, larger drying gas flow rates cause a small but significant increase of the H uptake (4%; $p \approx 0.02$). In the HDX/IMS-MS experiments below, we kept the values of drying gas flow (2 L/min), nebulizer pressure (12 psi), and drift tube voltage (650 V) constant.

The fragmentor is the only parameter to significantly trigger gas-phase exchange (Figure S7). The NUS values significantly drop at 400 V (~ 10%) and 440 V (~ 20%) for both 23TAG and T30177TT. At these fragmentor values, the analytes undergo CIU (Figure S74, Figure S93).[10,11] Their activation combined with an incomplete desolvation or the presence of water vapors in this part of the HP funnel may lead to gas-phase exchange. The decrease in NUS values is lower (< 5% at 440 V) for the tetramolecular G4 $[TG_4T]_4$, which does not disrupt at these fragmentor voltages (Figure S96-Figure S98), and thus maintains an efficient protection against exchange. The value at which gas-phase exchange becomes significant is therefore oligonucleotide dependent and likely linked to the voltage of CIU onset. To assess the maximum voltage that can be used without affecting the apparent exchange for any given species of the panel oligonucleotides, we systematically performed CIU experiments (Figure S55-Figure S98). Hereafter, the fragmentor voltage was set to 320 V, and increased to improve the S/N and IMS resolution, only where necessary (see section 0).

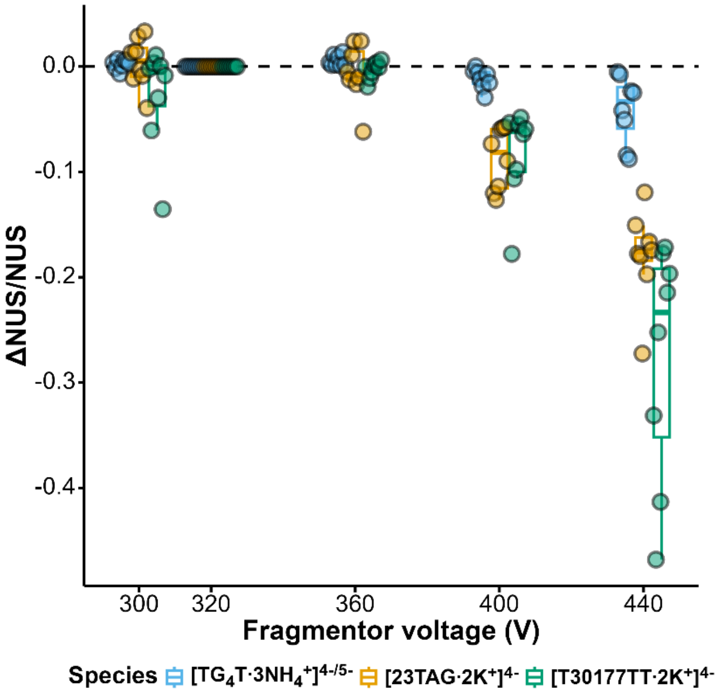


*Figure S7. Variation of the NUS values against the fragmentor voltage, relative to reference values at 320V.*

#### *Influence of IMS source parameters on MS signal and ATD*

The S/N is analyte-dependent and usually worse for non-parallel/polymorphic oligonucleotides (Figure S12), raising challenges for their HDX/IMS-MS analysis.

Increasing the drift voltage has no effect on the S/N ratio (Figure S12), but narrows the ATD peaks (Figure S13-Figure S16), which could be explained by a smaller number of collisions with the drift gas.[12] Increasing the drying gas flow results in a gradual improvement of the S/N up to a factor of two between 1 and 5 L/min (Figure S12). This improvement is also accompanied by a better resolution of the ATD populations, allowing to distinguish conformers of 23TAG more confidently (Figure S13). Similarly, an increase of nebulizer pressure improves the S/N up to a factor 2.5 from 4 to 24 psi (Figure S12). However, further increase causes the signal to drop. The ATD remains poorly resolved in the case of 23TAG, but a small shouldering appears in the case of T30177TT (Figure S14), possibly due to the better S/N ratio.

Increasing the fragmentor voltage significantly improves the S/N (Figure S12), and in some cases the ATD population resolution as well. This is the case for 23TAG, where a second population is more clearly visible at 360 V than at 300 V (Figure S13). However, too high fragmentor voltages lead to a compaction of the ions in the gas phase (Figure S188A),[10,11,13] and can trigger undesired gas-phase exchange (see section 0). If the S/N and/or IMS-resolution is not satisfactory at low fragmentor voltage, a compromise must be found. Monomorphic, parallel G4s such as T30177TT have narrow ATDs, with clear CIU onset. Conversely non-parallel G4s such as 23TAG generally have inherently broad ATDs, making the onset voltage of compaction less clear.

### Mass Spectra

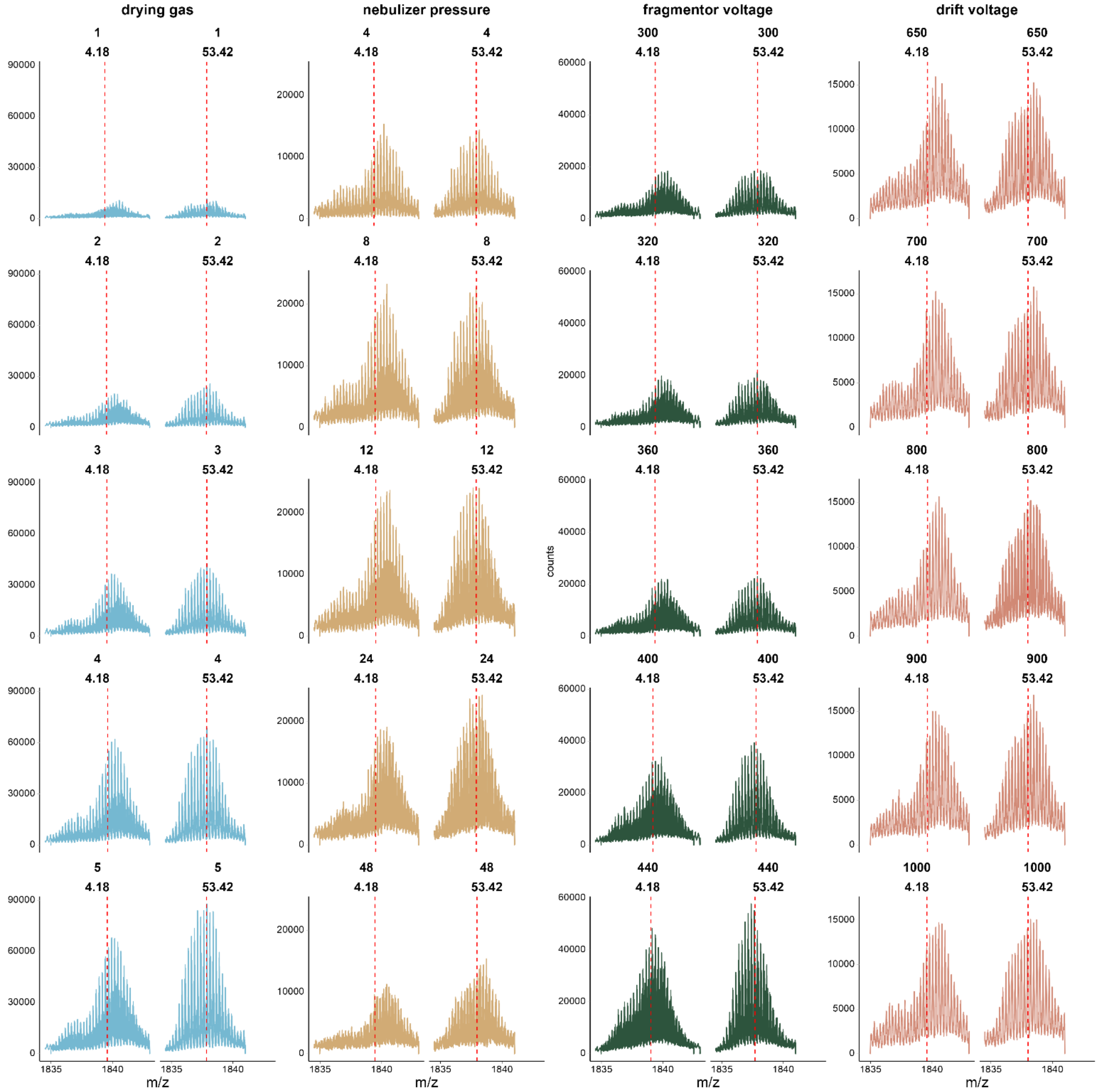


*Figure S8. MS distributions of 23TAG·2K⁺ for each parameter value (indicated in rows) and at 4.18 and 53.42s of mixing time (sub columns). The red dotted lines represent the centroid masses in each case.*

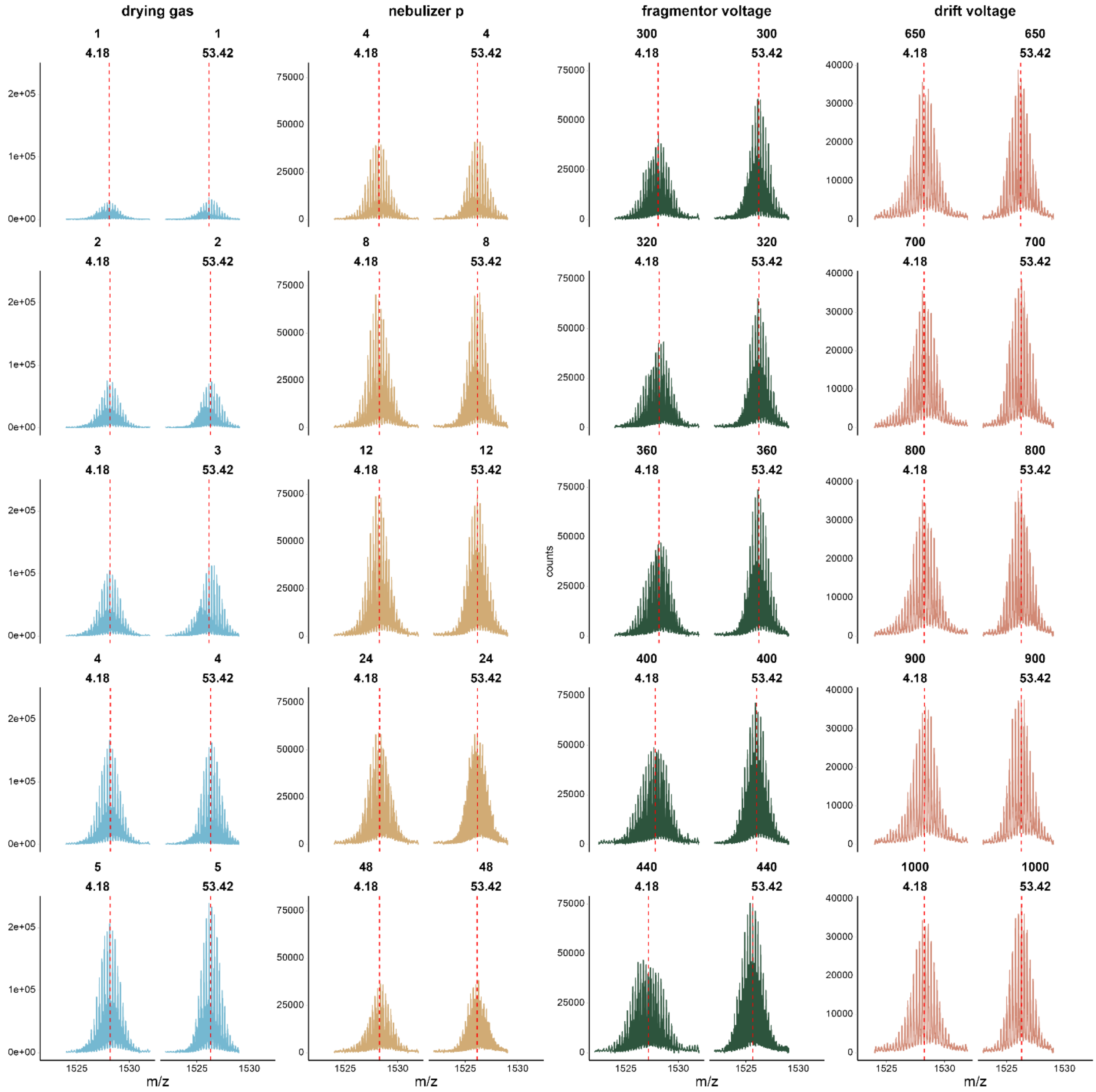


*Figure S9. MS distributions of T30177TT·2K⁺ for each parameter value (indicated in rows) and at 4.18 and 53.42s of mixing time (sub columns). The red dotted lines represent the centroid masses in each case.*

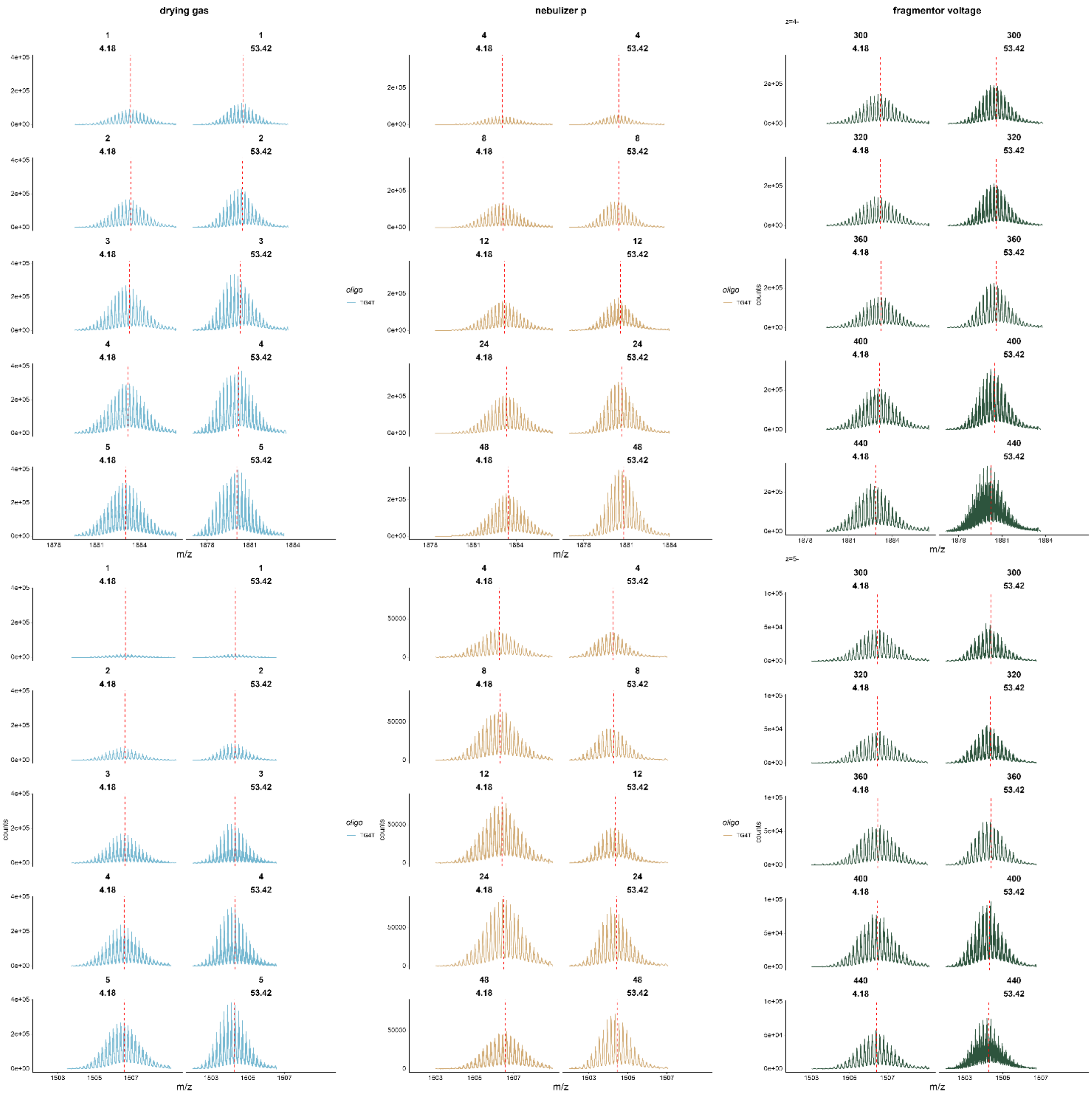


*Figure S10. MS distributions of the TG4T·3$NH_4^+$ for each parameter value (indicated in rows) and at 4.18 and 53.42s of mixing time (sub columns). The red dotted lines represent the centroid masses in each case. The top panel accounts for the 4- charge state and the bottom one for the 5- charge state.*

### *t-tests on source parameters variations*

*Table S1 Average data of the parameter section with the p-value associated to the following null hypothesis: H0: "the population mean is equal to the default value at a confidence level of 95%". Values at which the H0 is rejected are indicated in bold and the default parameter values are indicated with an asterisk.*

| Oligo | Parameter | Value | ΔNUS/NUS average (%) | p-value (NUS) |
|---|---|---|---|---|
| T30177TT | Fragmentor voltage (V) | 300 | -2.7672 | 0.1559 |
| | | 320* | 0 | n.c |
| | | 360 | -0.301 | 0.3483 |
| | | 400 | -8.279 | **0.0011** |
| | | 440 | -27.8127 | **0.0002** |
| | Drying gas flow rate (L/min) | 1 | -3.7811 | 0.0817 |
| | | 2* | 0 | n.c |
| | | 3 | -2.6273 | 0.4220 |
| | | 4 | 0.1689 | 0.8628 |
| | | 5 | 0.9261 | 0.0682 |
| | Nebulizer pressure (psi) | 4 | -2.3248 | 0.2521 |
| | | 8 | -0.0903 | 0.8659 |
| | | 12* | 0 | n.c |
| | | 24 | 0.4613 | 0.1363 |
| | | 48 | -0.3875 | 0.5419 |
| | Drift voltage (V) | 650* | 0 | n.c |
| | | 700 | 0.7133 | 0.3545 |
| | | 800 | 1.1848 | 0.2397 |
| | | 900 | 1.3576 | 0.2821 |
| | | 1000 | 1.6505 | 0.1974 |
| 23TAG | Fragmentor voltage (V) | 300 | 0.3371 | 0.7009 |
| | | 320* | 0 | n.c |
| | | 360 | -0.5904 | 0.5659 |
| | | 400 | **-8.7349** | **6.5756 E-5** |
| | | 440 | **-17.9704** | **8.2967 E-6** |
| | Drying gas flow rate (L/min) | 1 | -3.1261 | 0.1812 |
| | | 2* | 0 | n.c |
| | | 3 | 3.1228 | 0.1070 |
| | | 4 | **4.3396** | **0.0177** |
| | | 5 | **4.4216** | **0.0272** |
| | Nebulizer pressure (psi) | 4 | -2.8419 | 0.1816 |
| | | 8 | -0.0276 | 0.9744 |
| | | 12* | 0 | n.c |
| | | 24 | -1.1661 | 0.3660 |
| | | 48 | -1.2511 | 0.5944 |
| | Drift voltage (V) | 650* | 0 | n.c |
| | | 700 | -0.4580 | 0.6622 |
| | | 800 | 0.0293 | 0.9478 |
| | | 900 | 0.3393 | 0.5912 |
| | | 1000 | -0.5856 | 0.7106 |
| TG4T (z grouped) | Fragmentor voltage (V) | 300 | 0.2311 | 0.2039 |
| | | 320* | 0 | n.c |
| | | 360 | **0.5504** | **0.0185** |
| | | 400 | **-1.1573** | **0.0107** |

| | | | | |
|---|---|---|---|---|
| | | *440* | ***-4.0560*** | ***0.0087*** |
| | *Fragmentor voltage (V)* | *1* | *-0.1434* | *0.6755* |
| | | *2** | *0* | *n.c* |
| | | *3* | *-1.1633* | *0.1307* |
| | | *4* | *-3.1308* | *0.0742* |
| | | *5* | ***-4.5908*** | ***0.0459*** |
| | *Nebulizer pressure (psi)* | *4* | ***-2.6374*** | ***0.0021*** |
| | | *8* | ***-2.1095*** | ***0.0065*** |
| | | *12** | *0* | *n.c* |
| | | *24* | ***2.1611*** | ***4.4824 E-6*** |
| | | *48* | ***3.5675*** | ***1.9312 E-5*** |
| *TG4T (z 4-)* | *Fragmentor voltage (V)* | *300* | *0.1861* | *0.6031* |
| | | *320** | *0* | *n.c* |
| | | *360* | *0.5848* | *0.1079* |
| | | *400* | ***-1.6420*** | ***0.0483*** |
| | | *440* | ***-6.59775*** | ***0.0109*** |
| | *Drying gas flow rate (L/min)* | *1* | *-0.0568* | *0.9353* |
| | | *2** | *0* | *n.c* |
| | | *3* | *-2.2705* | *0.1435* |
| | | *4* | *-5.505* | *0.1215* |
| | | *5* | ***-8.3238*** | ***0.055*** |
| | *Nebulizer pressure (psi)* | *4* | ***-2.7263*** | ***0.0432*** |
| | | *8* | ***-2.4295*** | ***0.05523*** |
| | | *12** | *0* | *n.c* |
| | | *24* | ***2.4657*** | ***0.0013*** |
| | | *48* | ***4.3797*** | ***0.0011*** |
| *TG4T (z 5-)* | *Fragmentor voltage (V)* | *300* | *0.2760* | *0.1617* |
| | | *320** | *0* | *n.c* |
| | | *360* | ***0.516*** | *0.1737* |
| | | *400* | ***-0.6726*** | *0.1347* |
| | | *440* | ***-1.5143*** | ***0.0613*** |
| | *Drying gas flow rate (L/min)* | *1* | *-0.2300* | *0.4862* |
| | | *2** | *0* | *n.c* |
| | | *3* | *-0.056* | *0.6021* |
| | | *4* | *-0.7565* | ***0.0329*** |
| | | *5* | ***-0.8580*** | ***0.0432*** |
| | *Nebulizer pressure (psi)* | *4* | ***-2.5485*** | ***0.0634*** |
| | | *8* | ***-1.7895*** | ***0.1255*** |
| | | *12** | *0* | *n.c* |
| | | *24* | ***1.8565*** | ***0.0017*** |
| | | *48* | ***2.7552*** | ***0.0002*** |

#### *Influence of instrumental parameters of the apparent exchange*

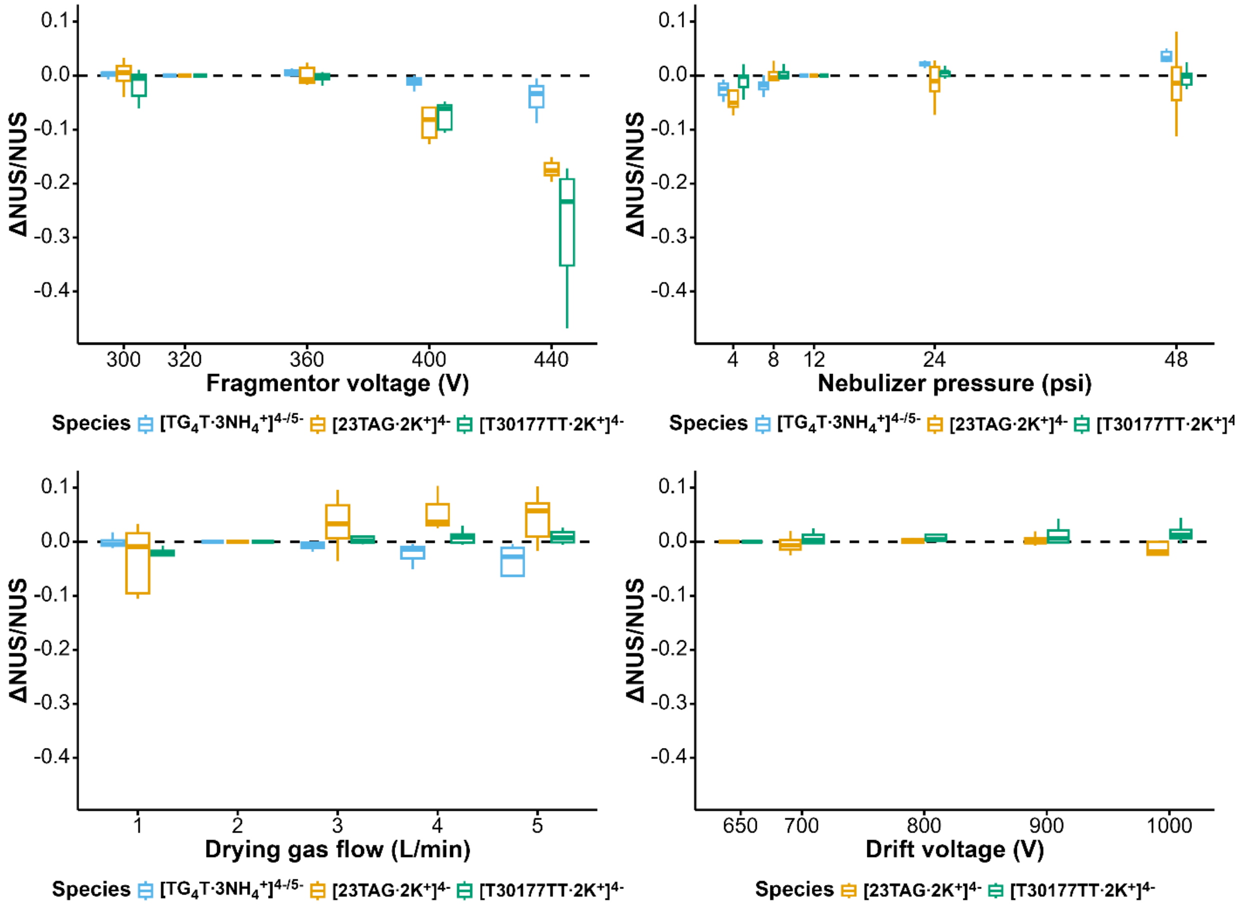


*Figure S11. Variation of the NUS values against the fragmentor voltage, nebulizer pressure, drying gas flow, and drift voltage, relative to reference values at the default setting; fragmentor voltage: 320 V, nebulizer pressure: 12 psi, drying gas flow: 2 L/min, , and drift tube voltage (650 V).*

### Influence of instrumental parameters of the signal-to-noise ratio

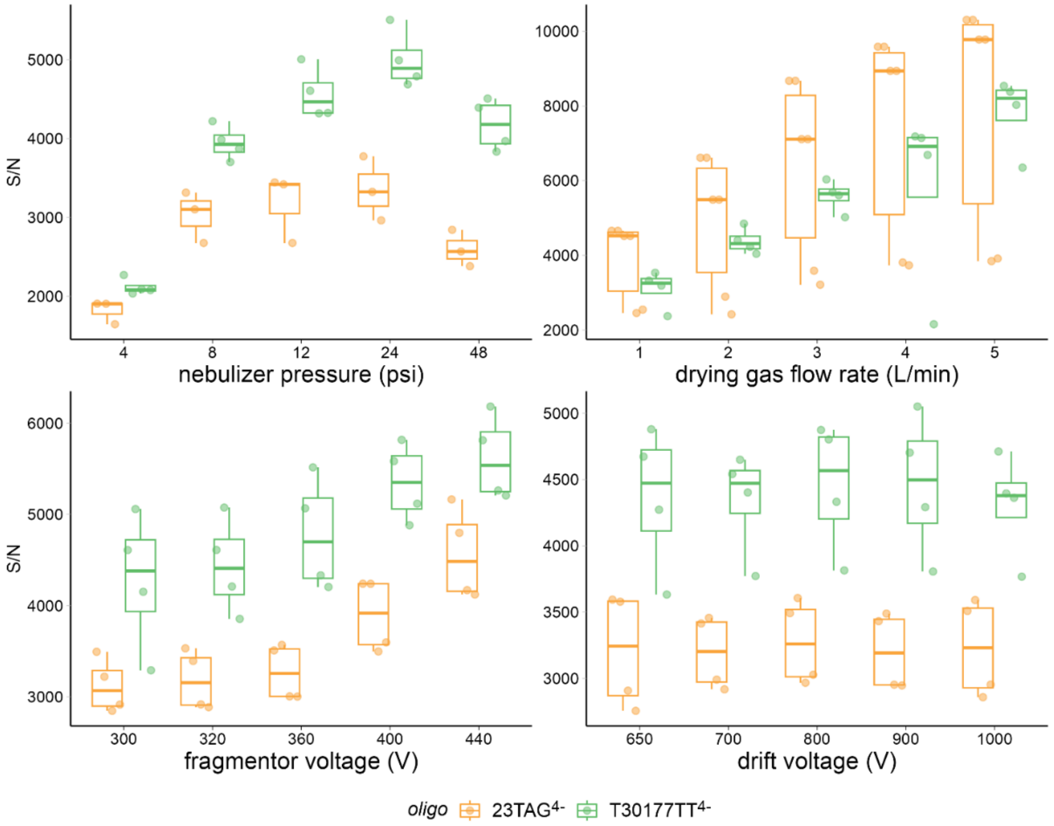


*Figure S12. Comparison of S/N ratio as a function of the different parameters of interest for the two oligos presented in the main text (23TAG·2K⁺ and T30177TT·2K⁺) indicated with different colors, both at charge state 4-.*

### *Arrival time distributions*

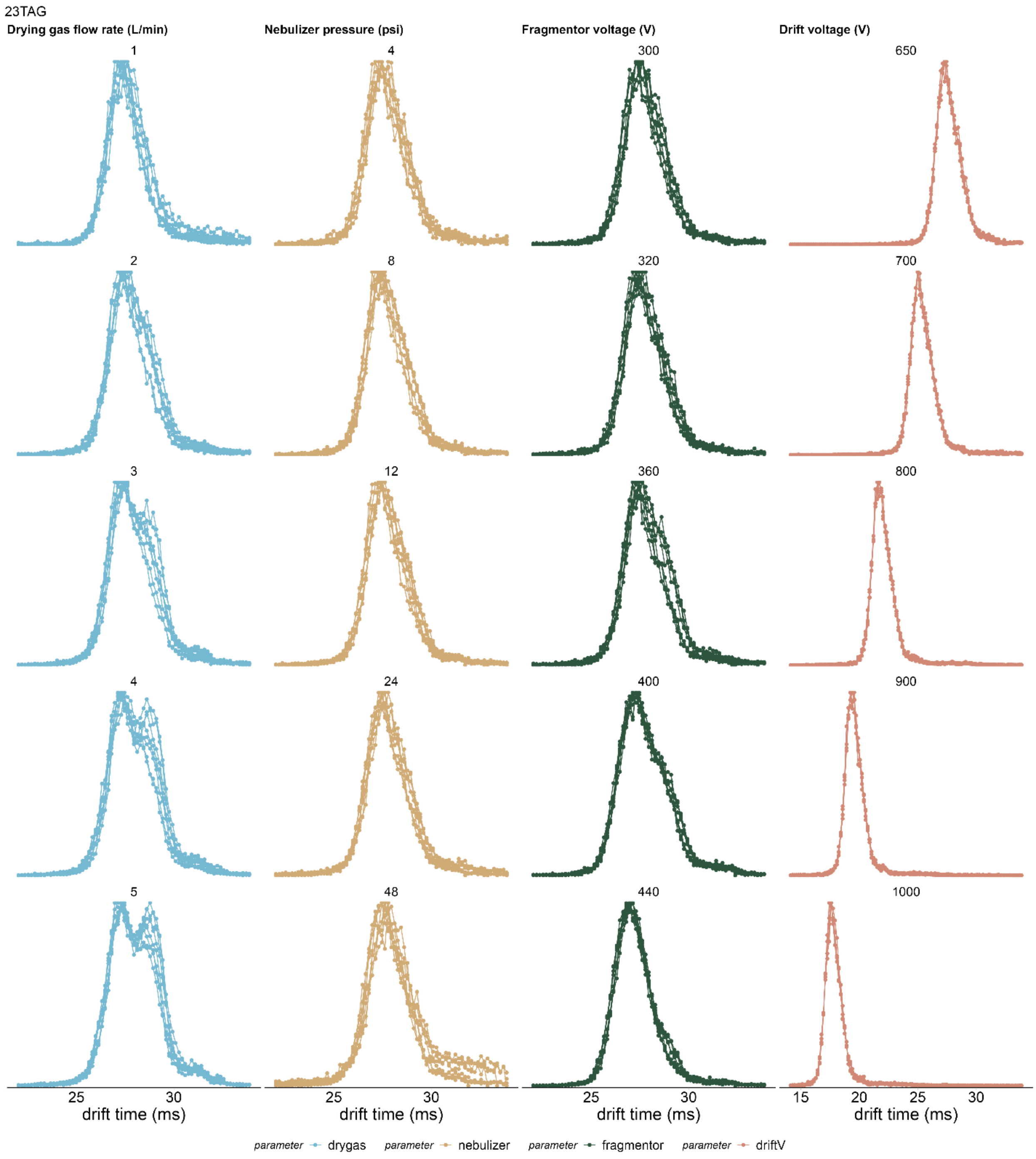


*Figure S13. Comparison of ATDs (intensity-normalized) of 23TAG·2K⁺ for the different parameters examined; each column corresponds to a parameter and each line to a different parameter value.*

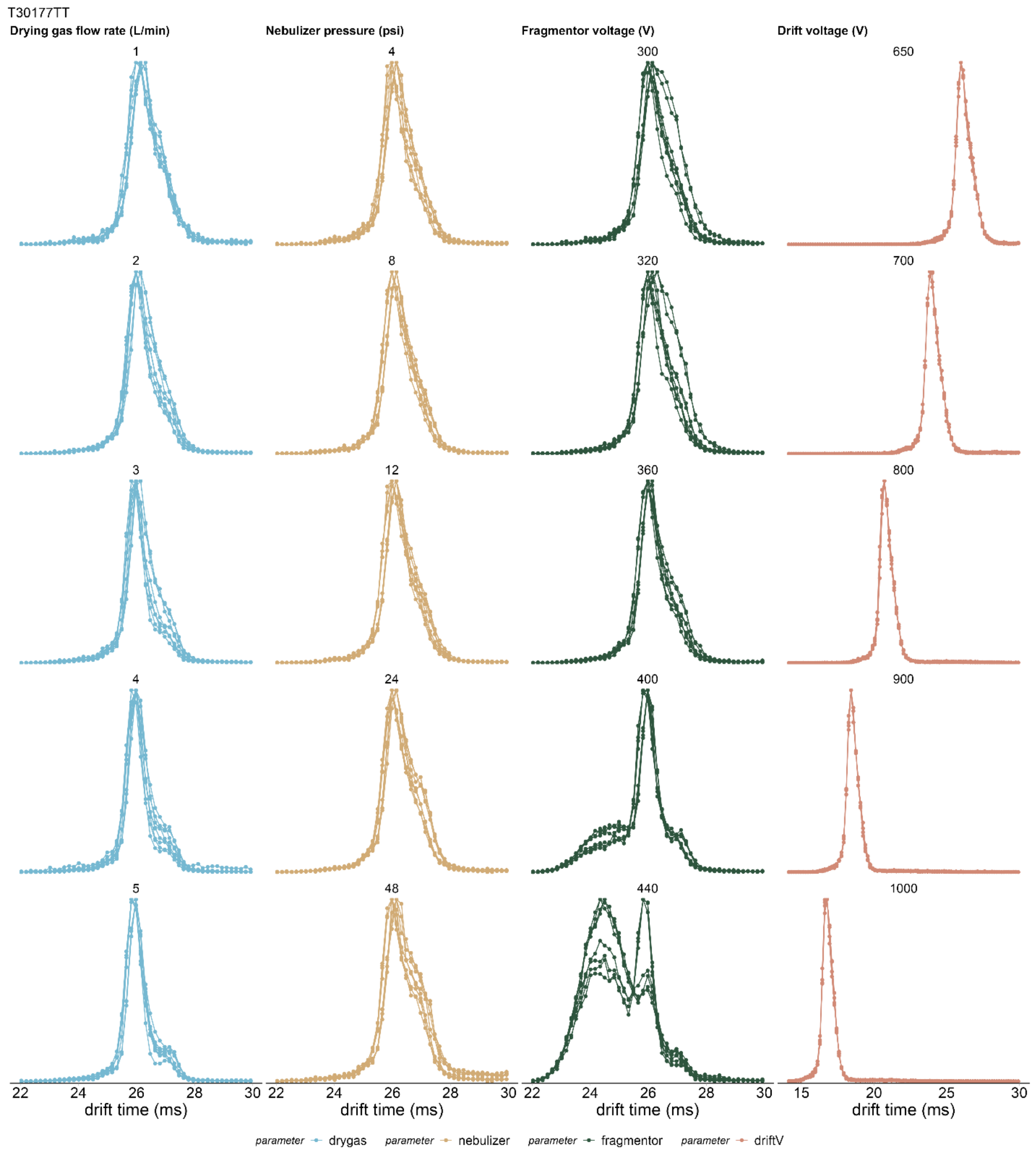


*Figure S14. Comparison of ATDs (intensity normalized) of T30177TT 2K⁺ for different values (rows) of different parameters (columns). The colors correspond to different H/D mixing times.*

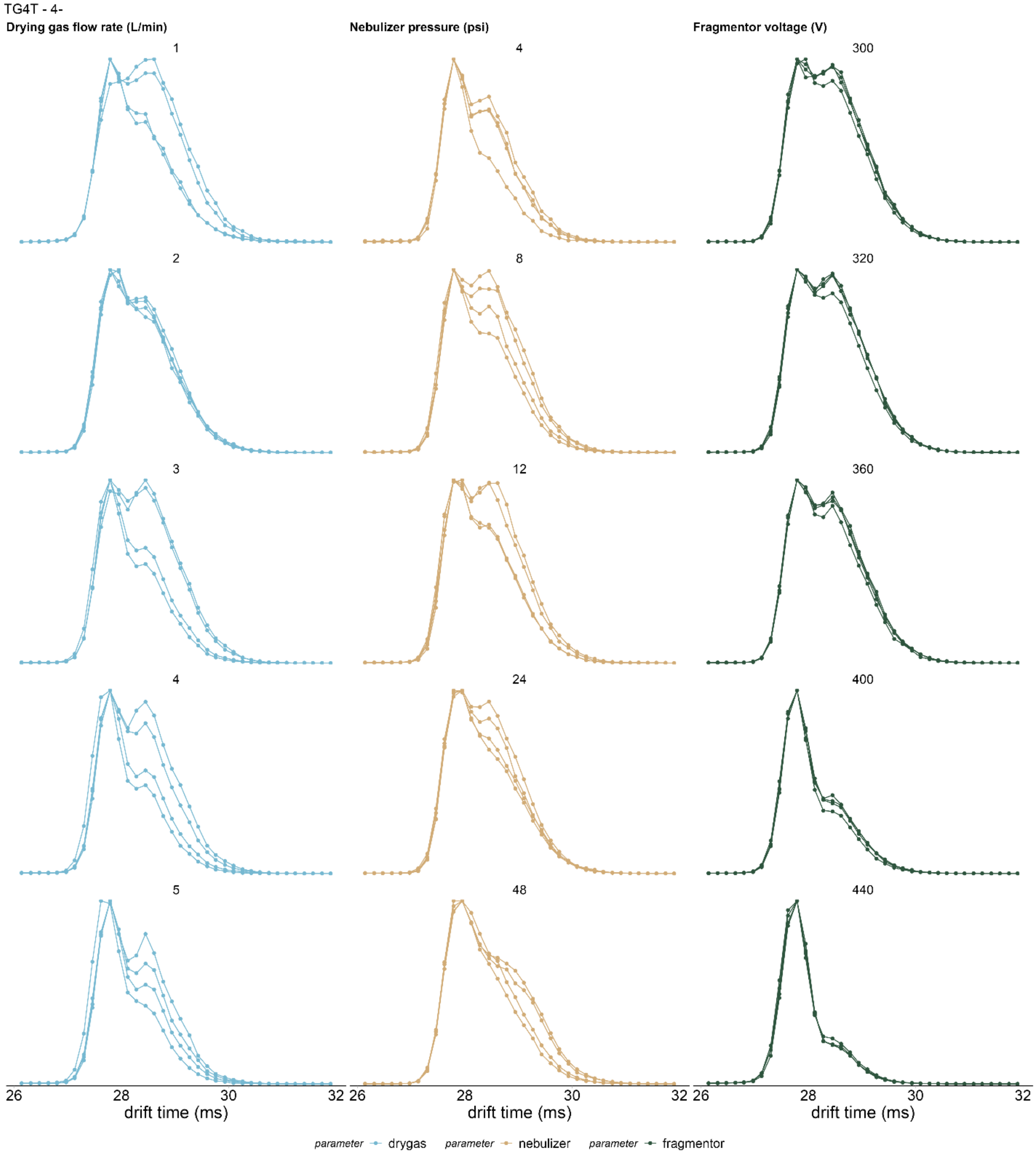


*Figure S15. Comparison of ATDs (intensity normalized) of TG4T 3NH$_4^+$ (4- charge state) for different values (rows) of different parameters (columns). The colors correspond to different H/D mixing times.*

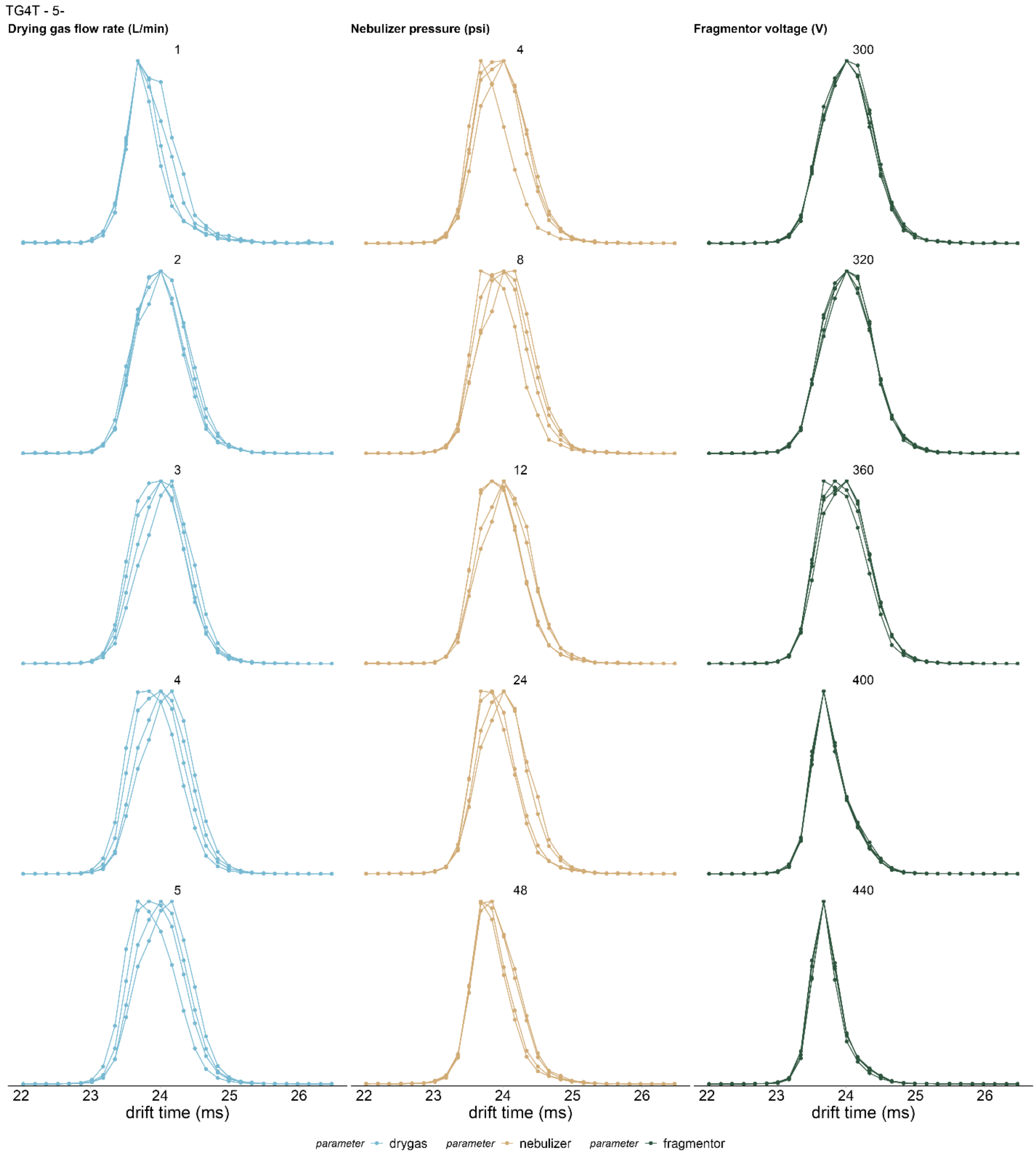


*Figure S16. Comparison of ATDs (intensity normalized) of TG4T 3NH$_4^+$ (5- charge state) for different values (rows) of different parameters (columns). The colors correspond to different H/D mixing times.*

#### *Influence of the buffer cation: case of TG4T*

The S/N ratio on TG4T in ammonium buffer is charge state dependent and was normalized by the average S/N (Eq. 5) of the default parameter value to compare more accurately this effect.

The trends observed on the nebulizer pressure for T30177TT and 23TAG seem to be reproduced with TG4T as it gradually improves the signal-to-noise ration except at 48 psi. The fragmentor, however, has a positive effect until 400V and is lost afterwards. The drying gas has a charge-state-dependent impact on S/N, increasing by a factor 2 for the 5- charge state but slightly decreasing for 4-.

Contrary to the other oligonucleotides in TMAA/KCl, this structure is much more resistant to the fragmentor voltage increase as no compaction was observed on the ATD, however a decrease of the S/N was observed at 440V.

The differences exhibited for the fragmentor is explained by ammonium loss in the gas-phase due to ion activation. This phenomenon causes difference in the NUS upon charge state. NUS values slightly increase with the nebulizer pressure and is constant with the fragmentor until 440V where the drop is smaller than 23TAG and T30177TT. The drying gas variation is constant for 5- but drops at 4-. The ammonium loss causes opening of the structure and might trigger exchange upon refolding with a fully protonated ammonium ion.[14]

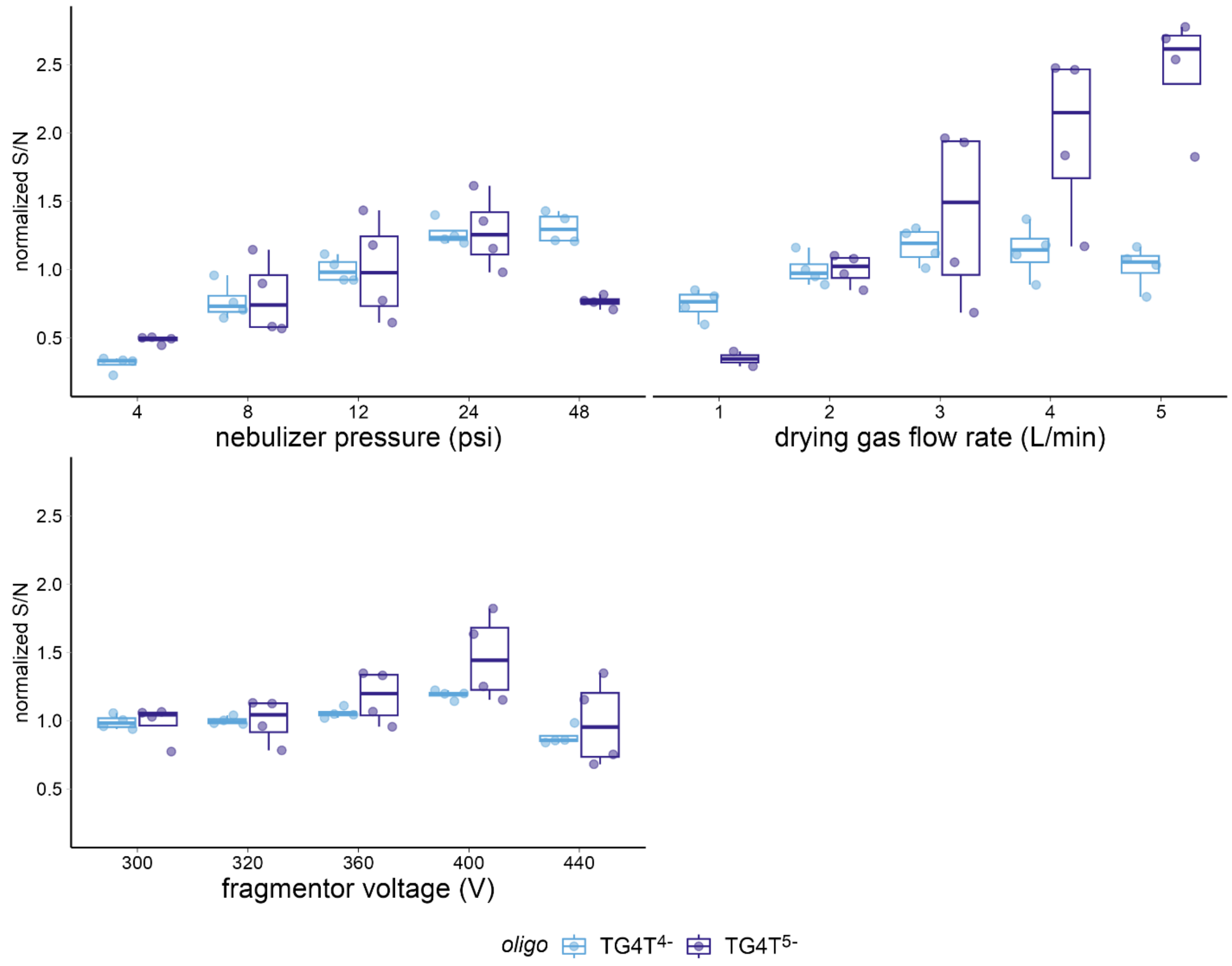


*Figure S17. Comparison of S/N ratio as a function of the different parameters of interest for the TG4T·3NH$_4^+$ G-quadruplex.*

### Agilent 6560 IMS Q-TOF base settings

1. *Source*:

- Gas temperature: 280°C
- Drying gas: 2.0 L/min
- Nebulizer pressure: 12 psig
- Capillary voltage: 3500 V
- Fragmentor: 320 V
- Skimmer: −27.0 V
- Sheath gas temperature: 1024°C

*Ion mobility front funnel:*

- High pressure funnel delta
- High pressure funnel RF:
- Trap funnel delta: −121 V
- Trap funnel RF: 89 V
- Trap funnel exit: −10.9 V

*Ion mobility trap:*

- Trap entrance grid low: −72 V
- Trap entrance grid delta: −70 V
- Trap entrance: −69 V
- Trap exit: −67.1 V
- Trap exit grid 1 low: −69.1 V
- Trap exit grid 1 delta: −64 V
- Trap exit grid 2 low: −72 V
- Trap exit grid 2 delta: −63 V

IM Drift tube:

- Drift gas: Helium
- Ion Mobility drift tube entrance: −650.0 V
- Ion Mobility drift tube exit: −209.5 V

*Ion mobility rear funnel:*

- Rear funnel RF: 179 V
- Rear funnel exit: −35.5 V
- IM Hex entrance: −31.8 V

*Optics 1:*

- Oct 1 DC: −25 V
- Lens 1: −23.0 V
- Lens 2: −16.5 V

*Quadrupole:*

- Quad DC: −21 V
- Postfilter DC: −20 V

*Cell:*

- Gas flow: 20.0 psig
- Cell entrance: −20 V
- Hex DC: −20.0 V
- Hex Delta: 3.0 V

*Optics 2:*

- Horiz Q: 1.50 V
- Oct2 DC: −14.6 V
- Ion Focus: −10 V
- VertQ: −12.90 V

*TOF:*

- Mirror Back: −1250 V
- Mirror Mid: 1674.9 V
- Mirror front: 7000 V
- Acc focus: 1950 V
- Puller offset: −32 V
- Puller: 700 V
- Pusher: −1200 V
- Min range: 62080 ns

### CCS conversion

The conversion from arrival time to CCS was performed on DNA samples containing 10 μM G4 at 9% D in the same buffer used for H/D experiments. The CCS calibration is performed by acquiring MS at 5 drift voltages: −650V, −700V, −800V, −900V, −1000V. The arrival time $t_A$ is proportional to the $\frac{p}{\Delta V}$ ratio $p$ being the pressure in the drift tube and $\Delta V$ the voltage difference applied at the extremities of the tube of length $L$. (Equation 6)

$$t_A = \frac{L^2}{K_0}\frac{T_0}{p_0 T}\left(\frac{p}{\Delta V}\right) + t_0 \tag{6}$$

Plotting $t_A$ as a function of $\frac{p}{\Delta V}$ gives access to the reduced ion-mobility $K_0$ from the slope and the time $t_0$ before detection from the intercept. The conversion from arrival time to CCS values was done by reinjecting $K_0$ in Equation 7.[12,15,16]

$$CCS = \frac{3}{16}\sqrt{\frac{2\pi}{\mu k_B T}}\frac{ze}{N_0}\frac{1}{K_0} \tag{7}$$

where $\mu$ is the reduced mass accounting for the DNA and the helium masses, $k_B$ the Boltzmann constant, $ze$ the charge of the ion, $T$ the drift tube temperature and $N_0$ the standard gas density (=2.687 × $10^{25}$ $m^{-3}$).

**23TAG•1K**

*100 mM TMAA + 1 mM KCl – fragmentor: 320 V*

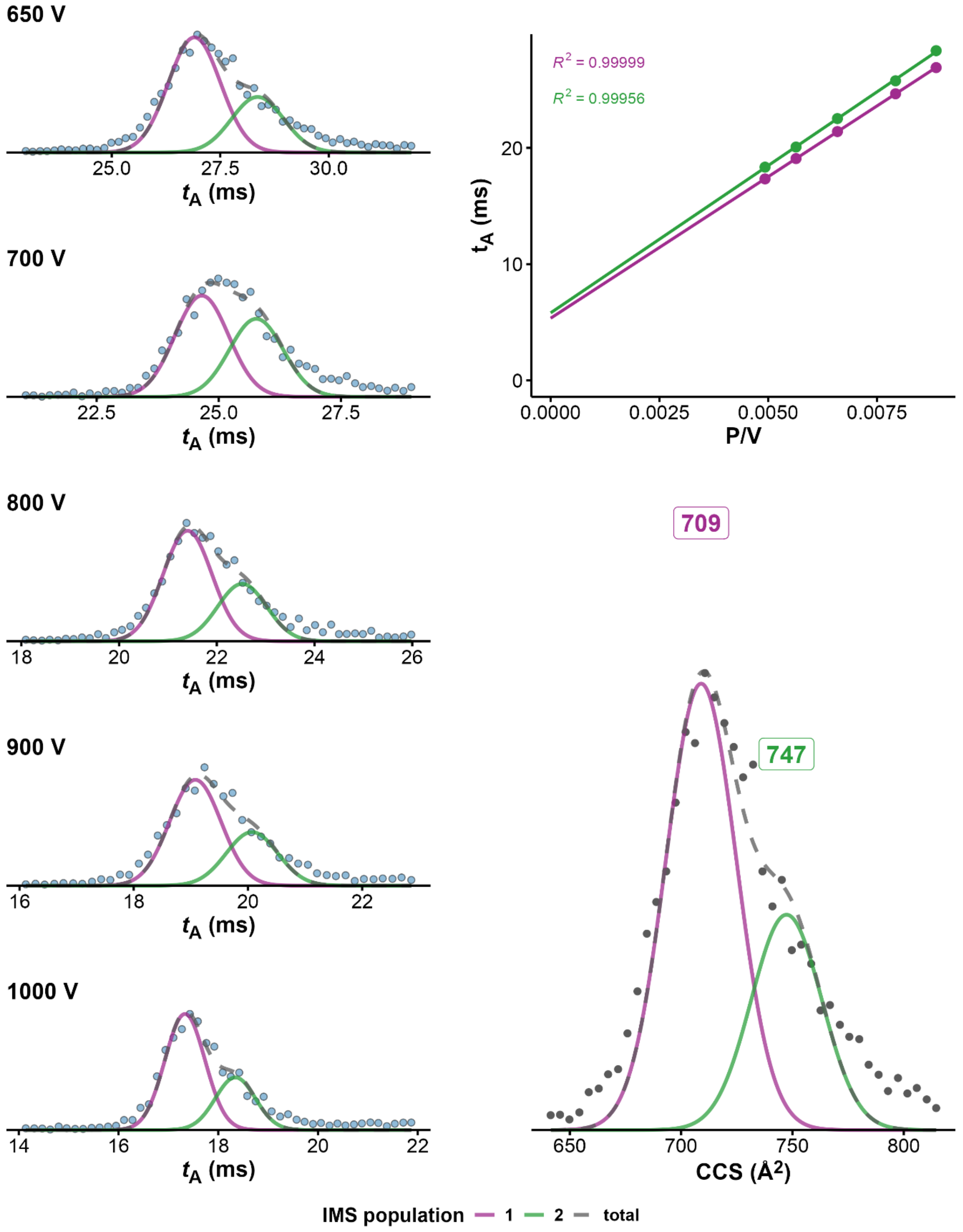


*Figure S18. CCS determination of 23TAG·1K⁺ (100mM TMAA + 1mM KCl fragmentor = 370V, z=4-). On the left panel ATDs at different drift voltages, on top right panel, $K_0$ determination from linear regression of arrival time at different voltages. Bottom right: CCS distribution obtained after conversion of the ATD at 650V*

**23TAG●2K**

*100 mM TMAA + 1 mM KCl – fragmentor: 320 V*

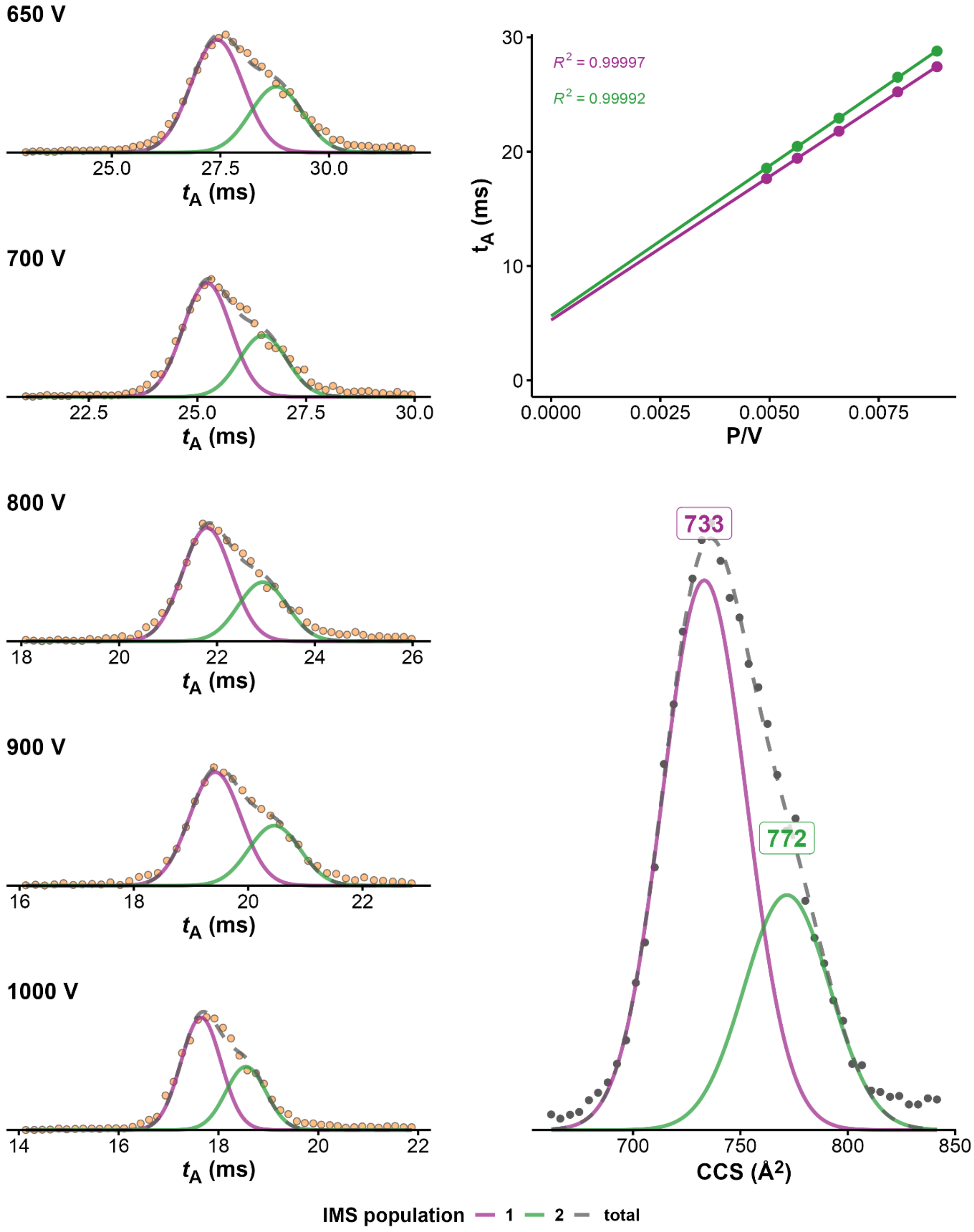


*Figure S19. CCS determination of 23TAG·2K$^+$ (100mM TMAA + 1mM KCl fragmentor = 320V, z=4-). On the left panel ATDs at different drift voltages, on top right panel, $K_0$ determination from linear regression of arrival time at different voltages. Bottom right: CCS distribution obtained after conversion of the ATD at 650V*

**23TAG•2K**

*100 mM TMAA + 1 mM KCl – fragmentor: 350 V*

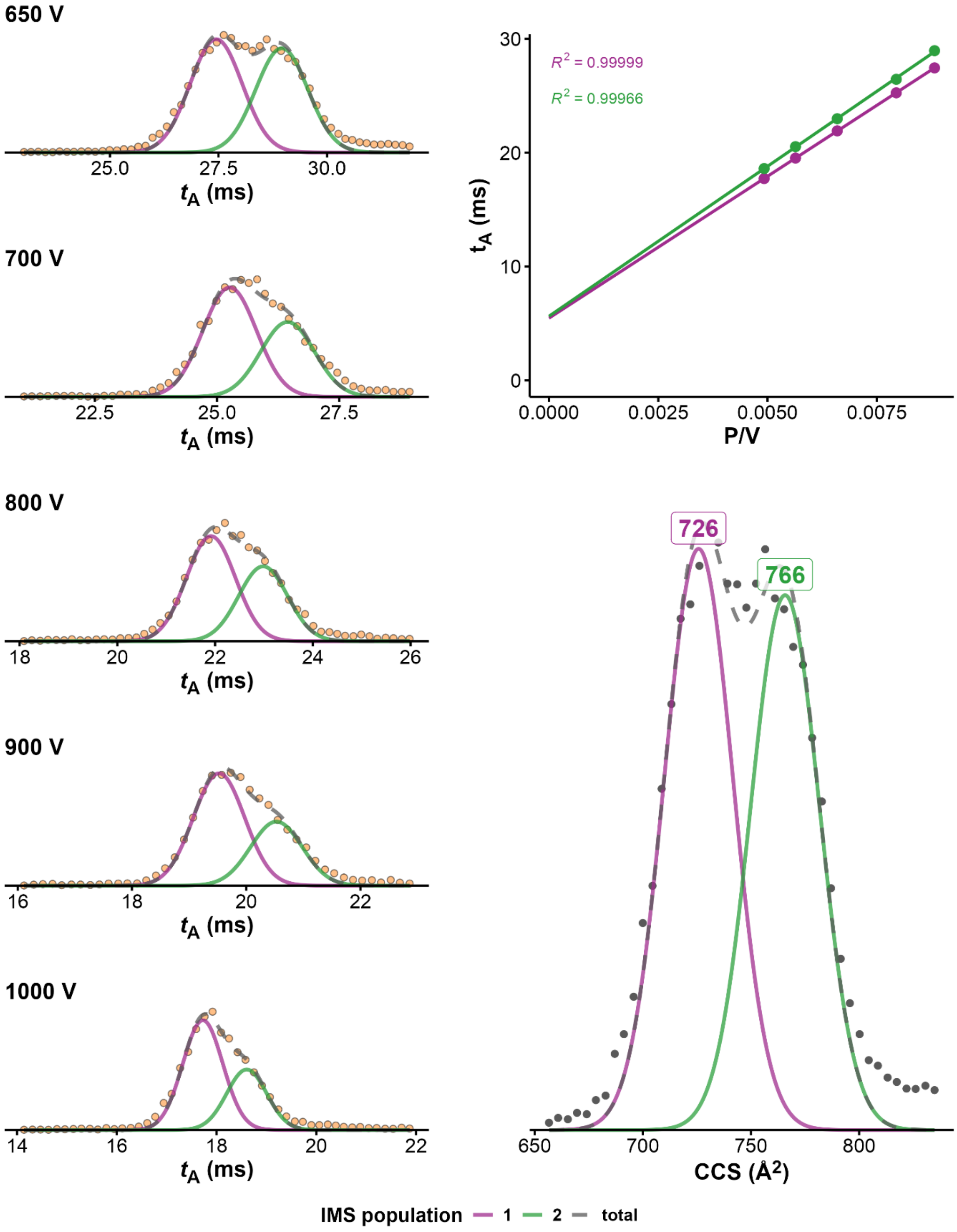


*Figure S20. CCS determination of 23TAG·2K⁺ (100mM TMAA + 1mM KCl fragmentor = 350V, z=4-). On the left panel ATDs at different drift voltages, on top right panel, $K_0$ determination from linear regression of arrival time at different voltages. Bottom right: CCS distribution obtained after conversion of the ATD at 650V*

**23TAG•2K**

*100 mM TMAA + 1 mM KCl – fragmentor: 370 V*

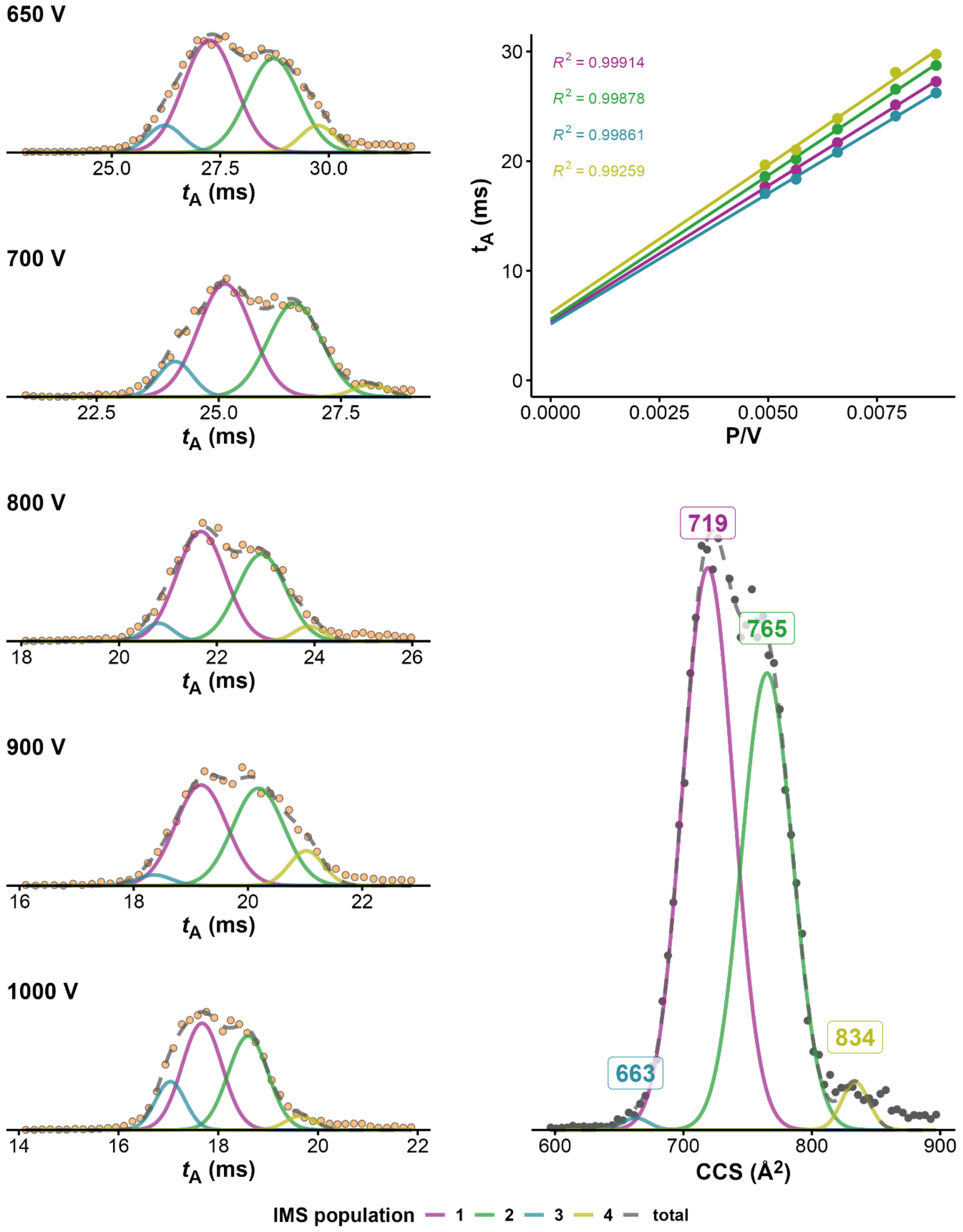


*Figure S21. CCS determination of 23TAG·2K⁺ (100mM TMAA + 1mM KCl fragmentor = 370V, z=4-). On the left panel ATDs at different drift voltages, on top right panel, $K_0$ determination from linear regression of arrival time at different voltages. Bottom right: CCS distribution obtained after conversion of the ATD at 650V*

**5YEY●1K**

*100 mM TMAA + 1 mM KCl – fragmentor: 320 V*

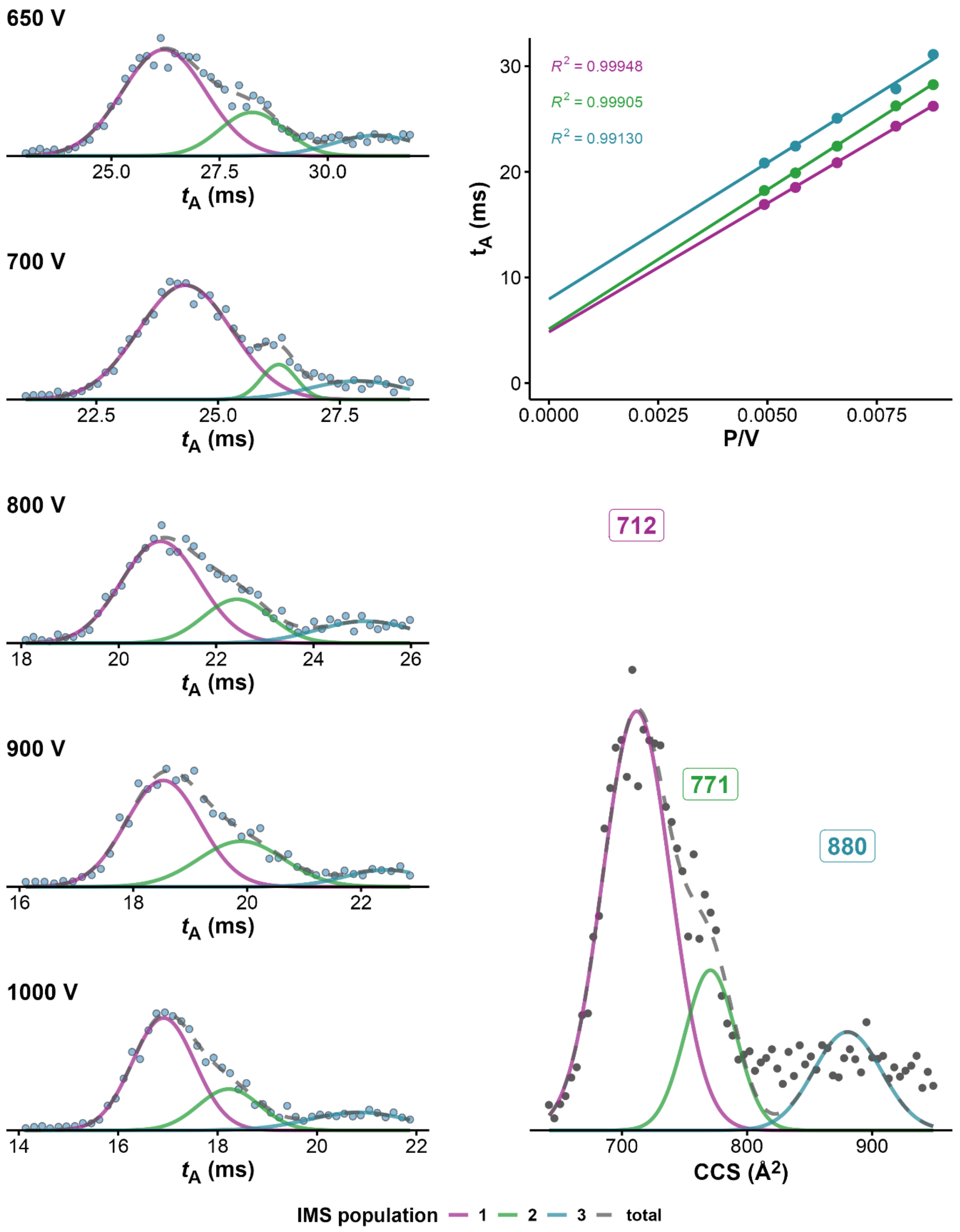


*Figure S22. CCS determination of 5YEY·1K⁺ (100mM TMAA + 1mM KCl fragmentor = 320V, z=4-). On the left panel ATDs at different drift voltages, on top right panel, $K_0$ determination from linear regression of arrival time at different voltages. Bottom right: CCS distribution obtained after conversion of the ATD at 650V*

**5YEY•2K**

*100 mM TMAA + 1 mM KCl – fragmentor: 320 V*

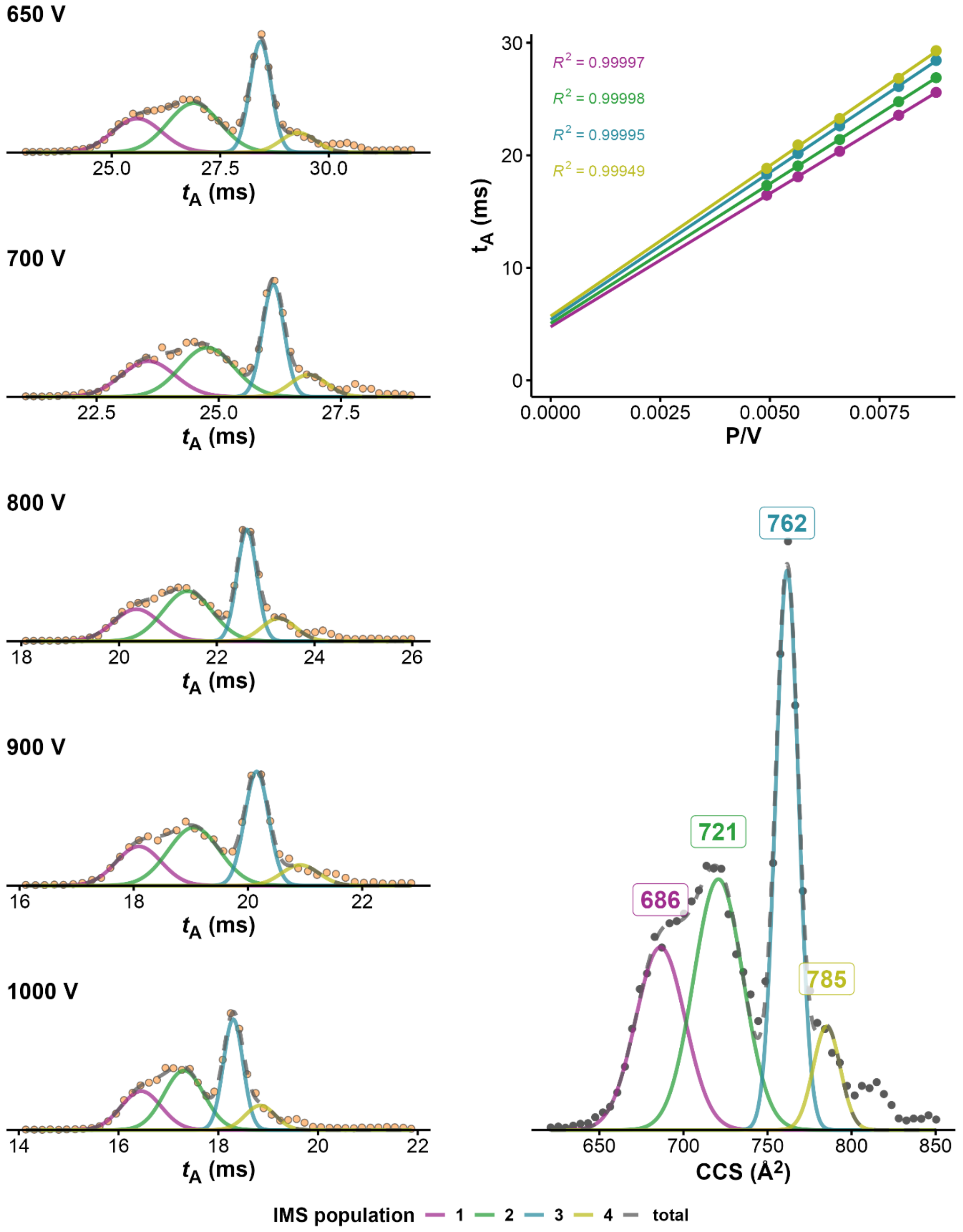


*Figure S23. CCS determination of 5YEY·2K⁺ (100mM TMAA + 1mM KCl, fragmentor = 320V, z=4-). On the left panel ATDs at different drift voltages, on top right panel, $K_0$ determination from linear regression of arrival time at different voltages. Bottom right: CCS distribution obtained after conversion of the ATD at 650V*

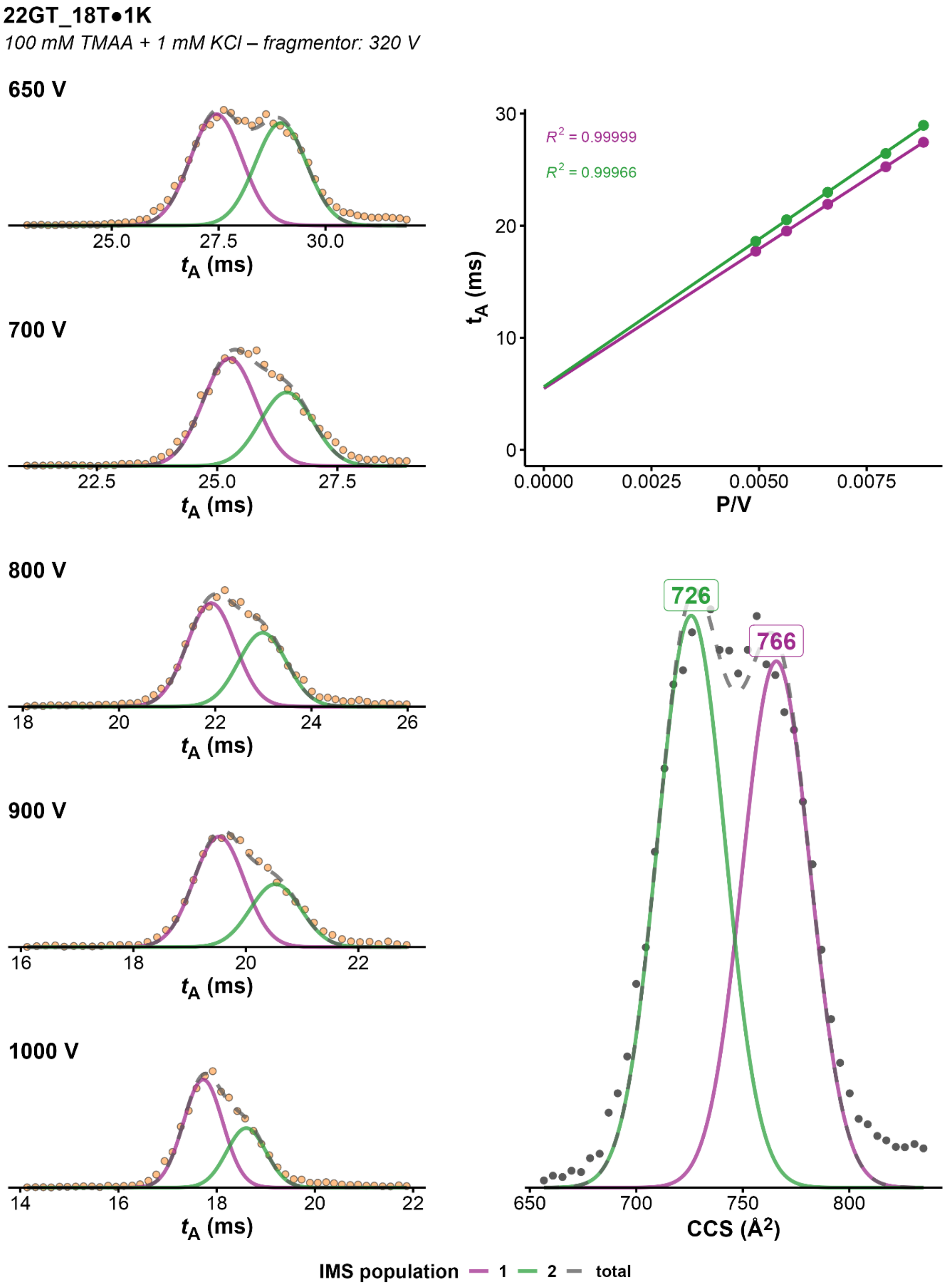


*Figure S24. CCS determination of 22GT_18T·2K⁺ (100mM TMAA + 1mM KCl, fragmentor = 320V, z=4-). On the left panel ATDs at different drift voltages, on top right panel, $K_0$ determination from linear regression of arrival time at different voltages. Bottom right: CCS distribution obtained after conversion of the ATD at 650V*

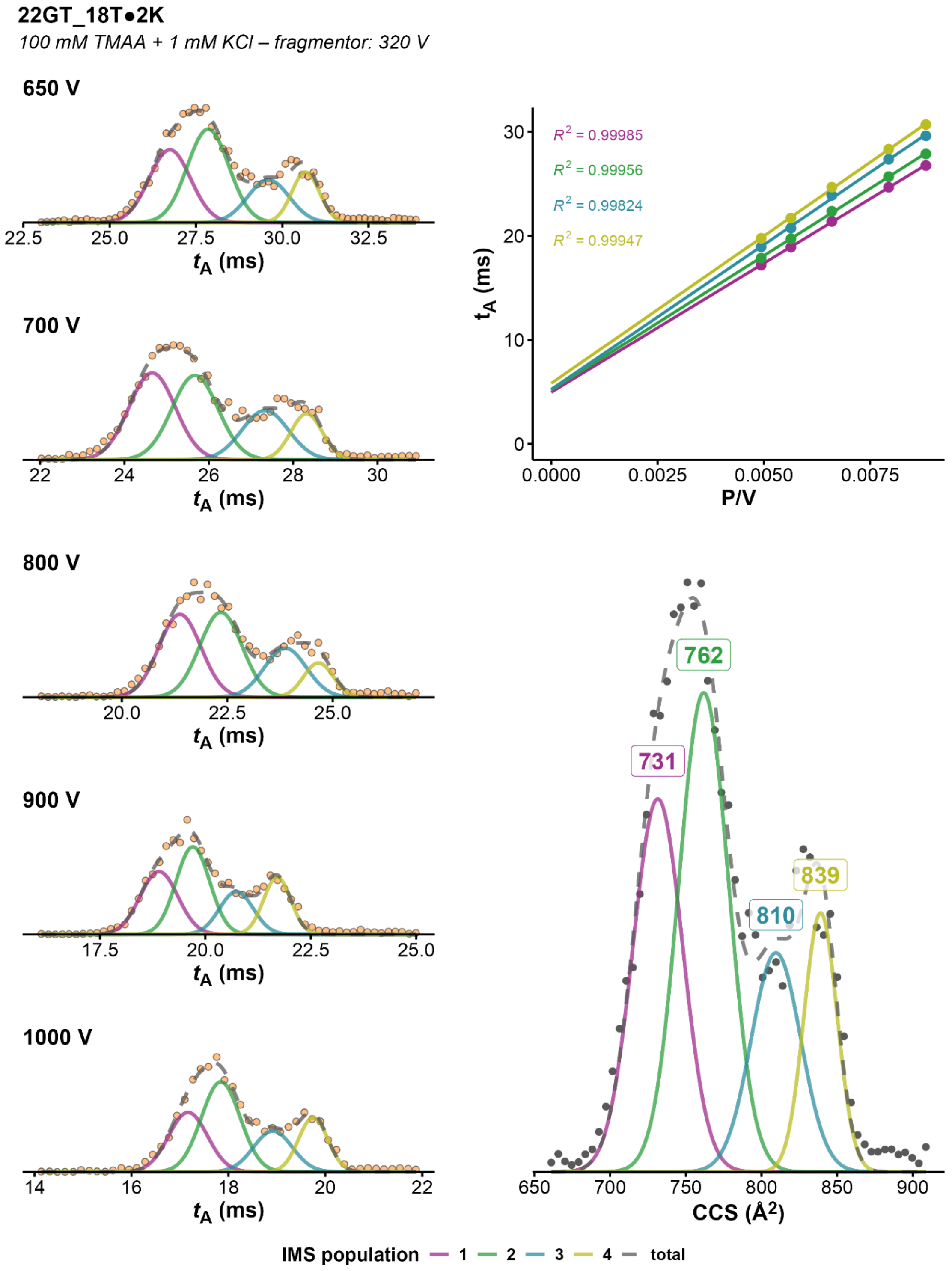


*Figure S25. CCS determination of 22GT_18T·2K⁺ (100mM TMAA + 1mM KCl, fragmentor = 320V, z=4-). On the left panel ATDs at different drift voltages, on top right panel, $K_0$ determination from linear regression of arrival time at different voltages. Bottom right: CCS distribution obtained after conversion of the ATD at 650V*

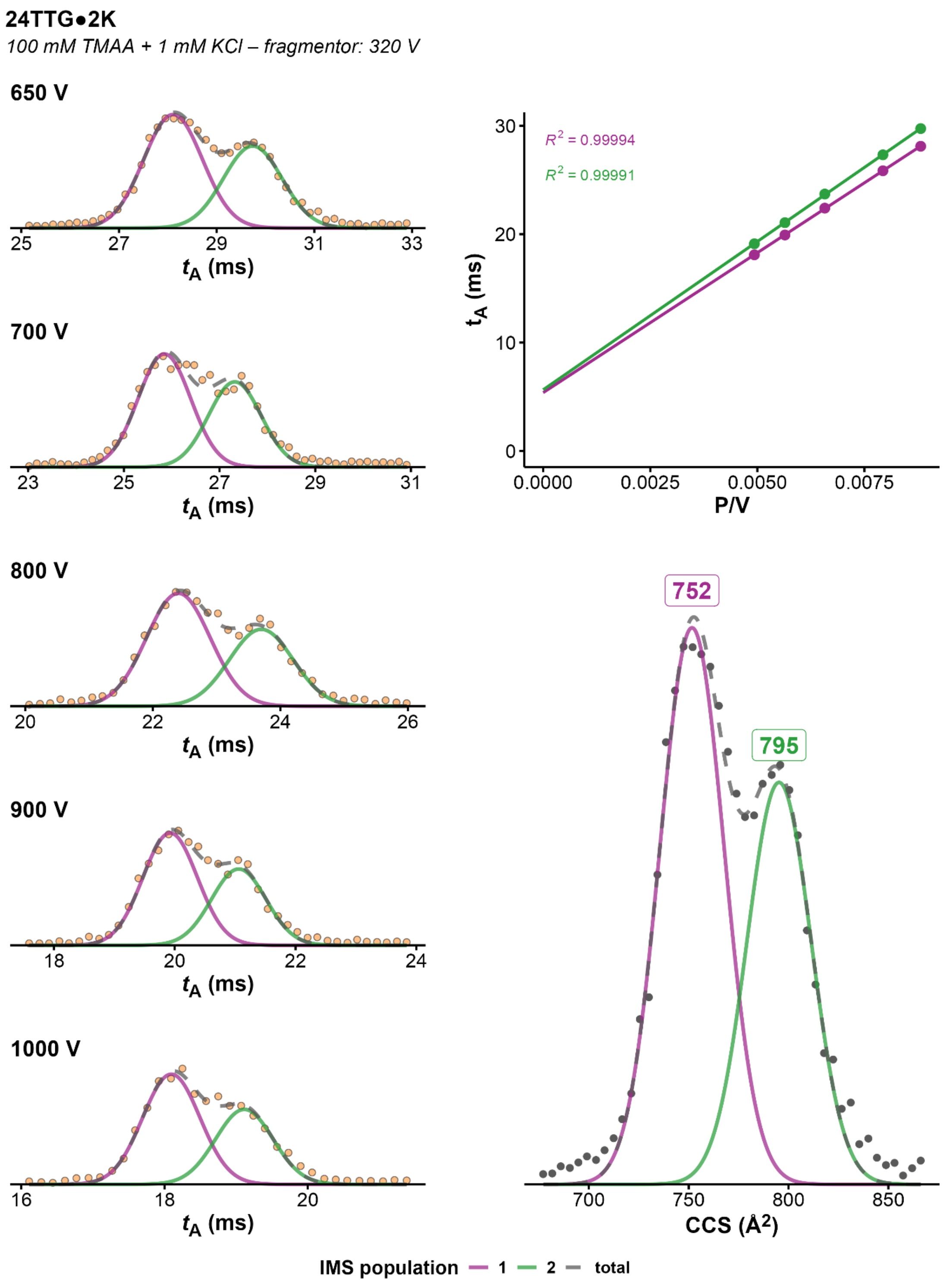


*Figure S26. CCS determination of 24TTG·2K+ (100mM TMAA + 1mM KCl, fragmentor = 320V, z=4-). On the left panel ATDs at different drift voltages, on top right panel, $K_0$ determination from linear regression of arrival time at different voltages. Bottom right: CCS distribution obtained after conversion of the ATD at 650V*

**24TTG•2K**

*100 mM TMAA + 0.5 mM KCl – fragmentor: 320 V*

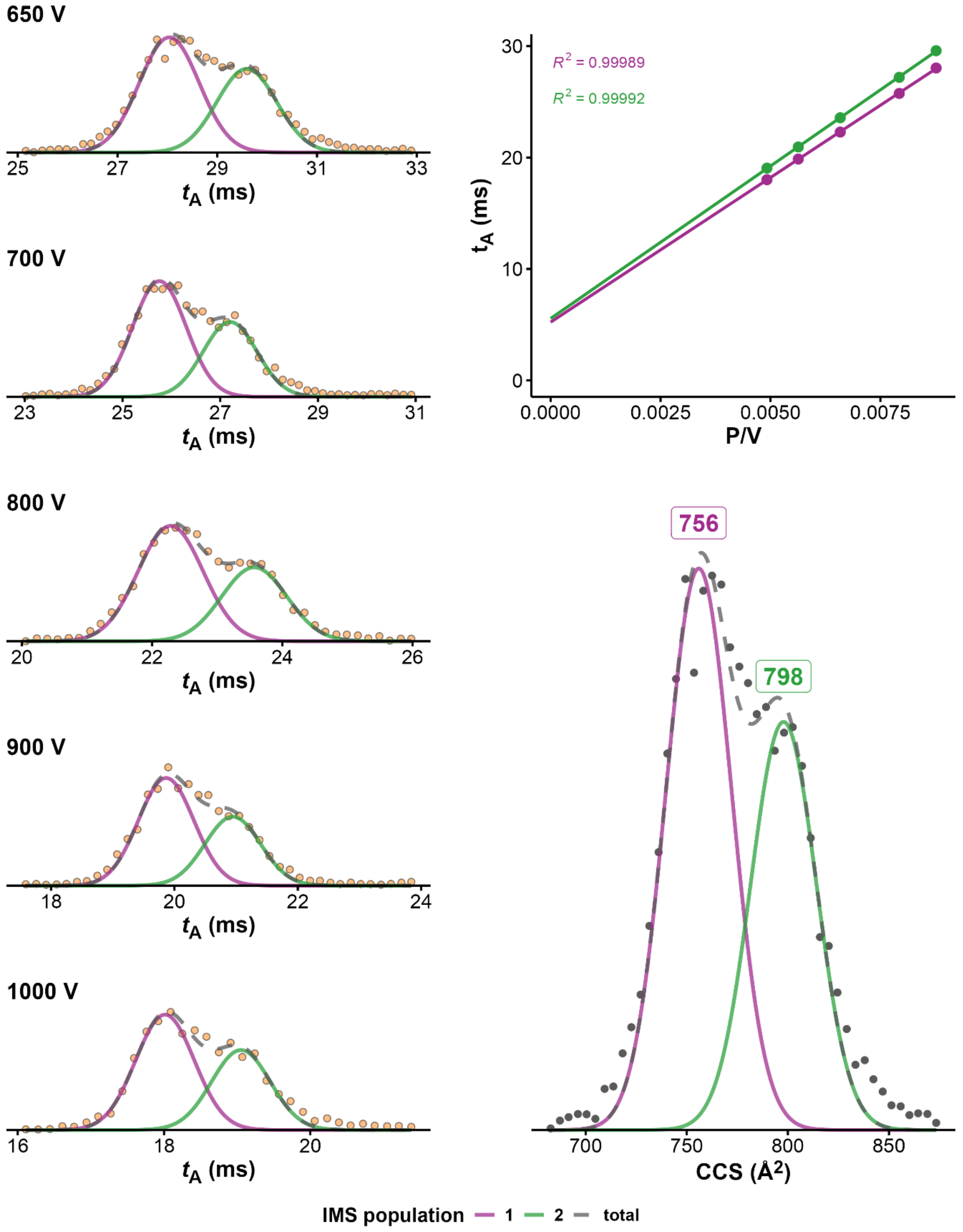


*Figure S27. CCS determination of 24TTG · 2K⁺ (100mM TMAA + 0.5mM KCl, fragmentor = 320V, z=4-). On the left panel ATDs at different drift voltages, on top right panel, $K_0$ determination from linear regression of arrival time at different voltages. Bottom right: CCS distribution obtained after conversion of the ATD at 650V*

**24TTG•2K**

*100 mM TMAA + 0.1 mM KCl – fragmentor: 320 V*

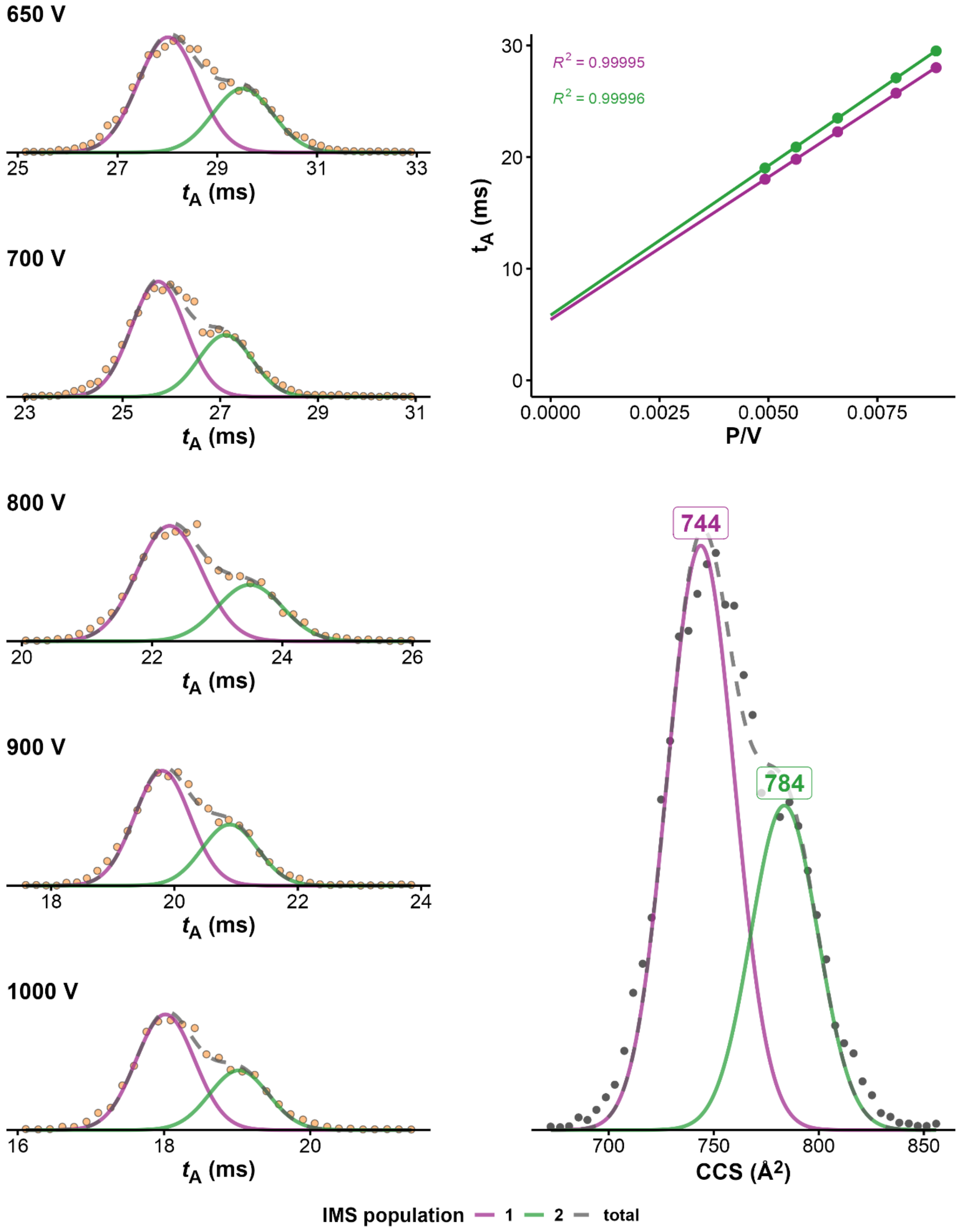


*Figure S28. CCS determination of 24TTG·2K⁺ (100mM TMAA + 0.1mM KCl, fragmentor = 320V, z=4-). On the left panel ATDs at different drift voltages, on top right panel, $K_0$ determination from linear regression of arrival time at different voltages. Bottom right: CCS distribution obtained after conversion of the ATD at 650V*

**222T•1K**

*100 mM TMAA + 0.1 mM KCl – fragmentor: 320 V*

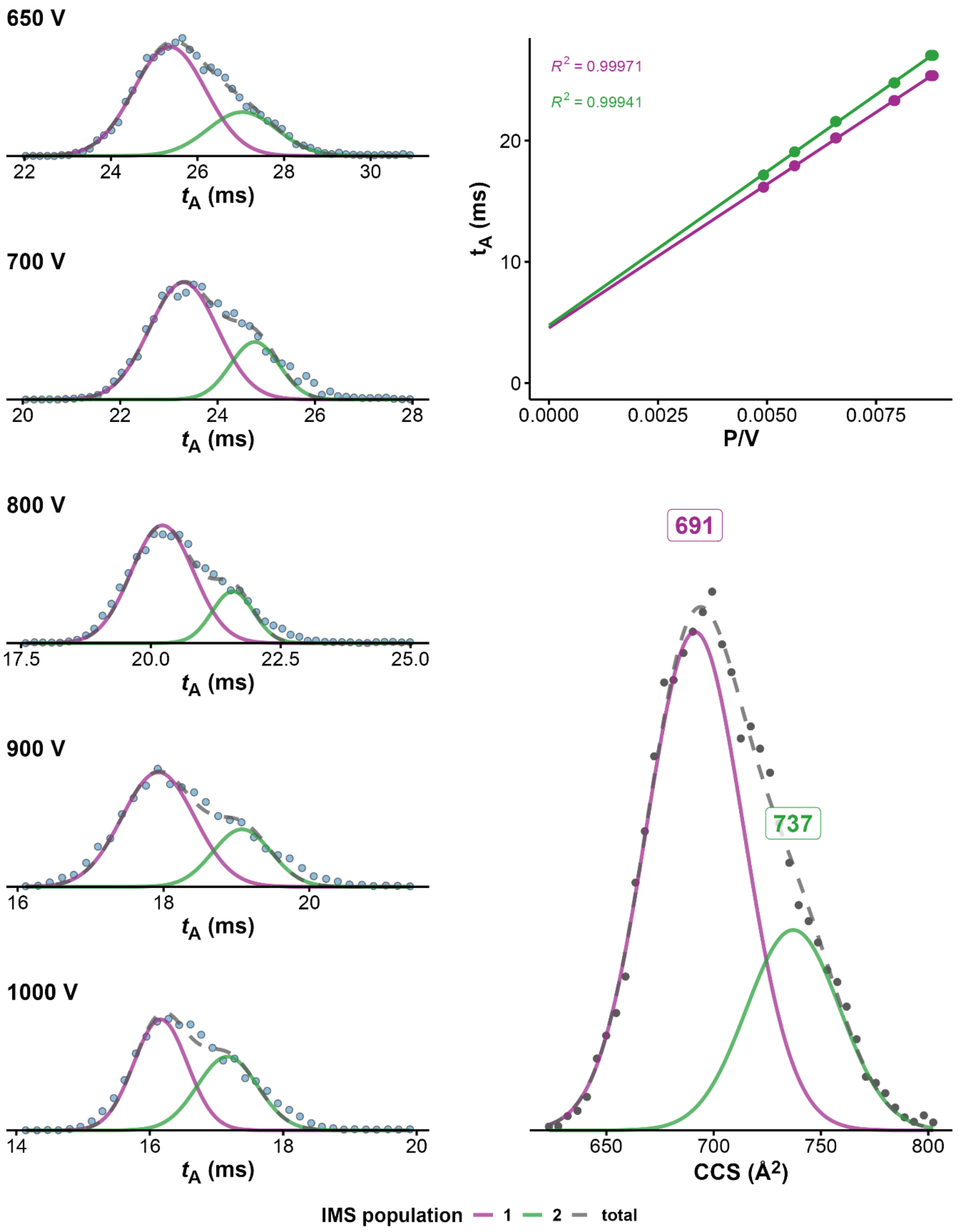


*Figure S29. CCS determination of 222T·1K⁺ (100mM TMAA + 0.1mM KCl, fragmentor = 320V, z=4-). On the left panel ATDs at different drift voltages, on top right panel, $K_0$ determination from linear regression of arrival time at different voltages. Bottom right: CCS distribution obtained after conversion of the ATD at 650V*

**222T•2K**

*100 mM TMAA + 0.1 mM KCl – fragmentor: 320 V*

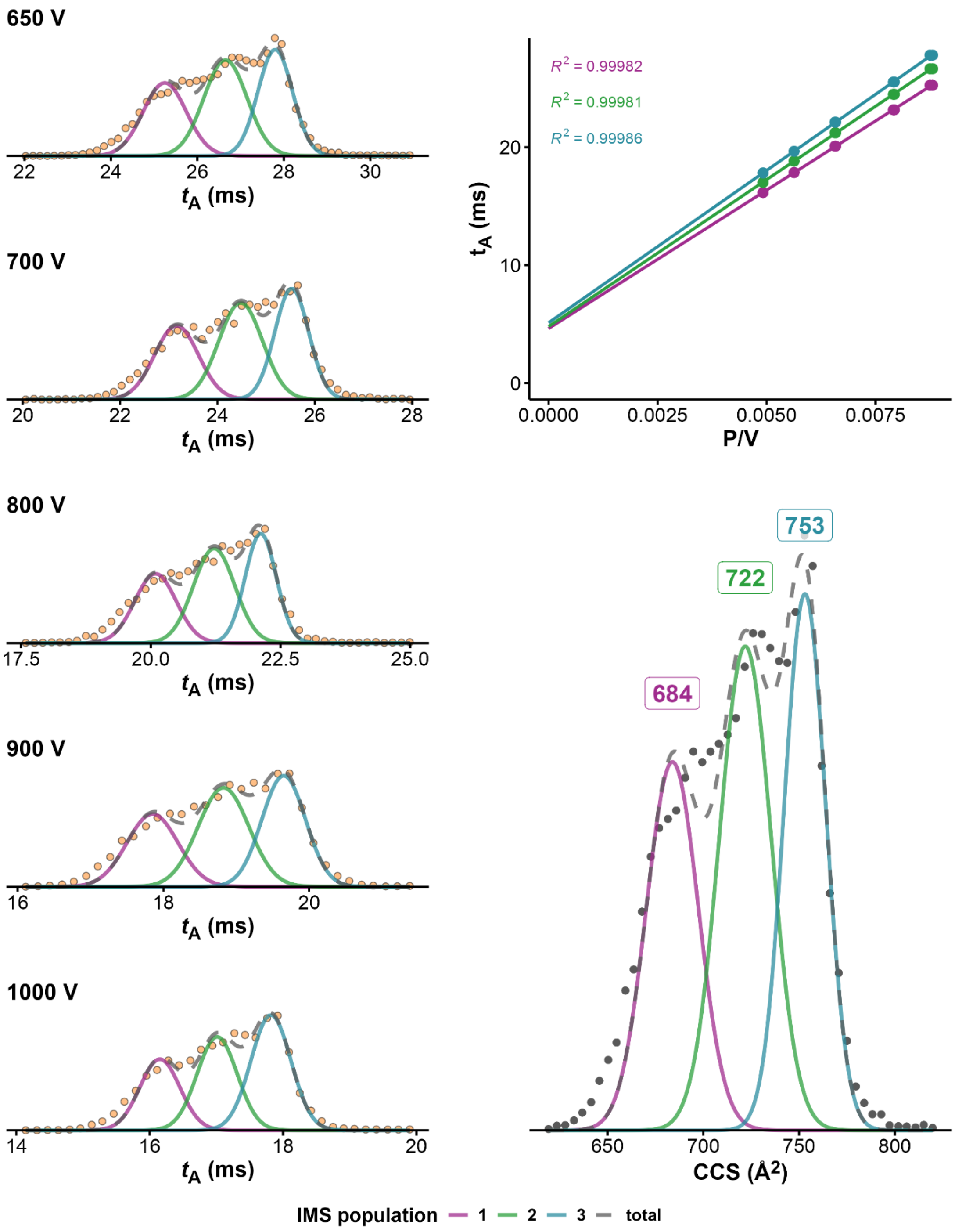


*Figure S30. CCS determination of 222T·2K⁺ (100mM TMAA + 0.1mM KCl, fragmentor = 320V, z=4-). On the left panel ATDs at different drift voltages, on top right panel, $K_0$ determination from linear regression of arrival time at different voltages. Bottom right: CCS distribution obtained after conversion of the ATD at 650V*

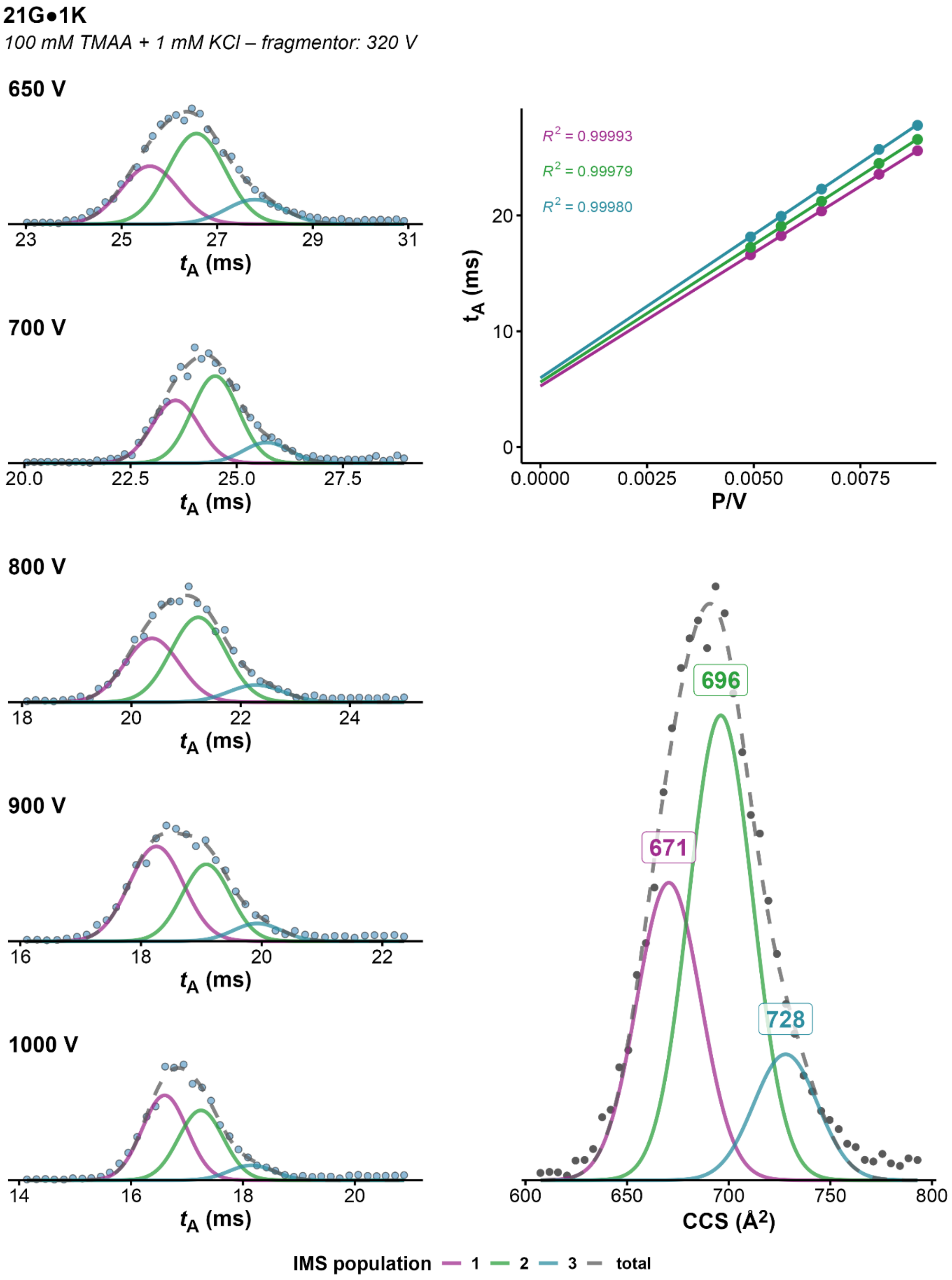


*Figure S31. CCS determination of 21G·1K⁺ (100mM TMAA + 1mM KCl, fragmentor = 320V, z=4-). On the left panel ATDs at different drift voltages, on top right panel, $K_0$ determination from linear regression of arrival time at different voltages. Bottom right: CCS distribution obtained after conversion of the ATD at 650V*

**21G•2K**

*100 mM TMAA + 1 mM KCl – fragmentor: 320 V*

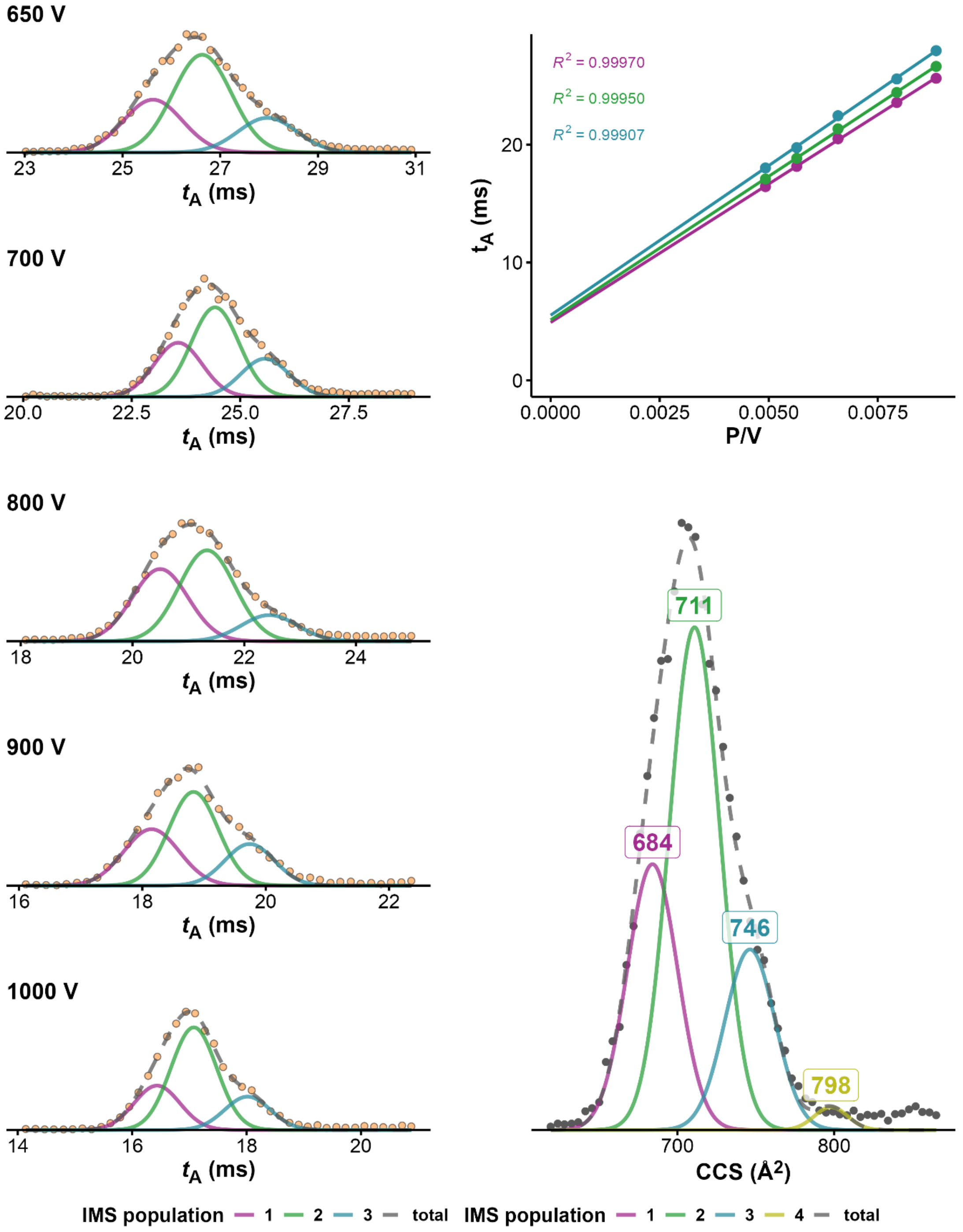


*Figure S32. CCS determination of $21G\cdot2K^+$ (100mM TMAA + 1mM KCl, fragmentor = 320V, z=4-). On the left panel ATDs at different drift voltages, on top right panel, $K_0$ determination from linear regression of arrival time at different voltages. Bottom right: CCS distribution obtained after conversion of the ATD at 650V*

**22AG•1K**

*100 mM TMAA + 1 mM KCl – fragmentor: 320 V*

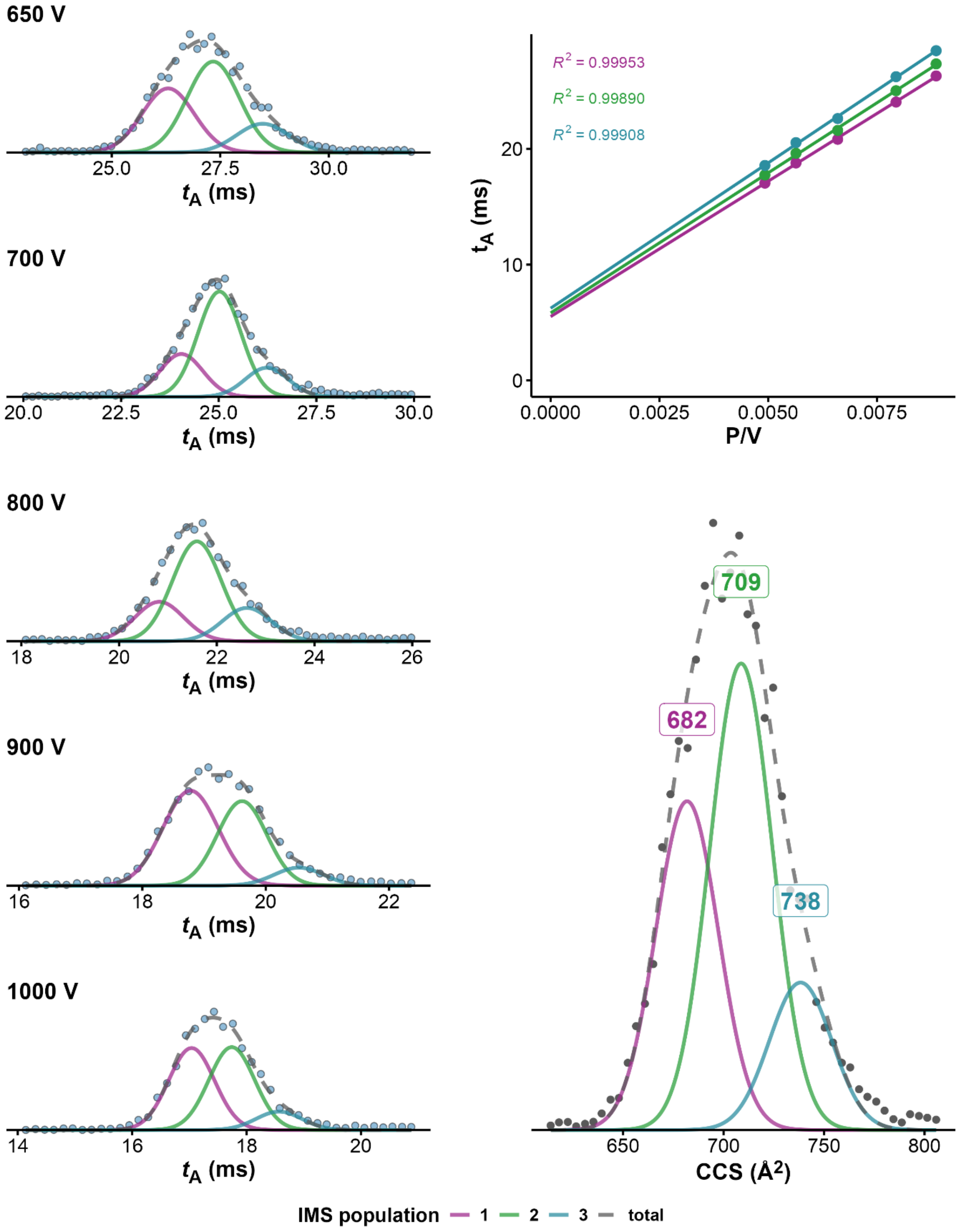


*Figure S33. CCS determination of 22AG · 1K⁺ (100mM TMAA + 1mM KCl, fragmentor = 320V, z=4-). On the left panel ATDs at different drift voltages, on top right panel, $K_0$ determination from linear regression of arrival time at different voltages. Bottom right: CCS distribution obtained after conversion of the ATD at 650V*

**22AG•2K**

*100 mM TMAA + 1 mM KCl – fragmentor: 320 V*

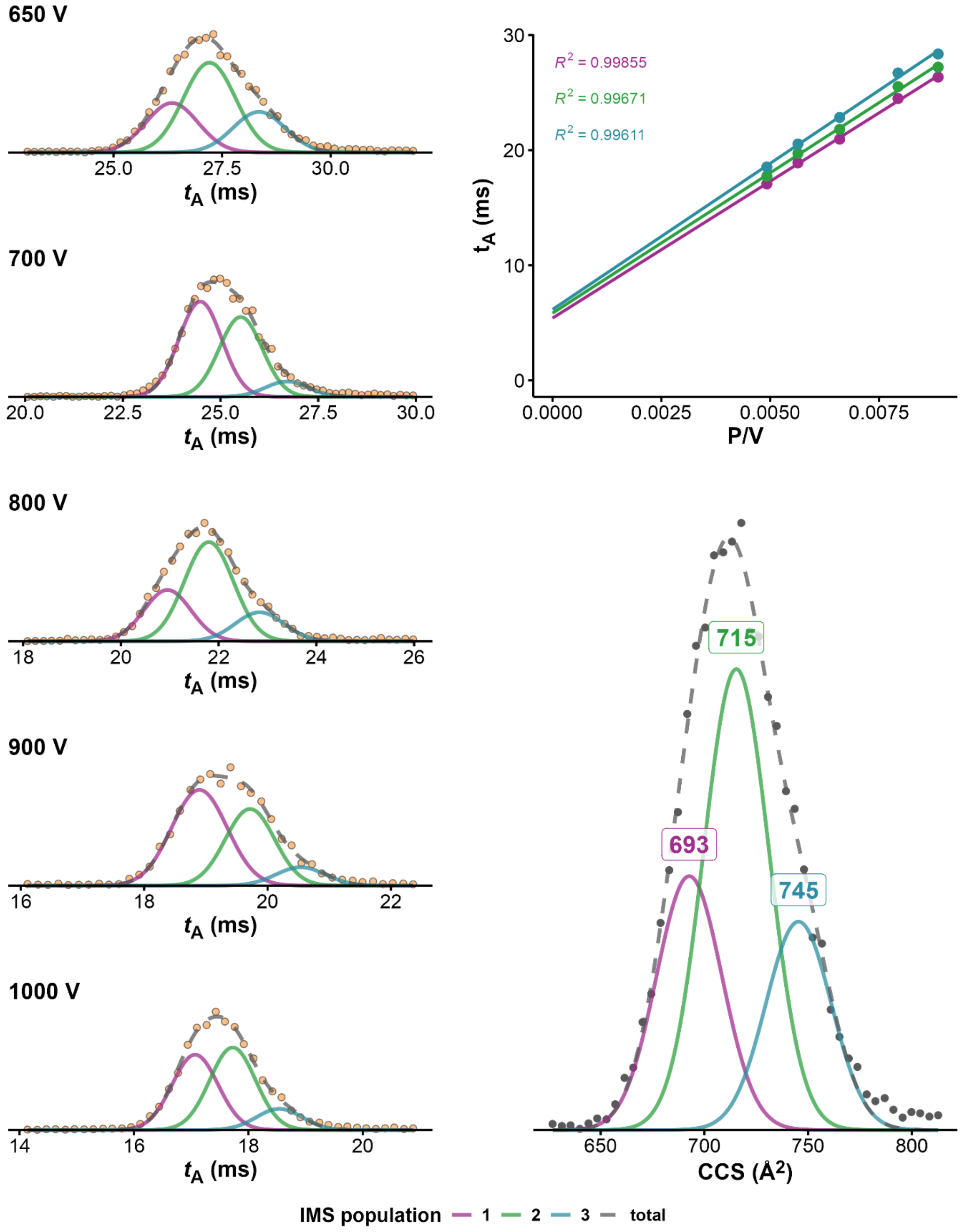


*Figure S34. CCS determination of 22AG·2K+ (100mM TMAA + 1mM KCl, fragmentor = 320V, z=4-). On the left panel ATDs at different drift voltages, on top right panel, $K_0$ determination from linear regression of arrival time at different voltages. Bottom right: CCS distribution obtained after conversion of the ATD at 650V*

**22GT•1K**

*100 mM TMAA + 1 mM KCl – fragmentor: 320 V*

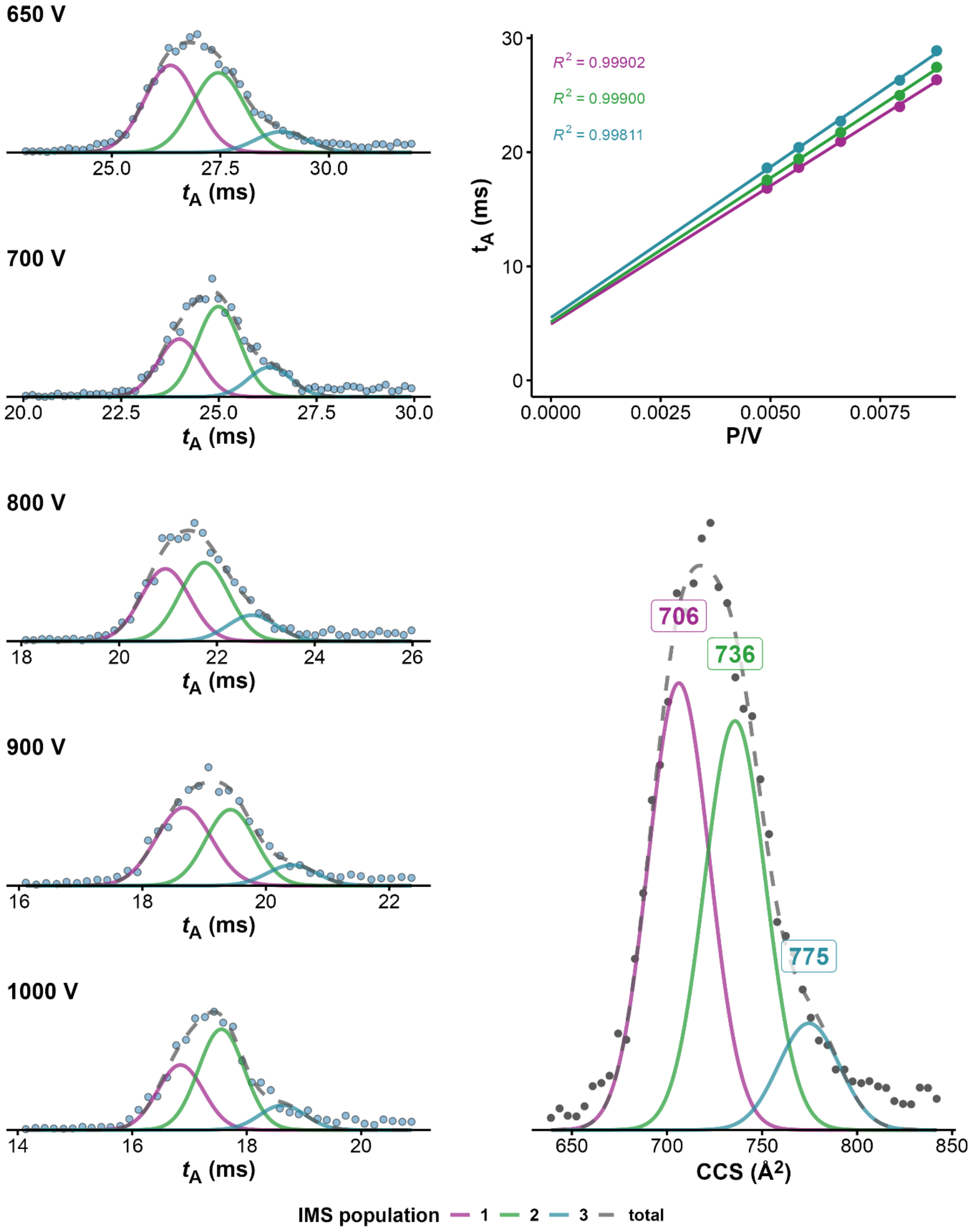


*Figure S35. CCS determination of 22GT·1K⁺ (100mM TMAA + 1mM KCl, fragmentor = 320V, z=4-). On the left panel ATDs at different drift voltages, on top right panel, $K_0$ determination from linear regression of arrival time at different voltages. Bottom right: CCS distribution obtained after conversion of the ATD at 650V*

**22GT•2K**

*100 mM TMAA + 1 mM KCl – fragmentor: 320 V*

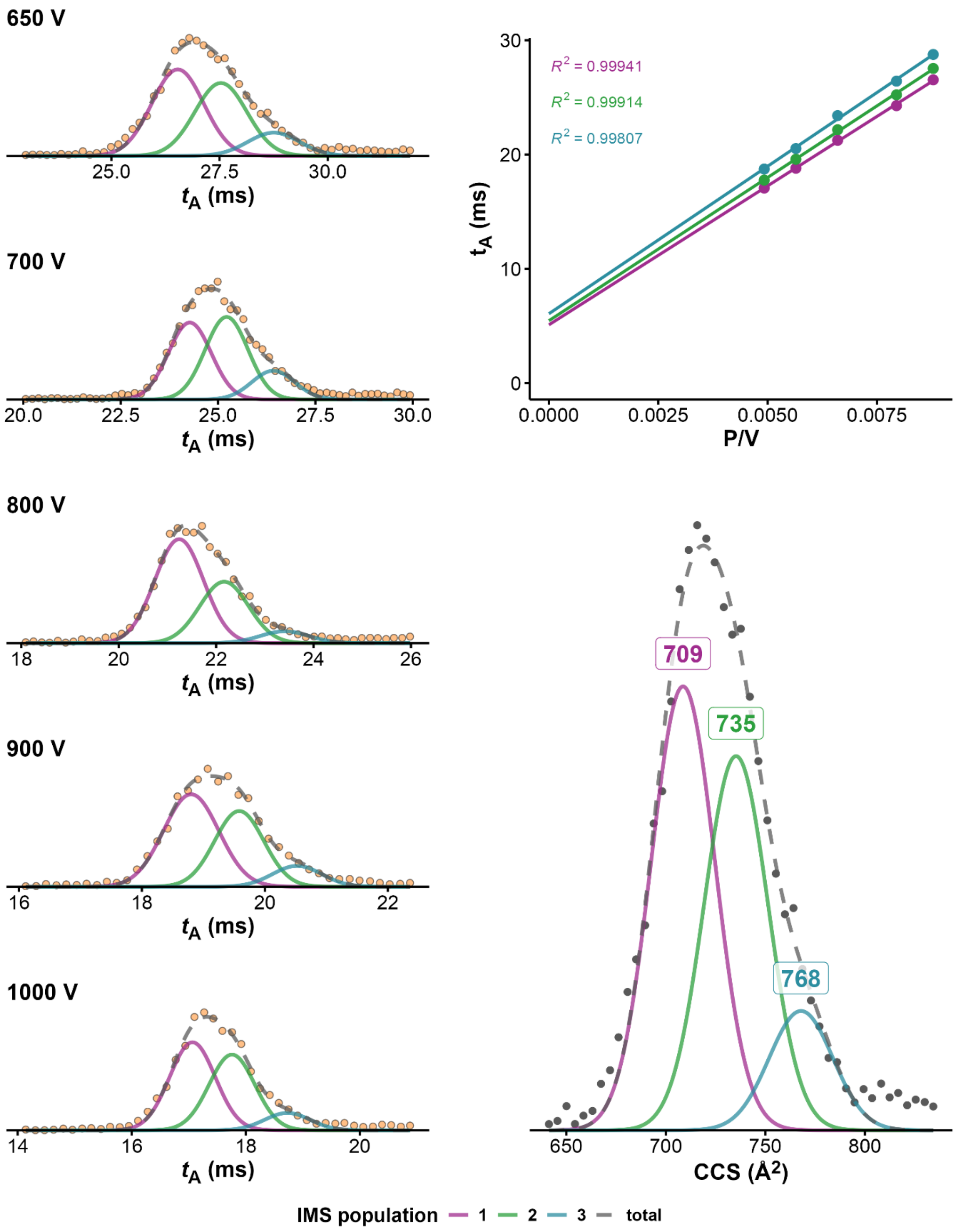


*Figure S36. CCS determination of 22GT·2K⁺ (100mM TMAA + 1mM KCl, fragmentor = 320V, z=4-). On the left panel ATDs at different drift voltages, on top right panel, $K_0$ determination from linear regression of arrival time at different voltages. Bottom right: CCS distribution obtained after conversion of the ATD at 650V*

**22CTA•1K**

*100 mM TMAA + 1 mM KCl – fragmentor: 320 V*

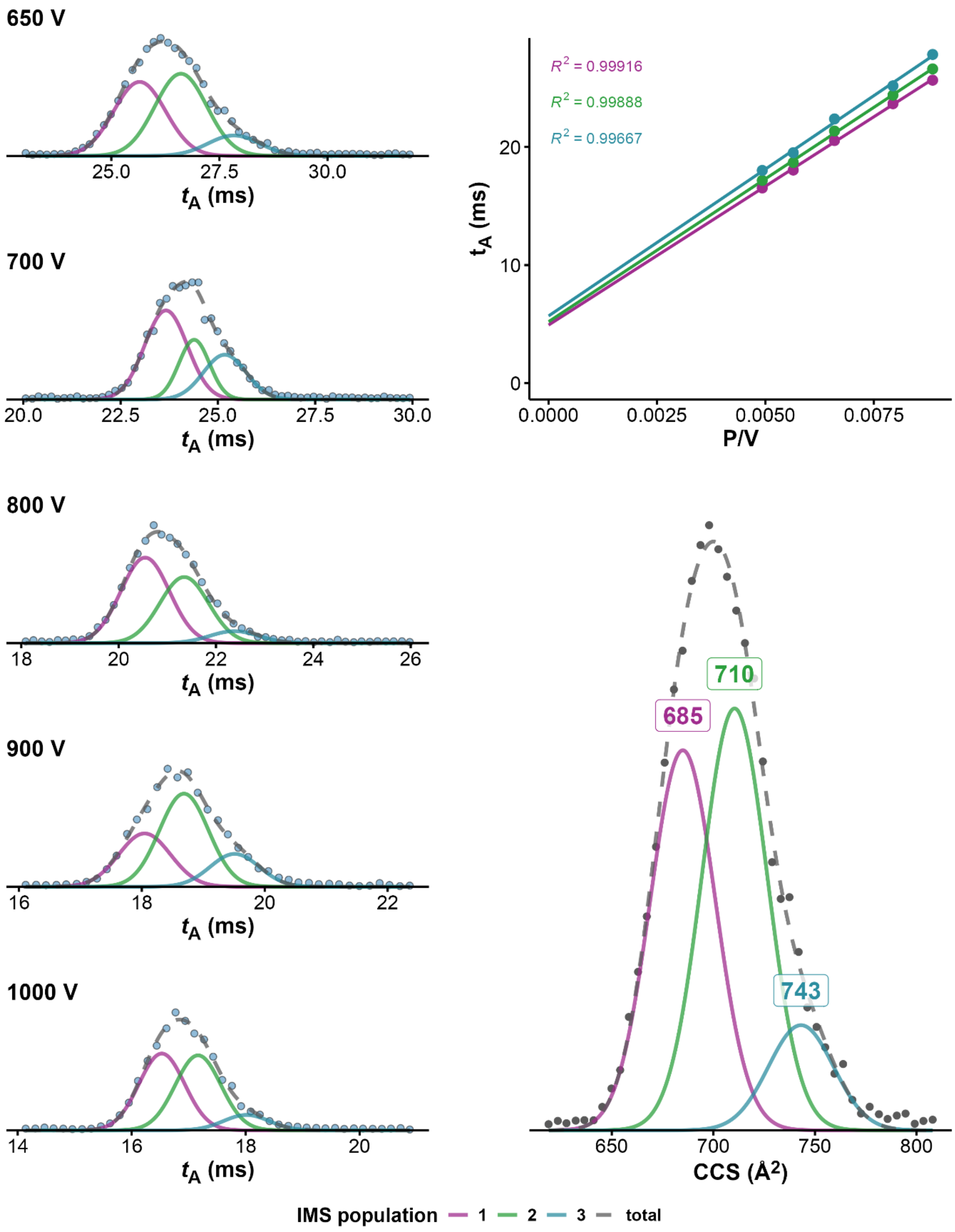


*Figure S37. CCS determination of 22CTA · 1K+ (100mM TMAA + 1mM KCl, fragmentor = 320V, z=4-). On the left panel ATDs at different drift voltages, on top right panel, $K_0$ determination from linear regression of arrival time at different voltages. Bottom right: CCS distribution obtained after conversion of the ATD at 650V*

**22CTA●2K**

*100 mM TMAA + 1 mM KCl – fragmentor: 320 V*

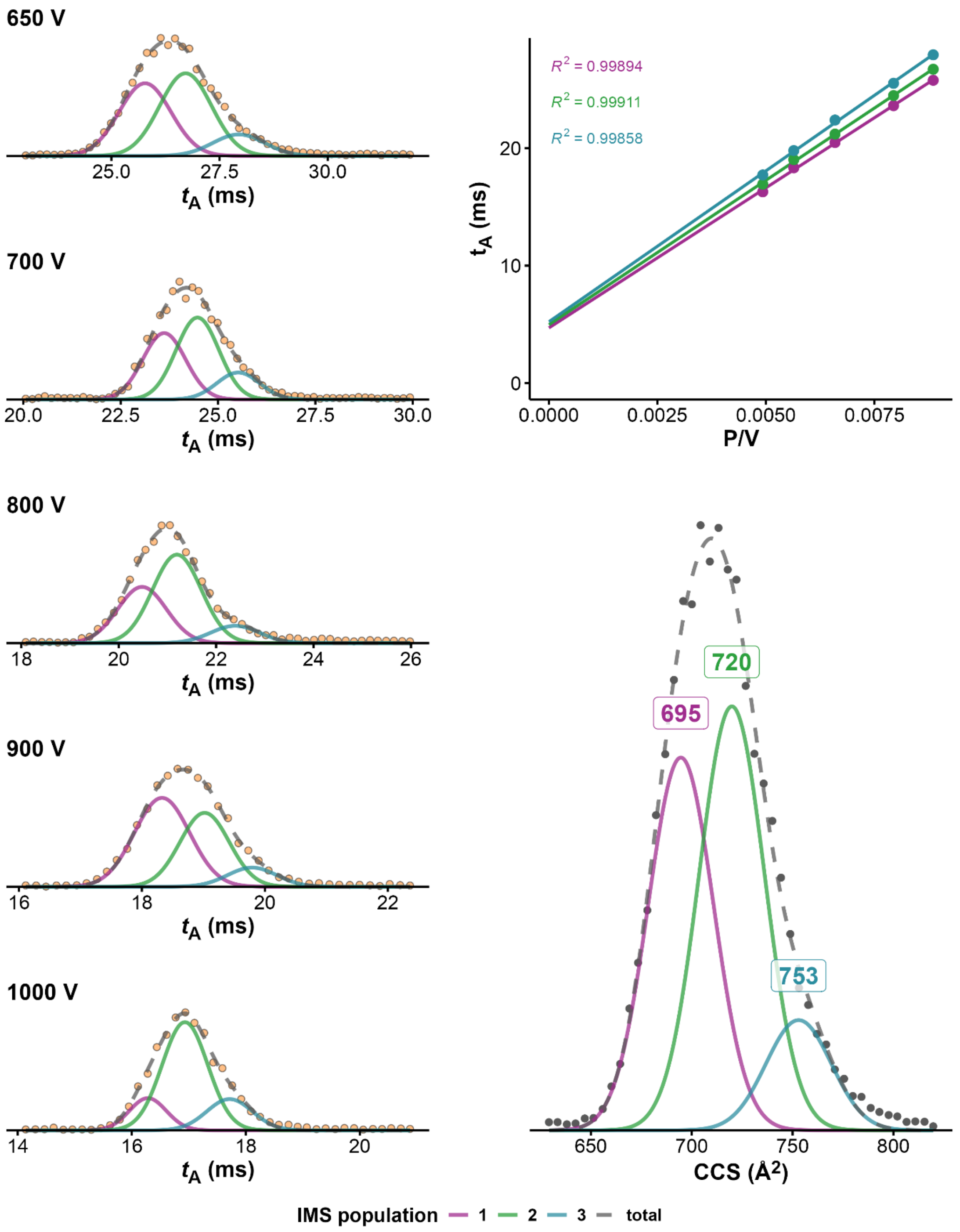


*Figure S38. CCS determination of 22CTA·2K⁺ (100mM TMAA + 1mM KCl, fragmentor = 320V, z=4-). On the left panel ATDs at different drift voltages, on top right panel, $K_0$ determination from linear regression of arrival time at different voltages. Bottom right: CCS distribution obtained after conversion of the ATD at 650V*

**23AG•2K**

*100 mM TMAA + 1 mM KCl – fragmentor: 320 V*

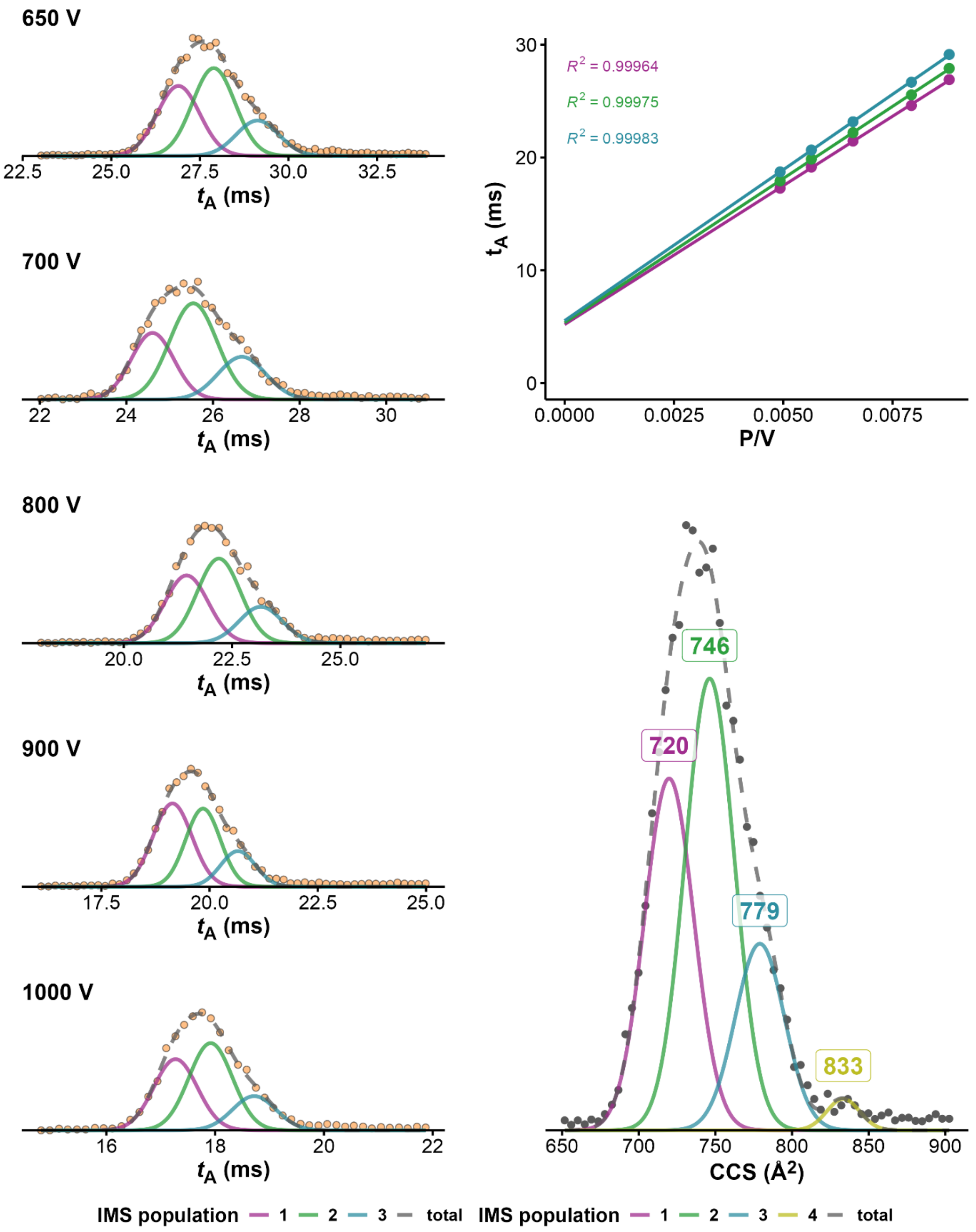


*Figure S39. CCS determination of 23AG·2K+ (100mM TMAA + 1mM KCl, fragmentor = 320V, z=4-). On the left panel ATDs at different drift voltages, on top right panel, $K_0$ determination from linear regression of arrival time at different voltages. Bottom right: CCS distribution obtained after conversion of the ATD at 650V*

**23AG•2K**

*100 mM TMAA + 1 mM KCl – fragmentor: 370 V*

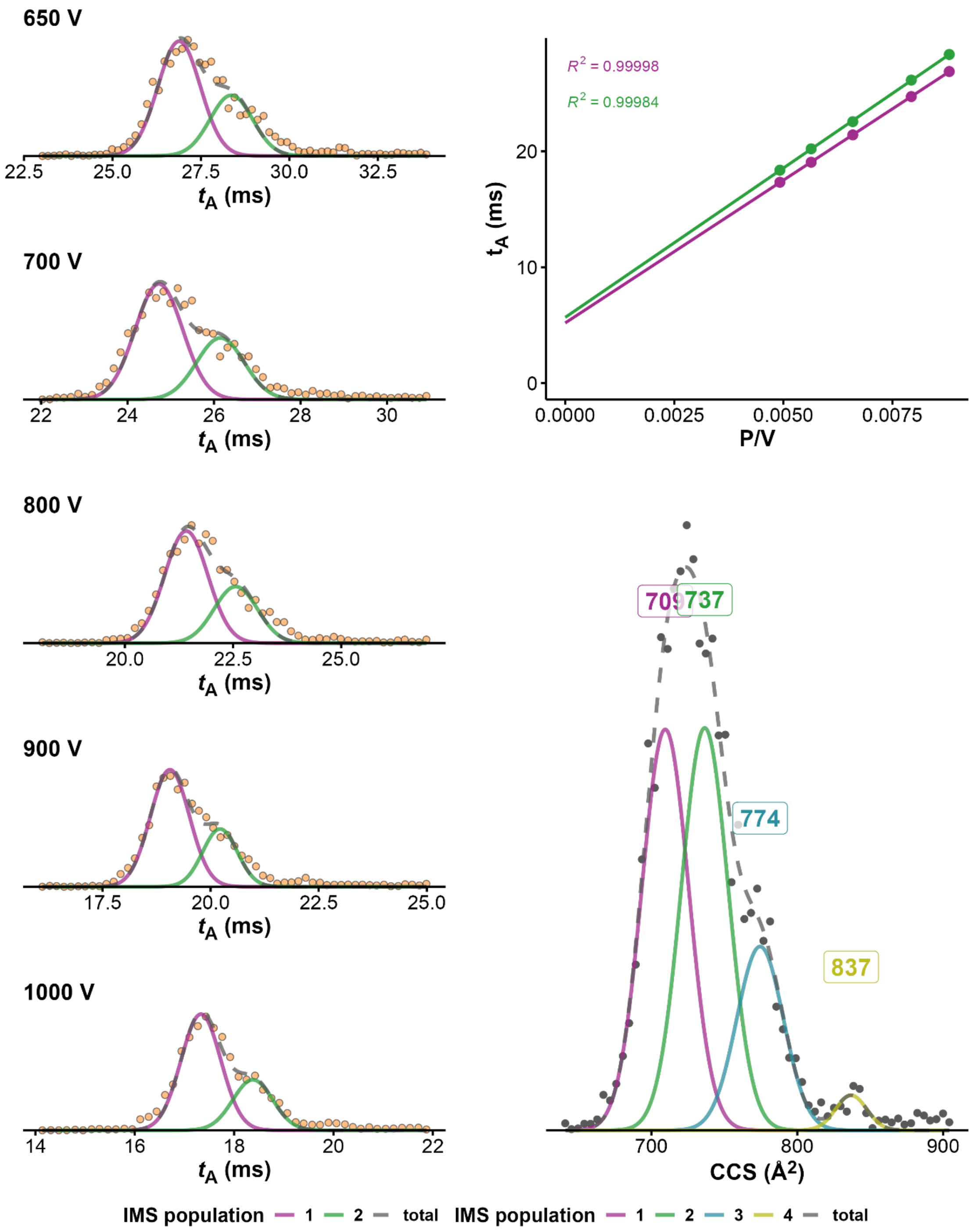


*Figure S40. CCS determination of 23AG·2K⁺ (100mM TMAA + 1mM KCl, fragmentor = 370V, z=4-). On the left panel ATDs at different drift voltages, on top right panel, $K_0$ determination from linear regression of arrival time at different voltages. Bottom right: CCS distribution obtained after conversion of the ATD at 650V*

**23AG_I14•1K**

*100 mM TMAA + 1 mM KCl – fragmentor: 320 V*

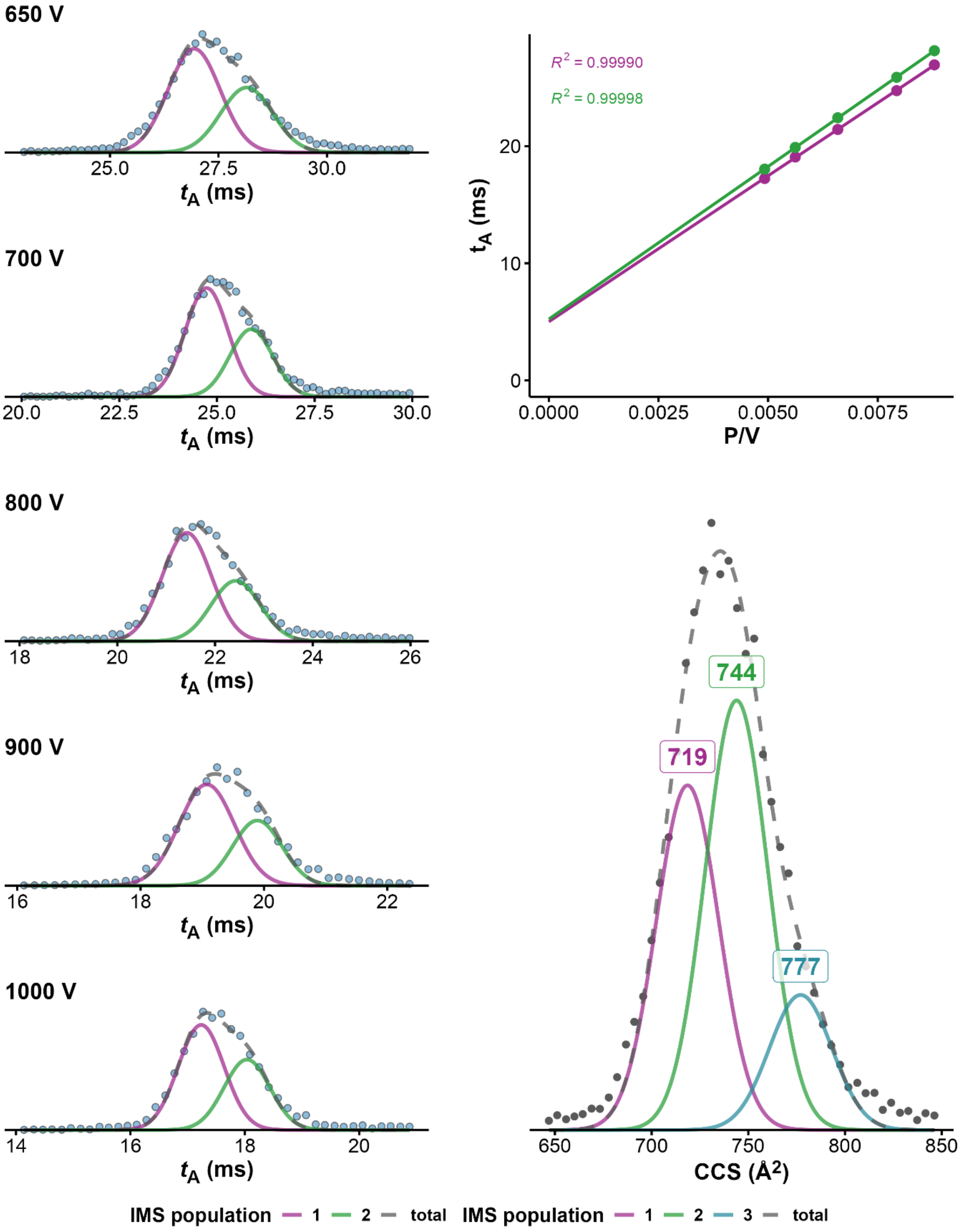


*Figure S41. CCS determination of 23AG_I14·1K⁺ (100mM TMAA + 1mM KCl, fragmentor = 320V, z=4-). On the left panel ATDs at different drift voltages, on top right panel, $K_0$ determination from linear regression of arrival time at different voltages. Bottom right: CCS distribution obtained after conversion of the ATD at 650V*

**23AG_I14•2K**
*100 mM TMAA + 1 mM KCl – fragmentor: 320 V*

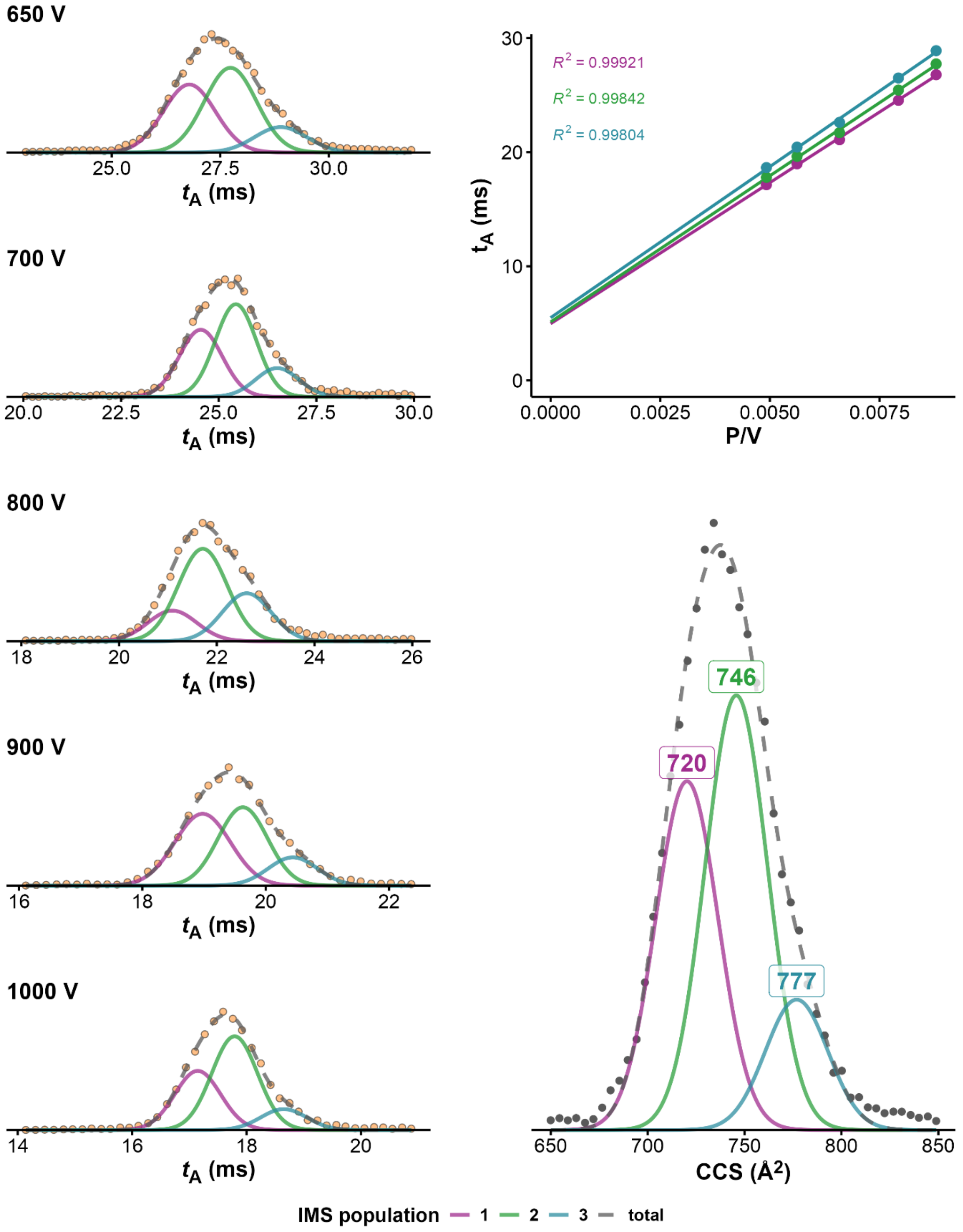


*Figure S42. CCS determination of 23AG_I14·2K⁺ (100mM TMAA + 1mM KCl, fragmentor = 320V, z=4-). On the left panel ATDs at different drift voltages, on top right panel, $K_0$ determination from linear regression of arrival time at different voltages. Bottom right: CCS distribution obtained after conversion of the ATD at 650V*

**23AG_I14●1K**

*100 mM TMAA + 1 mM KCl – fragmentor: 370 V*

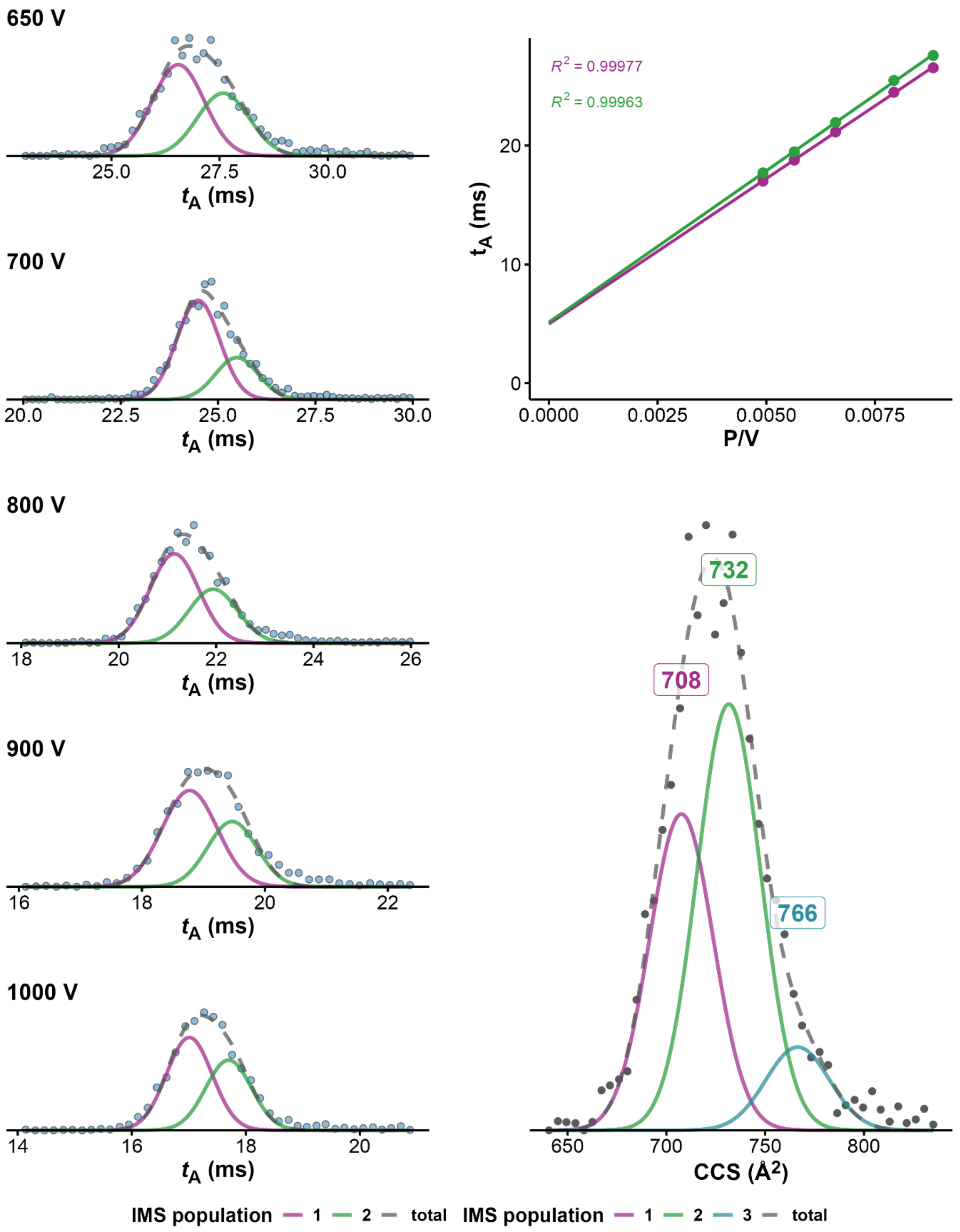


*Figure S43. CCS determination of 23AG_I14·1K⁺ (100mM TMAA + 1mM KCl, fragmentor = 370V, z=4-). On the left panel ATDs at different drift voltages, on top right panel, $K_0$ determination from linear regression of arrival time at different voltages. Bottom right: CCS distribution obtained after conversion of the ATD at 650V.*

**23AG_I14•2K**

*100 mM TMAA + 1 mM KCl – fragmentor: 370 V*

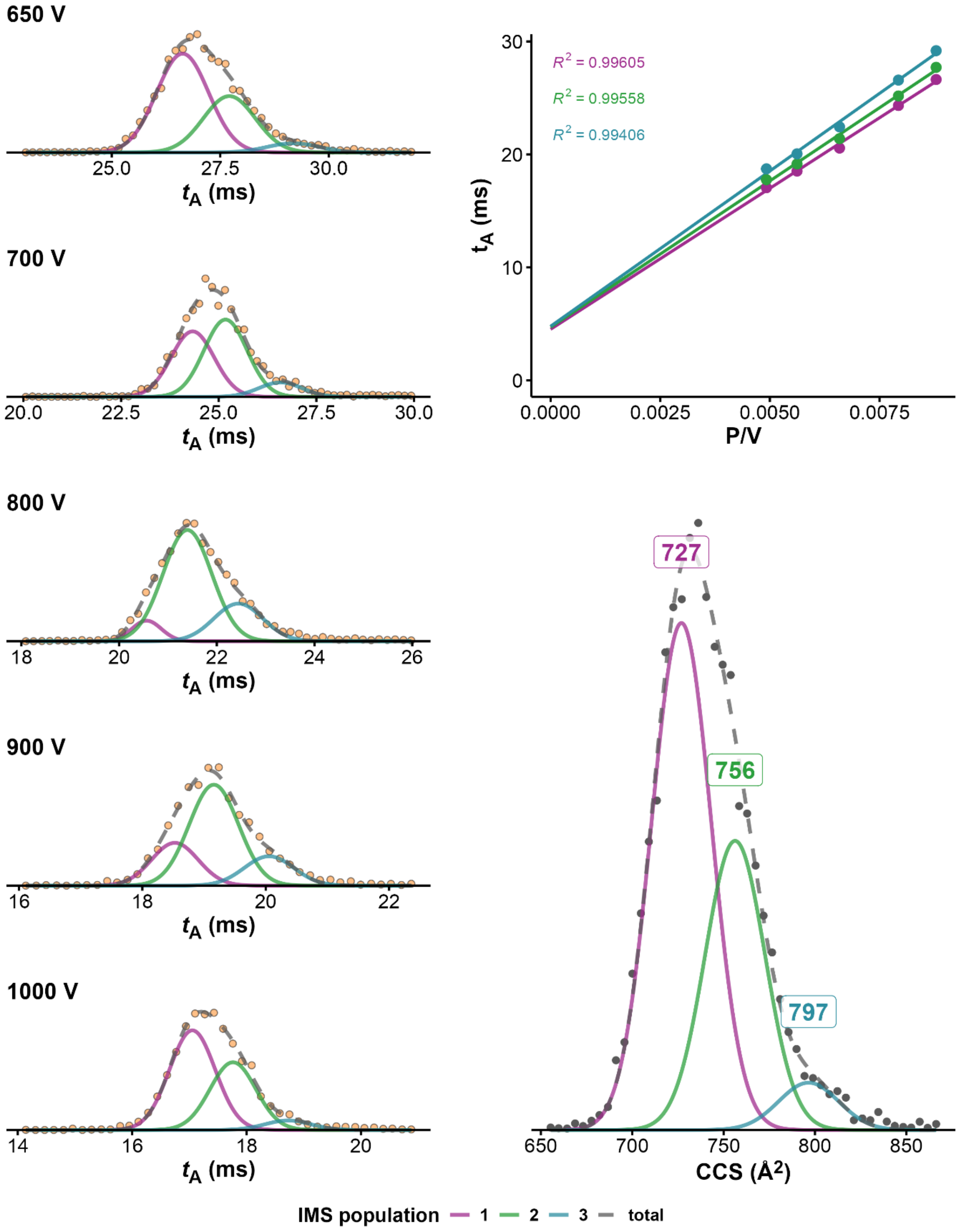


*Figure S44. CCS determination of 23AG_I14 · 2K⁺ (100mM TMAA + 1mM KCl, fragmentor = 370V, z=4-). On the left panel ATDs at different drift voltages, on top right panel, $K_0$ determination from linear regression of arrival time at different voltages. Bottom right: CCS distribution obtained after conversion of the ATD at 650V*

**26CEB•2K**

*100 mM TMAA + 1 mM KCl – fragmentor: 320 – z = 4- V*

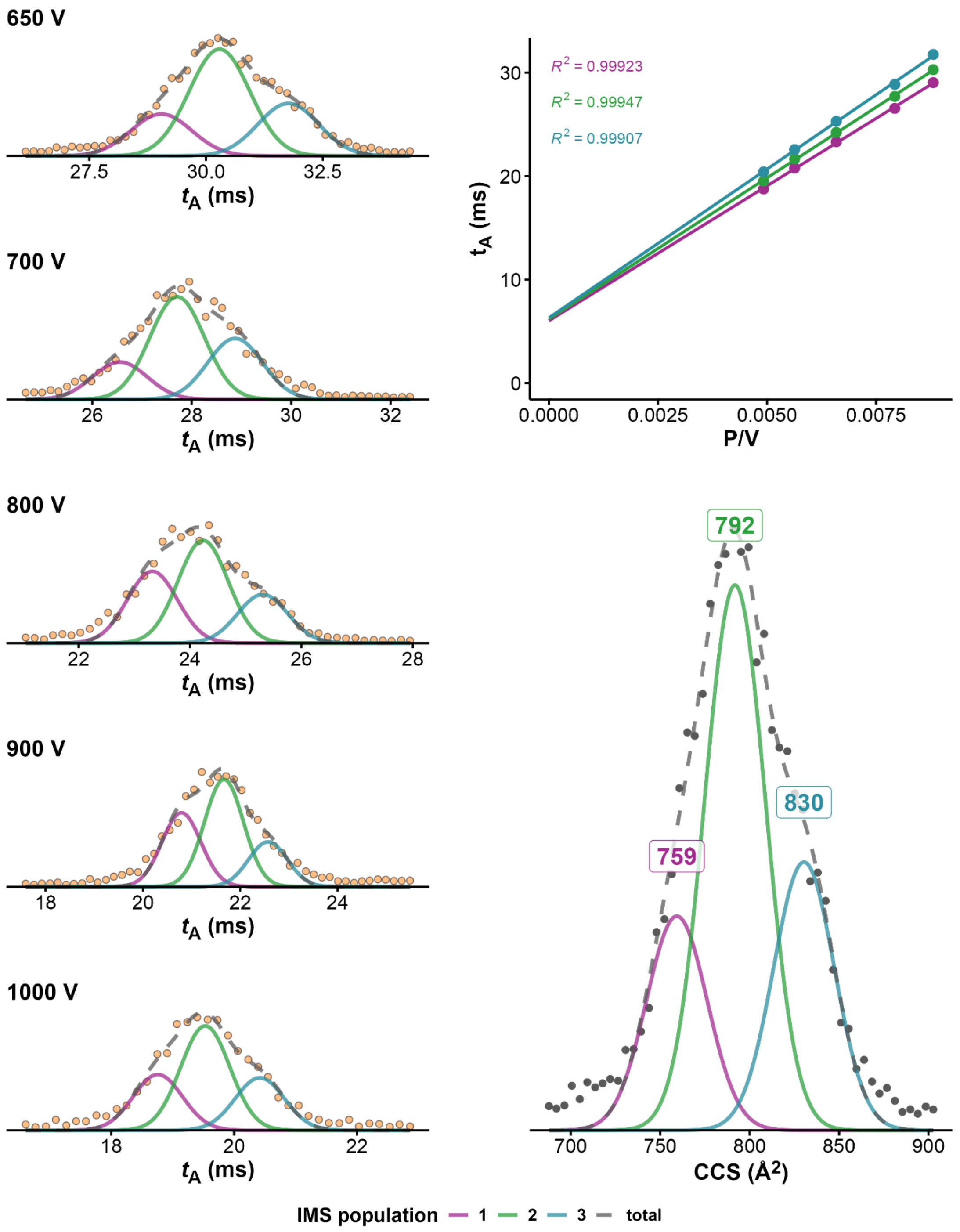


*Figure S45. CCS determination of 26CEB·2K⁺ (100mM TMAA + 1mM KCl, fragmentor = 320V, z=4-). On the left panel ATDs at different drift voltages, on top right panel, $K_0$ determination from linear regression of arrival time at different voltages. Bottom right: CCS distribution obtained after conversion of the ATD at 650V*

**26CEB•2K**

*100 mM TMAA + 1 mM KCl – fragmentor: 320 – z = 5- V*

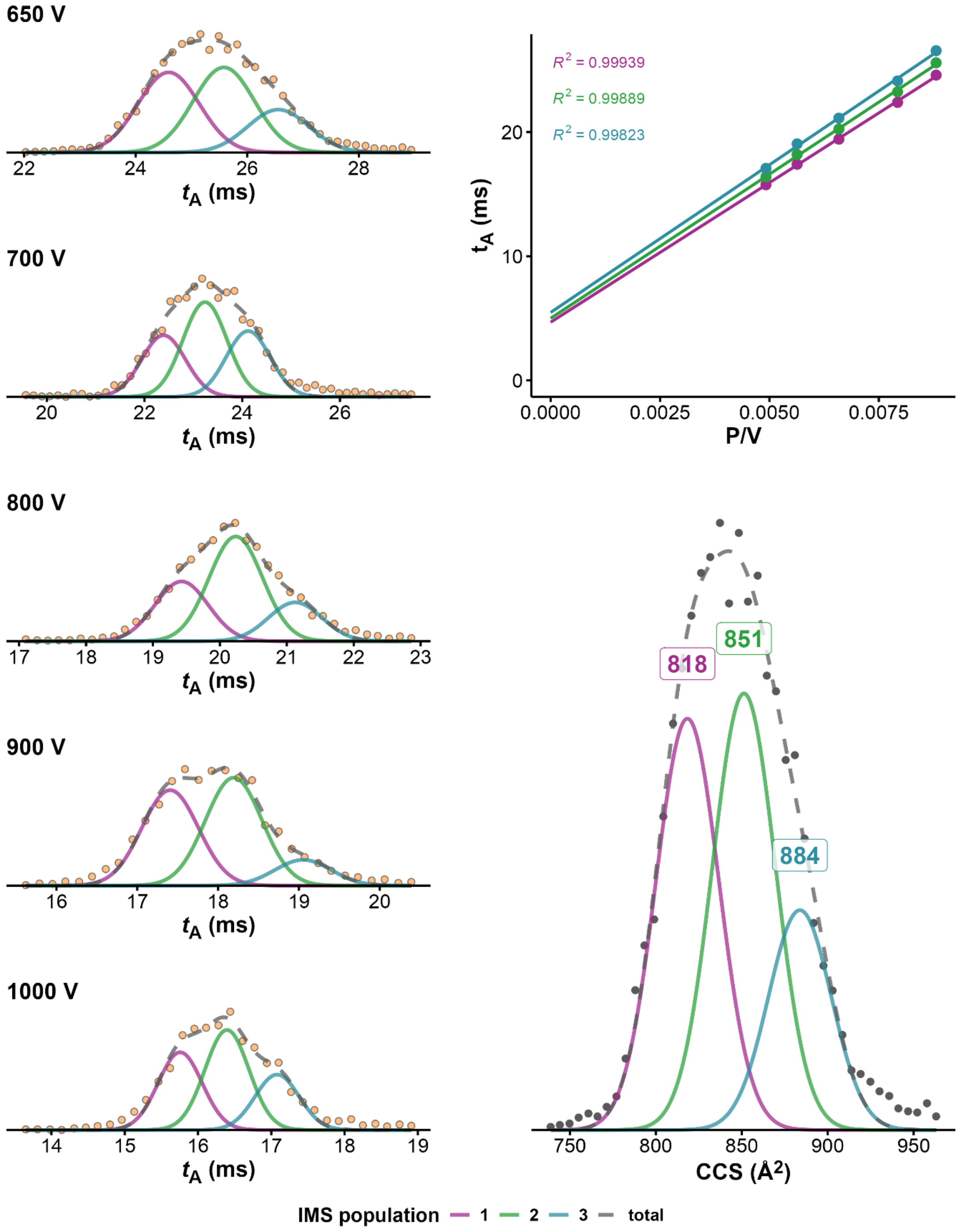


*Figure S46. CCS determination of 26CEB·2K⁺ (100mM TMAA + 1mM KCl, fragmentor = 320V, z=5-). On the left panel ATDs at different drift voltages, on top right panel, $K_0$ determination from linear regression of arrival time at different voltages. Bottom right: CCS distribution obtained after conversion of the ATD at 650V*

**222T•2K**

*100 mM TMAA + 1 mM KCl – fragmentor: 320 V*

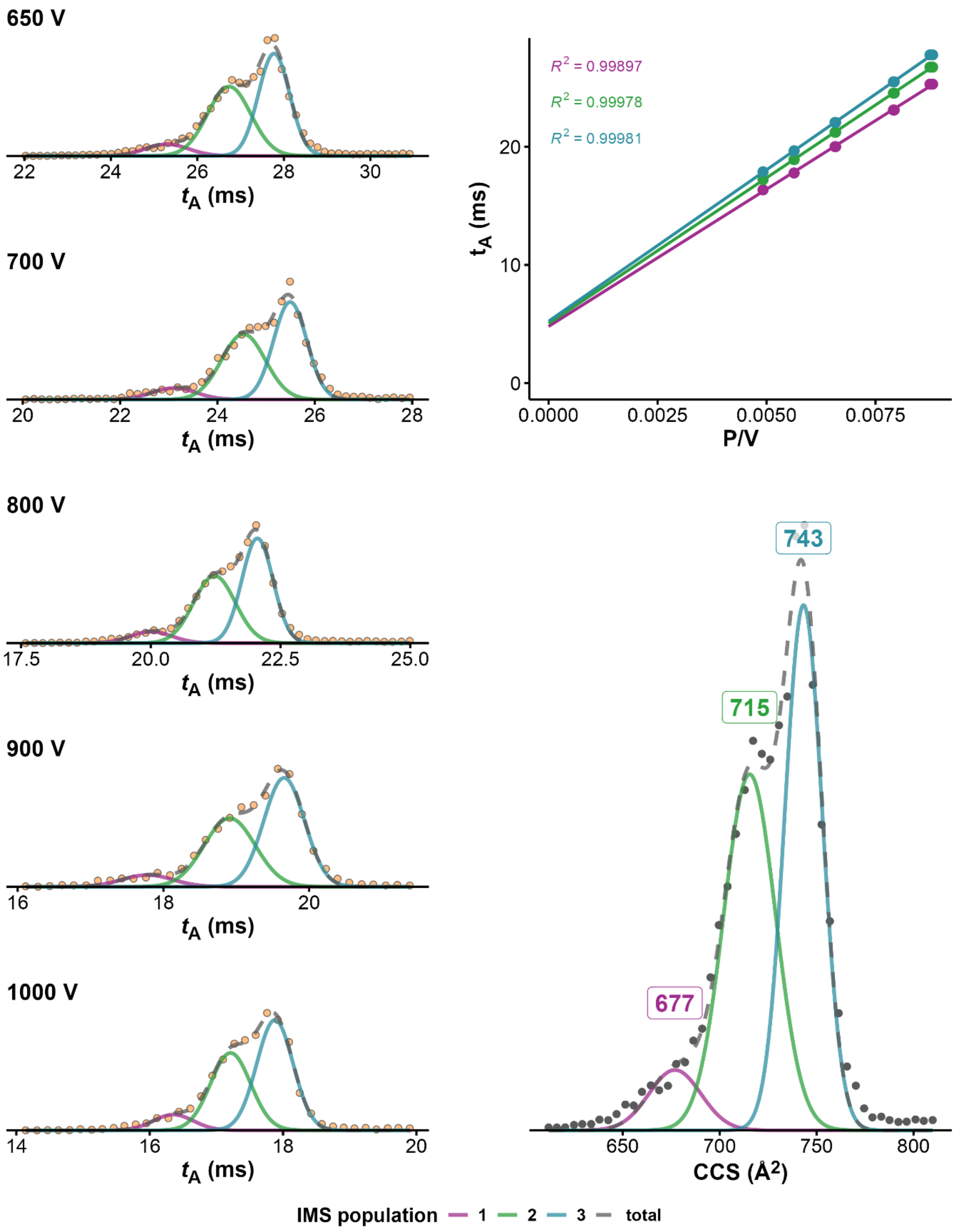


*Figure S47. CCS determination of 222T·2K⁺ (100mM TMAA + 1mM KCl, fragmentor = 320V, z=4-). On the left panel ATDs at different drift voltages, on top right panel, $K_0$ determination from linear regression of arrival time at different voltages. Bottom right: CCS distribution obtained after conversion of the ATD at 650V*

**T95TT•2K**

*100 mM TMAA + 0.5 mM KCl – fragmentor: 320 V*

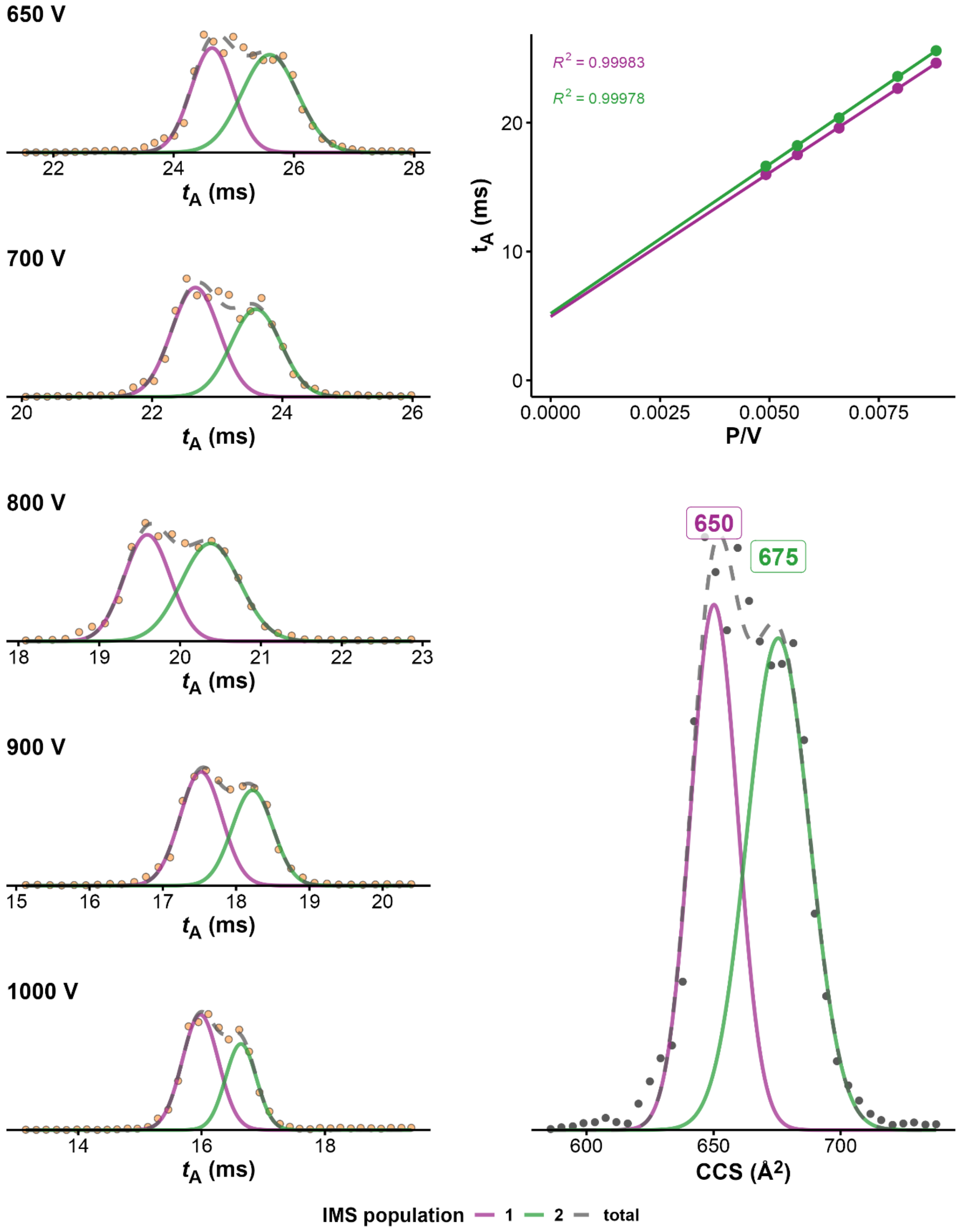


*Figure S48. CCS determination of T95TT·2K⁺ (100mM TMAA + 0.5mM KCl, fragmentor = 320V, z=4-). On the left panel ATDs at different drift voltages, on top right panel, $K_0$ determination from linear regression of arrival time at different voltages. Bottom right: CCS distribution obtained after conversion of the ATD at 650V*

**T30177TT•2K**

*100 mM TMAA + 0.5 mM KCl – fragmentor: 320 V*

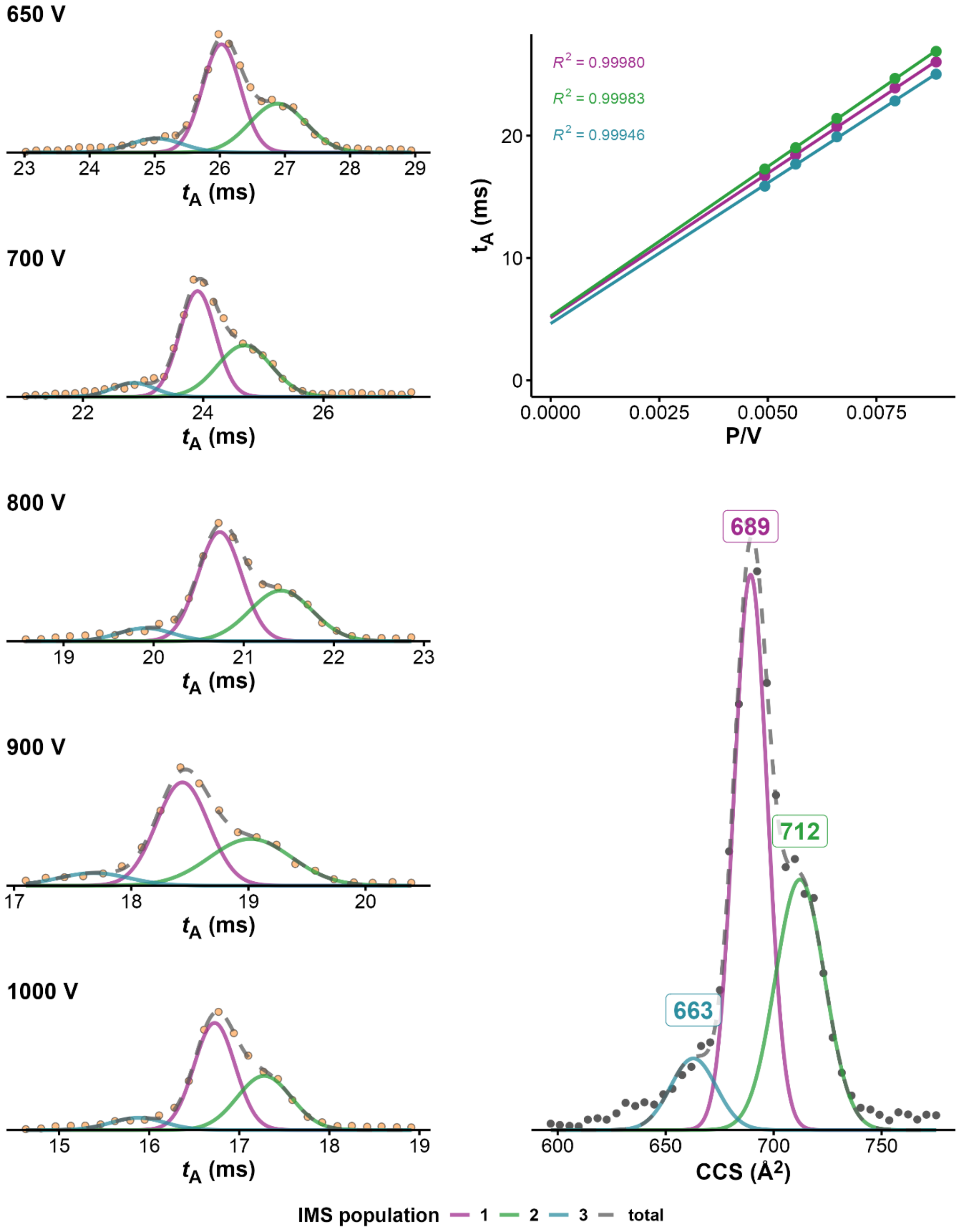


*Figure S49. CCS determination of T30177TT · 2K$^+$ (100mM TMAA + 0.5mM KCl, fragmentor = 320V, z=4-). On the left panel ATDs at different drift voltages, on top right panel, $K_0$ determination from linear regression of arrival time at different voltages. Bottom right: CCS distribution obtained after conversion of the ATD at 650V*

**2KPR•2K**

*100 mM TMAA + 1 mM KCl – fragmentor: 320 V*

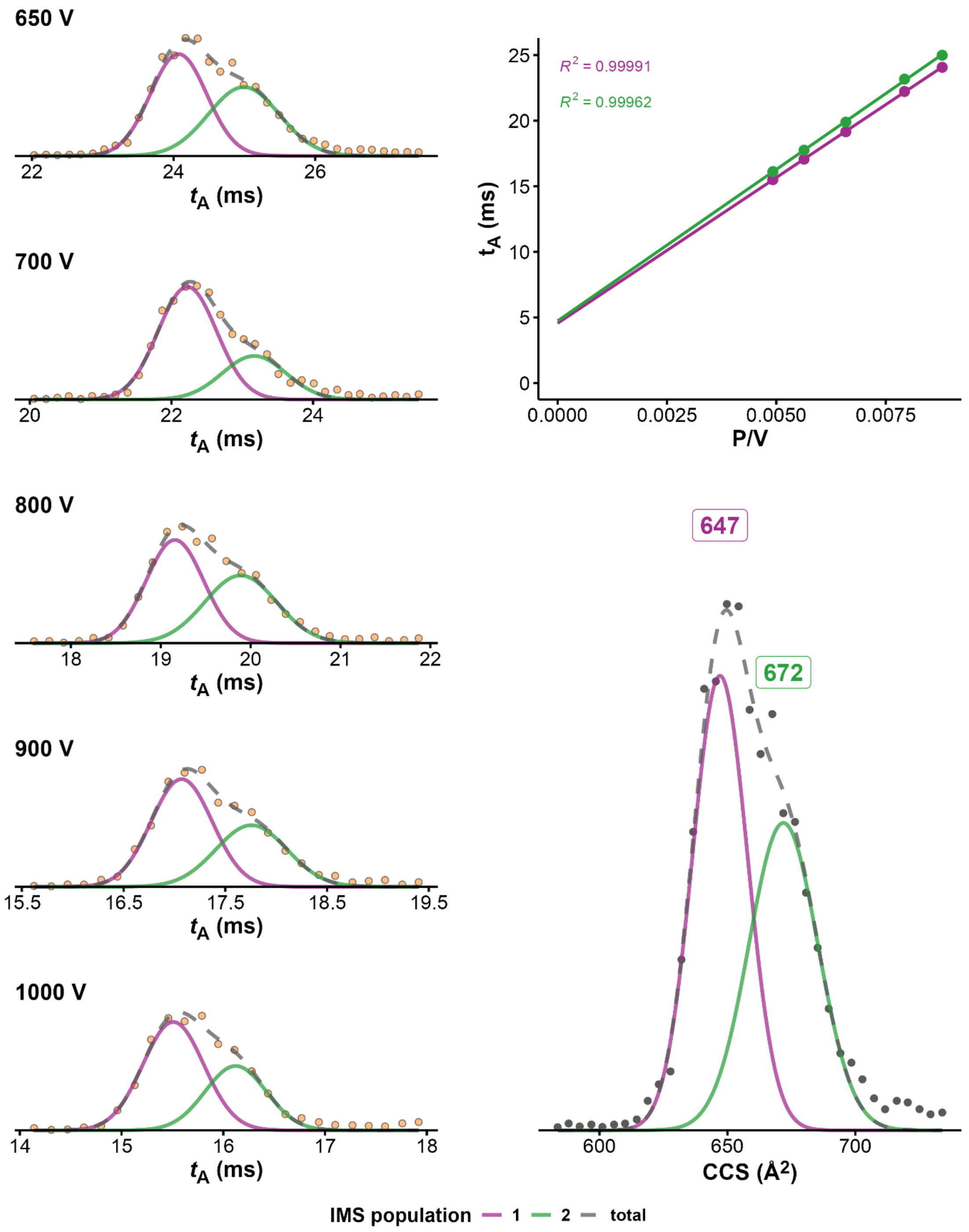


*Figure S50. CCS determination of 2KPR·2K⁺ (100mM TMAA + 1mM KCl, fragmentor = 320V, z=4-). On the left panel ATDs at different drift voltages, on top right panel, $K_0$ determination from linear regression of arrival time at different voltages. Bottom right: CCS distribution obtained after conversion of the ATD at 650V*

**TG4T•3K**

*100 mM TMAA + 1 mM KCl – fragmentor: 320 V*

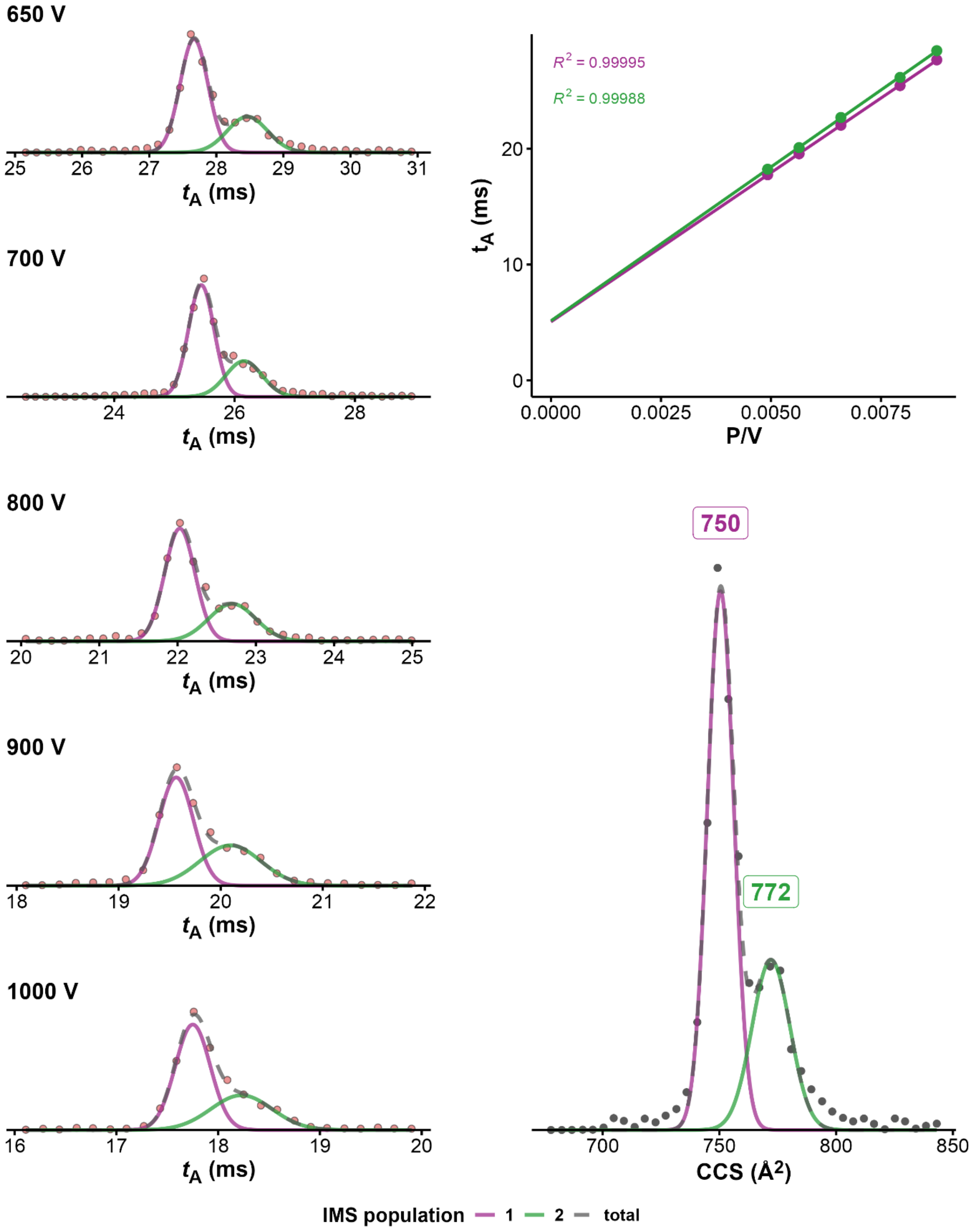


*Figure S51. CCS determination of [TG4T]$_4$ · 3K$^+$ (100mM TMAA + 1mM KCl, fragmentor = 320V, z=4-). On the left panel ATDs at different drift voltages, on top right panel, $K_0$ determination from linear regression of arrival time at different voltages. Bottom right: CCS distribution obtained after conversion of the ATD at 650V*

**TG4T•3NH$_4^+$**

*100 mM NH4OAc – fragmentor: 320 V – z = 4-*

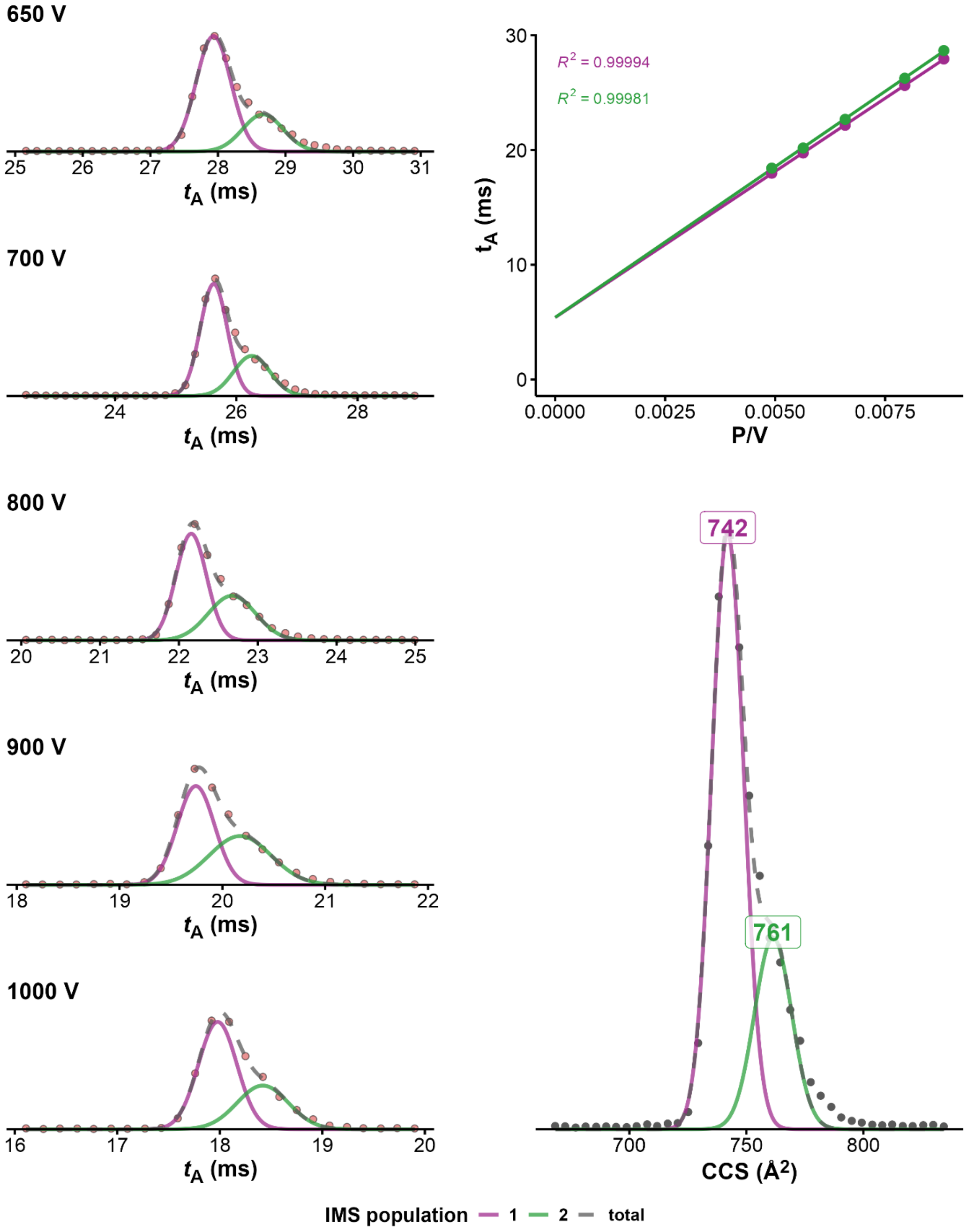


*Figure S52. CCS determination of [TG4T]$_4$ · 3NH$_4^+$+ (100mM NH$_4$OAc, fragmentor = 320V, z=4-). On the left panel ATDs at different drift voltages, on top right panel, $K_0$ determination from linear regression of arrival time at different voltages. Bottom right: CCS distribution obtained after conversion of the ATD at 650V*

**ckit87up•2K**

*100 mM TMAA + 1 mM KCl – fragmentor: 320 V*

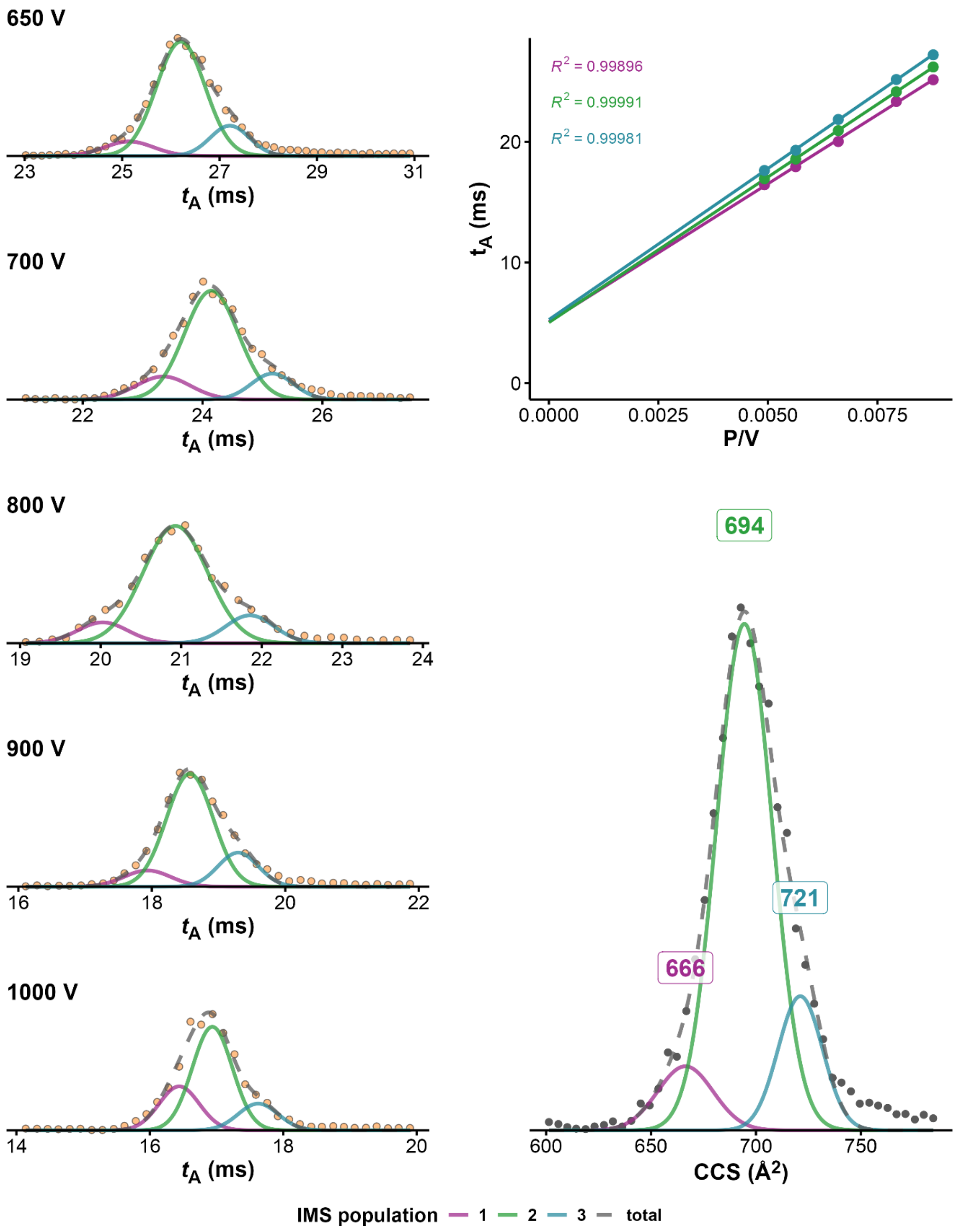


*Figure S53. CCS determination of ckit87up · 2K+ (100mM TMAA + 1mM KCl, fragmentor = 320V, z=4-). On the left panel ATDs at different drift voltages, on top right panel, $K_0$ determination from linear regression of arrival time at different voltages. Bottom right: CCS distribution obtained after conversion of the ATD at 650V*

**BCL2•2K**

*100 mM TMAA + 1 mM KCl – fragmentor: 320 V*

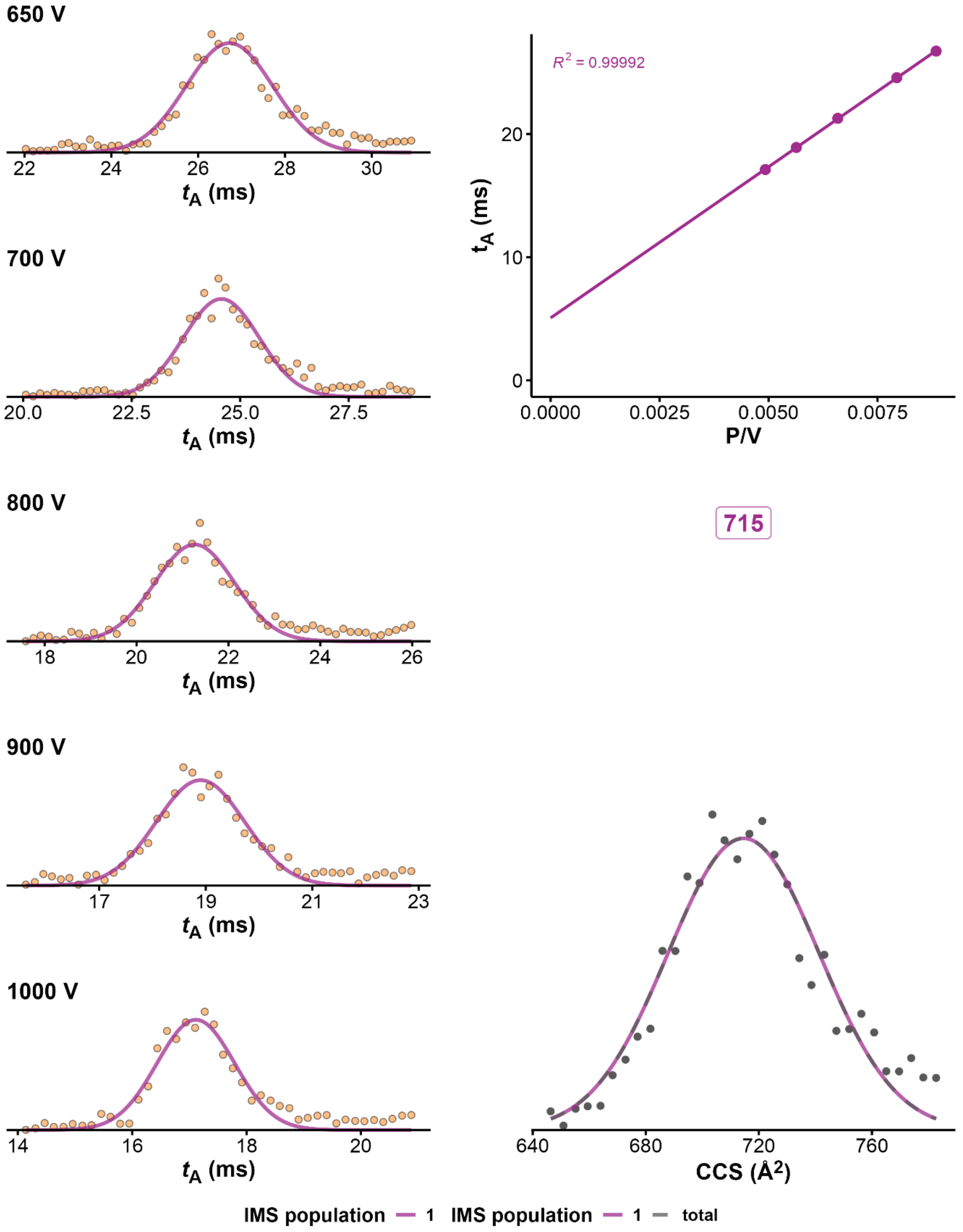


*Figure S54. CCS determination of BCL2·2K⁺ (100mM TMAA + 1mM KCl, fragmentor = 320V, z=4-). On the left panel ATDs at different drift voltages, on top right panel, $K_0$ determination from linear regression of arrival time at different voltages. Bottom right: CCS distribution obtained after conversion of the ATD at 650V*

### CIU heatmaps

*21G*

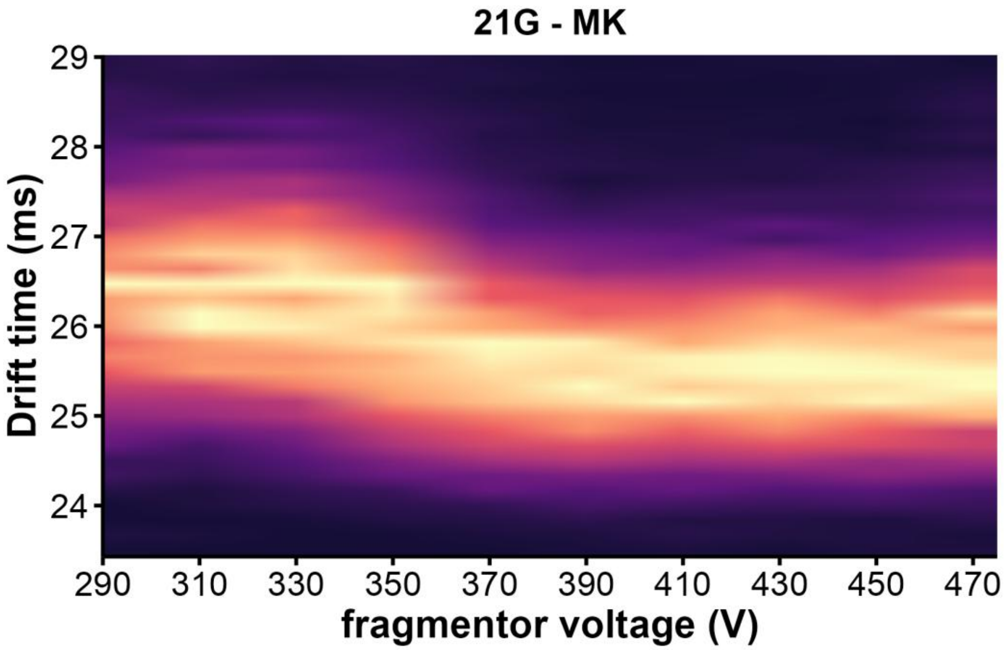


*Figure S55. CIU heatmap of 21G · 1K$^{+}$ (z= 4 -) in 100mM TMAA + 1 mM KCl*

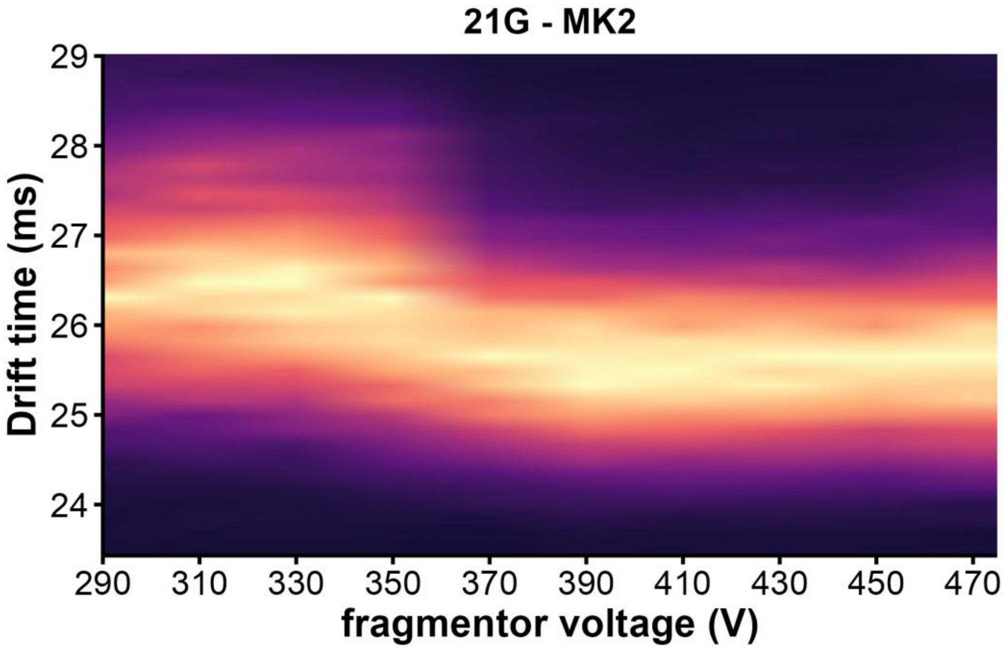


*Figure S56. CIU heatmap of 21G · 2K$^{+}$ (z= 4 -) in 100mM TMAA + 1 mM KCl*

*5YEY*

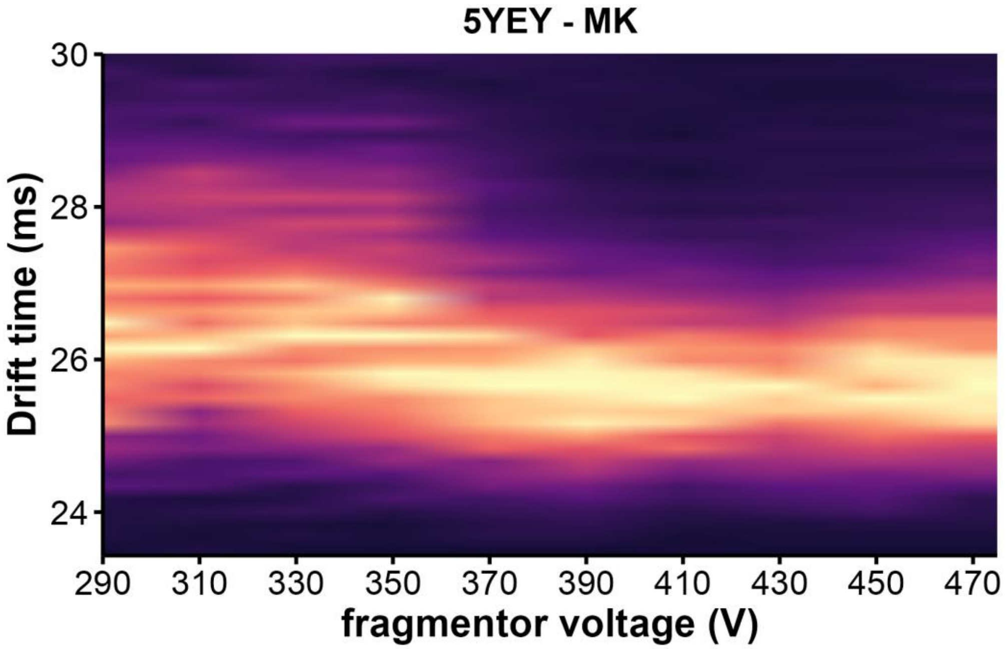


*Figure S57. CIU heatmap of 5YEY · 1K⁺ (z= 4 -) in 100mM TMAA + 1 mM KCl*

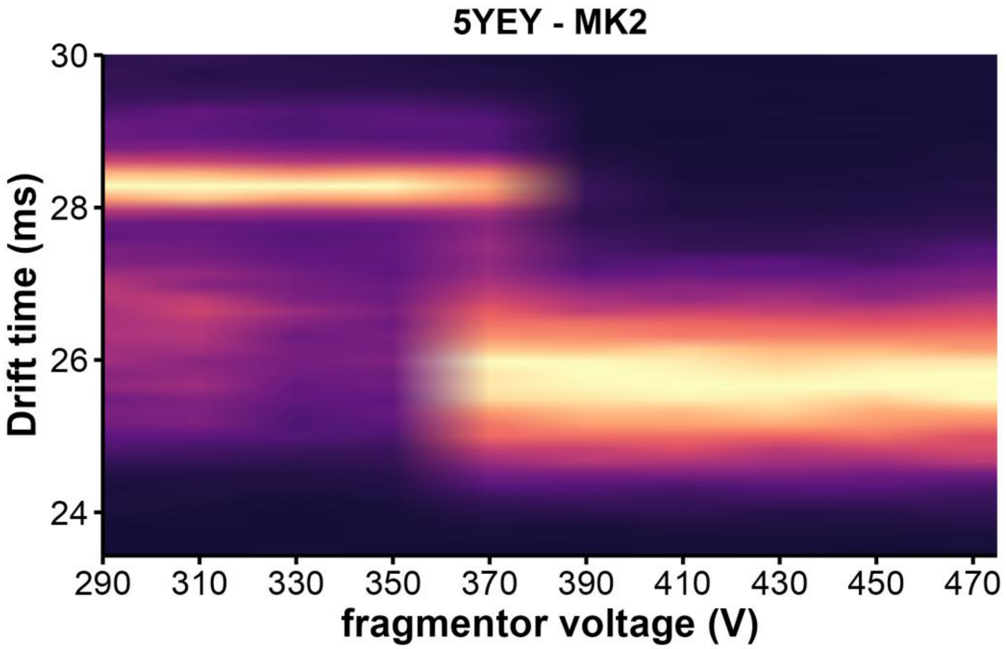


*Figure S58. CIU heatmap of 5YEY · 2K⁺ (z= 4 -) in 100mM TMAA + 1 mM KCl*

*22AG*

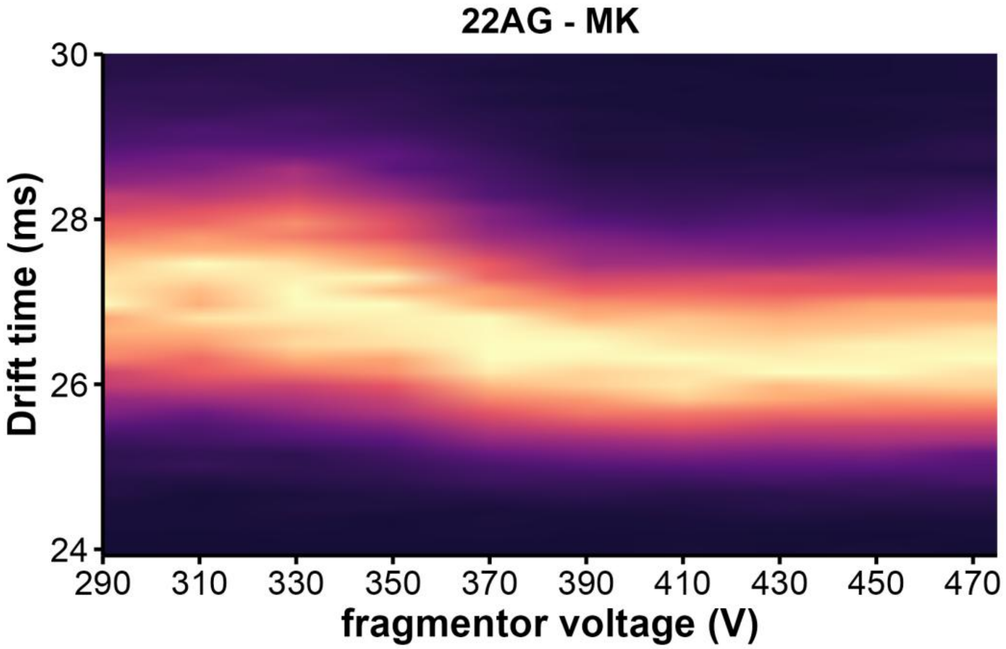


*Figure S59. CIU heatmap of 22AG · 1K⁺ (z= 4 -) in 100mM TMAA + 1 mM KCl*

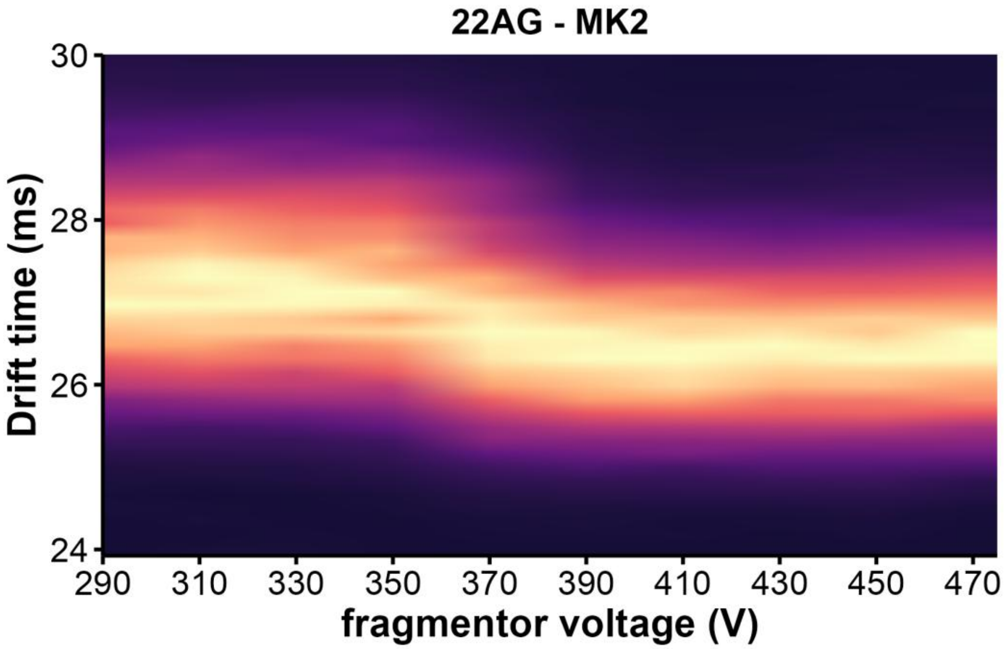


*Figure S60. CIU heatmap of 22AG · 2K⁺ (z= 4 -) in 100mM TMAA + 1 mM KCl*

*22GT*

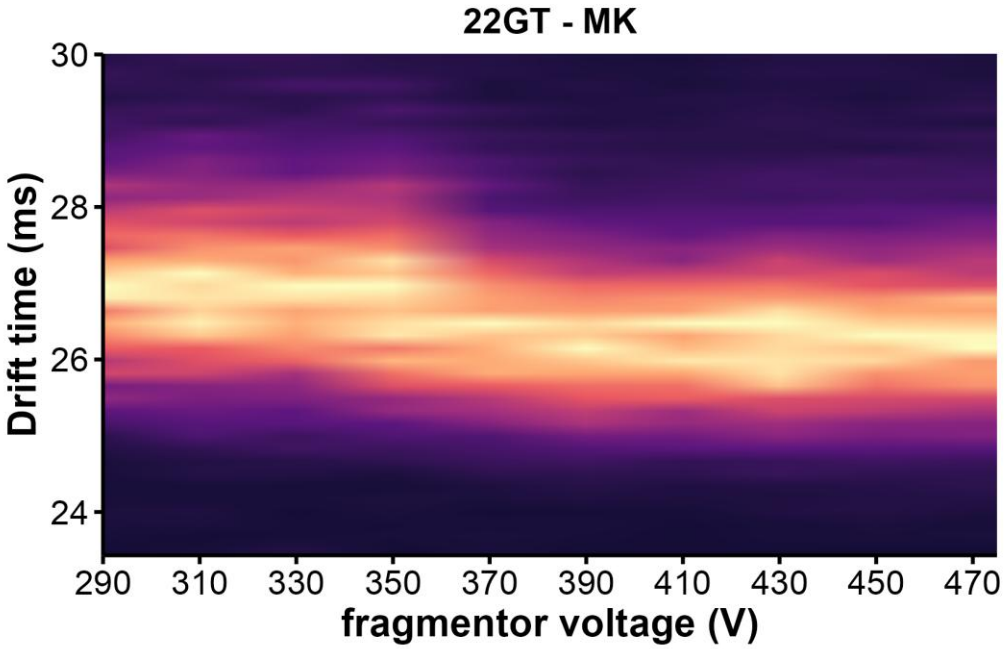


*Figure S61. CIU heatmap of 22GT· 1K⁺ (z= 4 -) in 100mM TMAA + 1 mM KCl*

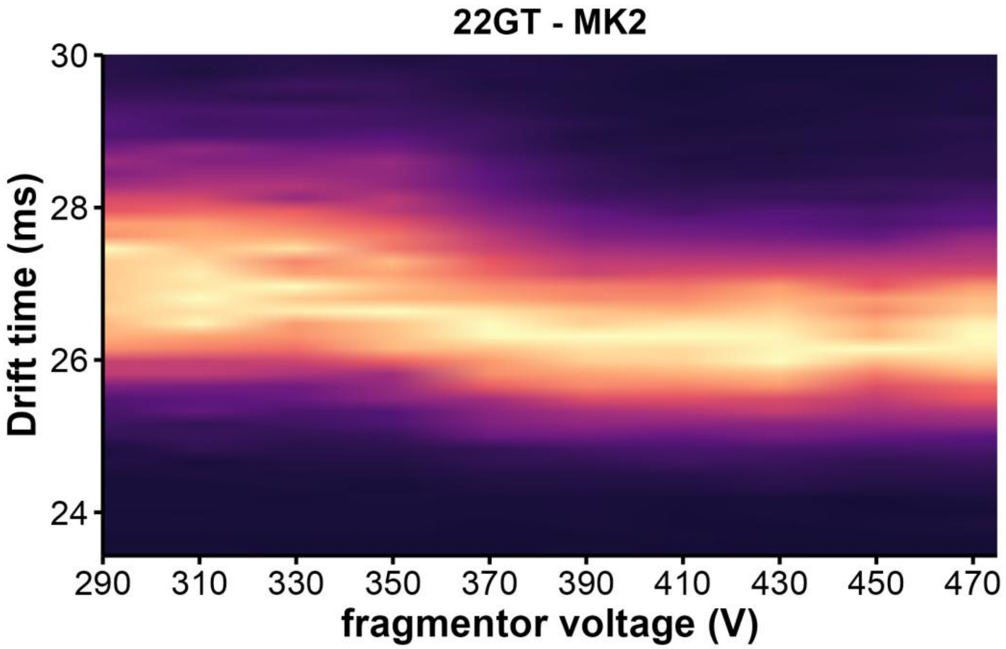


*Figure S62. CIU heatmap of 22GT· 2K⁺ (z= 4 -) in 100mM TMAA + 1 mM KCl*

*22GT_18T*

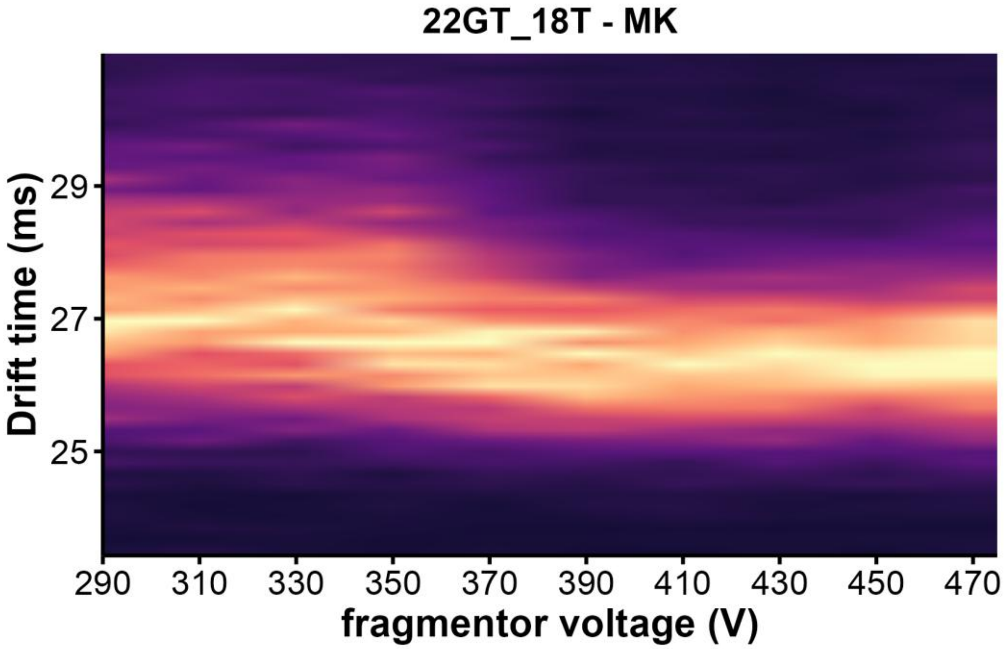


*Figure S63. CIU heatmap of 22GT_18T · 1K⁺ (z= 4 -) in 100mM TMAA + 1 mM KCl*

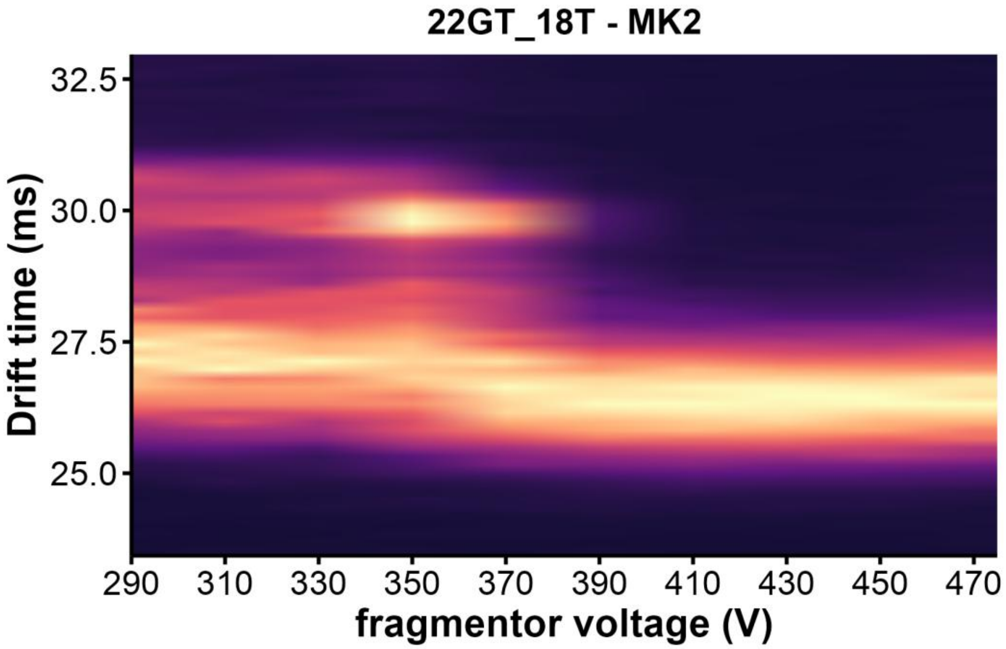


*Figure S64. CIU heatmap of 22GT_18T · 2K⁺ (z= 4 -) in 100mM TMAA + 1 mM KCl*

*22CTA*

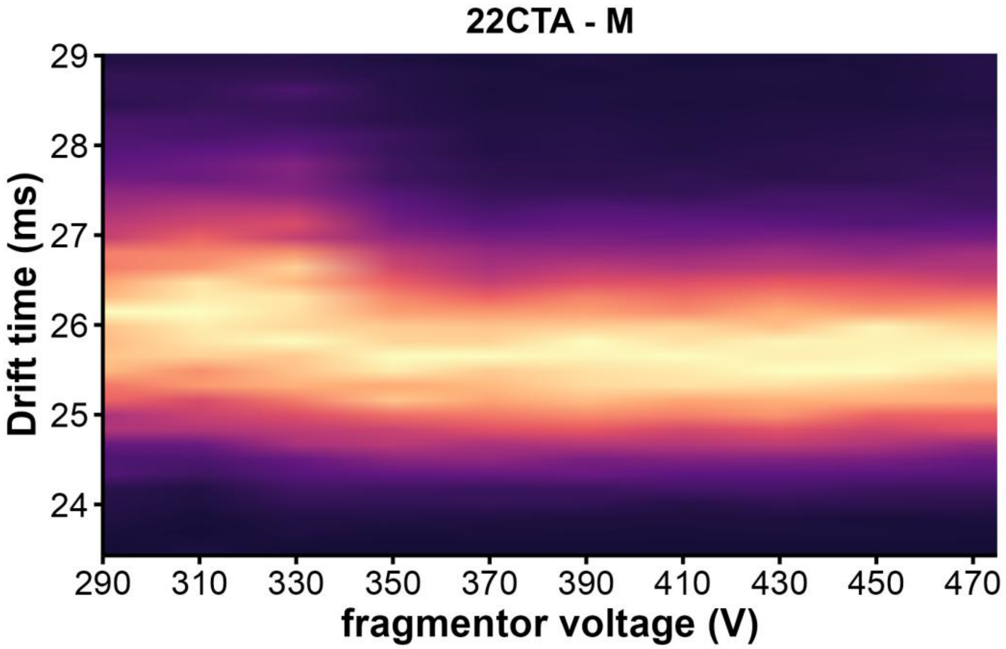


*Figure S65. CIU heatmap of 22CTA · 0K$^{+}$ (z= 4 -) in 100mM TMAA + 1 mM KCl*

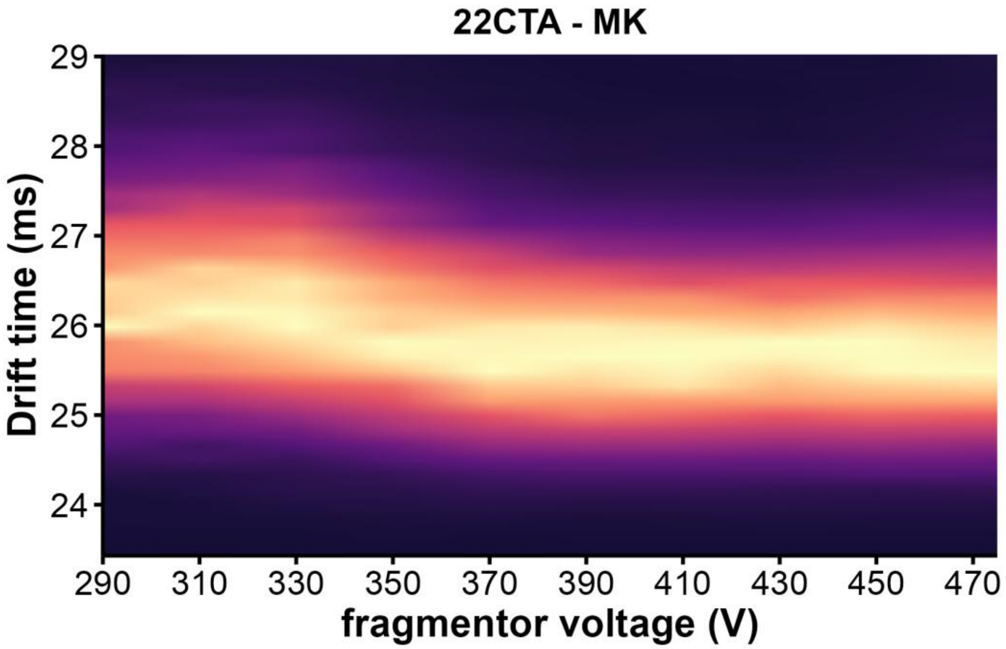


*Figure S66. CIU heatmap of 22CTA · 1K$^{+}$ (z= 4 -) in 100mM TMAA + 1 mM KCl*

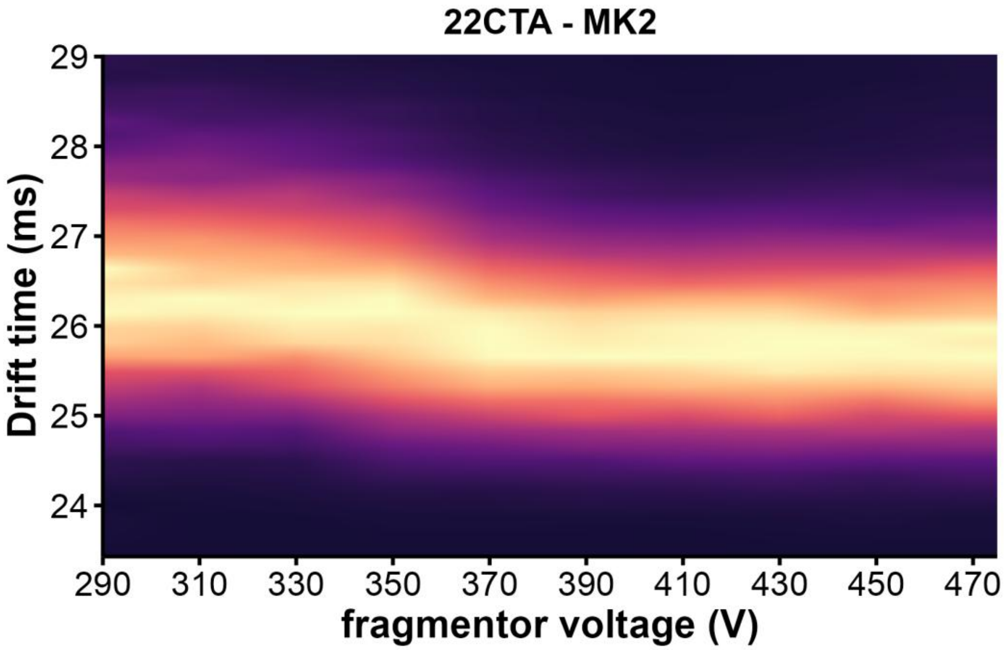


*Figure S67. CIU heatmap of 22CTA·2K⁺ (z= 4 -) in 100mM TMAA + 1 mM KCl*

*23AG*

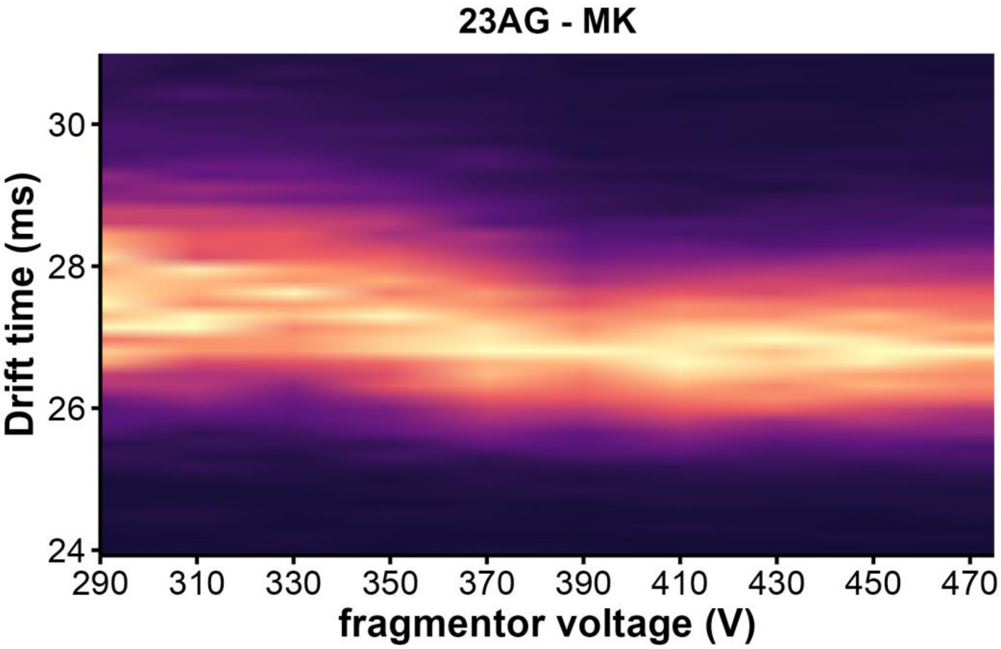


*Figure S68. CIU heatmap of 23AG · 1K$^+$ (z= 4 -) in 100mM TMAA + 1 mM KCl*

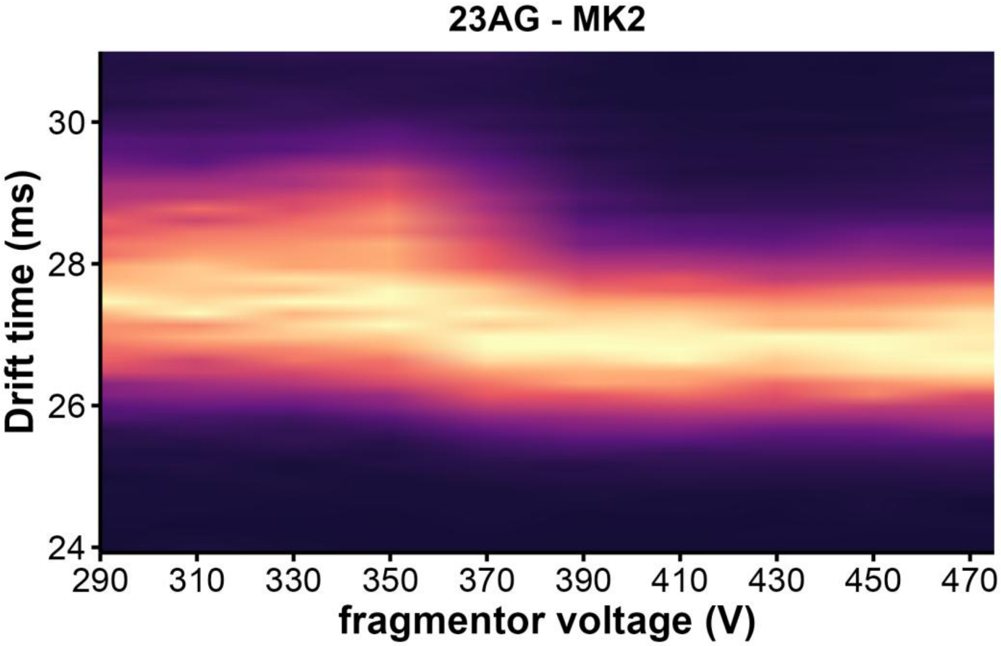


*Figure S69. CIU heatmap of 23AG · 2K$^+$ (z= 4 -) in 100mM TMAA + 1 mM KCl*

*23AG_I14*

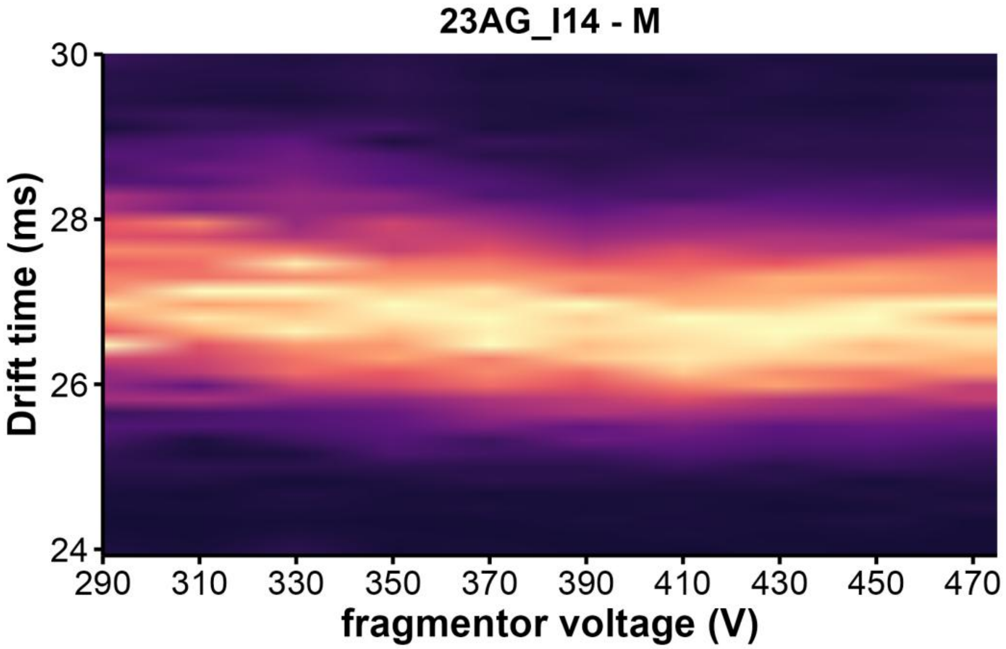


*Figure S70. CIU heatmap of 23AG_I14 · 0K⁺ (z= 4 -) in 100mM TMAA + 1 mM KCl*

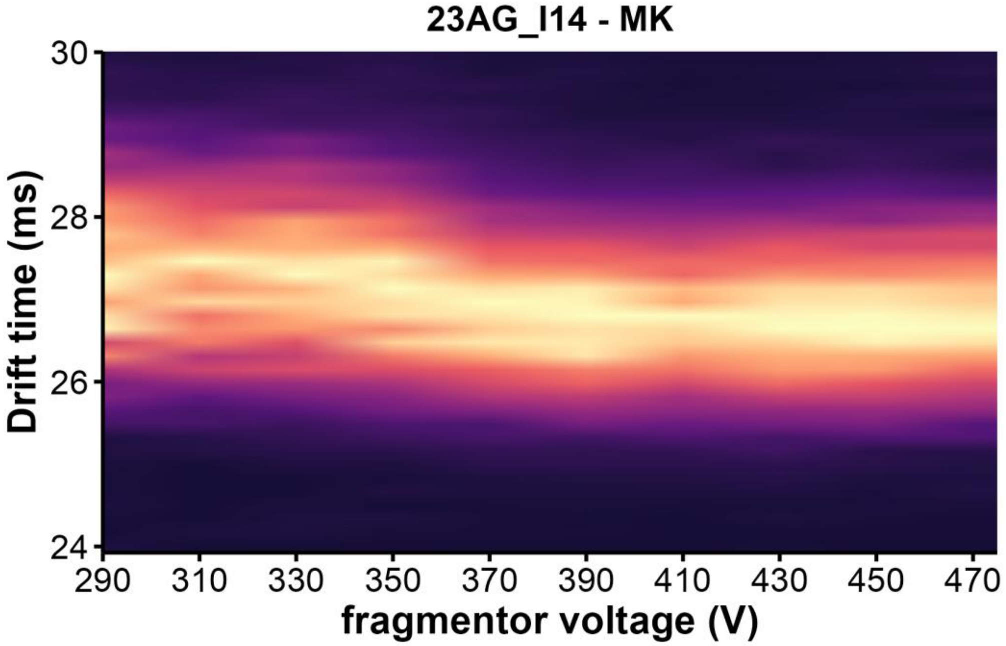


*Figure S71. CIU heatmap of 23AG_I14 · 1K⁺ (z= 4 -) in 100mM TMAA + 1 mM KCl*

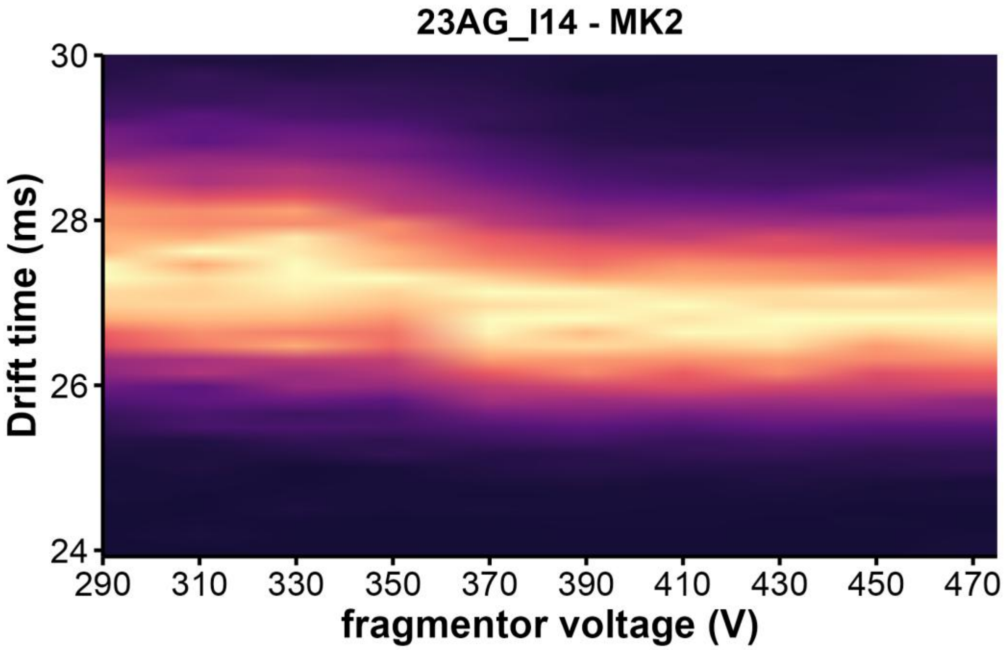


*Figure S72. CIU heatmap of 23AG_I14 · 2K⁺ (z= 4 -) in 100mM TMAA + 1 mM KCl*

*23TAG*

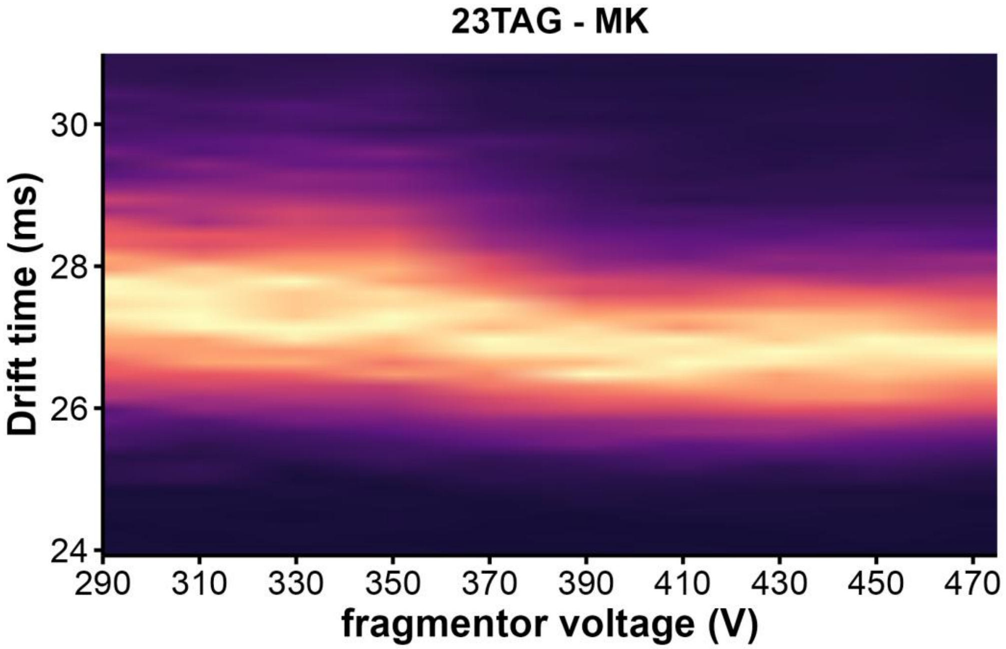


*Figure S73. CIU heatmap of 23TAG · 1K⁺ (z= 4 -) in 100mM TMAA + 1 mM KCl*

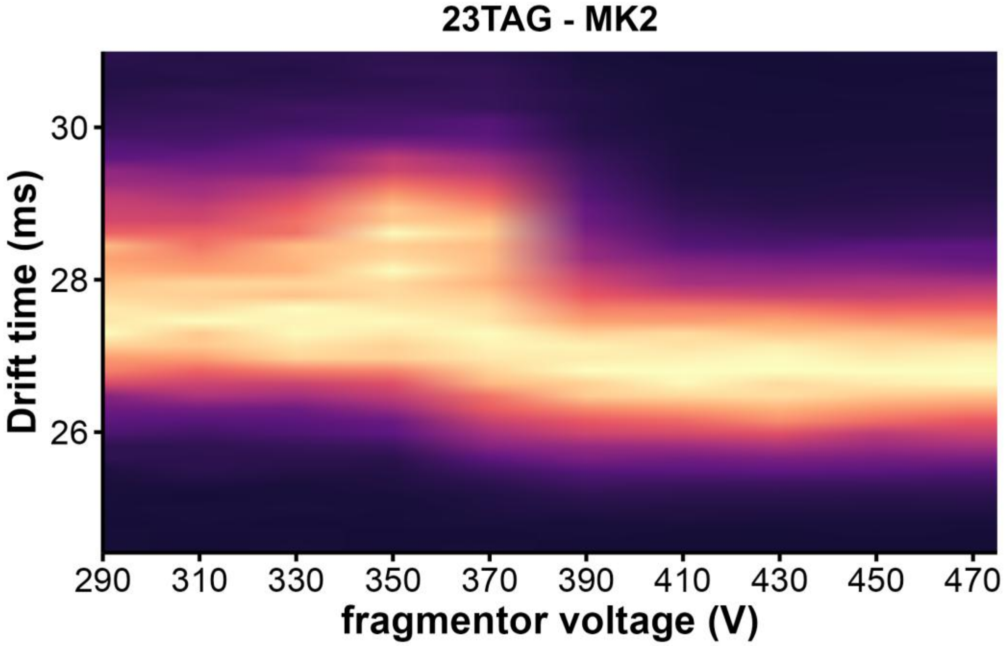


*Figure S74. CIU heatmap of 23TAG · 2K⁺ (z= 4 -) in 100mM TMAA + 1 mM KCl*

*24TTG*

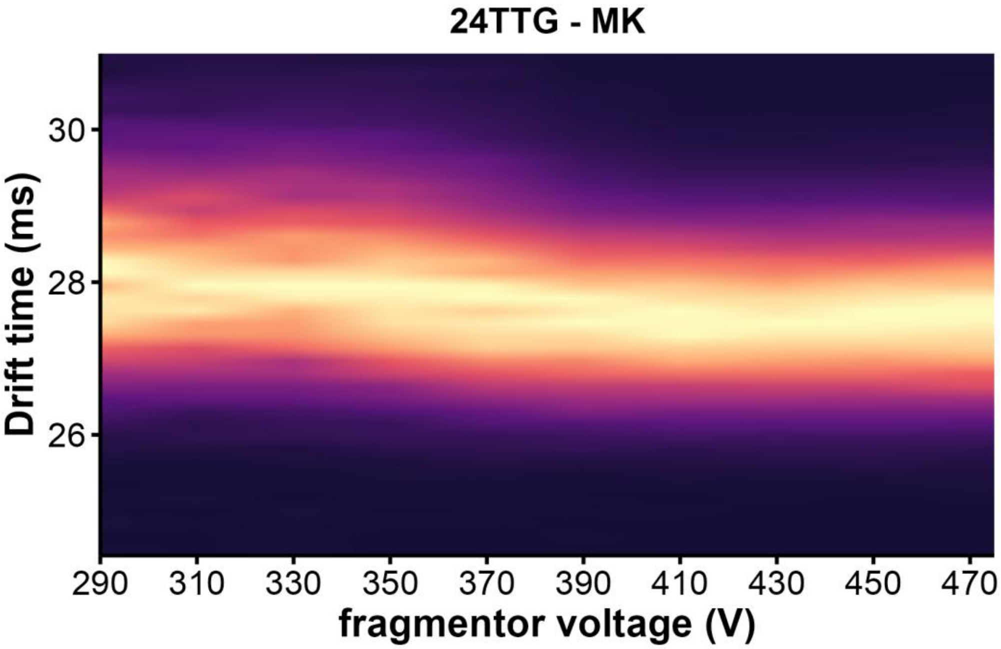


*Figure S75. CIU heatmap of 24TTG · 1K⁺ (z= 4 -) in 100mM TMAA + 0.1 mM KCl*

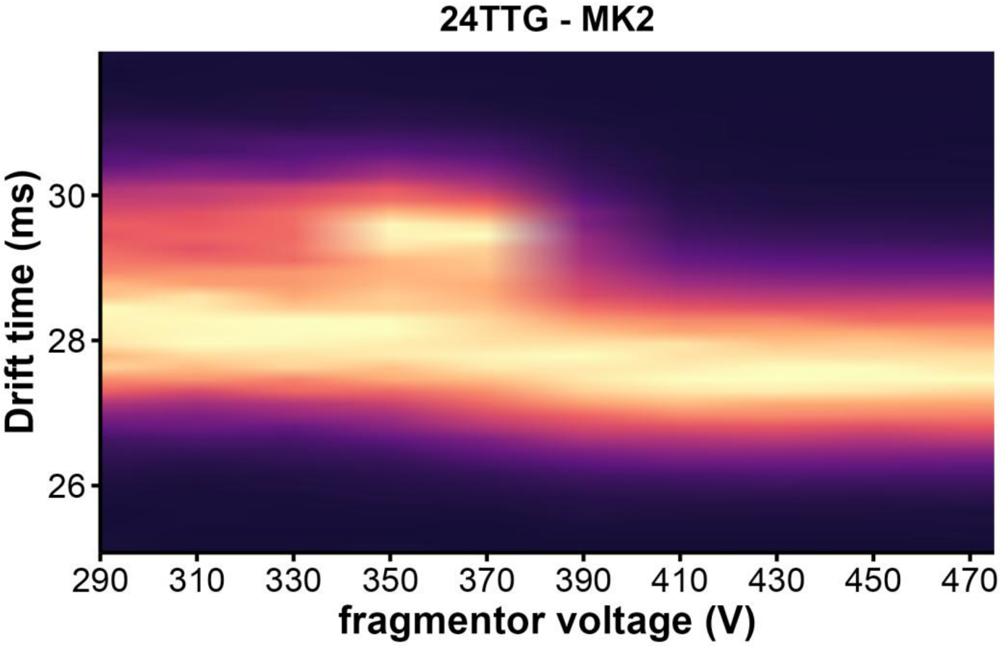


*Figure S76. CIU heatmap of 24TTG · 2K⁺ (z= 4 -) in 100mM TMAA + 0.1 mM KCl*

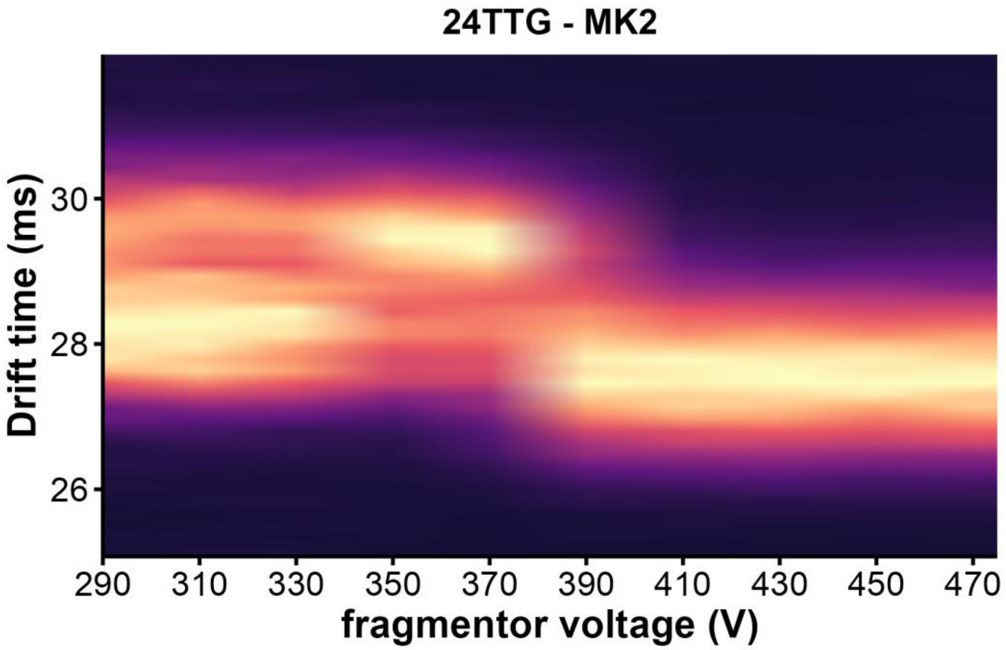


*Figure S77. CIU heatmap of 24TTG · 2K⁺ (z= 4 -) in 100mM TMAA + 1 mM KCl*

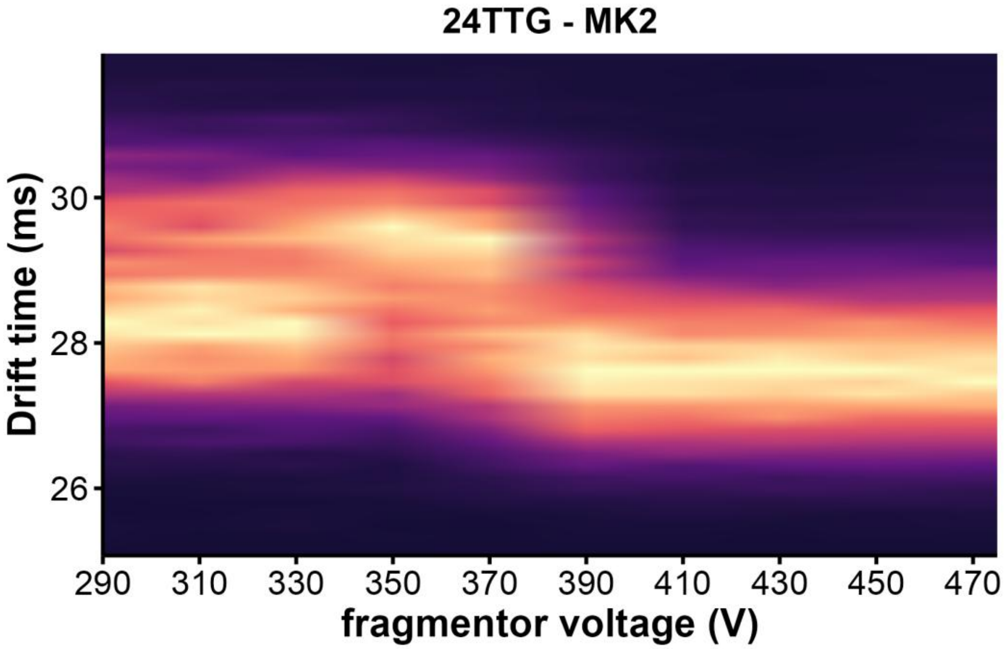


*Figure S78. CIU heatmap of 24TTG · 2K⁺ (z= 4 -) in 100mM TMAA + 0.5 mM KCl*

*25TAG*

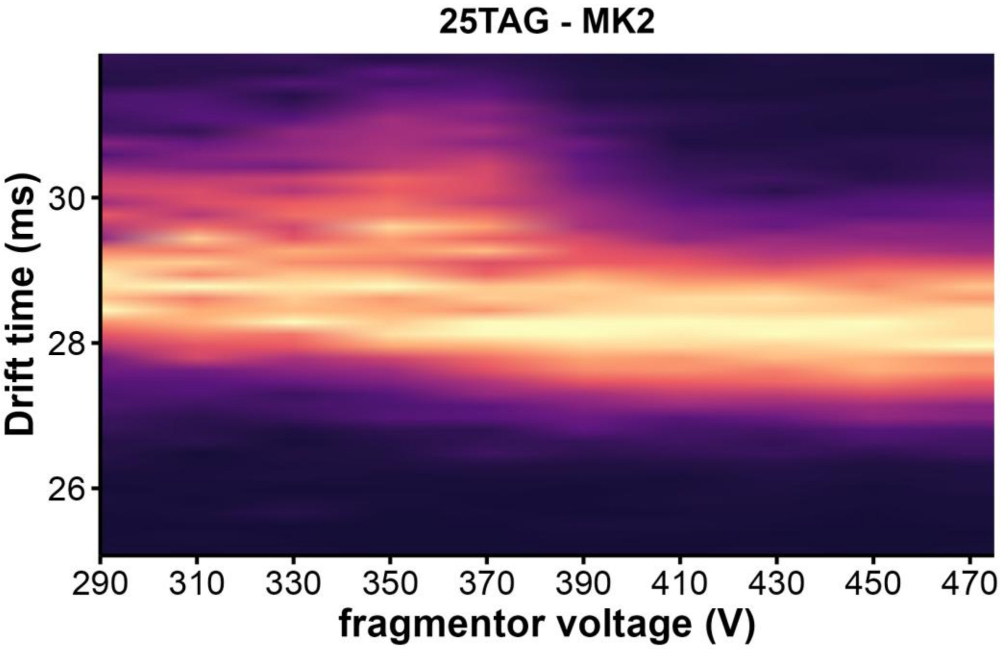


*Figure S79. CIU heatmap of 25TAG · $2K^{+}$ (z= 4 -) in 100mM TMAA + 1 mM KCl*

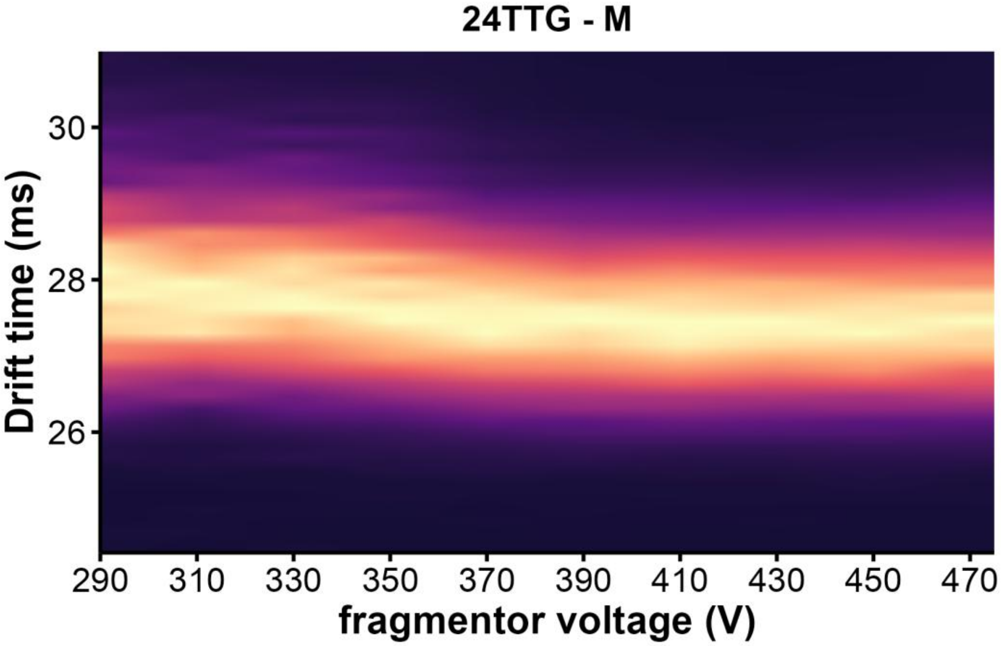


*Figure S80. CIU heatmap of 24TTG · $0K^{+}$ (z= 4 -) in 100mM TMAA + 0.1 mM KCl*

*222T*

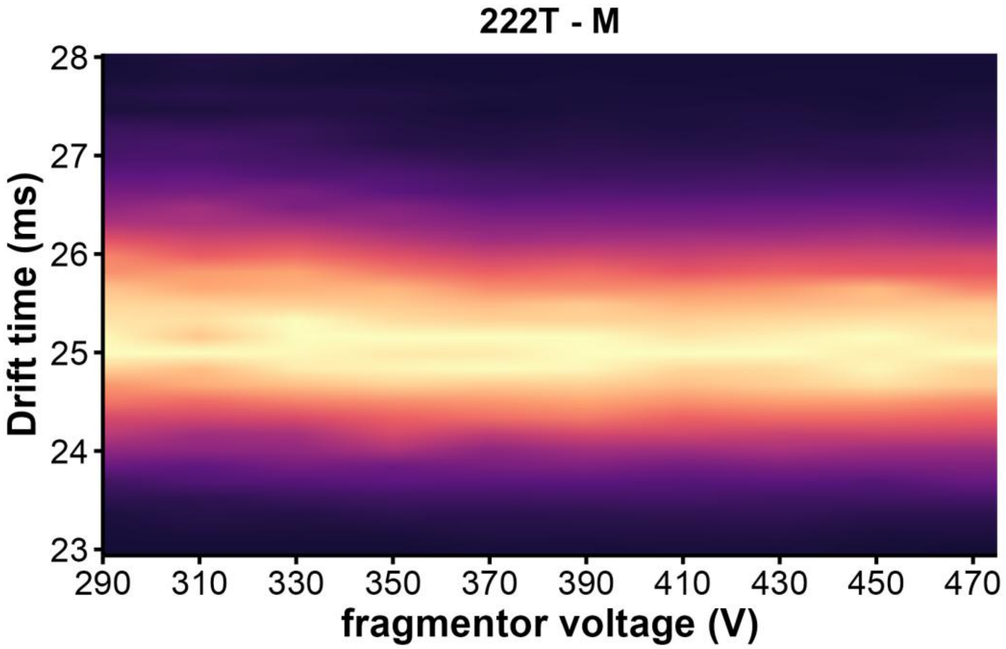


*Figure S81. CIU heatmap of 222T · 0K⁺ (z= 4 -) in 100mM TMAA + 0.1 mM KCl*

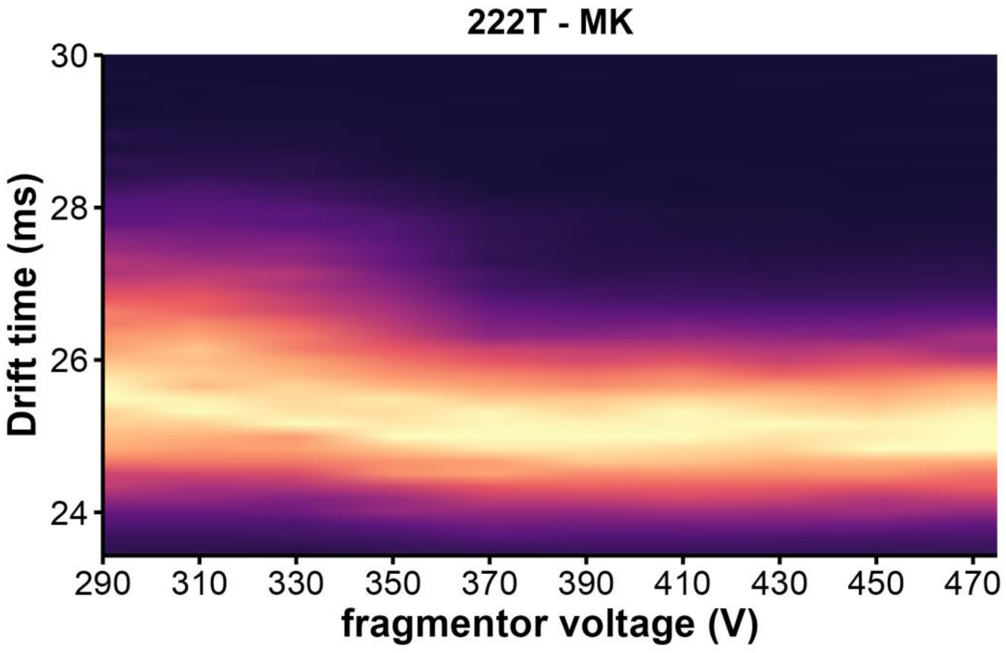


*Figure S82. CIU heatmap of 222T · 1K⁺ (z= 4 -) in 100mM TMAA + 0.1 mM KCl*

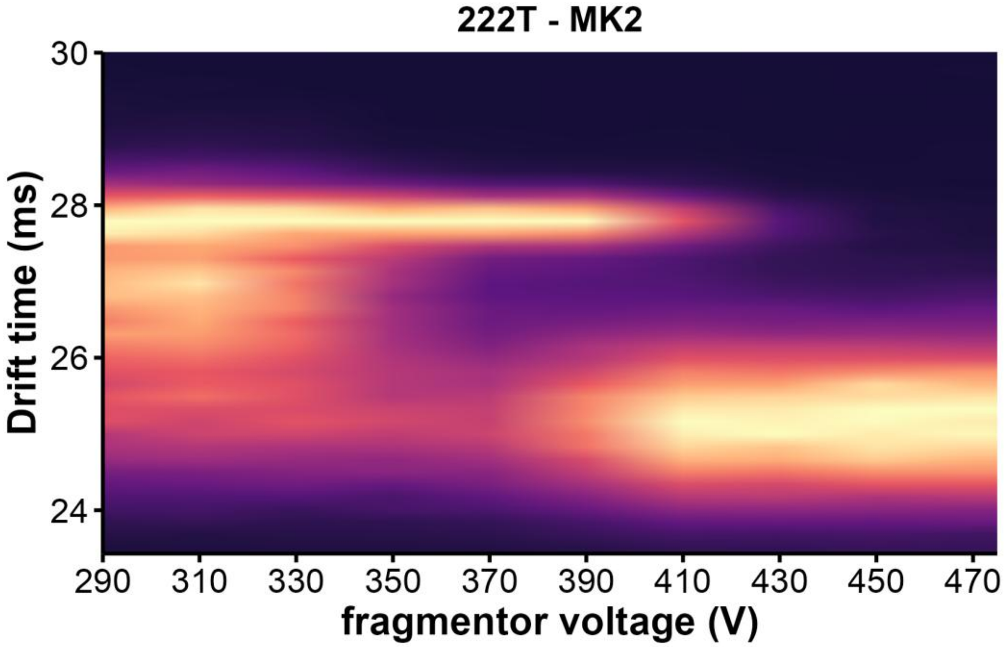


*Figure S83. CIU heatmap of 222T·2K⁺ (z= 4 -) in 100mM TMAA + 0.1 mM KCl*

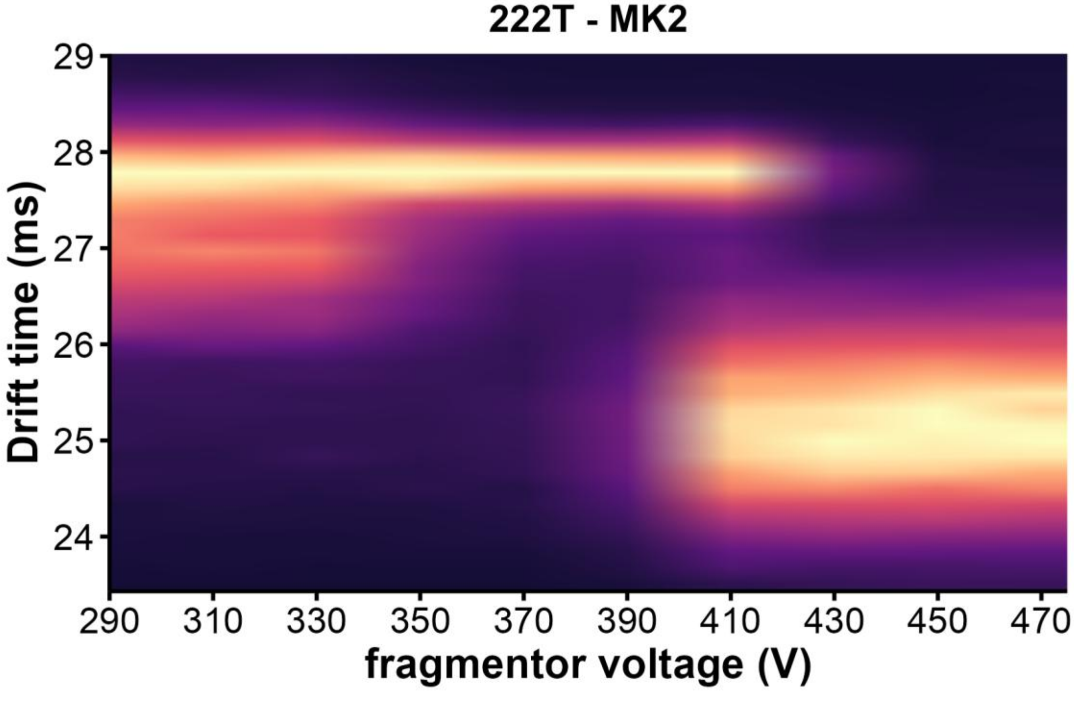


*Figure S84. CIU heatmap of 222T·2K⁺ (z= 4 -) in 100mM TMAA + 1 mM KCl*

*ckit-87up*

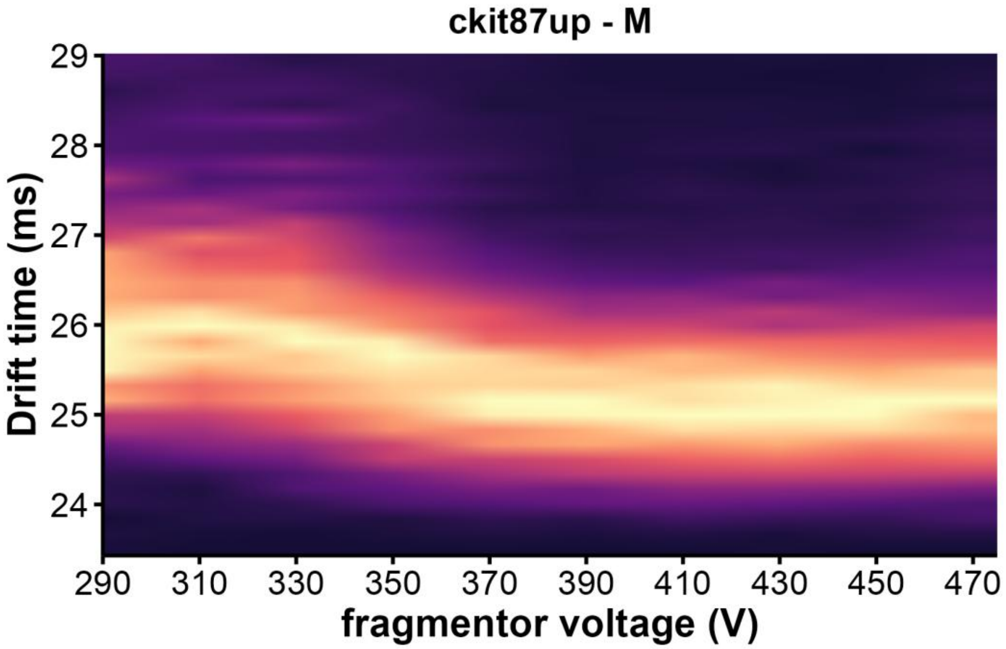


*Figure S85. CIU heatmap of ckit87up · 0K⁺ (z= 4 -) in 100mM TMAA + 1 mM KCl*

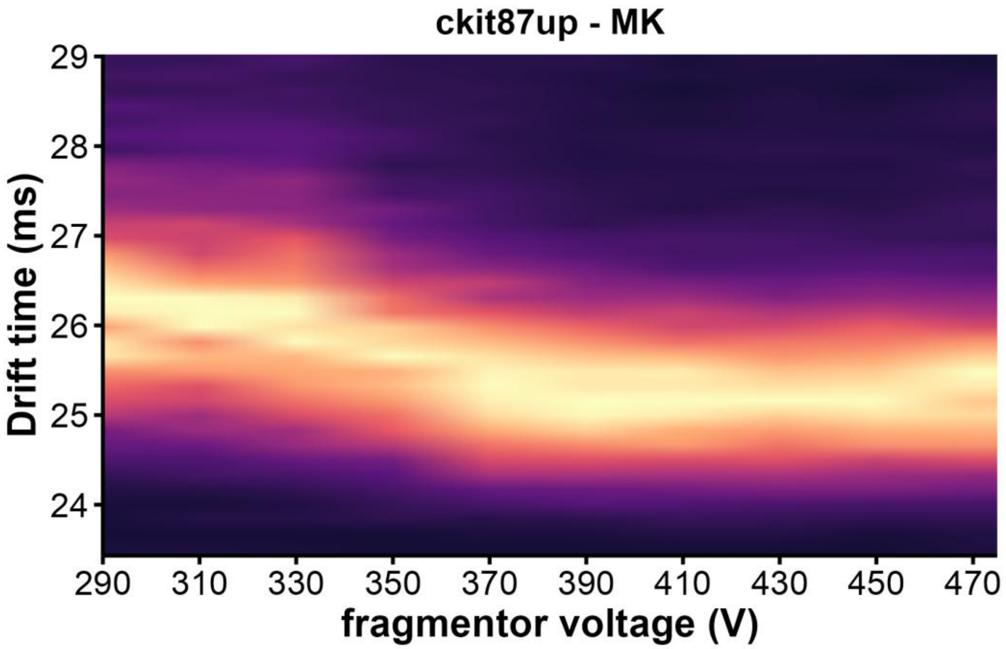


*Figure S86. CIU heatmap of ckit87up · 1K⁺ (z= 4 -) in 100mM TMAA + 1 mM KCl*

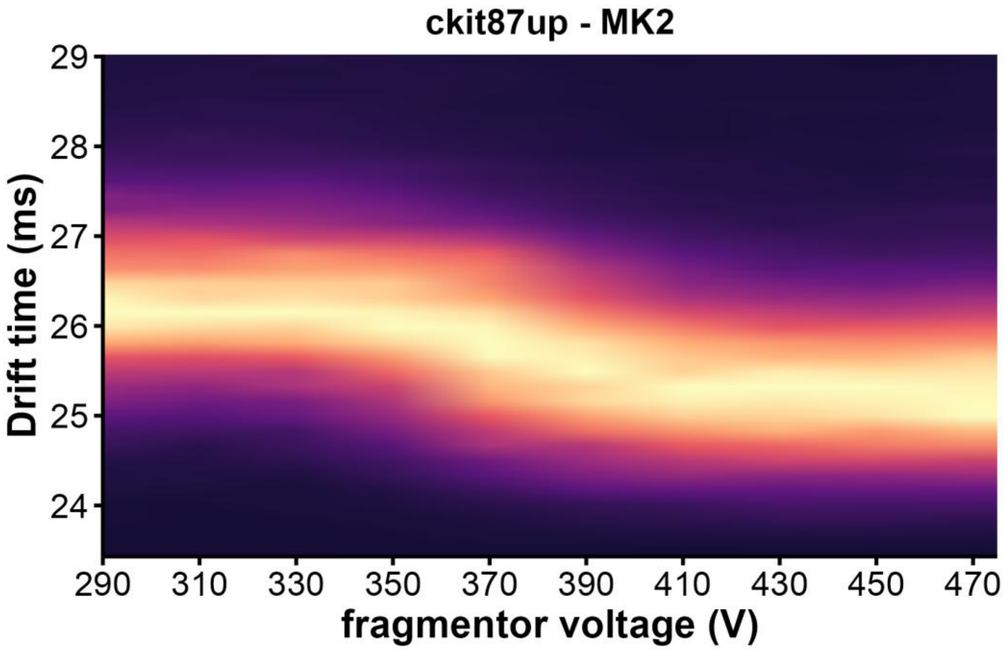


*Figure S87. CIU heatmap of ckit87up · 2K⁺ (z= 4 -) in 100mM TMAA + 1 mM KCl*

### *BCl2*

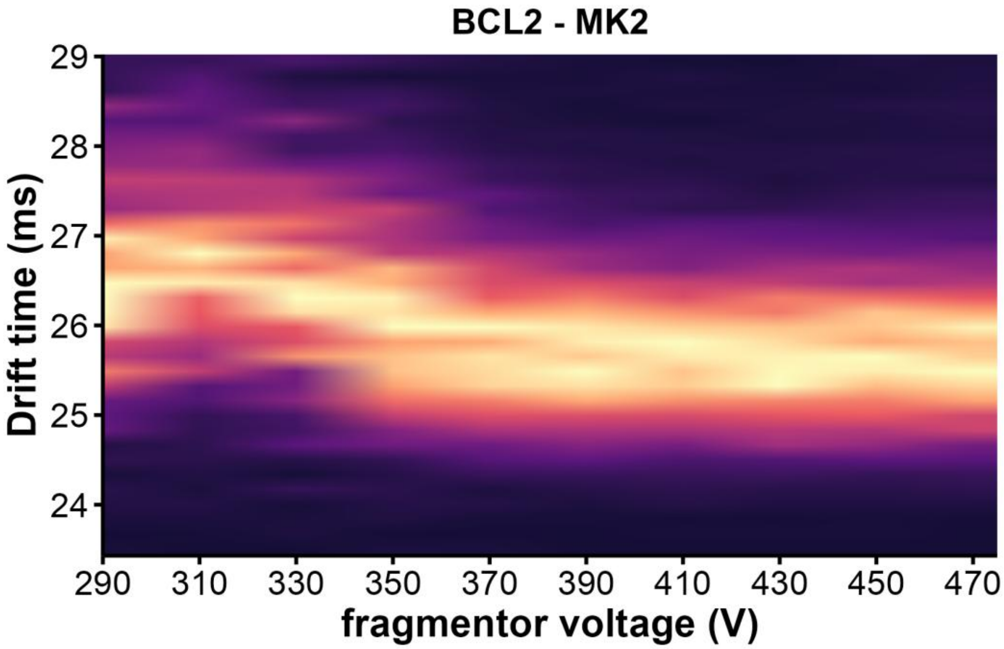


*Figure S88. CIU heatmap of BCL2 · $2K^+$ (z= 4 -) in 100mM TMAA + 1 mM KCl*

### 2KPR

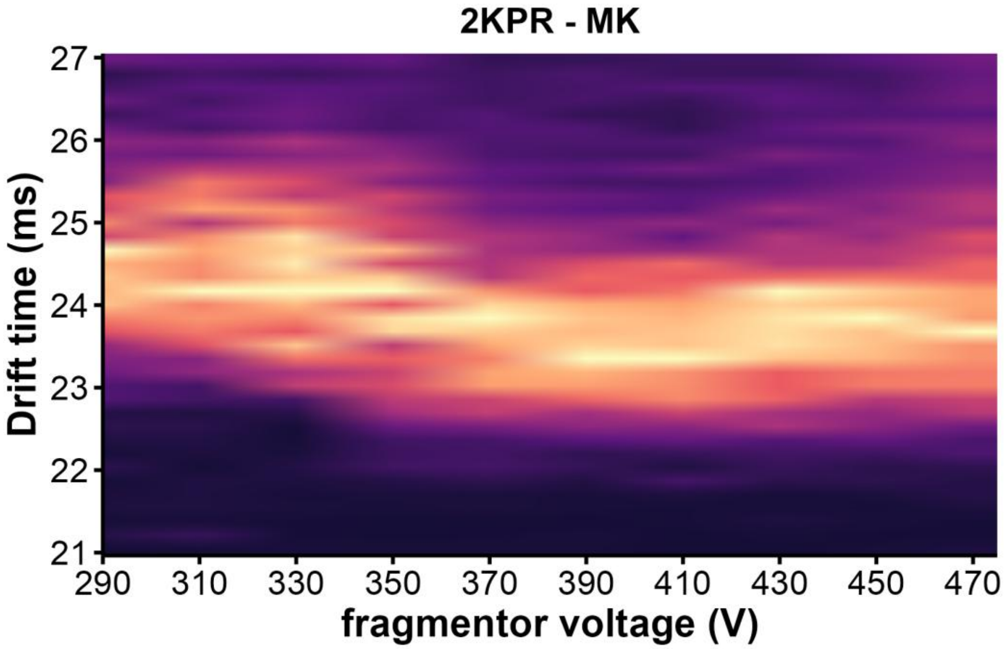


*Figure S89. CIU heatmap of 2KPR · 1K⁺ (z= 4 -) in 100mM TMAA + 1 mM KCl*

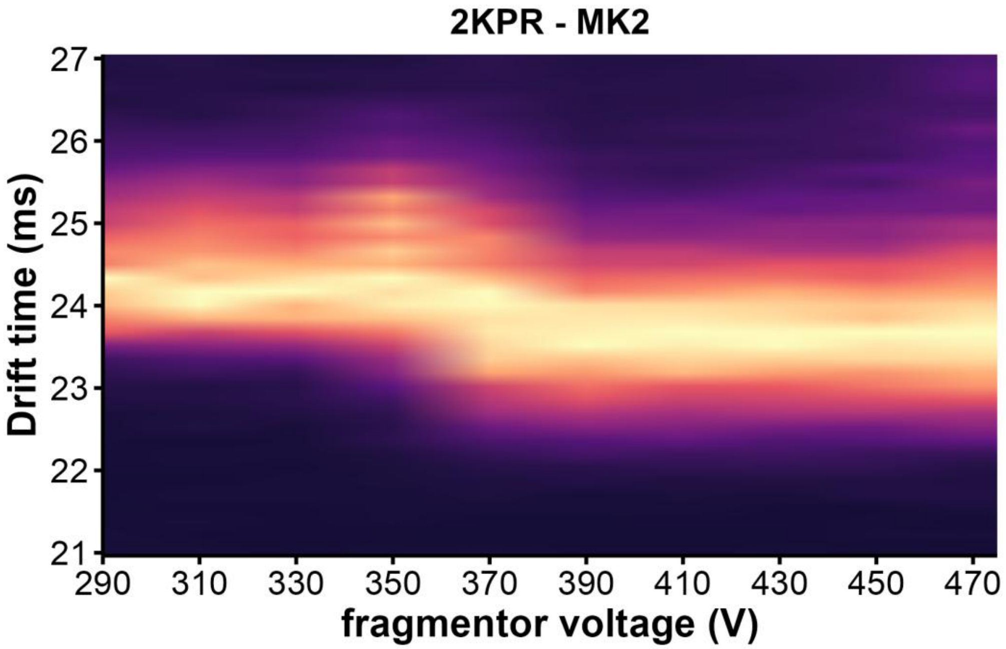


*Figure S90. CIU heatmap of 2KPR · 2K⁺ (z= 4 -) in 100mM TMAA + 1 mM KCl*

*26CEB*

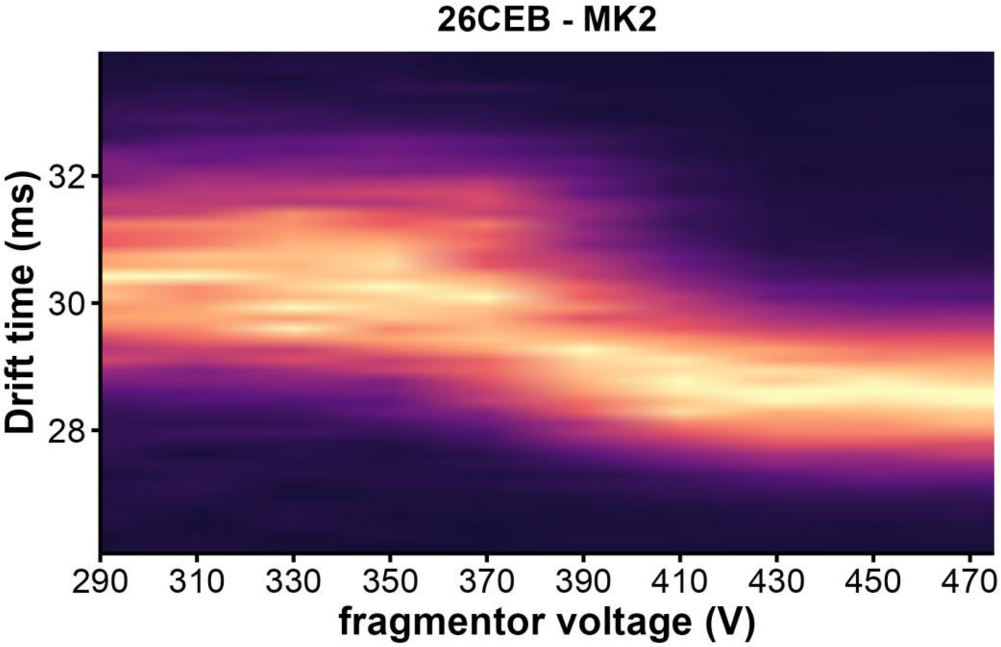


*Figure S91. CIU heatmap of 26CEB · 2K⁺ (z= 4 -) in 100mM TMAA + 1 mM KCl*

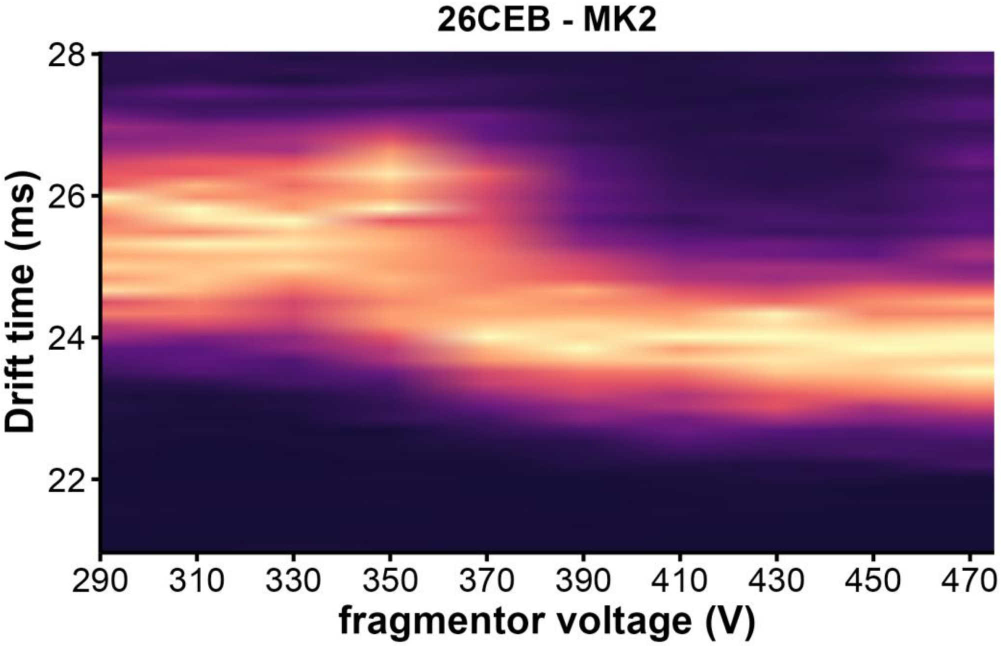


*Figure S92. CIU heatmap of 26CEB · 2K⁺ (z= 5 -) in 100mM TMAA + 1 mM KCl*

*T30177TT*

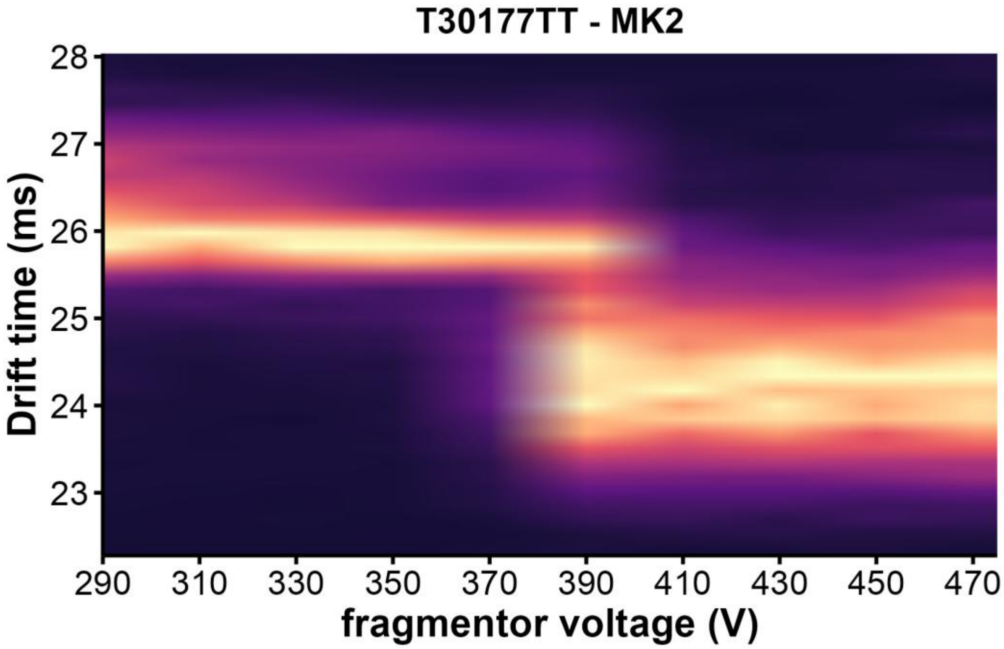


*Figure S93. CIU heatmap of T30177TT·2K⁺ (z= 4 -) in 100mM TMAA + 0.5 mM KCl*

*T95TT*

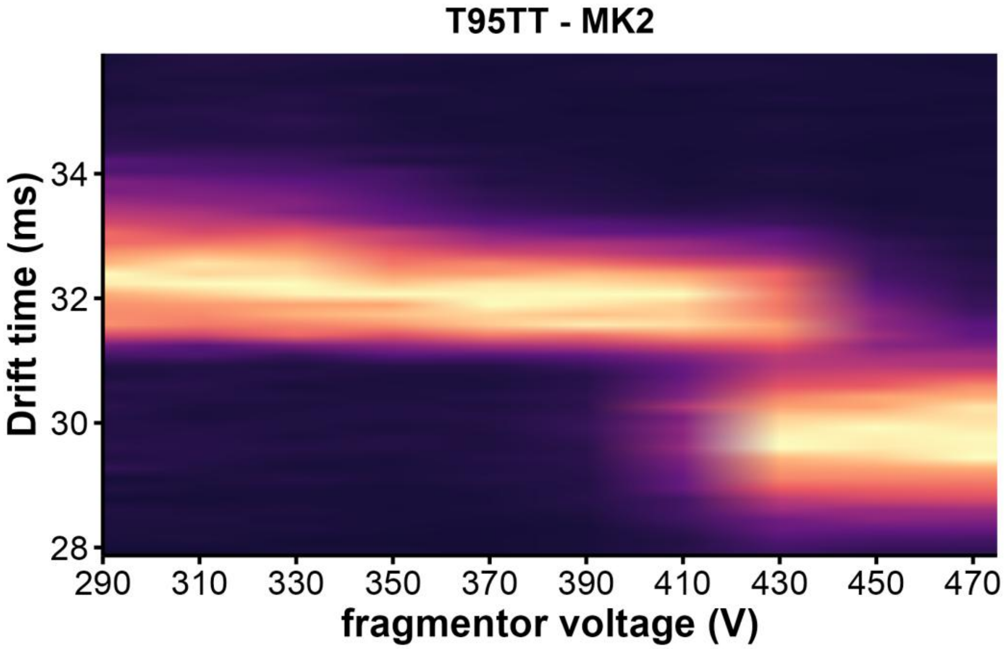


*Figure S94. CIU heatmap of T95TT · 2K⁺ (z= 3 -) in 100mM TMAA + 0.5 mM KCl*

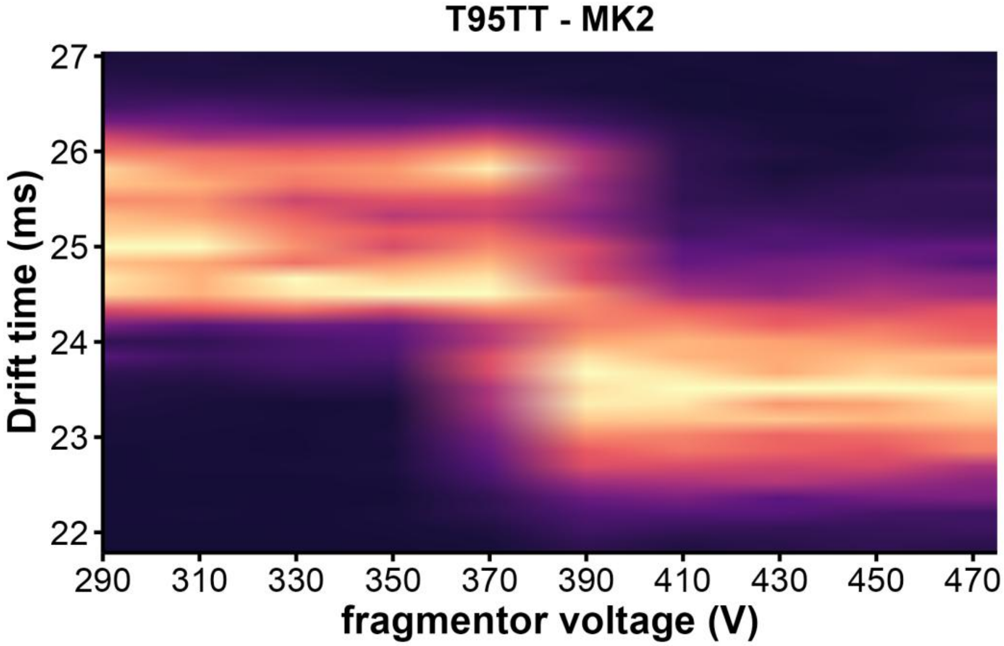


*Figure S95. CIU heatmap of T95TT · 2K⁺ (z= 4 -) in 100mM TMAA + 0.5 mM KCl*

*TG4T*

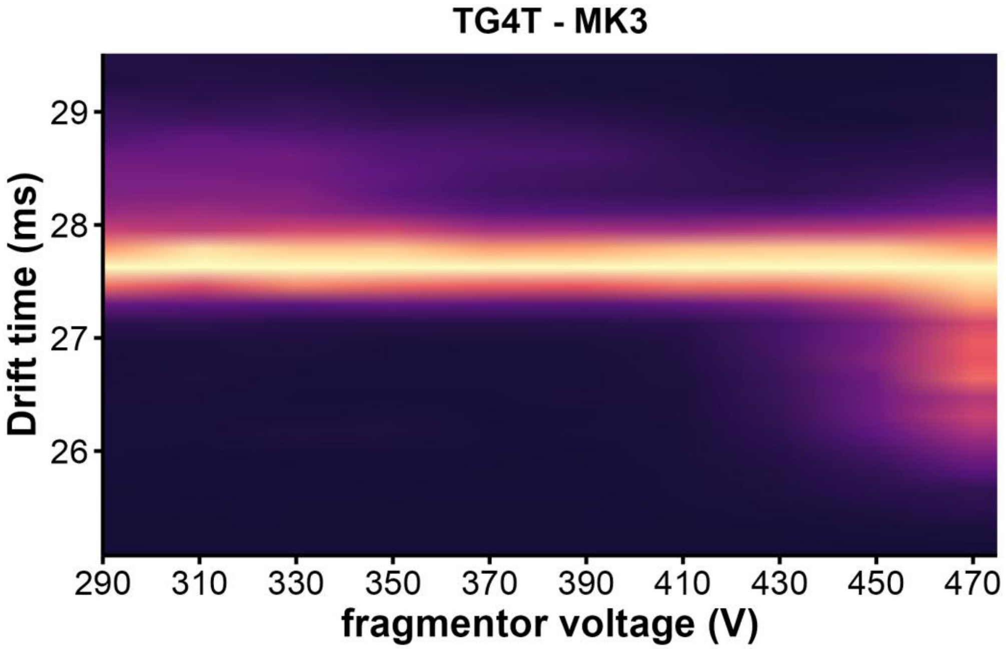


*Figure S96. CIU heatmap of TG4T·3K$^{+}$ (z= 4 -) in 100mM TMAA + 1 mM KCl*

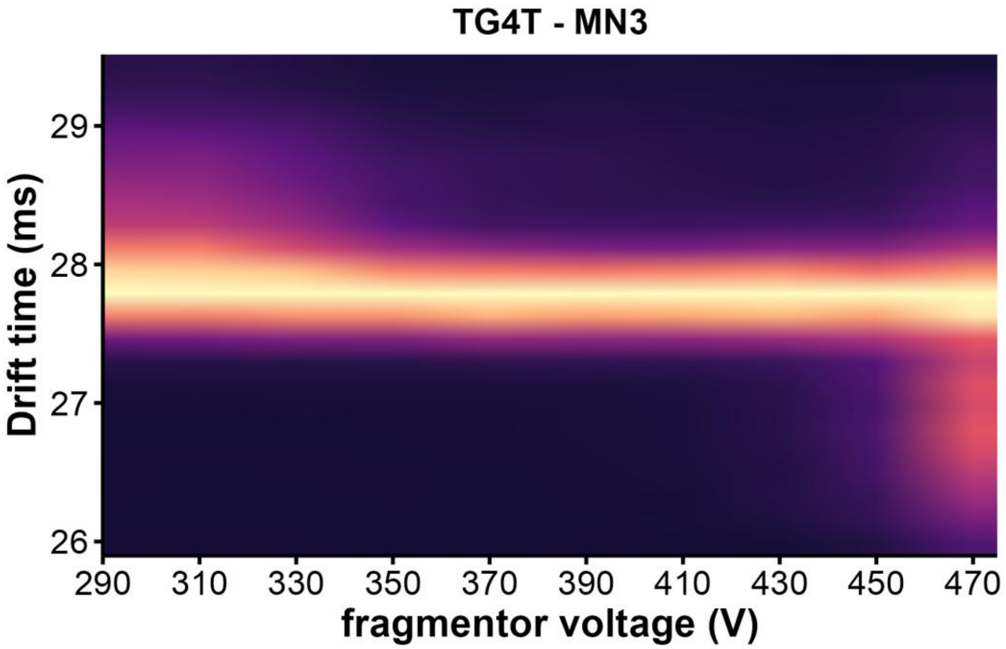


*Figure S97. CIU heatmap of TG4T·M($NH_4^+$)$_3$ (z= 4 -) in 100 mM $NH_4OAc$*

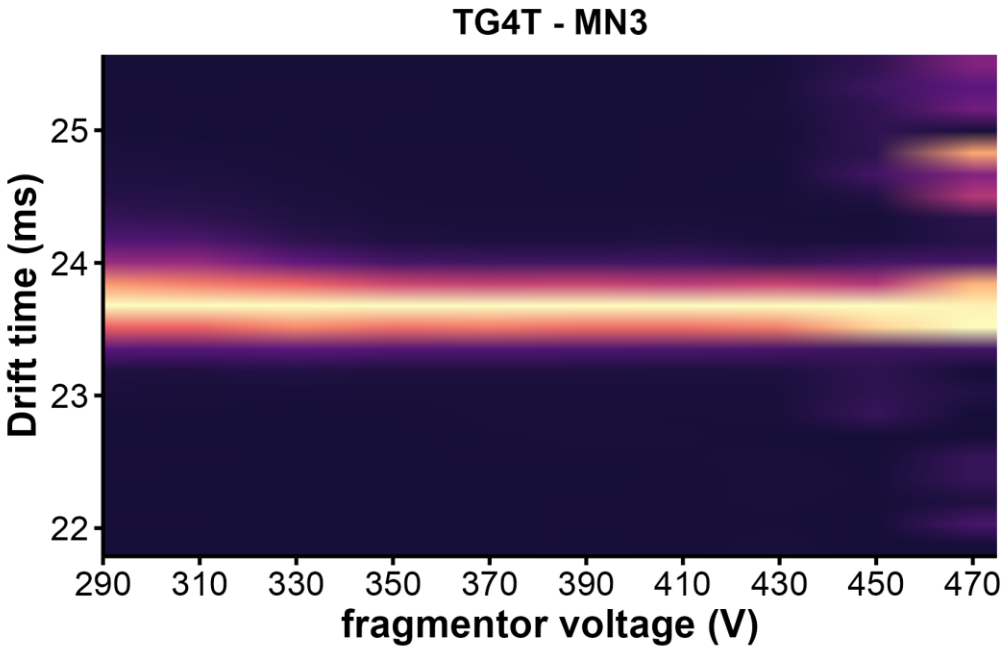


Figure S98. CIU heatmap of TG4T · $M(NH_4^+)_3$ (z= 5 -) in 100 mM $NH_4OAc$

### HDX/IMS setup

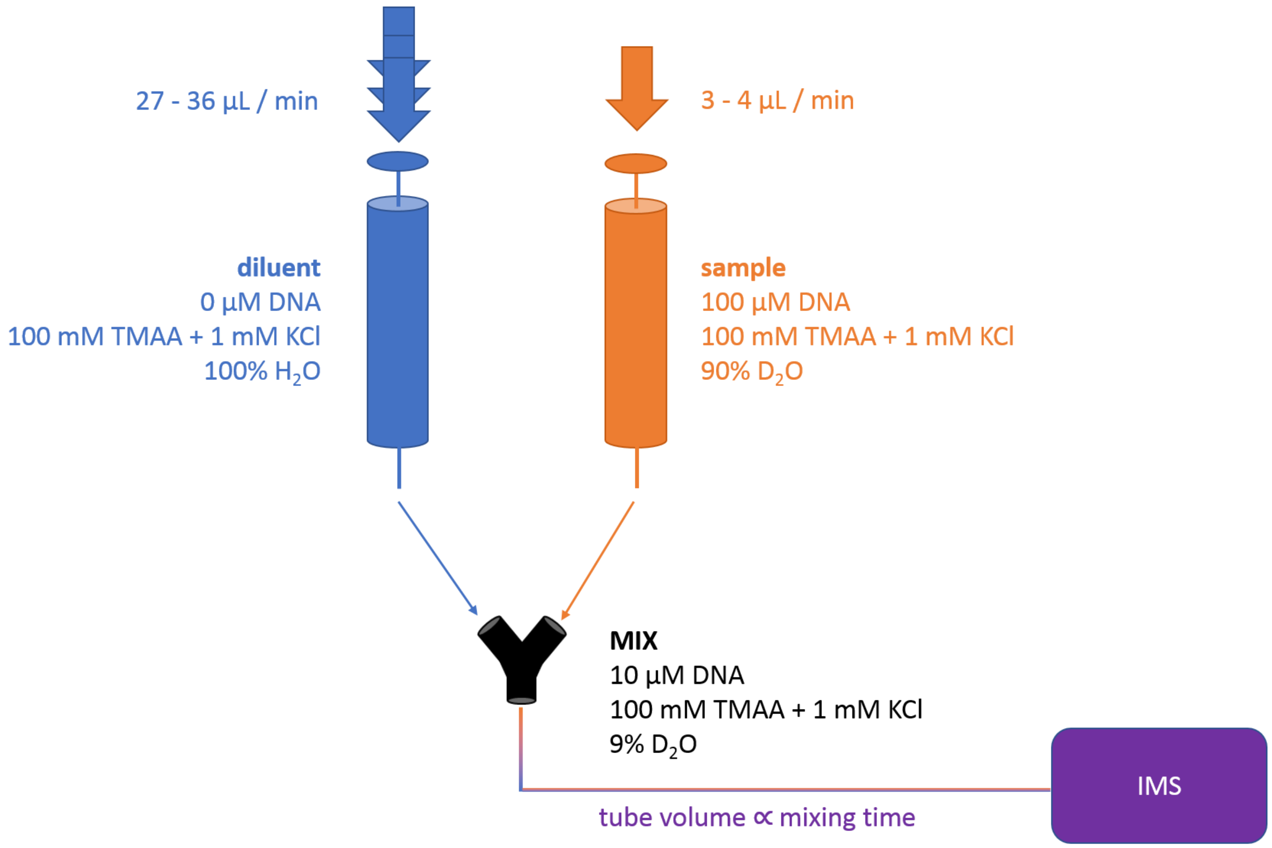


*Figure S99. HDX/MS continuous flow setup. Two syringes set on a two-channel syringe pump. In the mixing tee the sample is diluted by 10 and the $D_2O$ content drops to 9%. Increasing the tube volumes between the tee and the MS induces a longer mixing time that ranges from 3s to 5min.*

### HDX kinetics and density maps

*21G / MK / z = 4- / 1 mM KCl / 320 V*

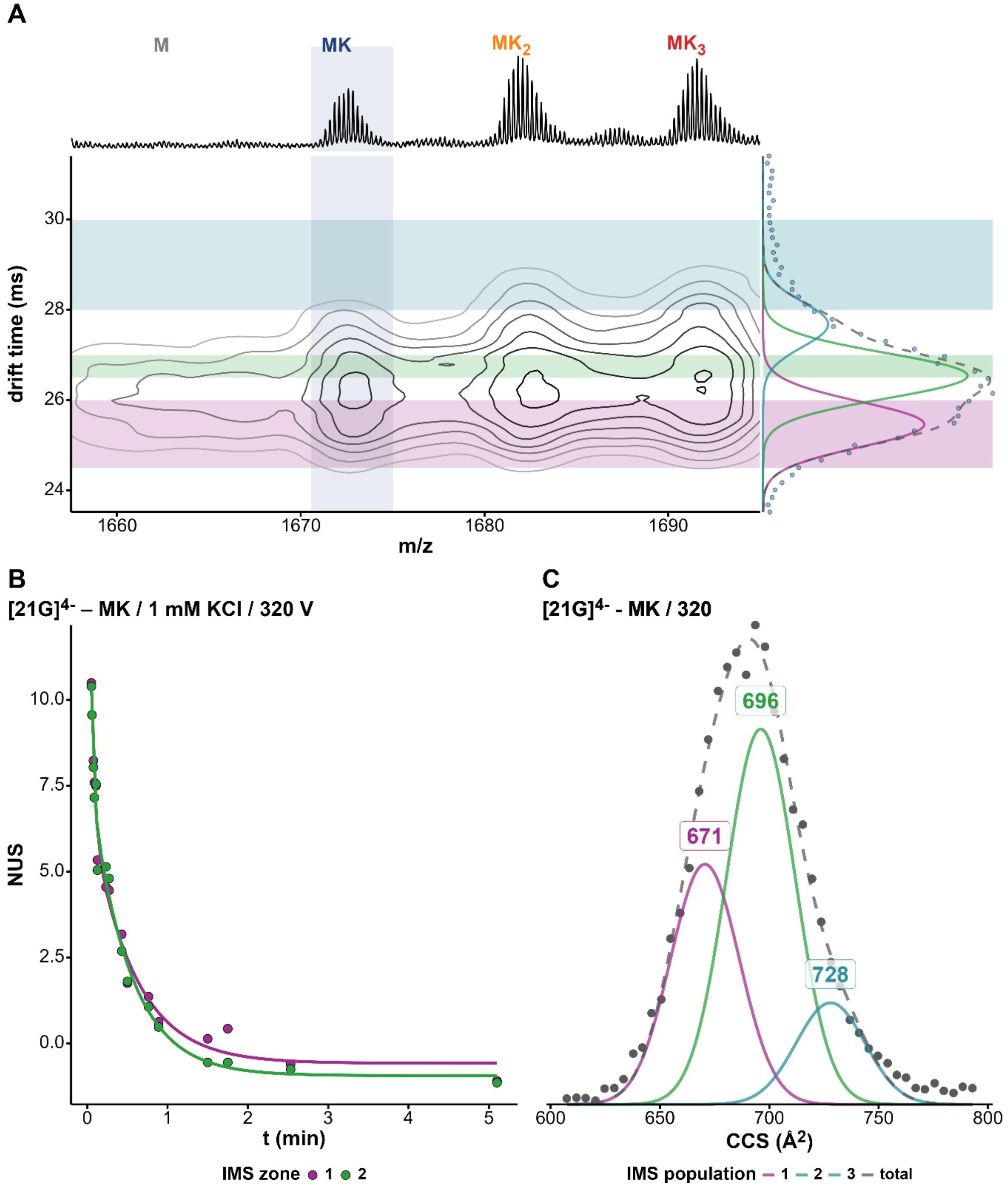


*Figure S100. (A) Density map of 21G in 100 mM TMAA + 1 mM KCl at fragmentor 320V and charge state 4-. The x-axis represents the mass spectrum and the y-axis the arrival-time distribution (ATD) of the 1K⁺ adduct with the Gaussian fits and the different zones selected for HDX/IMS. (B) HDX/IMS kinetics with colors matching the IMS zones of the density map for the same K⁺ adduct. (C) CCS distribution of the 1K⁺ adduct of 21G with the CCS values of each population.*

*21G / $MK_2$ / z = 4- / 1 mM KCl / 320 V*

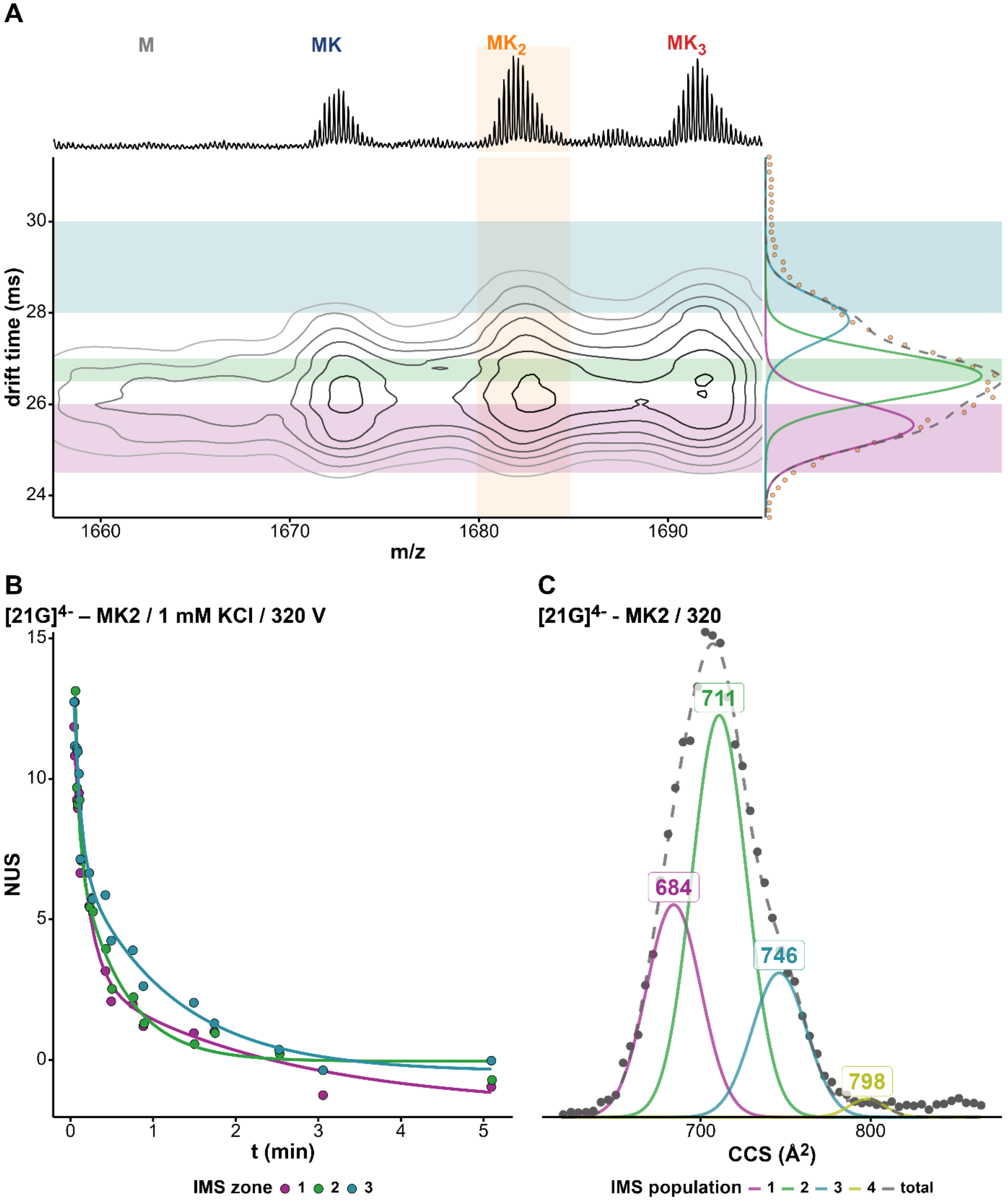


*Figure S101. (A) Density map of 21G in 100 mM TMAA + 1 mM KCl at fragmentor 320V and charge state 4-. The x-axis represents the mass spectrum and the y-axis the arrival-time distribution (ATD) of the $2K^+$ adduct with the Gaussian fits and the different zones selected for HDX/IMS. (B) HDX/IMS kinetics with colors matching the IMS zones of the density map for the same $K^+$ adduct. (C) CCS distribution of the $2K^+$ adduct of 21G with the CCS values of each population.*

*5YEY / MK / z = 4- / 1 mM KCl / 320 V*

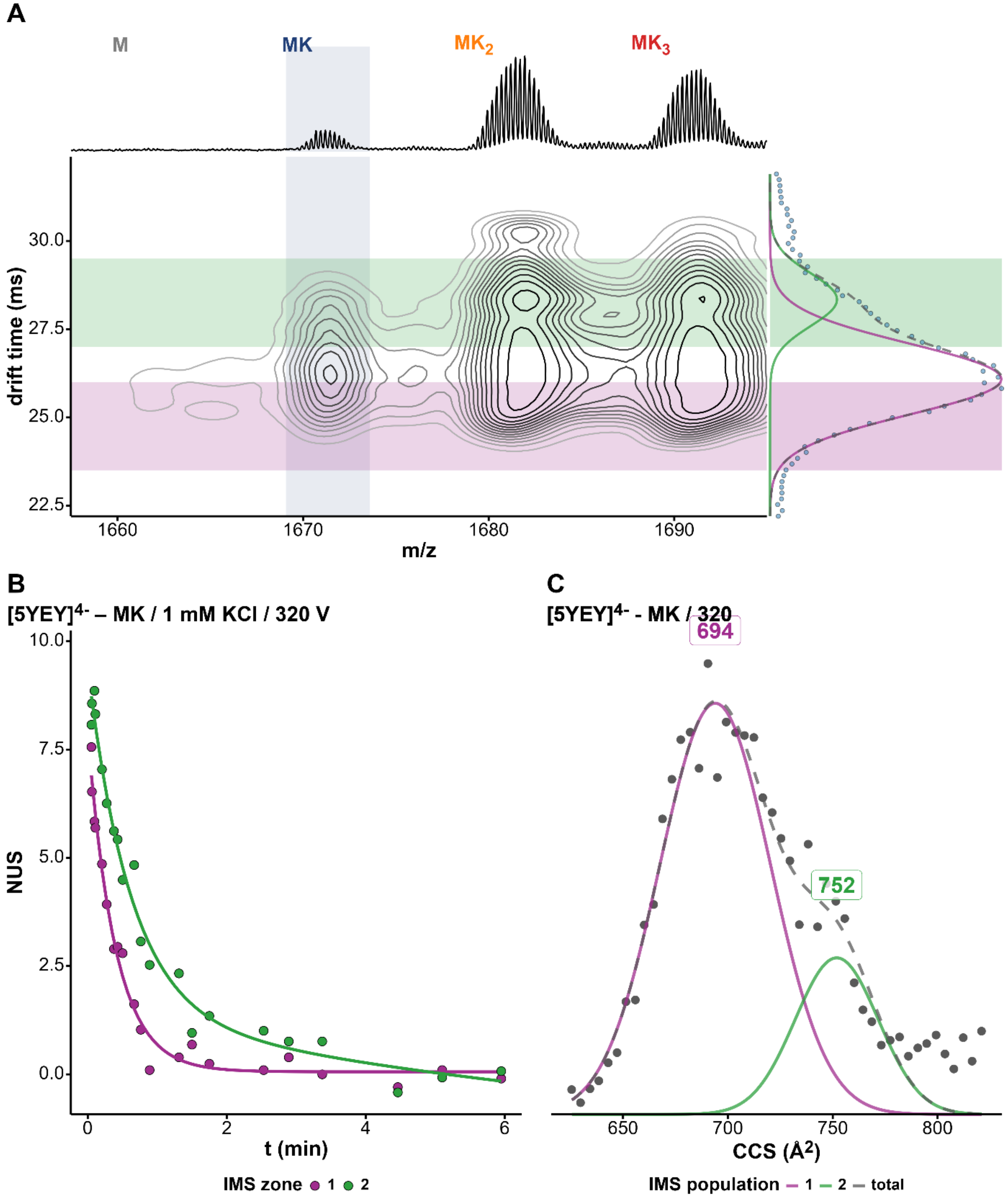


*Figure S102. (A) Density map of 5YEY in 100 mM TMAA + 1 mM KCl at fragmentor 320V and charge state 4-. The x-axis represents the mass spectrum and the y-axis the arrival-time distribution (ATD) of the 1K$^+$ adduct with the Gaussian fits and the different zones selected for HDX/IMS. (B) HDX/IMS kinetics with colors matching the IMS zones of the density map for the same K$^+$ adduct. (C) CCS distribution of the 1K$^+$ adduct of 5YEY with the CCS values of each population.*

*5YEY / $MK_2$ / z = 4- / 1 mM KCl / 320 V*

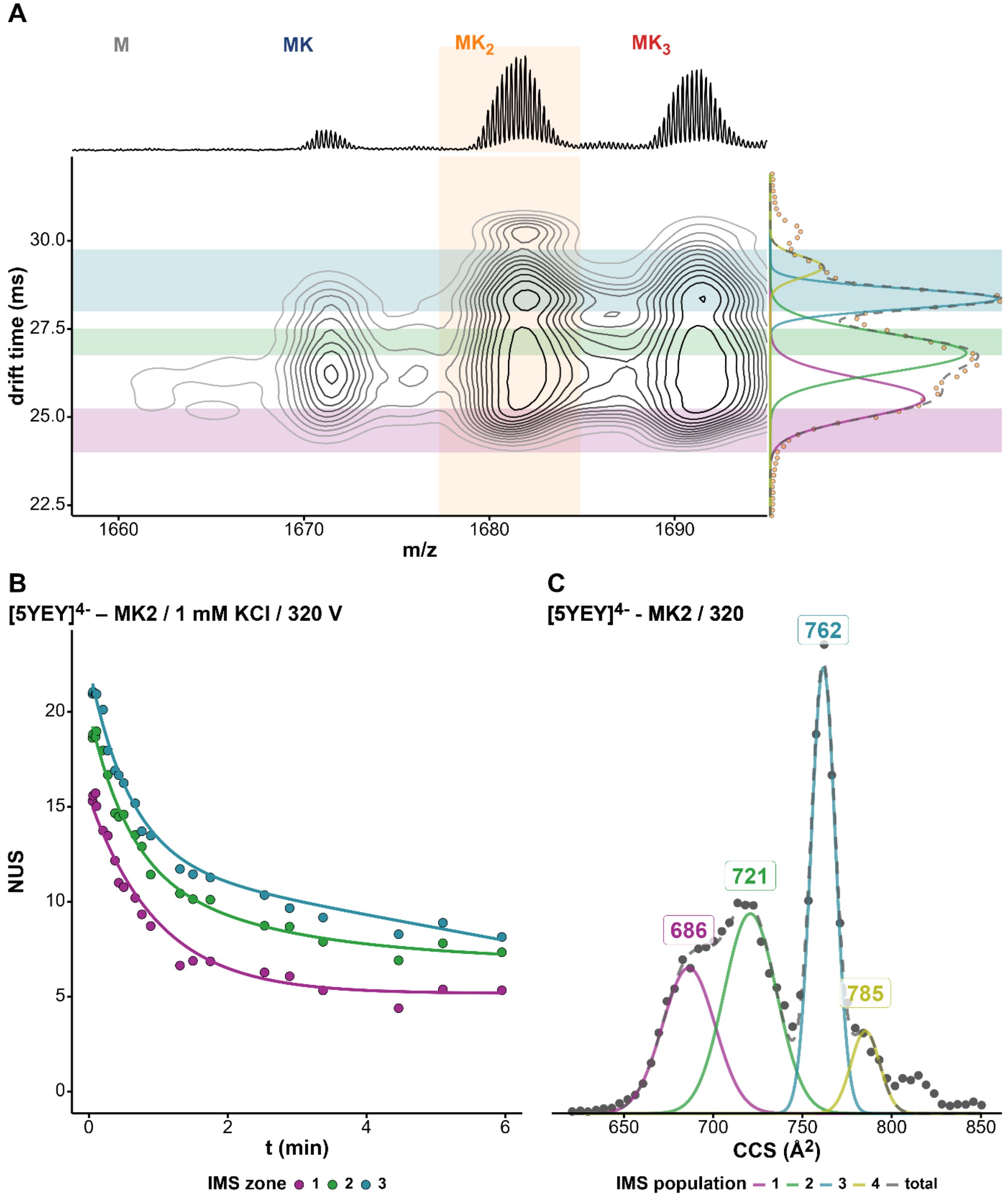


*Figure S103. (A) Density map of 5YEY in 100 mM TMAA + 1 mM KCl at fragmentor 320V and charge state 4-. The x-axis represents the mass spectrum and the y-axis the arrival-time distribution (ATD) of the $2K^+$ adduct with the Gaussian fits and the different zones selected for HDX/IMS. (B) HDX/IMS kinetics with colors matching the IMS zones of the density map for the same $K^+$ adduct. (C) CCS distribution of the $2K^+$ adduct of 5YEY with the CCS values of each population.*

*22AG / MK / z = 4- / 1 mM KCl / 320 V*

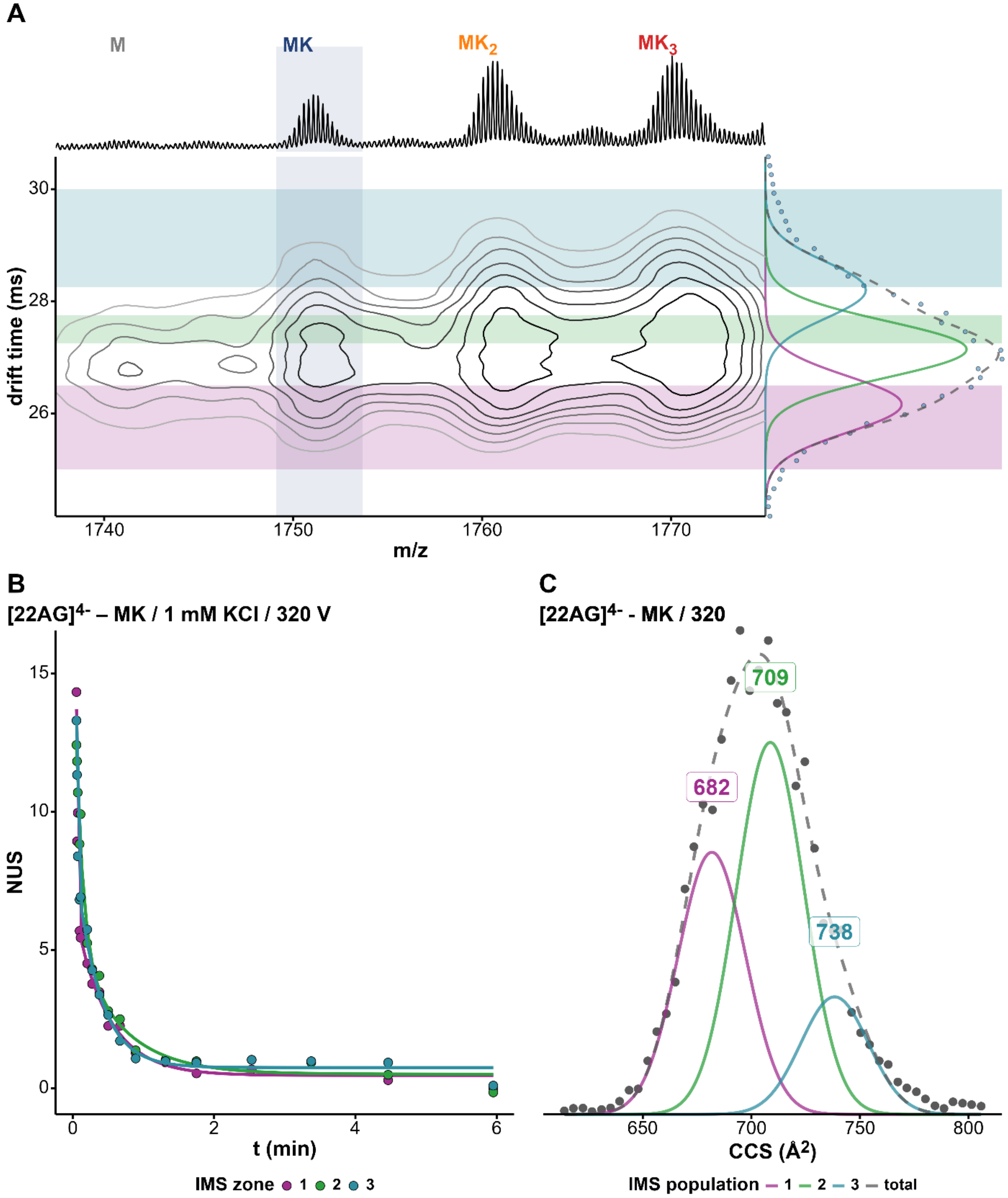


*Figure S104. (A) Density map of 22AG in 100 mM TMAA + 1 mM KCl at fragmentor 320V and charge state 4-. The x-axis represents the mass spectrum and the y-axis the arrival-time distribution (ATD) of the 1K⁺ adduct with the Gaussian fits and the different zones selected for HDX/IMS. (B) HDX/IMS kinetics with colors matching the IMS zones of the density map for the same K⁺ adduct. (C) CCS distribution of the 1K⁺ adduct of 22AG with the CCS values of each population.*

*22AG / $MK_2$ / z = 4- / 1 mM KCl / 320 V*

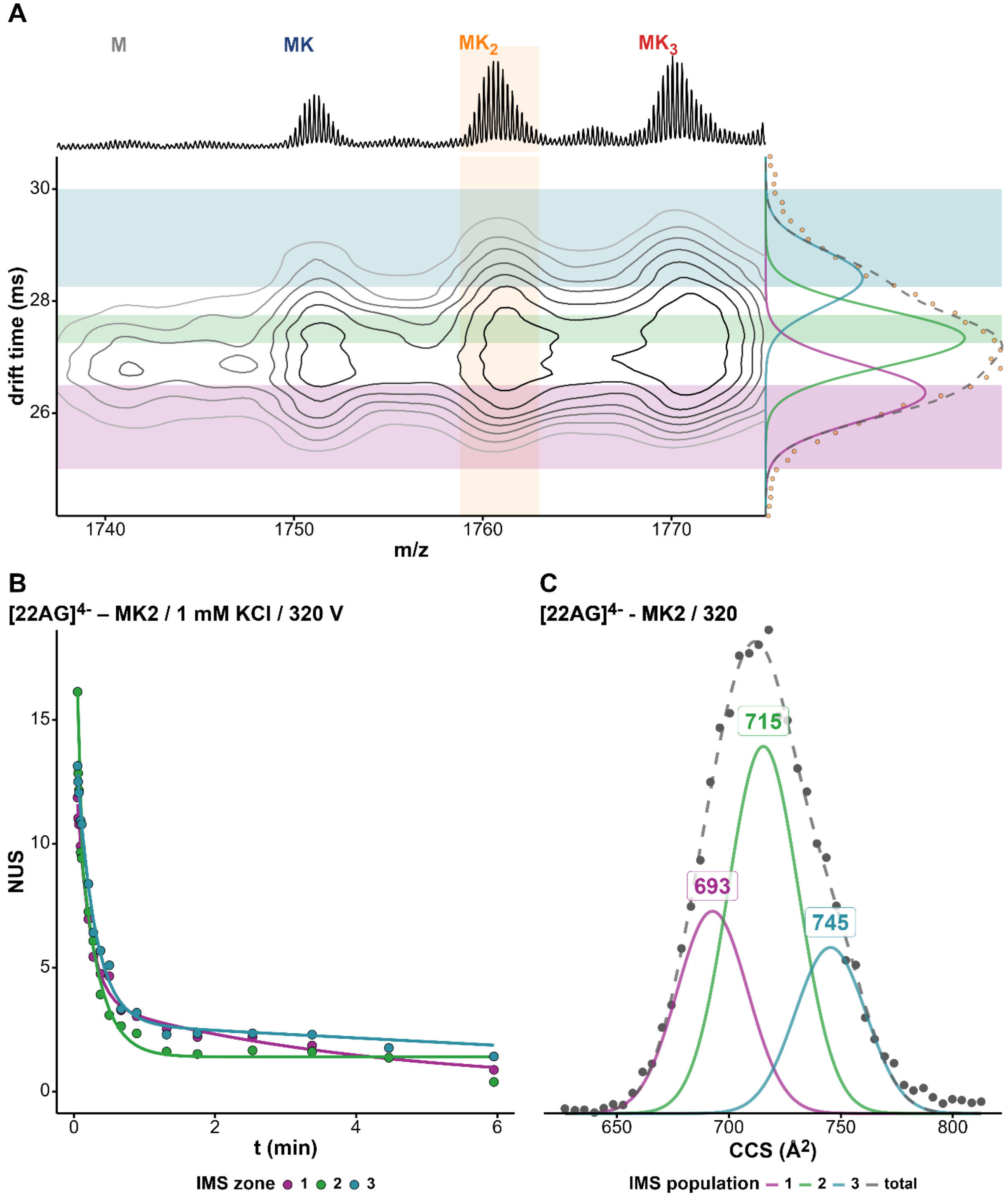


*Figure S105. (A) Density map of 22AG in 100 mM TMAA + 1 mM KCl at fragmentor 320V and charge state 4-. The x-axis represents the mass spectrum and the y-axis the arrival-time distribution (ATD) of the $2K^+$ adduct with the Gaussian fits and the different zones selected for HDX/IMS. (B) HDX/IMS kinetics with colors matching the IMS zones of the density map for the same $K^+$ adduct. (C) CCS distribution of the $2K^+$ adduct of 22AG with the CCS values of each population.*

*22GT / MK / z = 4- / 1 mM KCl / 320 V*

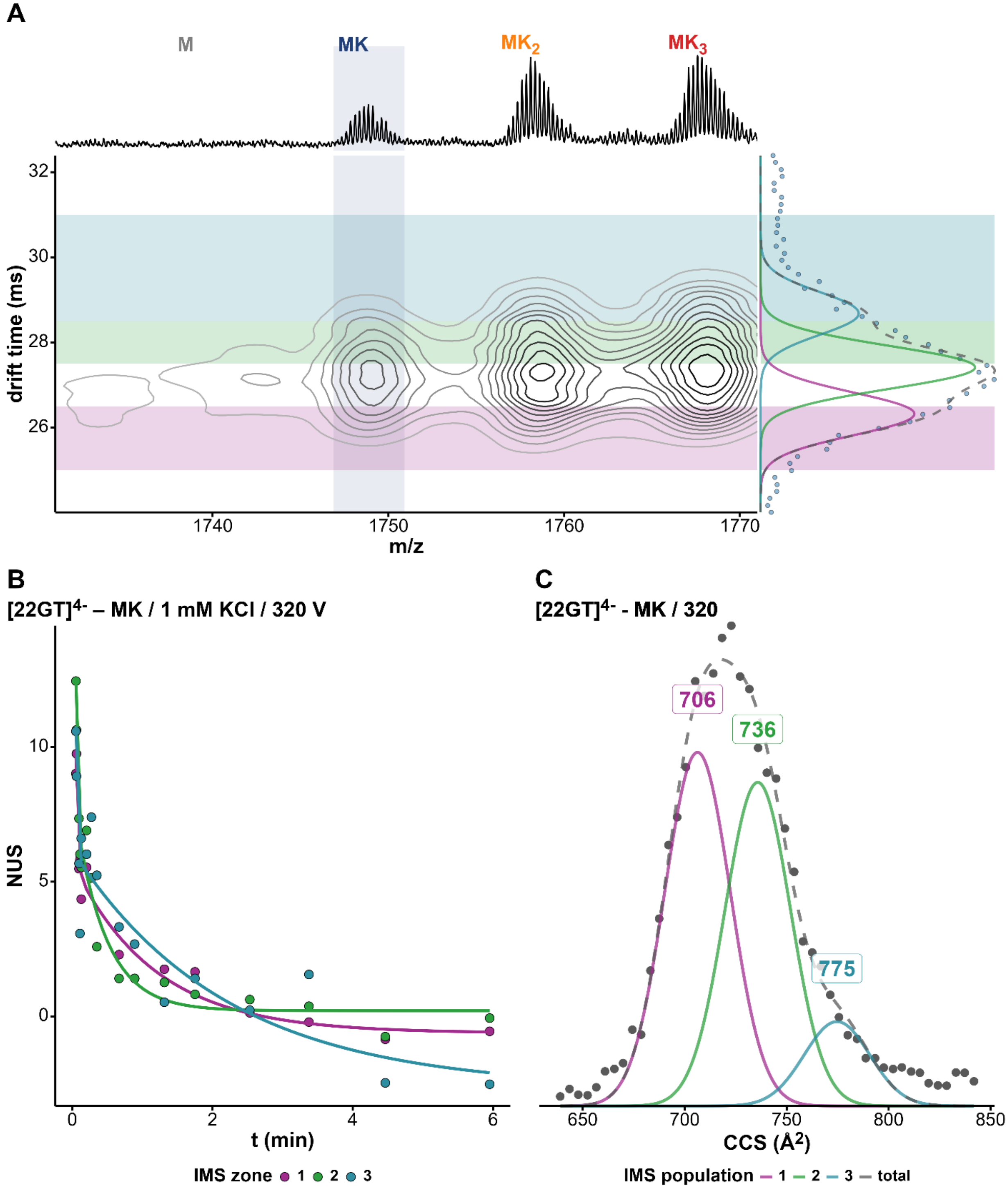


*Figure S106. (A) Density map of 22GT in 100 mM TMAA + 1 mM KCl at fragmentor 320V and charge state 4-. The x-axis represents the mass spectrum and the y-axis the arrival-time distribution (ATD) of the 1K$^+$ adduct with the Gaussian fits and the different zones selected for HDX/IMS. (B) HDX/IMS kinetics with colors matching the IMS zones of the density map for the same K$^+$ adduct. (C) CCS distribution of the 1K$^+$ adduct of 22GT with the CCS values of each population.*

*22GT / MK₂ / z = 4- / 1 mM KCl / 320 V*

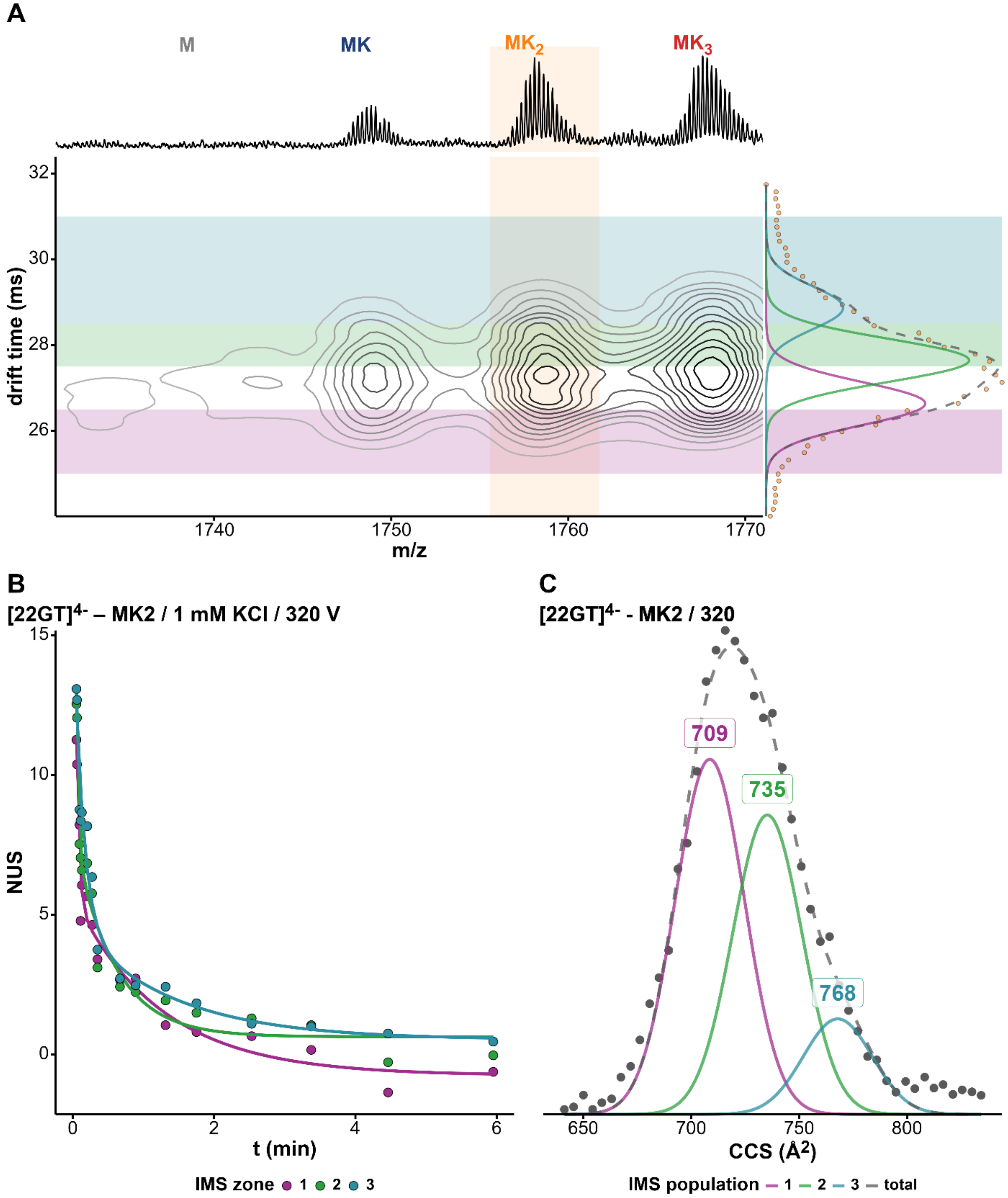


*Figure S107. (A) Density map of 22GT in 100 mM TMAA + 1 mM KCl at fragmentor 320V and charge state 4-. The x-axis represents the mass spectrum and the y-axis the arrival-time distribution (ATD) of the 2K⁺ adduct with the Gaussian fits and the different zones selected for HDX/IMS. (B) HDX/IMS kinetics with colors matching the IMS zones of the density map for the same K⁺ adduct. (C) CCS distribution of the 2K⁺ adduct of 22GT with the CCS values of each population.*

*22GT_18T / MK / z = 4- / 1 mM KCl / 320 V*

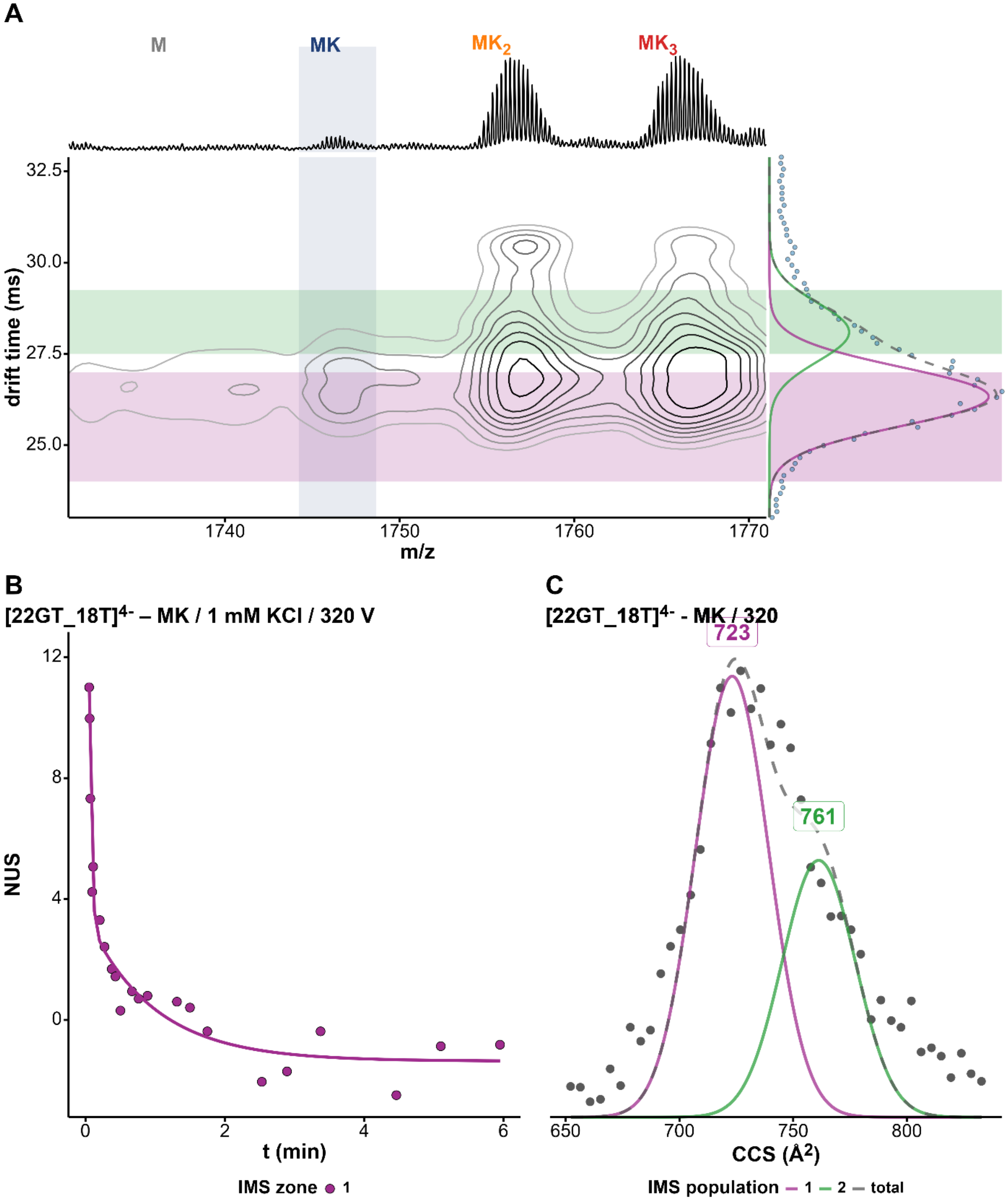


*Figure S108. (A) Density map of 22GT_18T in 100 mM TMAA + 1 mM KCl at fragmentor 320V and charge state 4-. The x-axis represents the mass spectrum and the y-axis the arrival-time distribution (ATD) of the 1K⁺ adduct with the Gaussian fits and the different zones selected for HDX/IMS. (B) HDX/IMS kinetics with colors matching the IMS zones of the density map for the same K⁺ adduct. (C) CCS distribution of the 1K⁺ adduct of 22GT_18T with the CCS values of each population.*

*22GT_18T / MK₂ / z = 4- / 1 mM KCl / 320 V*

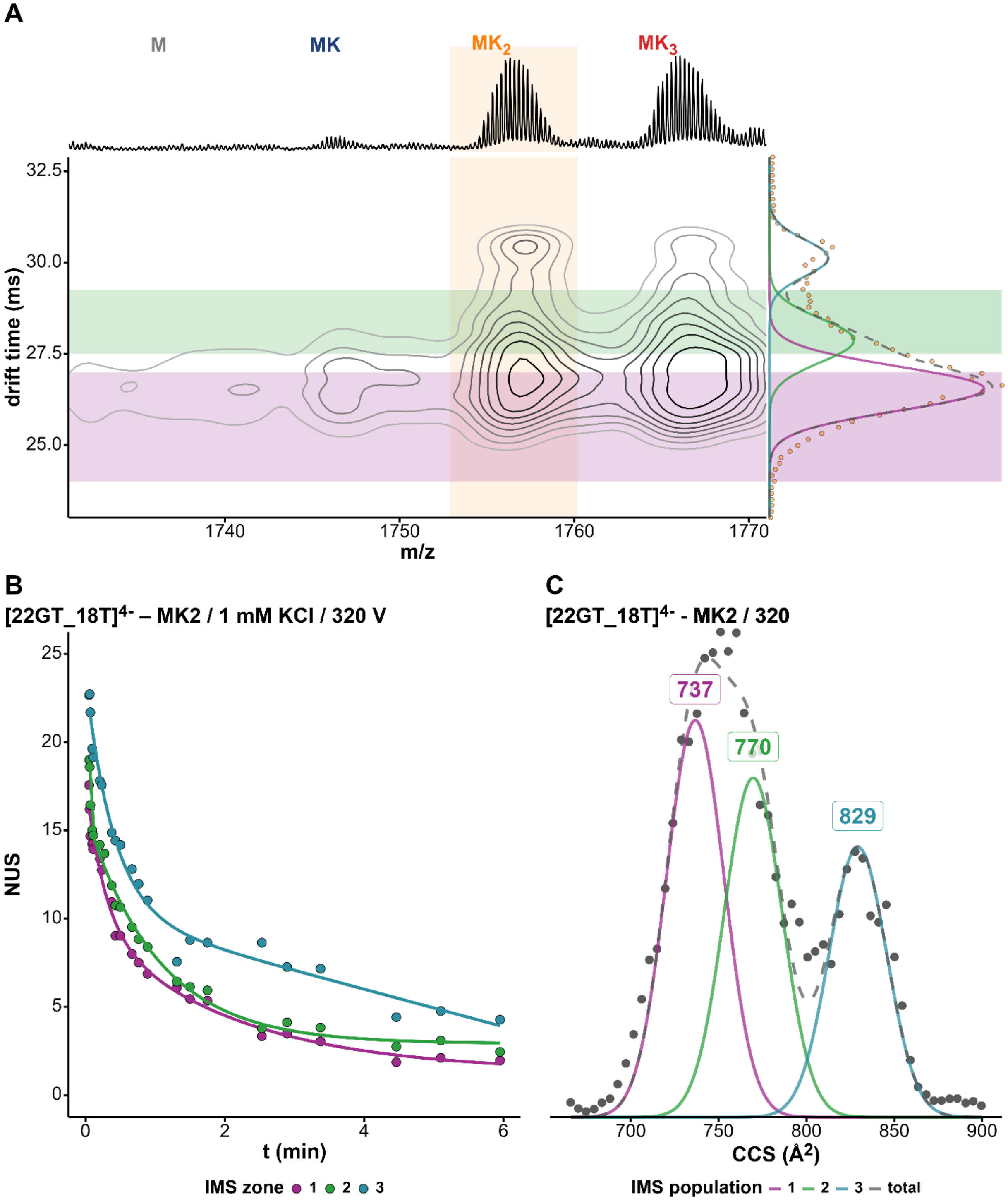


*Figure S109. (A) Density map of 22GT_18T in 100 mM TMAA + 1 mM KCl at fragmentor 320V and charge state 4-. The x-axis represents the mass spectrum and the y-axis the arrival-time distribution (ATD) of the 2K⁺ adduct with the Gaussian fits and the different zones selected for HDX/IMS. (B) HDX/IMS kinetics with colors matching the IMS zones of the density map for the same K⁺ adduct. (C) CCS distribution of the 2K⁺ adduct of 22GT_18T with the CCS values of each population.*

*22CTA / MK / z = 4- / 1 mM KCl / 320 V*

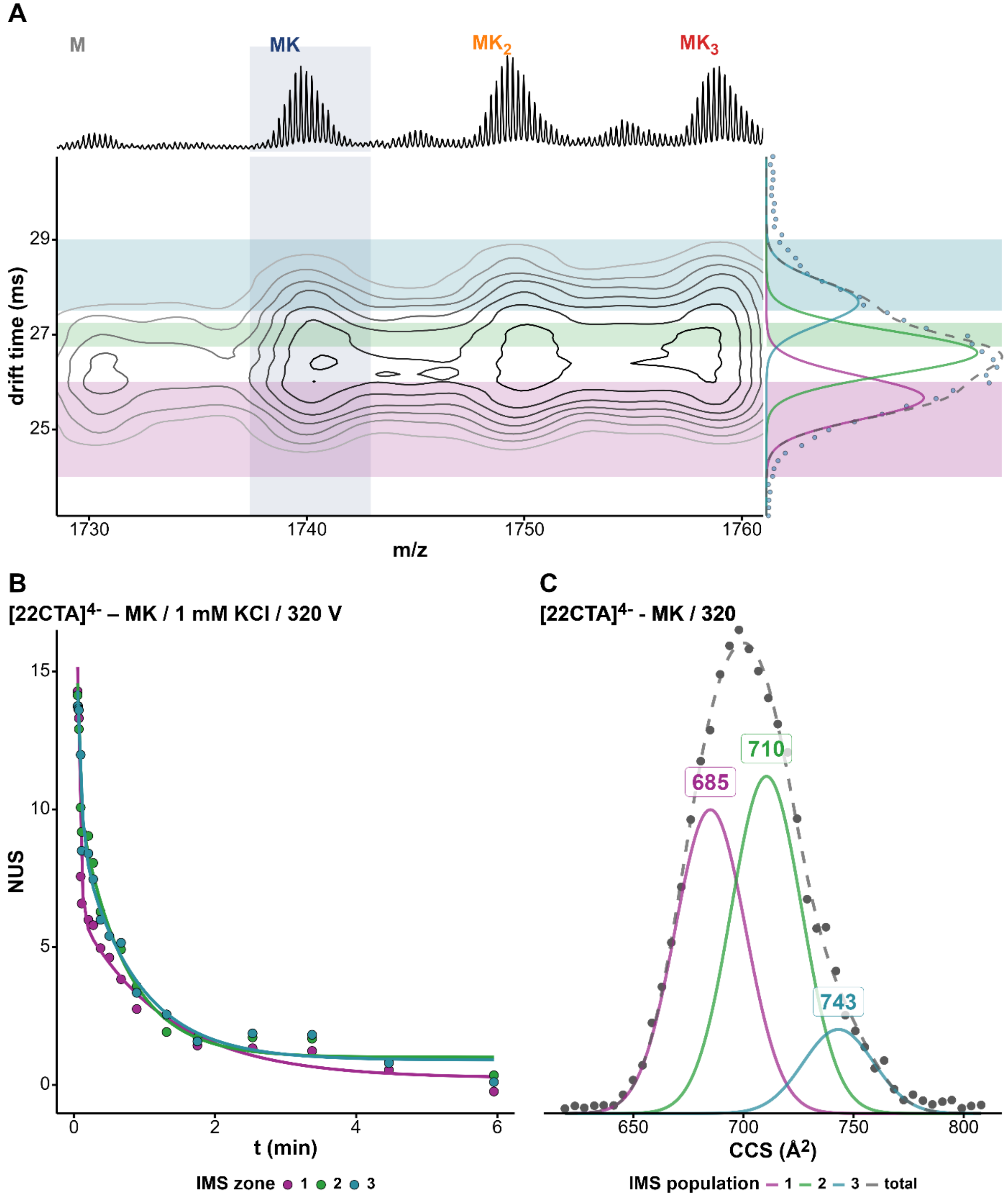


*Figure S110. (A) Density map of 22CTA in 100 mM TMAA + 1 mM KCl at fragmentor 320V and charge state 4-. The x-axis represents the mass spectrum and the y-axis the arrival-time distribution (ATD) of the 1K⁺ adduct with the Gaussian fits and the different zones selected for HDX/IMS. (B) HDX/IMS kinetics with colors matching the IMS zones of the density map for the same K⁺ adduct. (C) CCS distribution of the 1K⁺ adduct of 22CTA with the CCS values of each population.*

*22CTA / MK$_2$ / z = 4- / 1 mM KCl / 320 V*

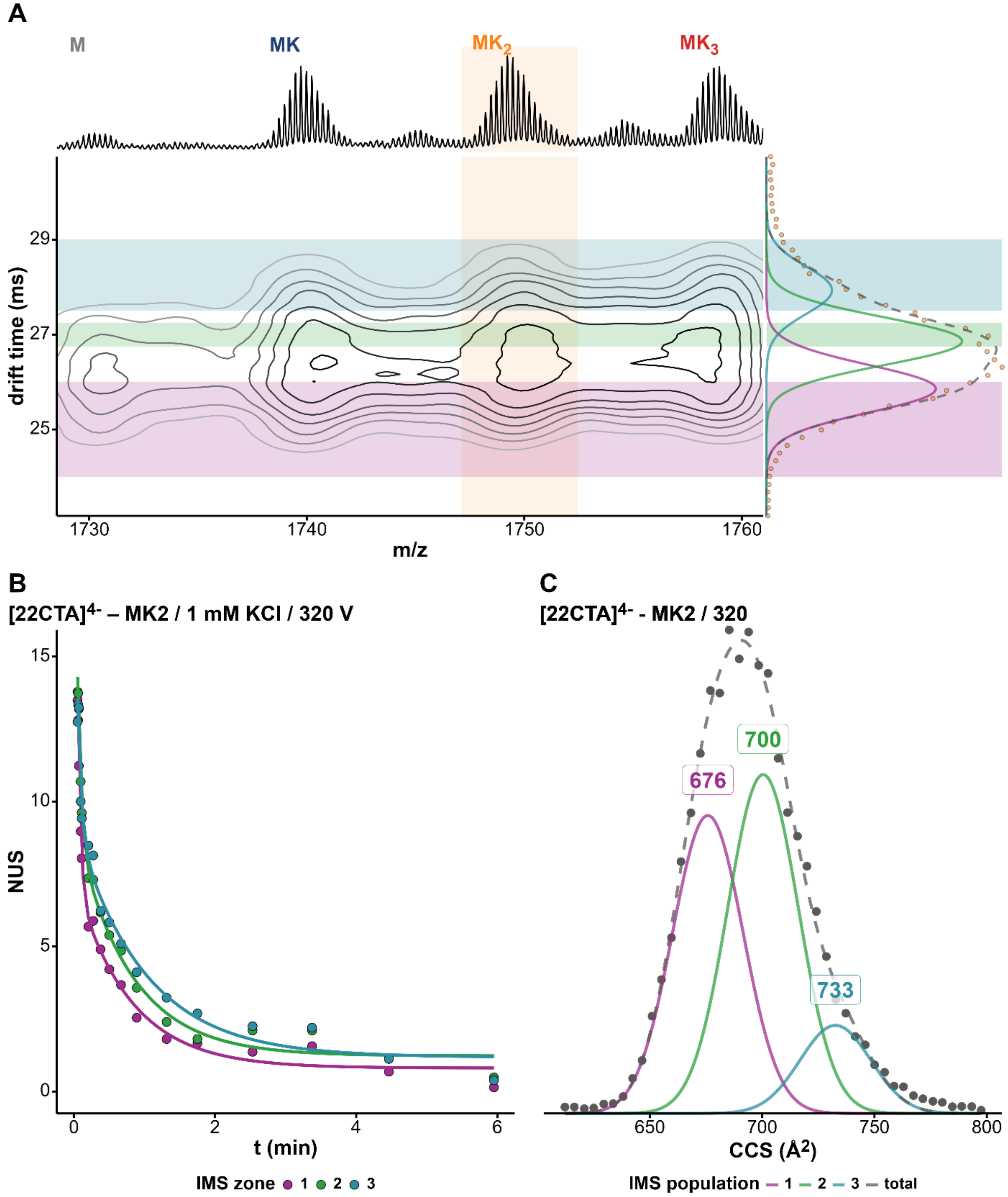


*Figure S111. (A) Density map of 22CTA in 100 mM TMAA + 1 mM KCl at fragmentor 320V and charge state 4-. The x-axis represents the mass spectrum and the y-axis the arrival-time distribution (ATD) of the 2K$^+$ adduct with the Gaussian fits and the different zones selected for HDX/IMS. (B) HDX/IMS kinetics with colors matching the IMS zones of the density map for the same K$^+$ adduct. (C) CCS distribution of the 2K$^+$ adduct of 22CTA with the CCS values of each population.*

*23AG / $MK_2$ / z = 4- / 1 mM KCl / 320 V*

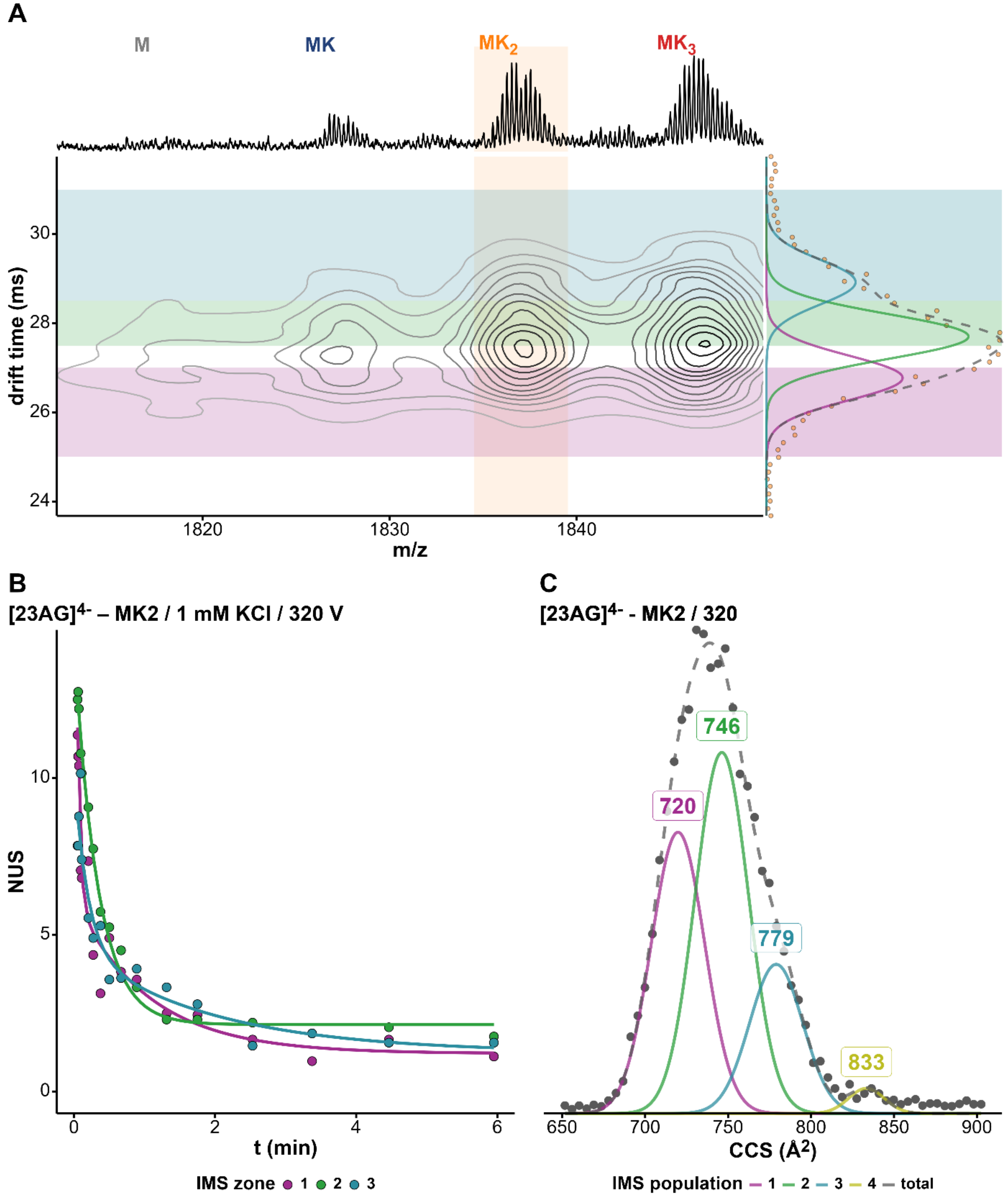


*Figure S112. (A) Density map of 23AG in 100 mM TMAA + 1 mM KCl at fragmentor 320V and charge state 4-. The x-axis represents the mass spectrum and the y-axis the arrival-time distribution (ATD) of the 2K⁺ adduct with the Gaussian fits and the different zones selected for HDX/IMS. (B) HDX/IMS kinetics with colors matching the IMS zones of the density map for the same K⁺ adduct. (C) CCS distribution of the 1K⁺ adduct of 23AG with the CCS values of each population.*

*23AG / $MK_2$ / z = 4- / 1 mM KCl / 370 V*

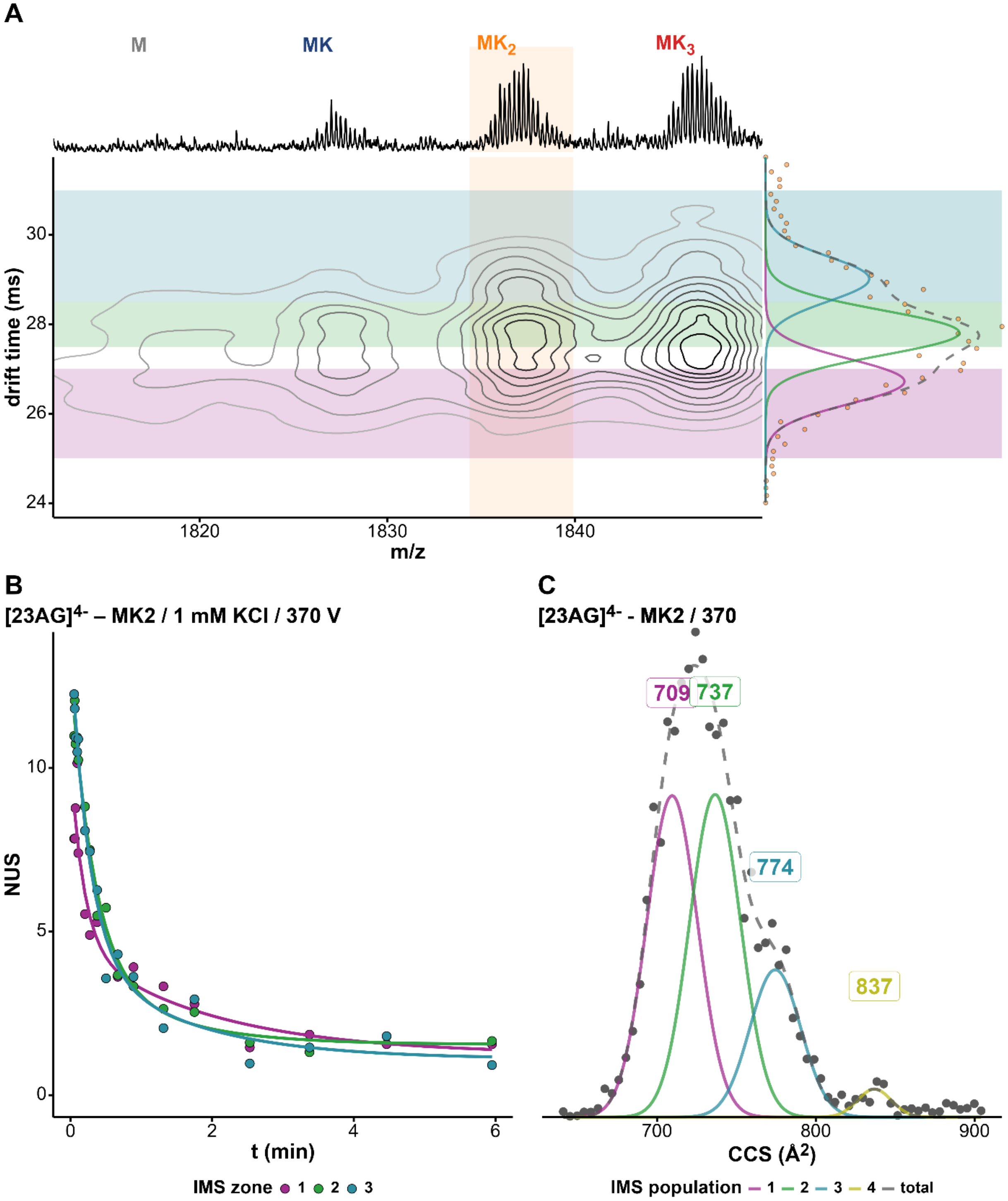


*Figure S113. (A) Density map of 23AG in 100 mM TMAA + 1 mM KCl at fragmentor 370V and charge state 4-. The x-axis represents the mass spectrum and the y-axis the arrival-time distribution (ATD) of the 2K$^+$ adduct with the Gaussian fits and the different zones selected for HDX/IMS. (B) HDX/IMS kinetics with colors matching the IMS zones of the density map for the same K$^+$ adduct. (C) CCS distribution of the 2K$^+$ adduct of 23AG corresponding adduct with the CCS values of each population.*

*23AG_I14 / MK / z = 4- / 1 mM KCl / 320 V*

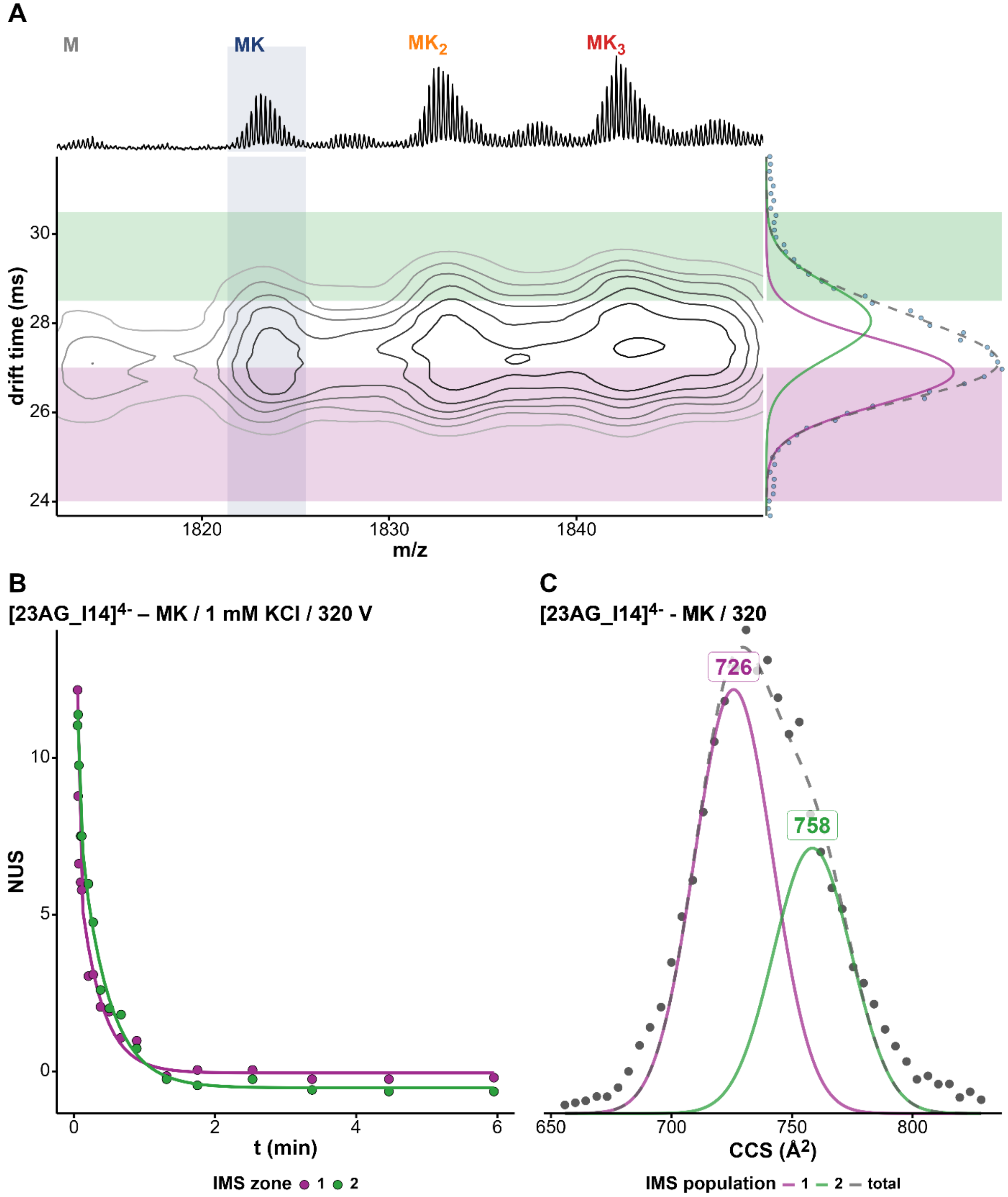


*Figure S114. (A) Density map of 23AG_I14 in 100 mM TMAA + 1 mM KCl at fragmentor 320V and charge state 4-. The x-axis represents the mass spectrum and the y-axis the arrival-time distribution (ATD) of the 1K⁺ adduct with the Gaussian fits and the different zones selected for HDX/IMS. (B) HDX/IMS kinetics with colors matching the IMS zones of the density map for the same K⁺ adduct. (C) CCS distribution of the 1K⁺ adduct of 23AG_I14 with the CCS values of each population.*

*23AG_I14 / $MK_2$ / z = 4- / 1 mM KCl / 320 V*

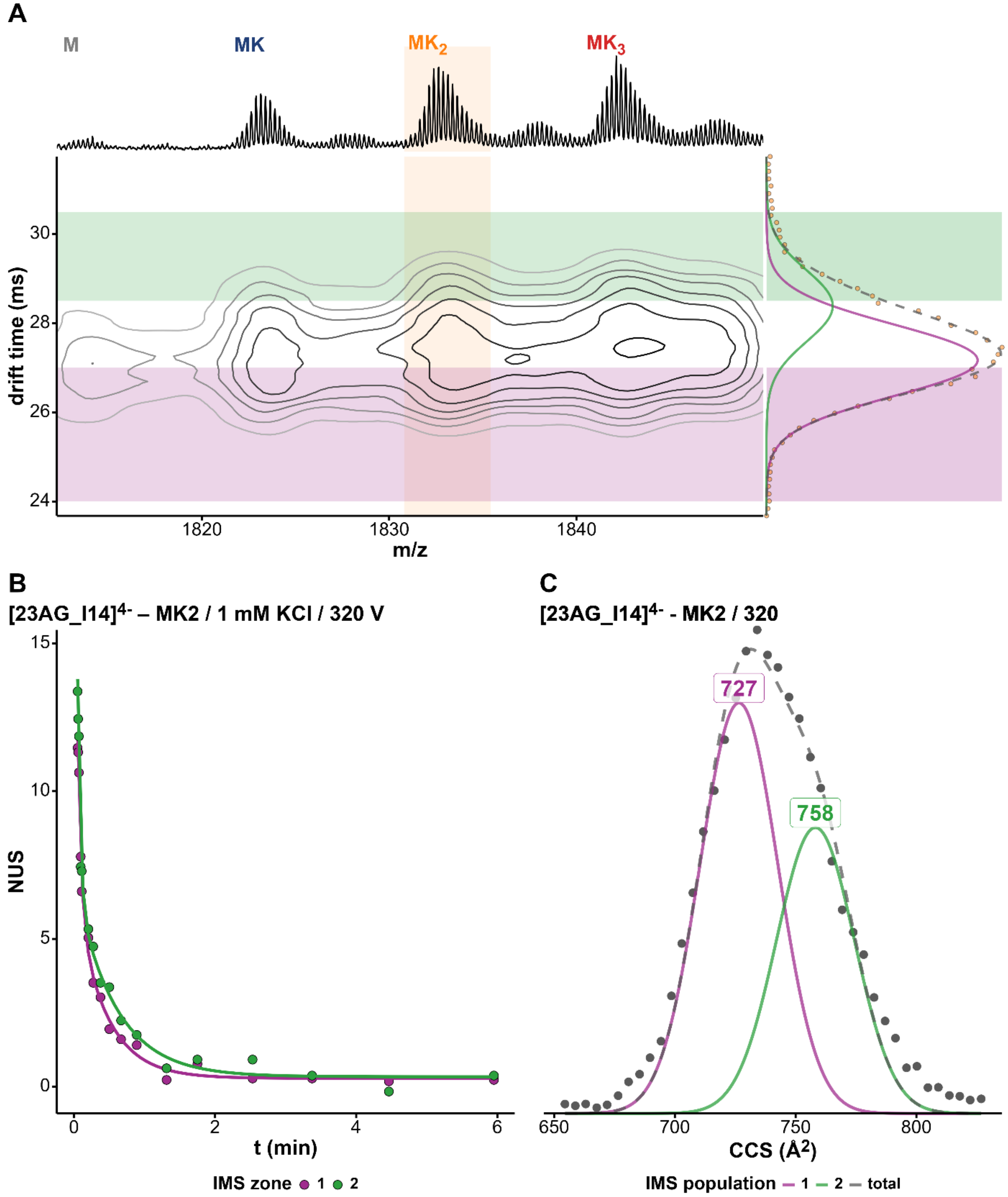


*Figure S115. (A) Density map of 23AG_I14 in 100 mM TMAA + 1 mM KCl at fragmentor 320V and charge state 4-. The x-axis represents the mass spectrum and the y-axis the arrival-time distribution (ATD) of the 2K⁺ adduct with the Gaussian fits and the different zones selected for HDX/IMS. (B) HDX/IMS kinetics with colors matching the IMS zones of the density map for the same K⁺ adduct. (C) CCS distribution of the 2K⁺ adduct of 23AG_I14 with the CCS values of each population.*

*23AG_I14 / MK / z = 4- / 1 mM KCl / 370 V*

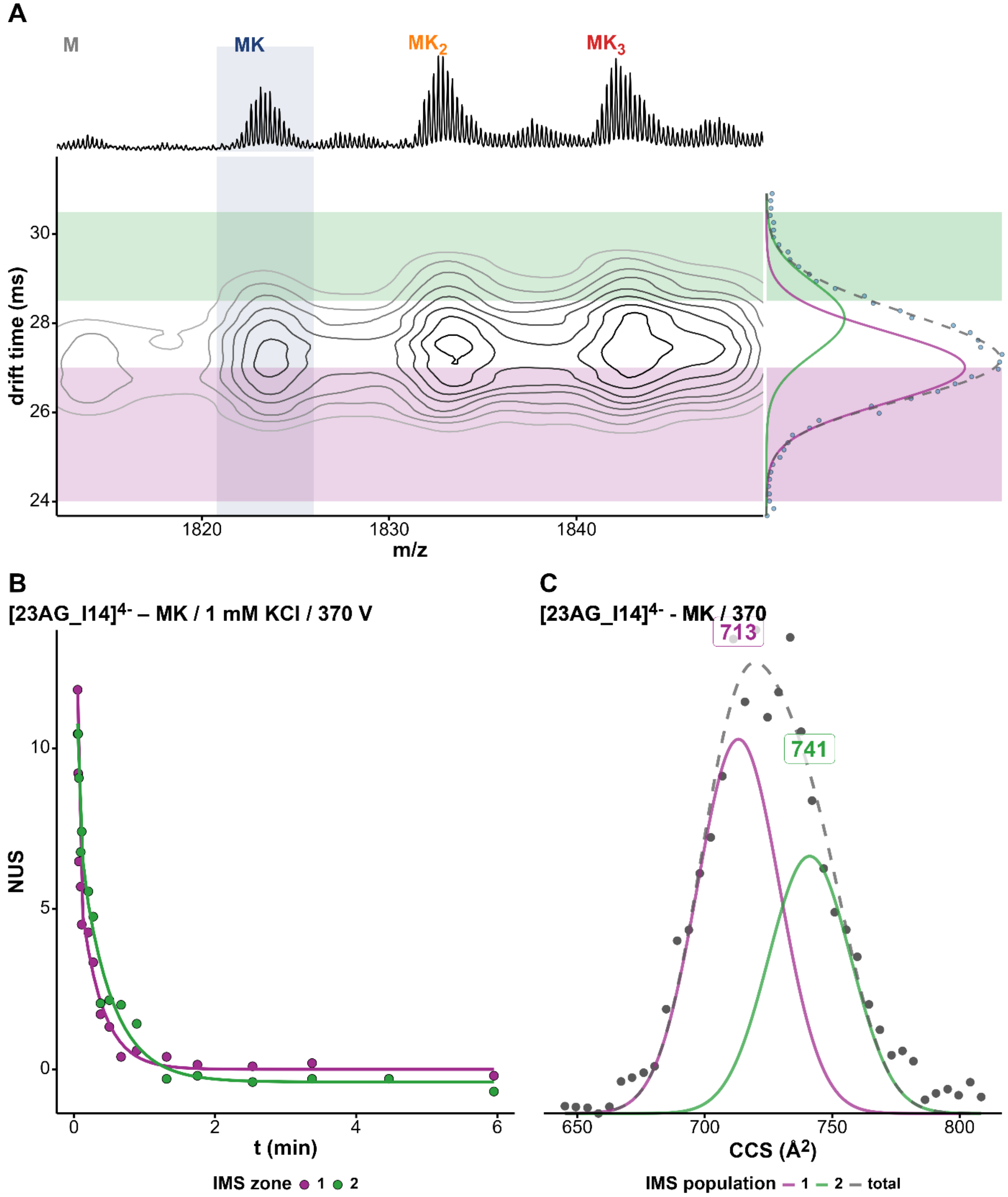


*Figure S116. (A) Density map of 23AG_I14 in 100 mM TMAA + 1 mM KCl at fragmentor 370V and charge state 4-. The x-axis represents the mass spectrum and the y-axis the arrival-time distribution (ATD) of the 1K⁺ adduct with the Gaussian fits and the different zones selected for HDX/IMS. (B) HDX/IMS kinetics with colors matching the IMS zones of the density map for the same K⁺ adduct. (C) CCS distribution of the 1K⁺ adduct of 23AG_I14 with the CCS values of each population.*

*23AG_I14 / $MK_2$ / z = 4- / 1 mM KCl / 320 V*

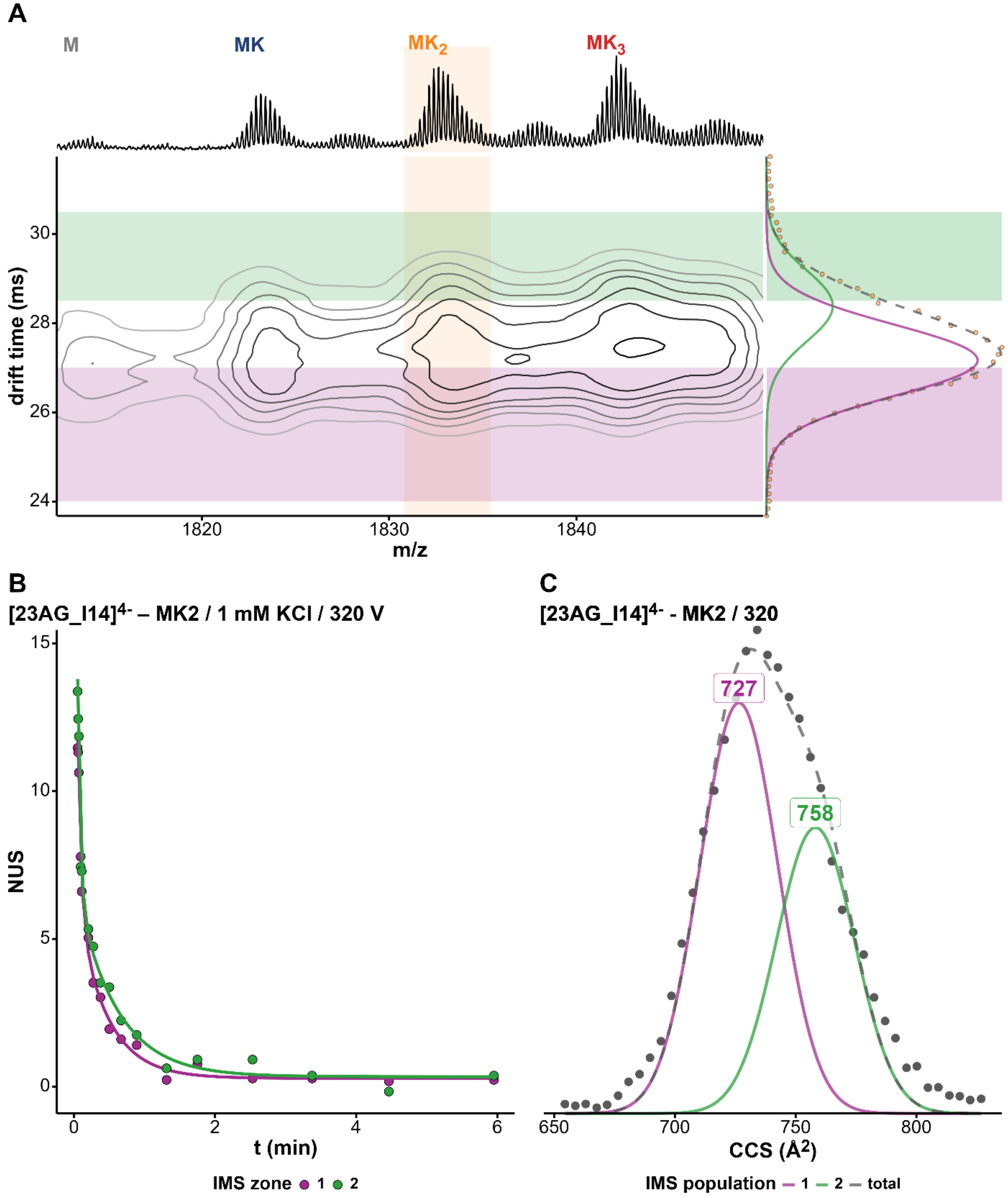


*Figure S117. (A) Density map of 23AG_I14 in 100 mM TMAA + 1 mM KCl at fragmentor 370V and charge state 4-. The x-axis represents the mass spectrum and the y-axis the arrival-time distribution (ATD) of the 2K⁺ adduct with the Gaussian fits and the different zones selected for HDX/IMS. (B) HDX/IMS kinetics with colors matching the IMS zones of the density map for the same K⁺ adduct. (C) CCS distribution of the 2 K⁺ adduct of 23AG_I14 with the CCS values of each population.*

*23TAG / MK / z = 4- / 1 mM KCl / 370 V*

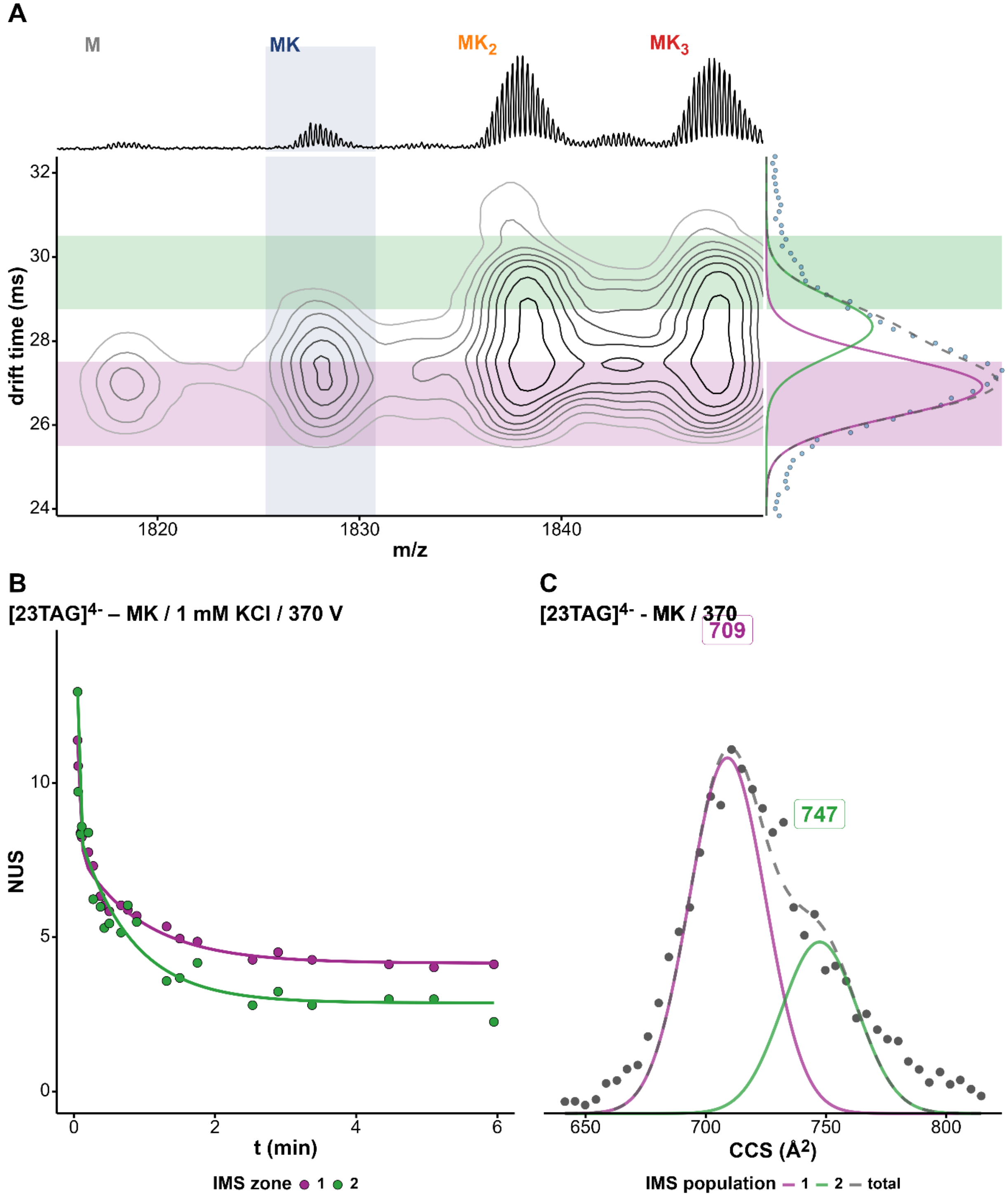


*Figure S118. (A) Density map of 23TAG in 100 mM TMAA + 1 mM KCl at fragmentor 370V and charge state 4-. The x-axis represents the mass spectrum and the y-axis the arrival-time distribution (ATD) of the 1K⁺ adduct with the Gaussian fits and the different zones selected for HDX/IMS. (B) HDX/IMS kinetics with colors matching the IMS zones of the density map for the same K⁺ adduct. (C) CCS distribution of the 1K⁺ adduct of 23TAG with the CCS values of each population.*

*23TAG / $MK_2$ / z = 4- / 1 mM KCl / 320 V*

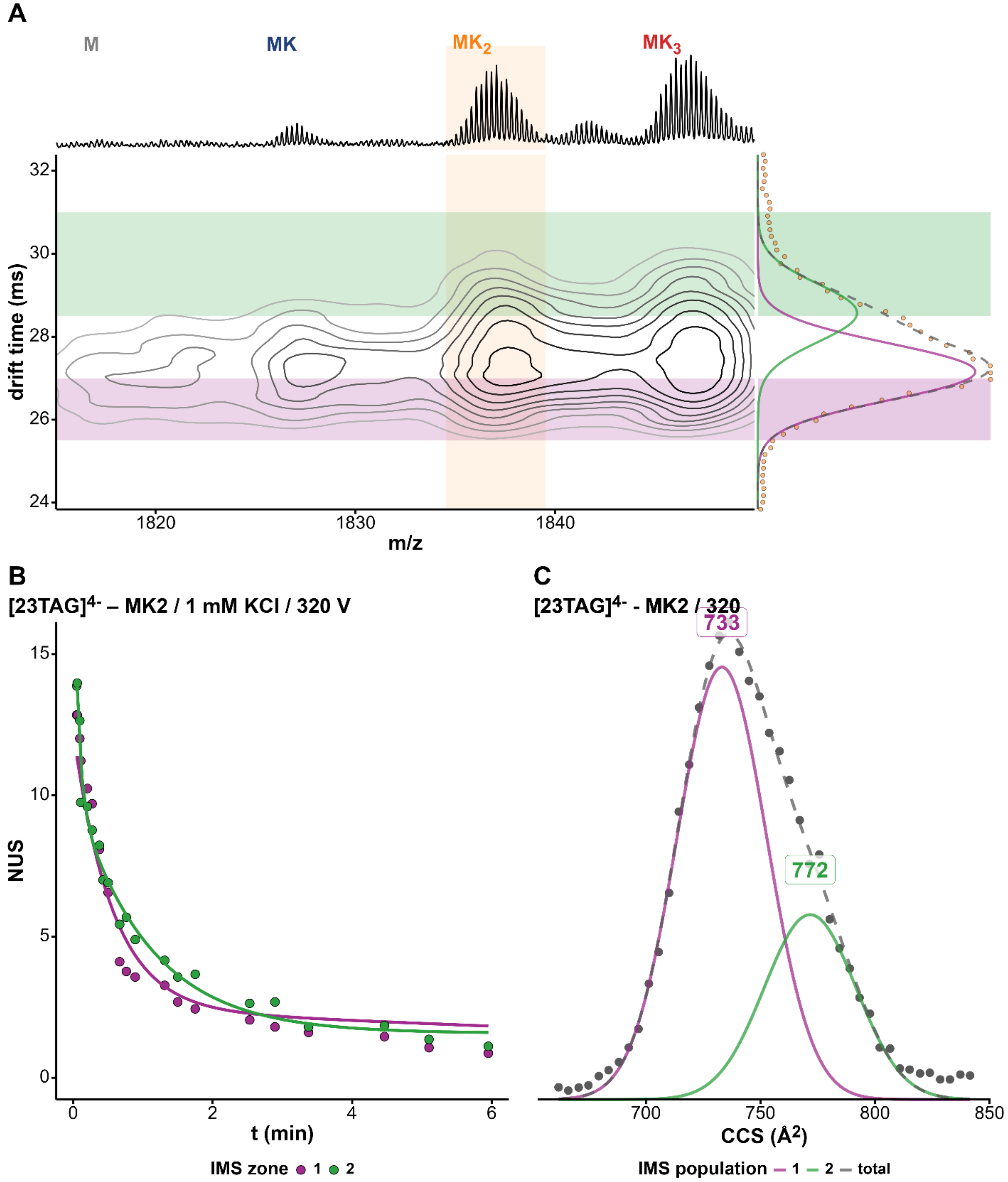


*Figure S119. (A) Density map of 23TAG in 100 mM TMAA + 1 mM KCl at fragmentor 320V and charge state 4-. The x-axis represents the mass spectrum and the y-axis the arrival-time distribution (ATD) of the nK⁺ adduct with the Gaussian fits and the different zones selected for HDX/IMS. (B) HDX/IMS kinetics with colors matching the IMS zones of the density map for the same K⁺ adduct. (C) CCS distribution of the 2K⁺ adduct of 23TAG with the CCS values of each population.*

*23TAG / $MK_2$ / z = 4- / 1 mM KCl / 350 V*

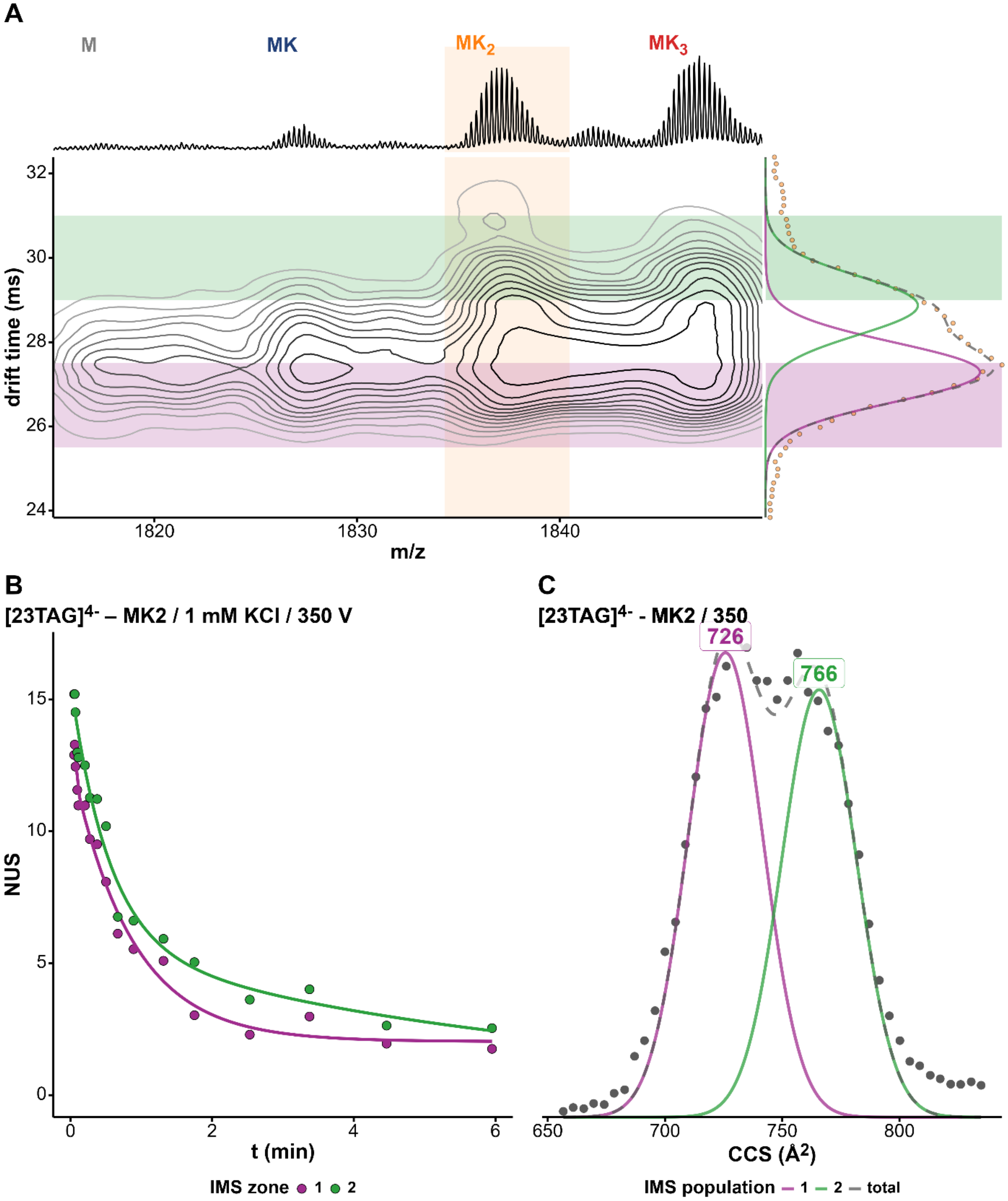


*Figure S120. (A) Density map of 23TAG in 100 mM TMAA + 1 mM KCl at fragmentor 350V and charge state 4-. The x-axis represents the mass spectrum and the y-axis the arrival-time distribution (ATD) of the 2K⁺ adduct with the Gaussian fits and the different zones selected for HDX/IMS. (B) HDX/IMS kinetics with colors matching the IMS zones of the density map for the same K⁺ adduct. (C) CCS distribution of the 2K⁺ adduct of 23TAG with the CCS values of each population.*

*23TAG / MK₂ / z = 4- / 1 mM KCl / 370 V*

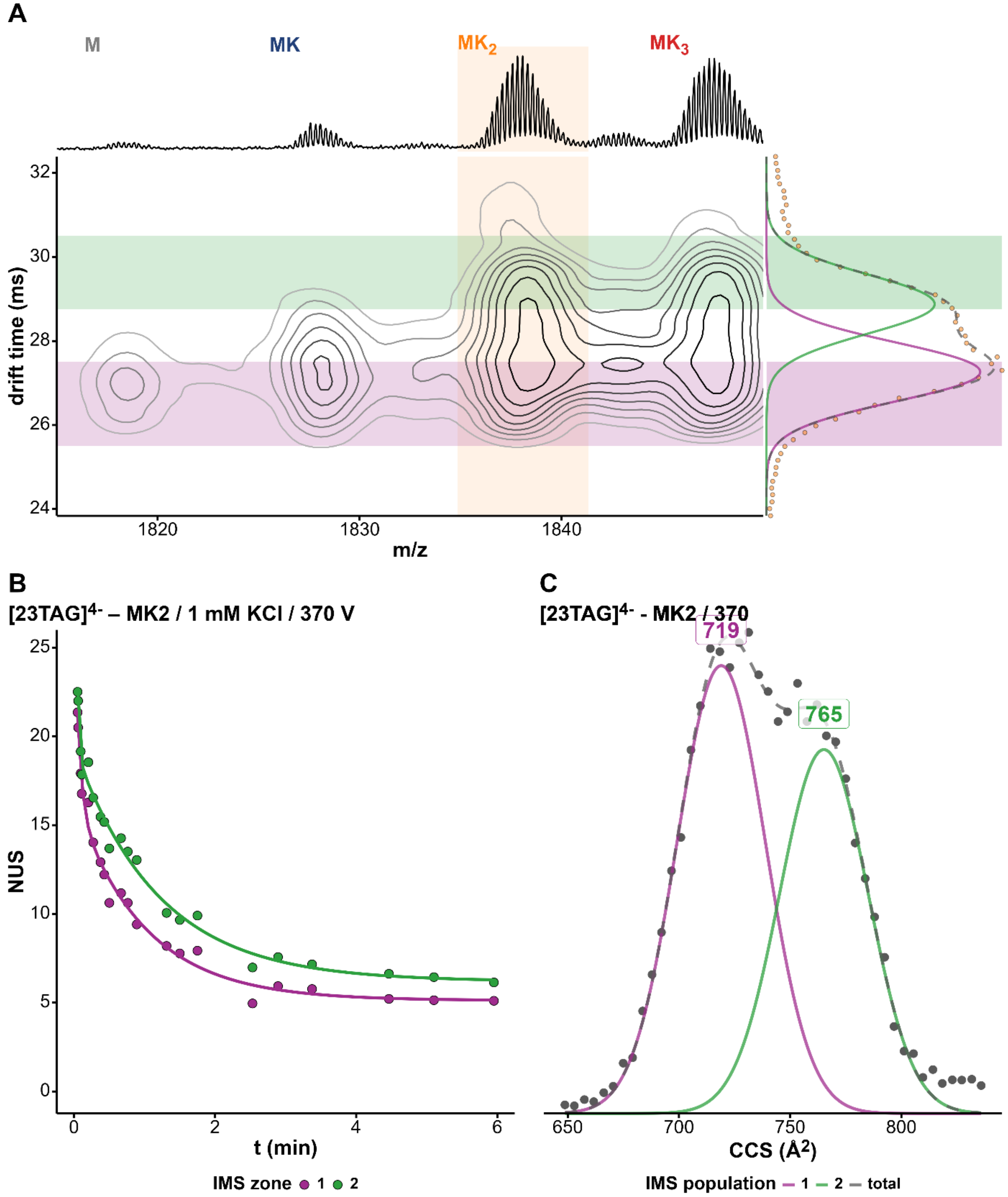


*Figure S121. (A) Density map of 23TAG in 100 mM TMAA + 1 mM KCl at fragmentor 370V and charge state 4-. The x-axis represents the mass spectrum and the y-axis the arrival-time distribution (ATD) of the 2K$^+$ adduct with the Gaussian fits and the different zones selected for HDX/IMS. (B) HDX/IMS kinetics with colors matching the IMS zones of the density map for the same K$^+$ adduct. (C) CCS distribution of the 2K$^+$ adduct of 23TAG with the CCS values of each population.*

### *24TTG / $MK_2$ / z = 4- / 1 mM KCl / 320 V*

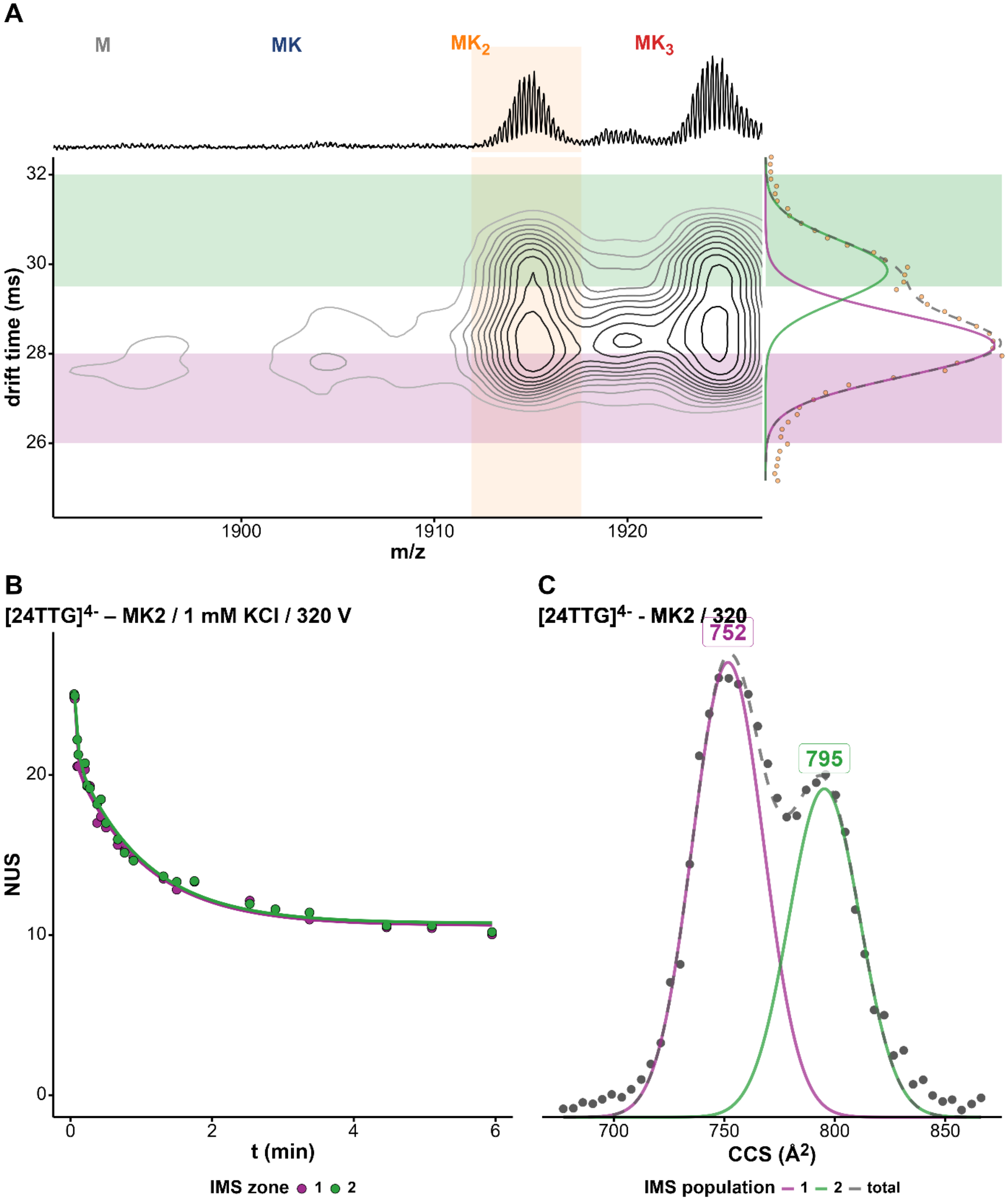


*Figure S122. (A) Density map of 24TTG in 100 mM TMAA + 1 mM KCl at fragmentor 320V and charge state 4-. The x-axis represents the mass spectrum and the y-axis the arrival-time distribution (ATD) of the 2$K^+$ adduct with the Gaussian fits and the different zones selected for HDX/IMS. (B) HDX/IMS kinetics with colors matching the IMS zones of the density map for the same $K^+$ adduct. (C) CCS distribution of the 2$K^+$ adduct of 24TTG with the CCS values of each population.*

*24TTG / $MK_2$ / z = 4- / 0.5 mM KCl / 320 V*

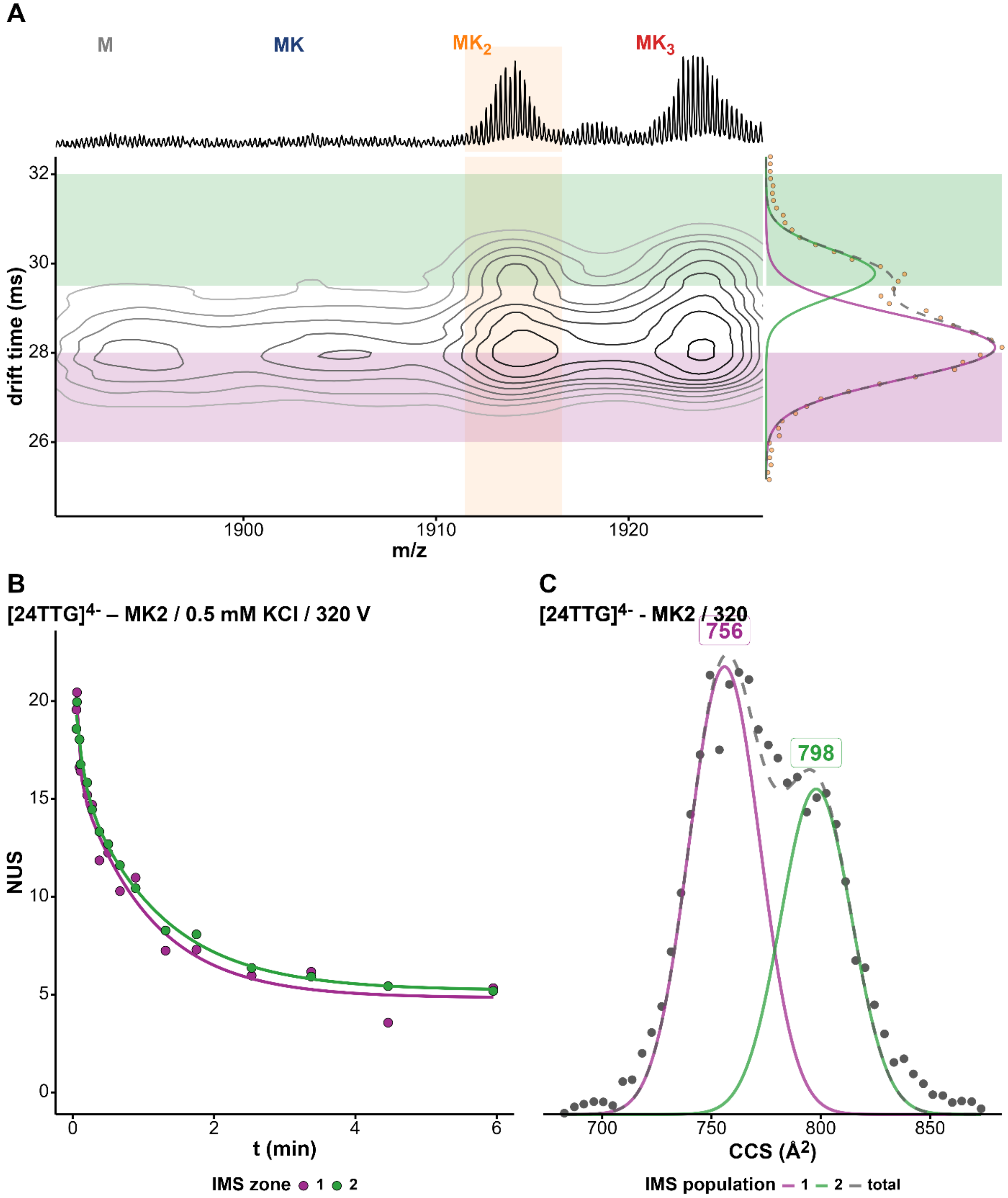


*Figure S123. (A) Density map of 24TTG in 100 mM TMAA + 0.5 mM KCl at fragmentor 320V and charge state 4-. The x-axis represents the mass spectrum and the y-axis the arrival-time distribution (ATD) of the 2K⁺ adduct with the Gaussian fits and the different zones selected for HDX/IMS. (B) HDX/IMS kinetics with colors matching the IMS zones of the density map for the same K⁺ adduct. (C) CCS distribution of the 2K⁺ adduct of 24TTG with the CCS values of each population.*

*24TTG / MK₂ / z = 4- / 0.1 mM KCl / 320 V*

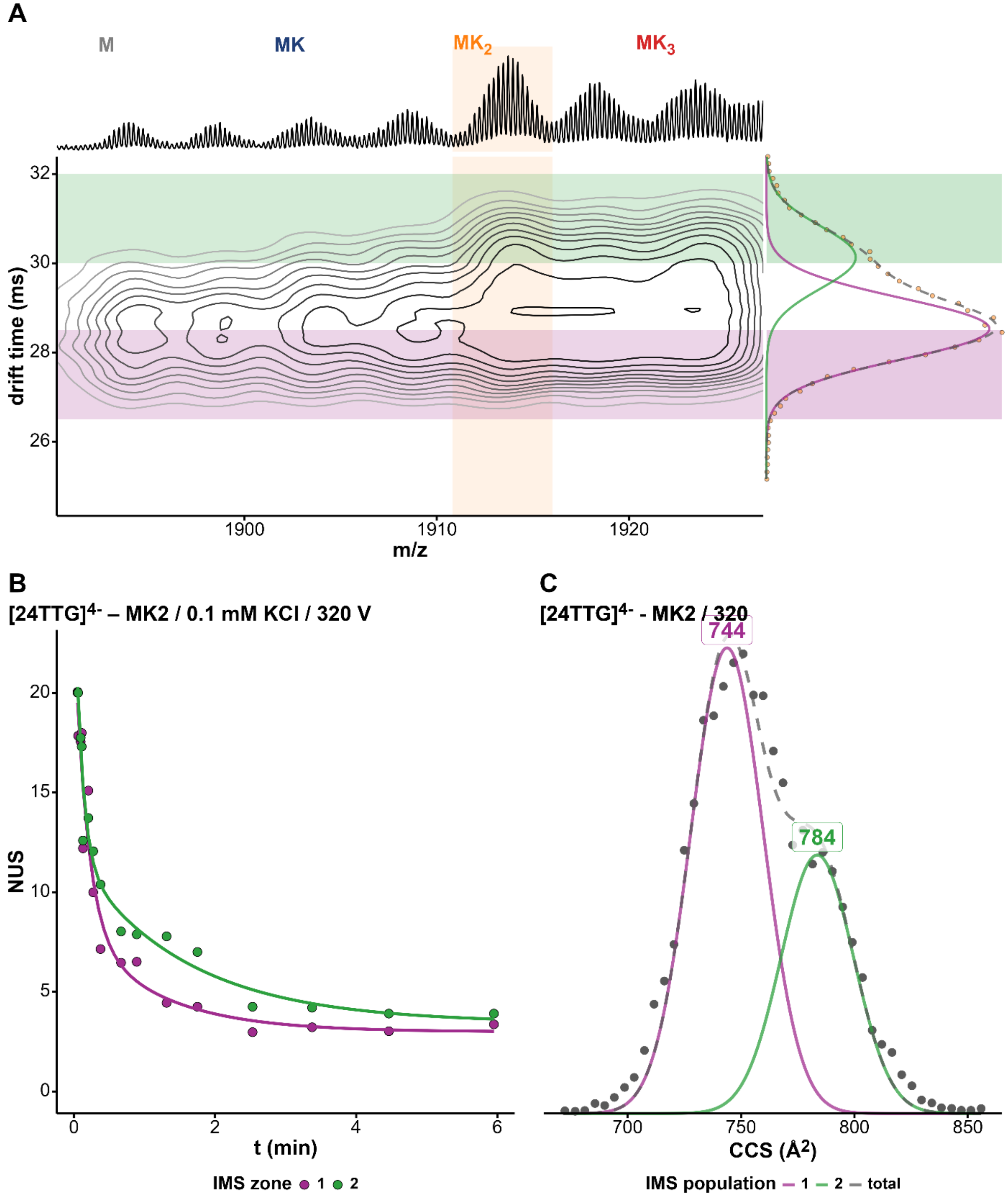


*Figure S124. (A) Density map of 24TTG in 100 mM TMAA + 0.1 mM KCl at fragmentor 320V and charge state 4-. The x-axis represents the mass spectrum and the y-axis the arrival-time distribution (ATD) of the 2K⁺ adduct with the Gaussian fits and the different zones selected for HDX/IMS. (B) HDX/IMS kinetics with colors matching the IMS zones of the density map for the same K⁺ adduct. (C) CCS distribution of the 2K⁺ adduct of 24TTG with the CCS values of each population.*

*222T / MK / z = 4- / 0.1 mM KCl / 320 V*

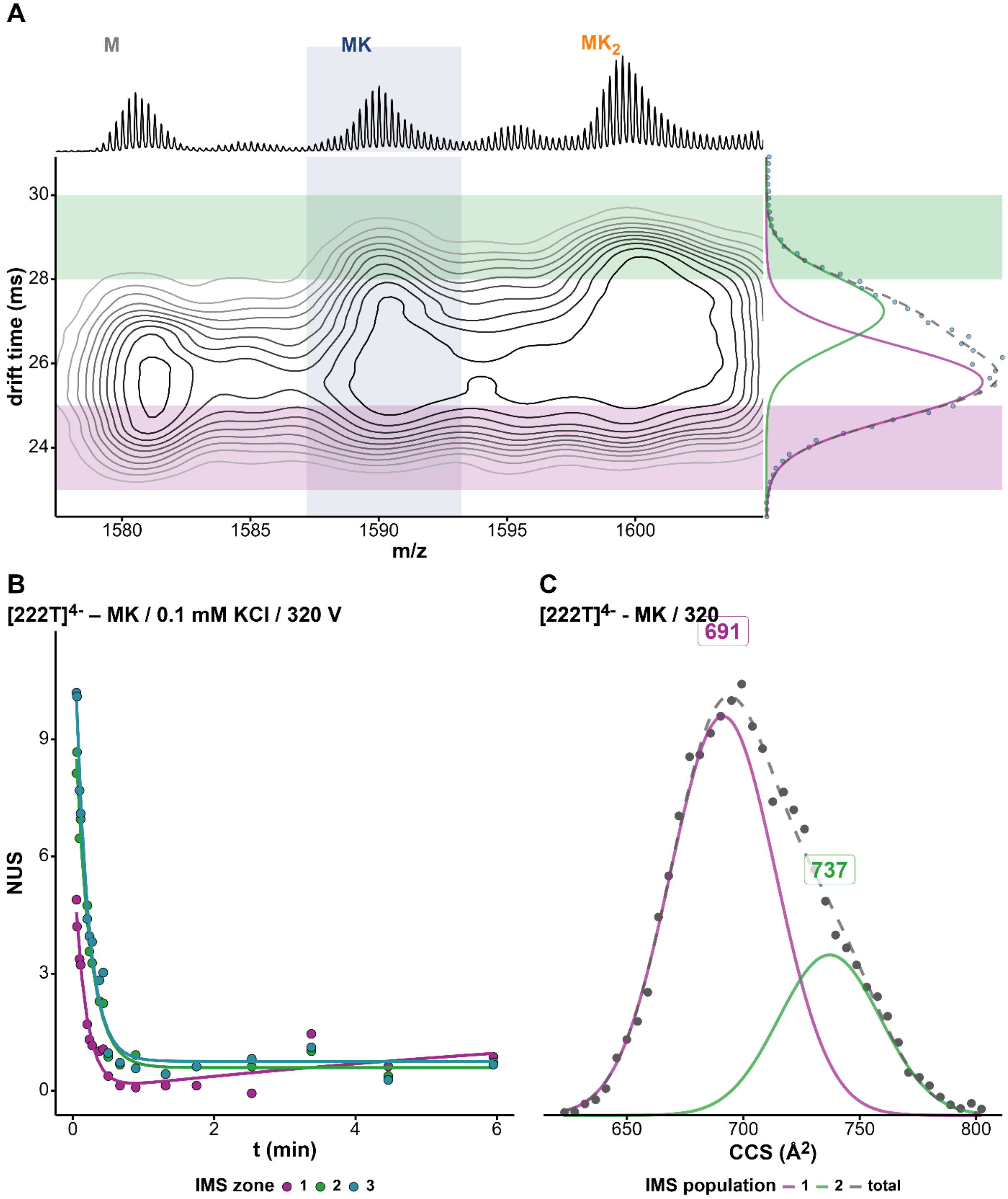


*Figure S125. (A) Density map of 222T in 100 mM TMAA + 0.1 mM KCl at fragmentor 320V and charge state 4-. The x-axis represents the mass spectrum and the y-axis the arrival-time distribution (ATD) of the 1K⁺ adduct with the Gaussian fits and the different zones selected for HDX/IMS. (B) HDX/IMS kinetics with colors matching the IMS zones of the density map for the same K⁺ adduct. (C) CCS distribution of the 1K⁺ adduct of 222T with the CCS values of each population.*

*222T / MK₂ / z = 4- / 0.1 mM KCl / 320 V*

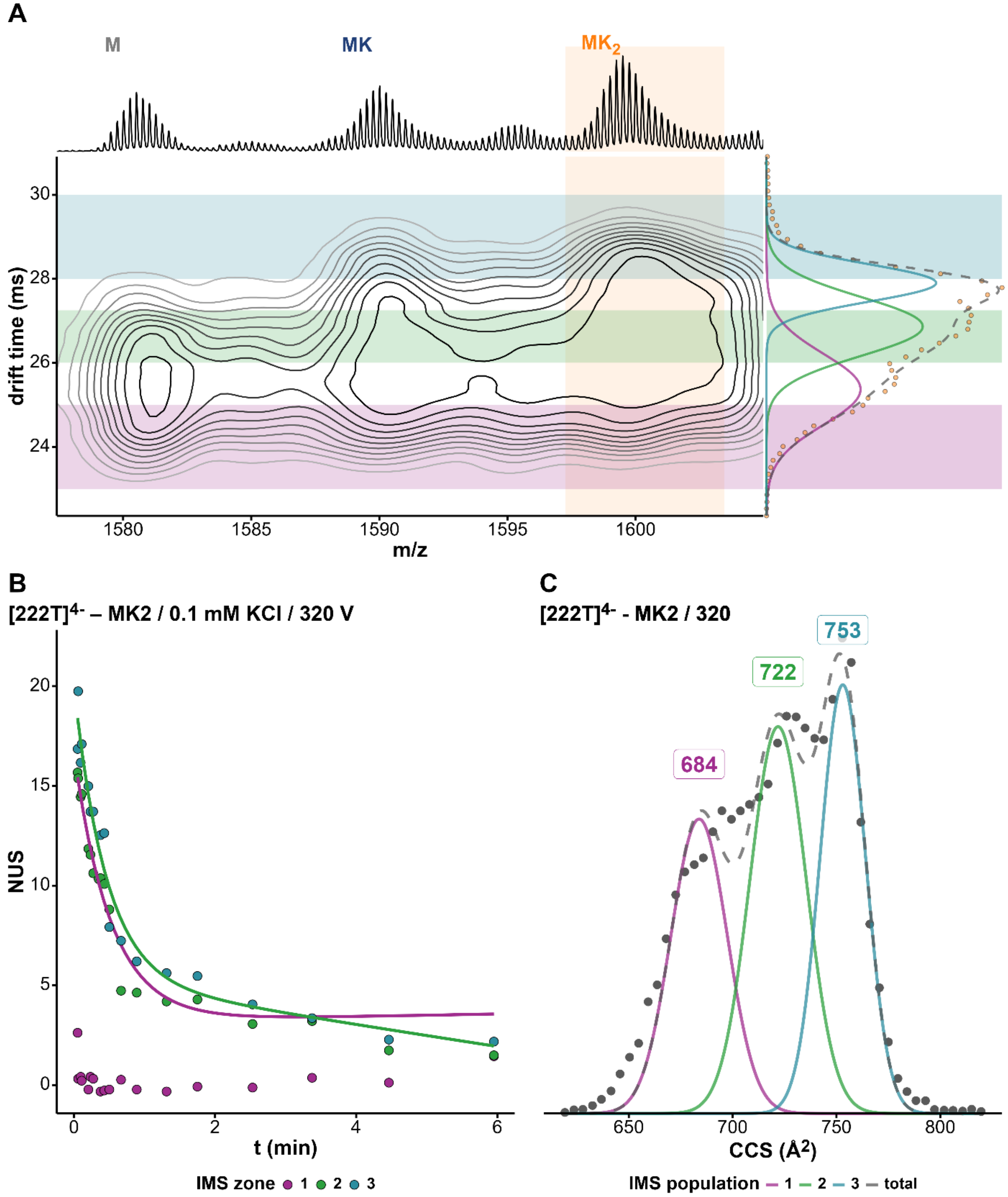


*Figure S126. (A) Density map of 222T in 100 mM TMAA + 0.1 mM KCl at fragmentor 320V and charge state 4-. The x-axis represents the mass spectrum and the y-axis the arrival-time distribution (ATD) of the 2K⁺ adduct with the Gaussian fits and the different zones selected for HDX/IMS. (B) HDX/IMS kinetics with colors matching the IMS zones of the density map for the same K⁺ adduct. (C) CCS distribution of the 2K⁺ adduct of 222T with the CCS values of each population.*

*222T / $MK_2$ / z = 4- / 1 mM KCl / 320 V*

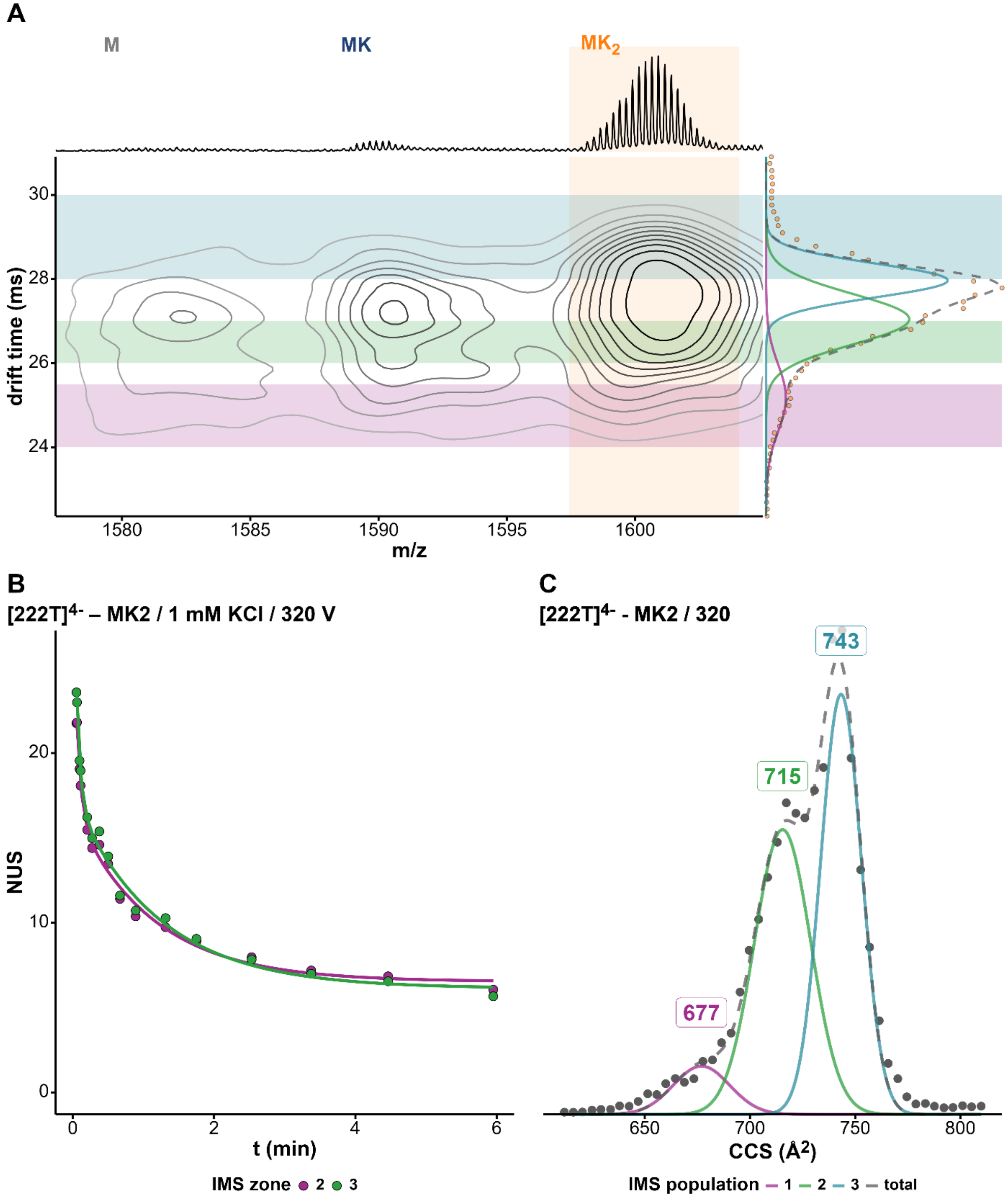


*Figure S127. (A) Density map of 222T in 100 mM TMAA + 1 mM KCl at fragmentor 320V and charge state 4-. The x-axis represents the mass spectrum and the y-axis the arrival-time distribution (ATD) of the $2K^+$ adduct with the Gaussian fits and the different zones selected for HDX/IMS. (B) HDX/IMS kinetics with colors matching the IMS zones of the density map for the same $K^+$ adduct. (C) CCS distribution of the $2K^+$ adduct of 222T with the CCS values of each population.*

### *ckit87up / $MK_2$ / z = 4- / 1 mM KCl / 320 V*

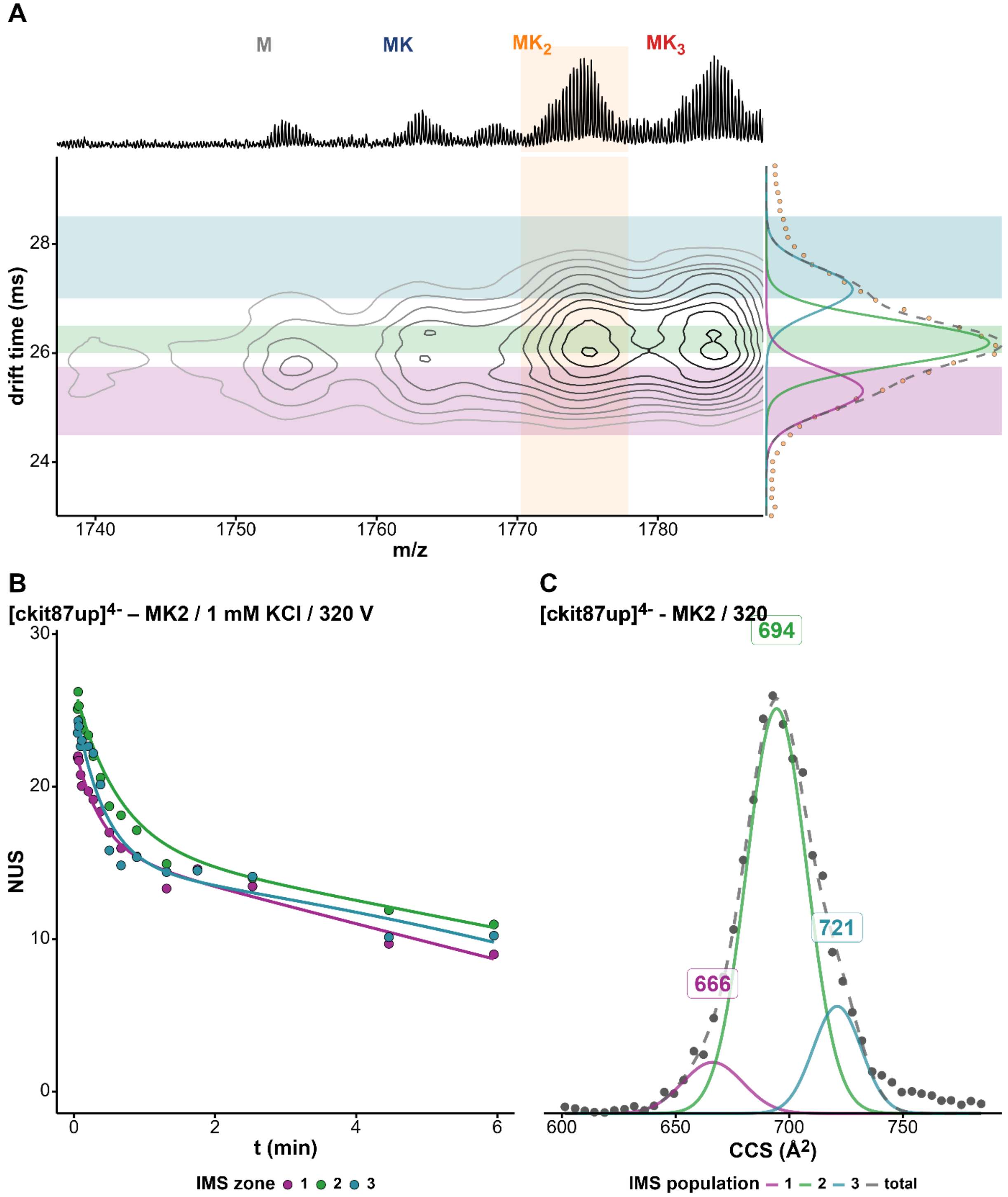


*Figure S128. (A) Density map of ckit87up in 100 mM TMAA + 1 mM KCl at fragmentor 320V and charge state 4-. The x-axis represents the mass spectrum and the y-axis the arrival-time distribution (ATD) of the 2K⁺ adduct with the Gaussian fits and the different zones selected for HDX/IMS. (B) HDX/IMS kinetics with colors matching the IMS zones of the density map for the same K⁺ adduct. (C) CCS distribution of the 2K⁺ adduct of ckit87up with the CCS values of each population.*

*BCl2 / $MK_2$ / z = 4- / 1 mM KCl / 320 V*

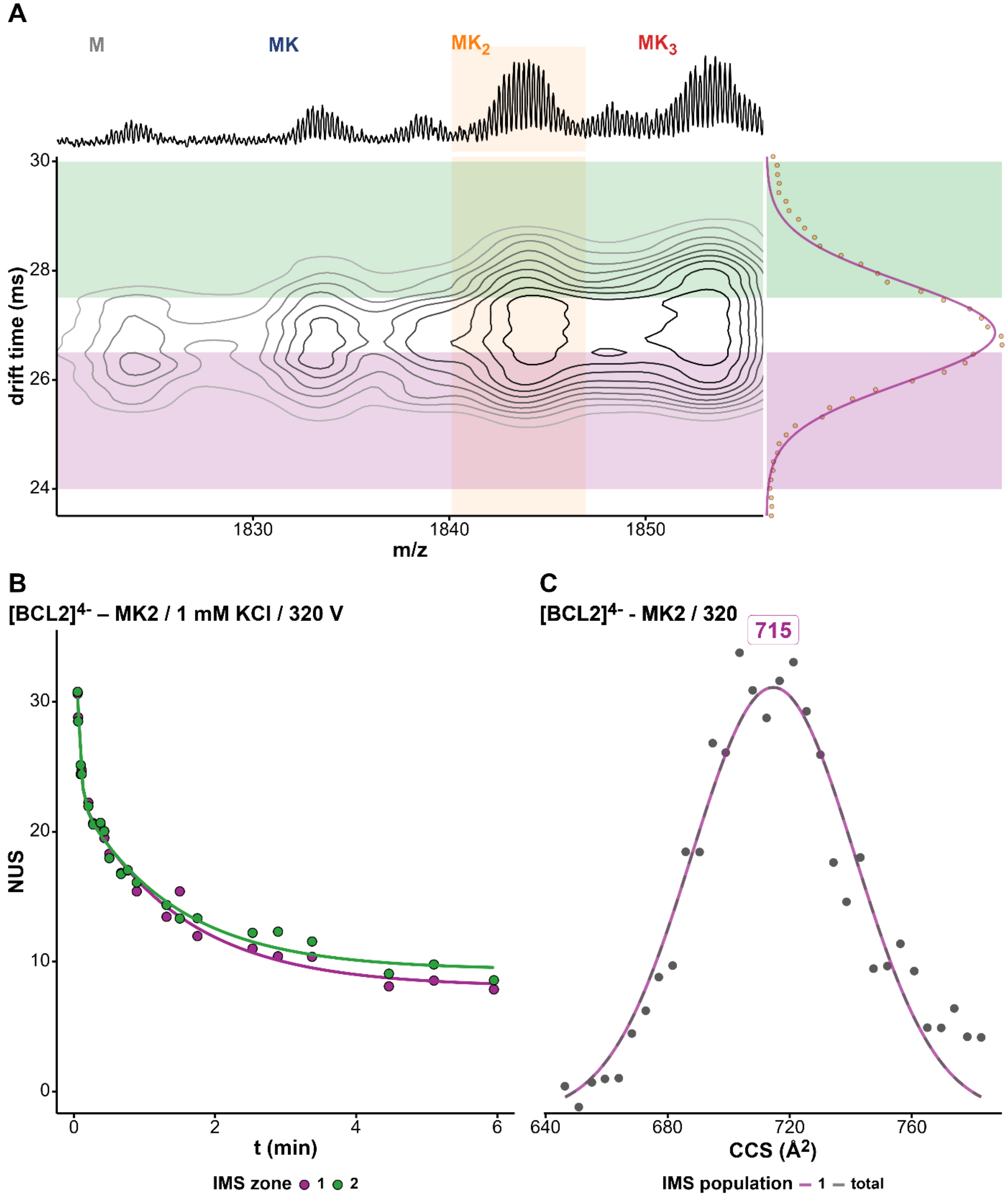


*Figure S129. (A) Density map of BCL2 in 100 mM TMAA + 1 mM KCl at fragmentor 320V and charge state 4-. The x-axis represents the mass spectrum and the y-axis the arrival-time distribution (ATD) of the $2K^+$ adduct with the Gaussian fits and the different zones selected for HDX/IMS. (B) HDX/IMS kinetics with colors matching the IMS zones of the density map for the same $K^+$ adduct. (C) CCS distribution of the $2K^+$ adduct of BCL2 with the CCS values of each population.*

*2KPR / $MK_2$ / z = 4- / 1 mM KCl / 320 V*

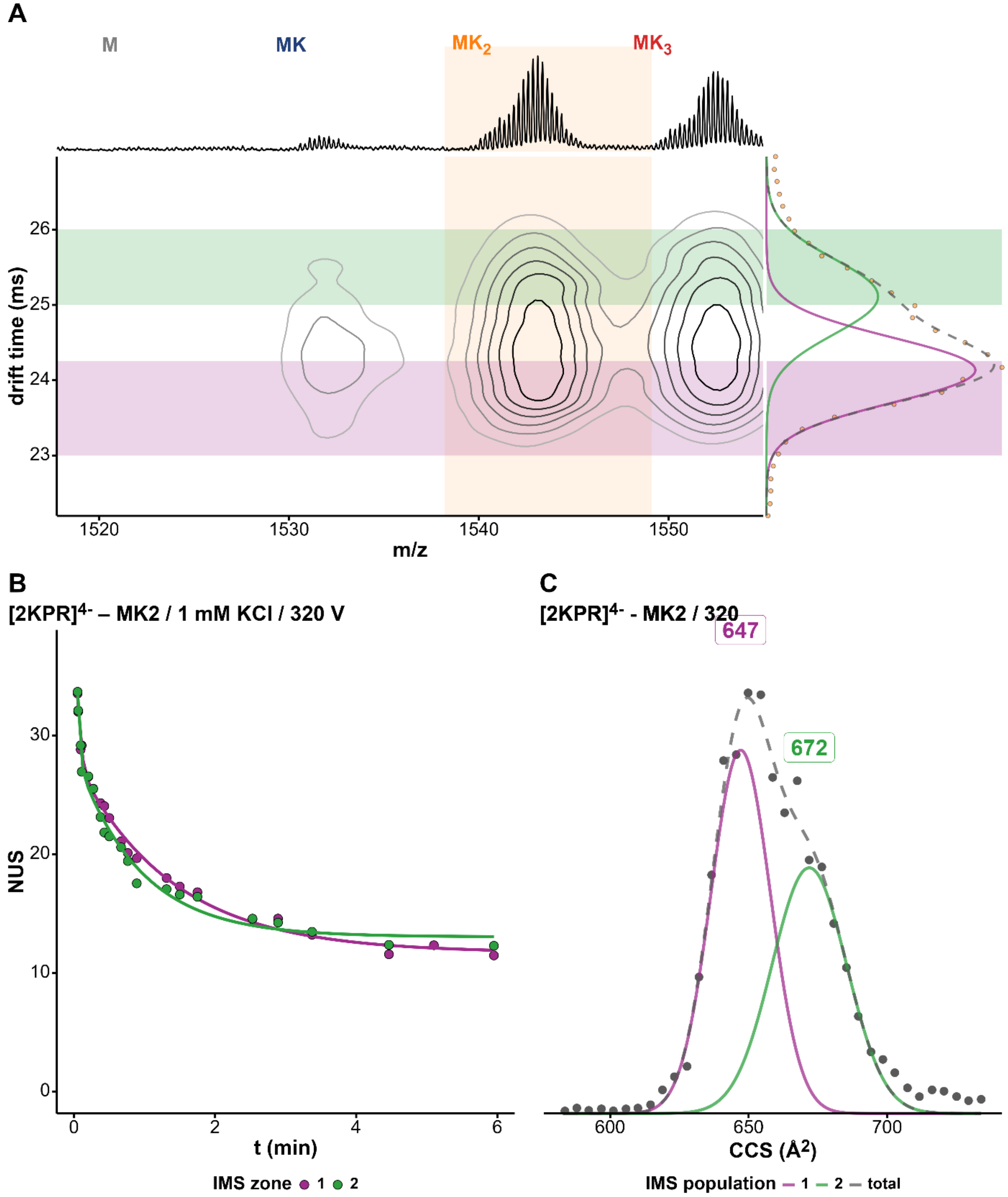


*Figure S130. (A) Density map of 2KPR in 100 mM TMAA + 1 mM KCl (z=4-). The x-axis represents the mass spectrum and the y-axis the arrival-time distribution (ATD) of the 2K⁺ adduct with the Gaussian fits and the different zones selected for HDX/IMS. (B) HDX/IMS kinetics with colors matching the IMS zones of the density map for the same 2K⁺ adduct. (C) CCS distribution of the 2K⁺ adduct of 2KPR adduct with the CCS values of each population.*

*26CEB / $MK_2$ / z = 4- / 1 mM KCl / 320 V*

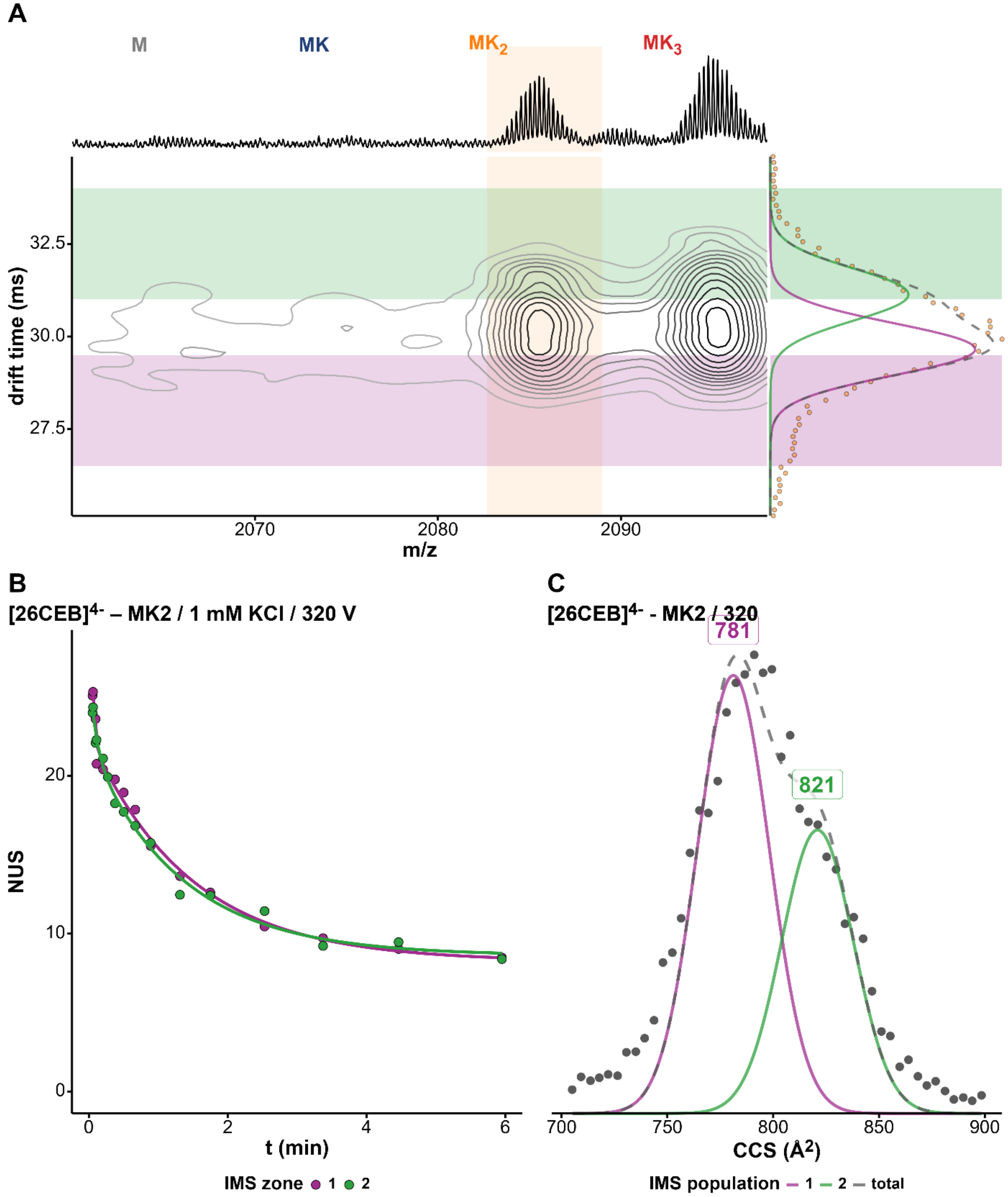


*Figure S131. (A) Density map of 26CEB in 100 mM TMAA + 1 mM KCl at fragmentor 320V and charge state 4-. The x-axis represents the mass spectrum and the y-axis the arrival-time distribution (ATD) of the 2K⁺ adduct with the Gaussian fits and the different zones selected for HDX/IMS. (B) HDX/IMS kinetics with colors matching the IMS zones of the density map for the same K⁺ adduct. (C) CCS distribution of the 2K⁺ adduct of 26CEB with the CCS values of each population.*

*26CEB / $MK_2$ / z = 5- / 1 mM KCl / 320 V*

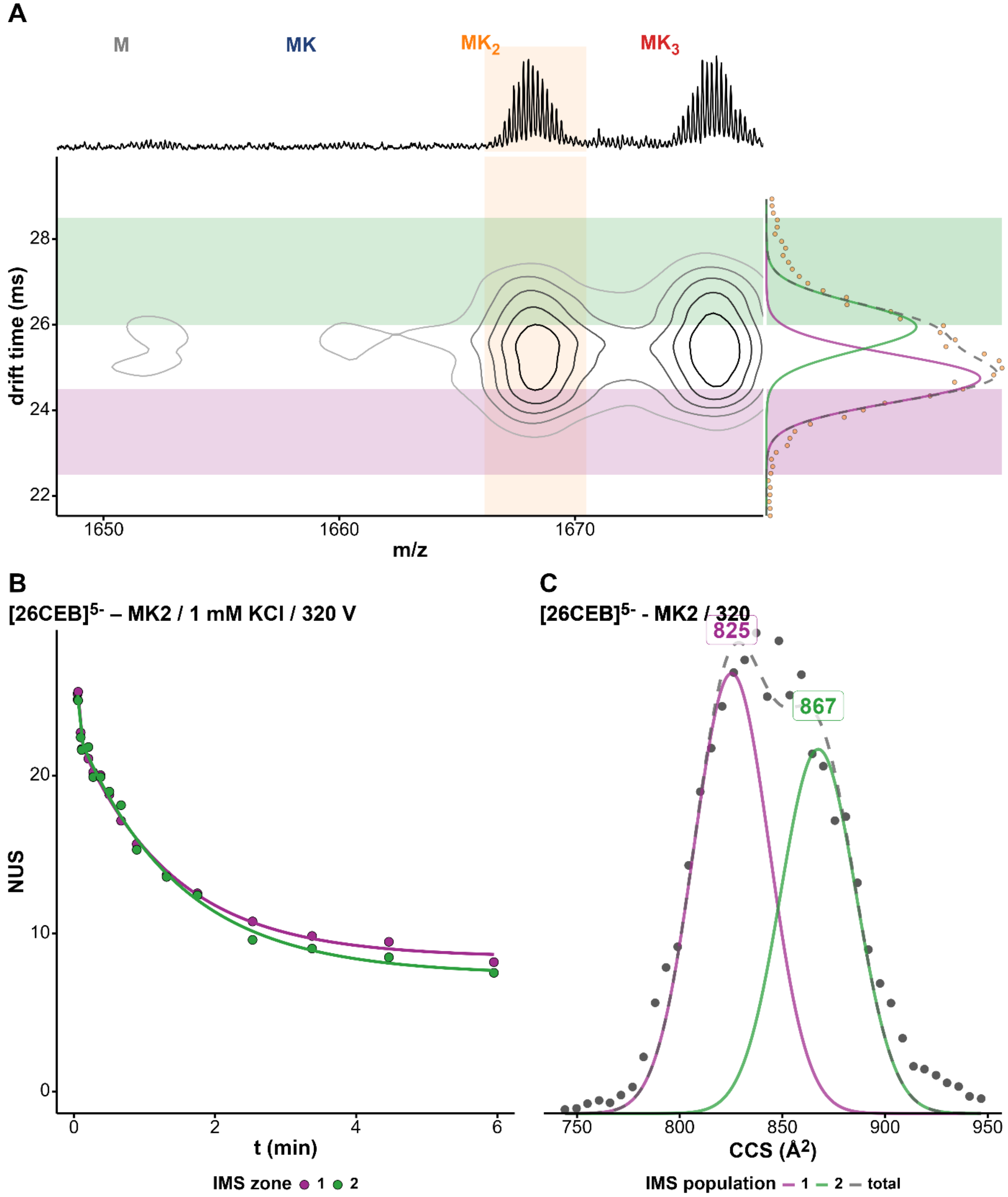


*Figure S132. (A) Density map of 26CEB in 100 mM TMAA + 1 mM KCl at fragmentor 320V and charge state 5-. The x-axis represents the mass spectrum and the y-axis the arrival-time distribution (ATD) of the $2K^+$ adduct with the Gaussian fits and the different zones selected for HDX/IMS. (B) HDX/IMS kinetics with colors matching the IMS zones of the density map for the same $K^+$ adduct. (C) CCS distribution of the $2K^+$ adduct of 26CEB with the CCS values of each population.*

*T95TT / $MK_2$ / z = 4- / 0.5 mM KCl / 320 V*

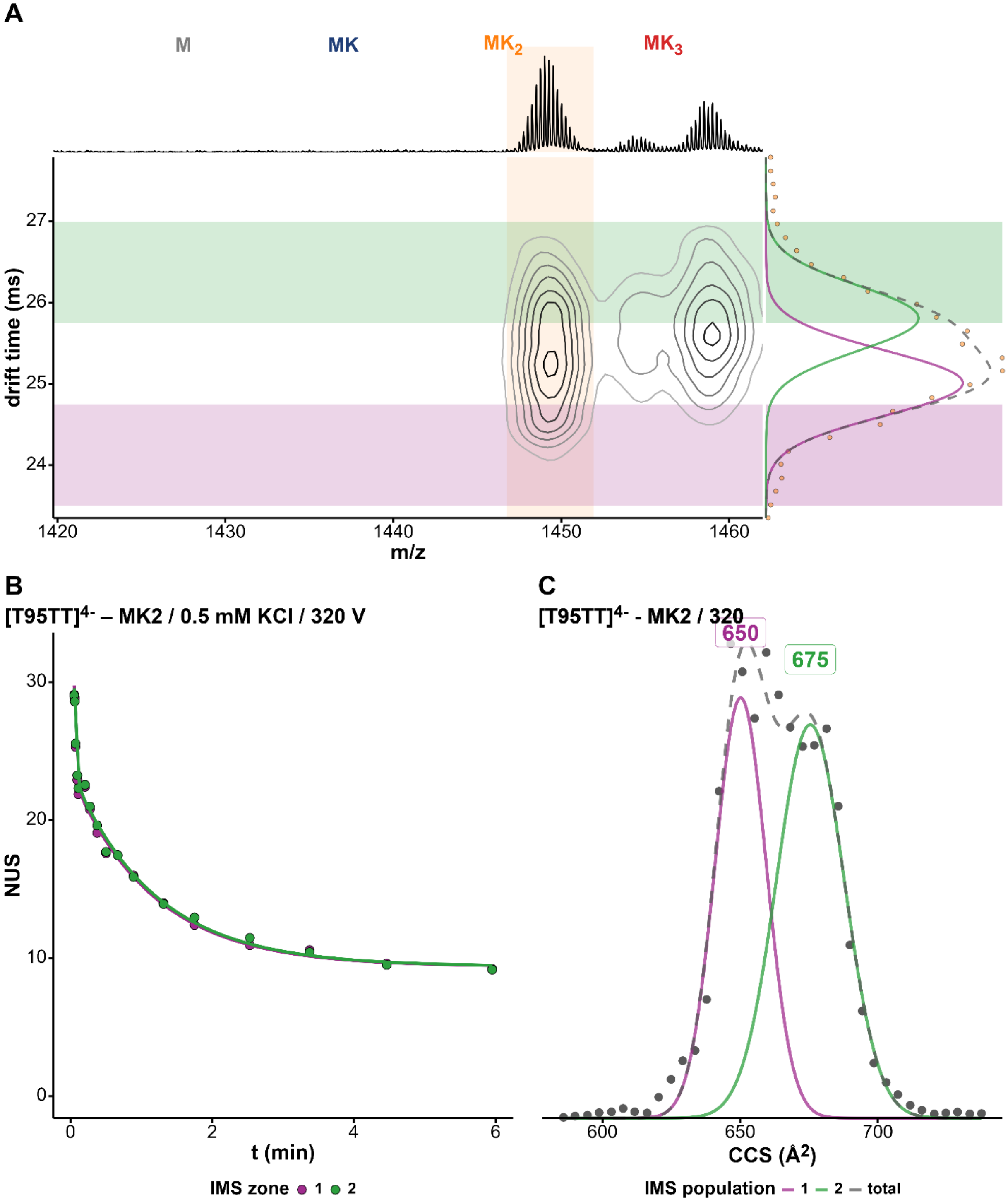


*Figure S133. (A) Density map of T95TT in 100 mM TMAA + 0.5 mM KCl at fragmentor 320V and charge state 4-. The x-axis represents the mass spectrum and the y-axis the arrival-time distribution (ATD) of the 2K⁺ adduct with the Gaussian fits and the different zones selected for HDX/IMS. (B) HDX/IMS kinetics with colors matching the IMS zones of the density map for the same K⁺ adduct. (C) CCS distribution of the 2K⁺ adduct of T95TT with the CCS values of each population.*

*T30177TT / $MK_2$ / z = 4- / 0.5 mM KCl / 320 V*

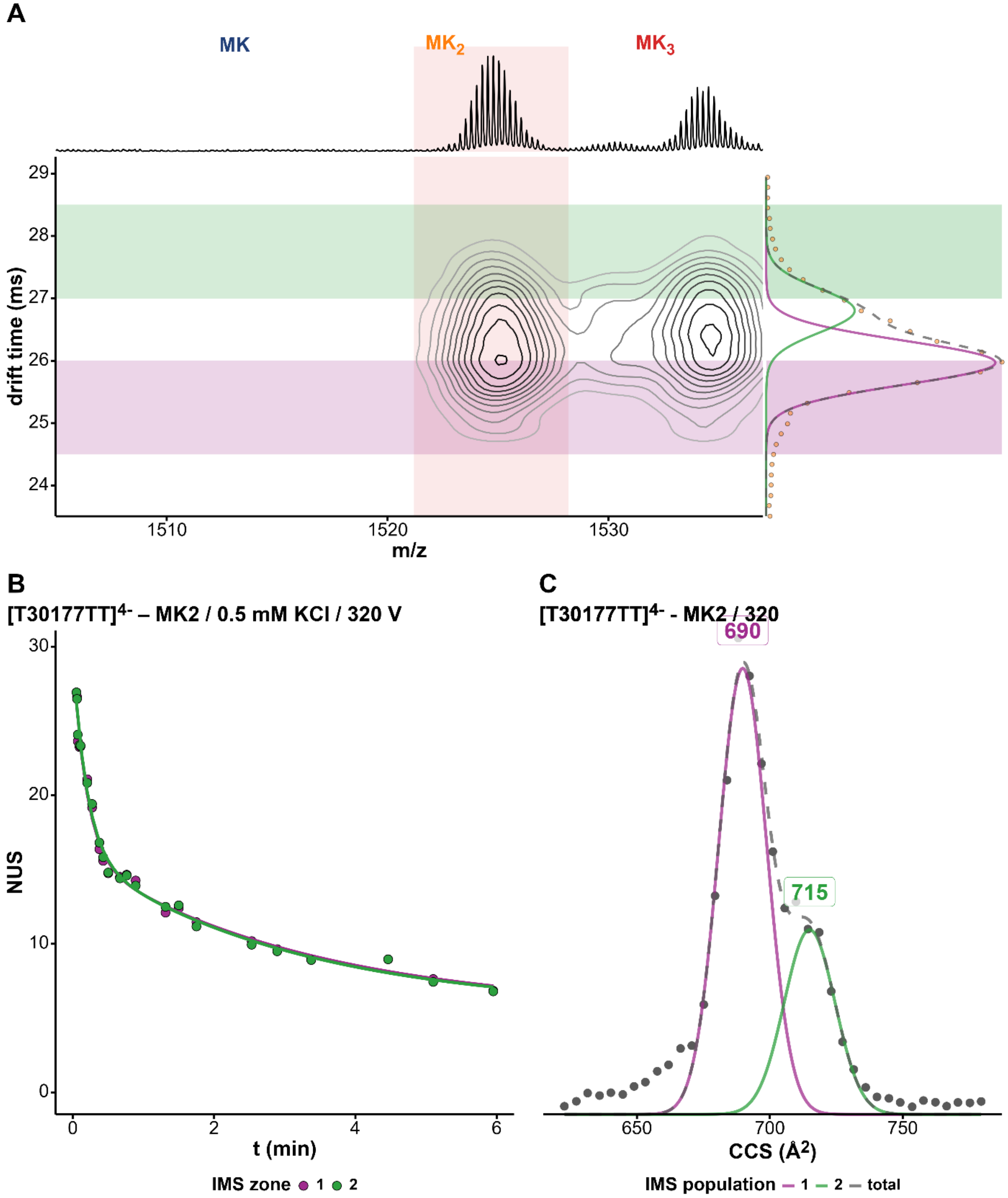


*Figure S134. (A) Density map of T30177TT in 100 mM TMAA + 0.5 mM KCl at fragmentor 320V and charge state 4-. The x-axis represents the mass spectrum and the y-axis the arrival-time distribution (ATD) of the $2K^{+}$ adduct with the Gaussian fits and the different zones selected for HDX/IMS. (B) HDX/IMS kinetics with colors matching the IMS zones of the density map for the same $K^{+}$ adduct. (C) CCS distribution of the $2K^{+}$ adduct of T30177TT with the CCS values of each population.*

*TG4T / $MK_3$ / z = 4- / 1 mM KCl / 320 V*

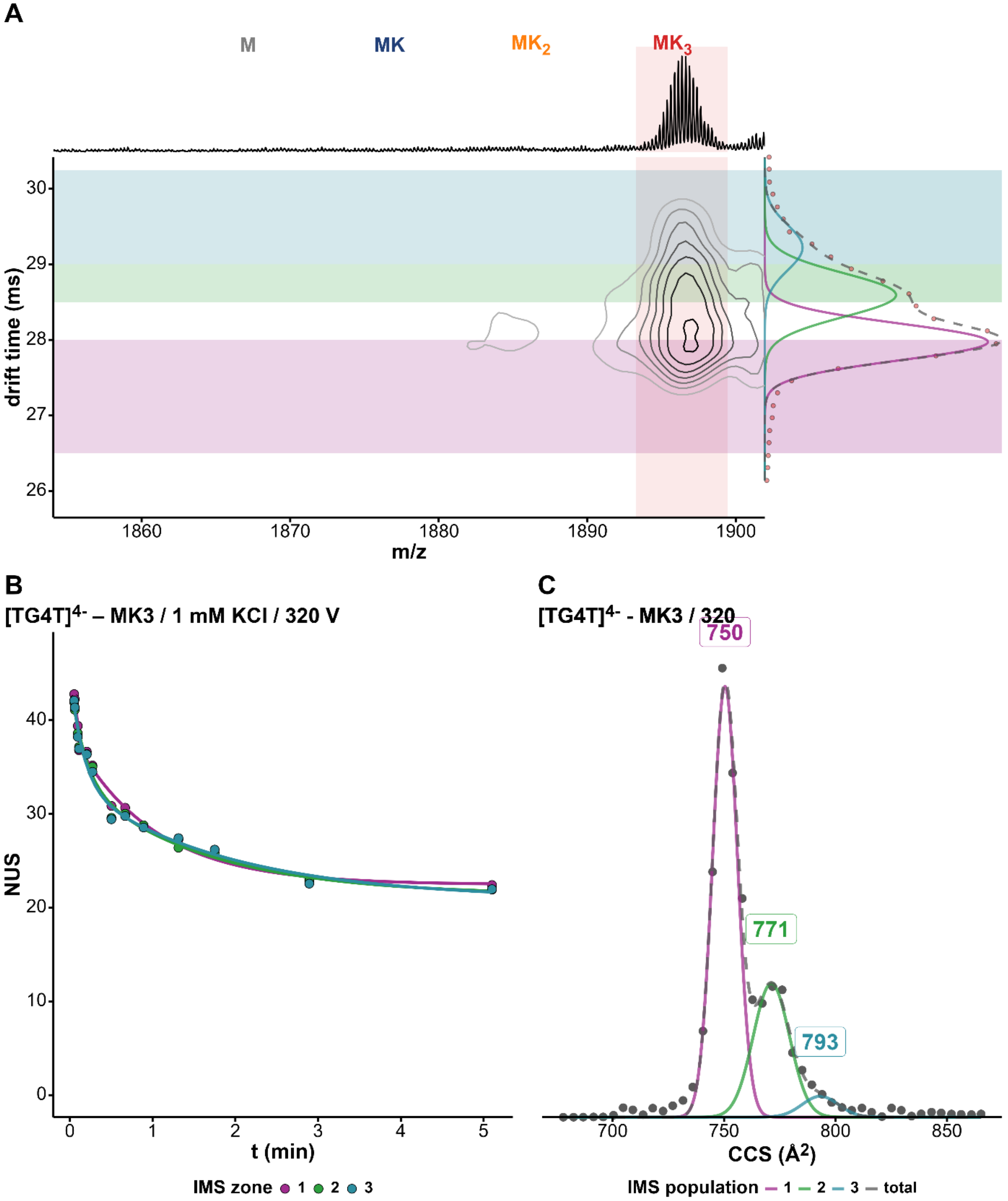


*Figure S135. (A) Density map of $[TG4T]_4$ in 100 mM TMAA + 1 mM KCl. The x-axis represents the mass spectrum and the y-axis the arrival-time distribution (ATD) of the $3K^+$ adduct with the Gaussian fits and the different zones selected for HDX/IMS. (B) HDX/IMS kinetics with colors matching the IMS zones of the density map for the same $K^+$ adduct. (C) CCS distribution of the $3K^+$ adduct of TG4T adduct with the CCS values of each population.*

*TG4T / M($NH_4^+$)$_3$ / z = 4- / 1 mM KCl / 320 V*

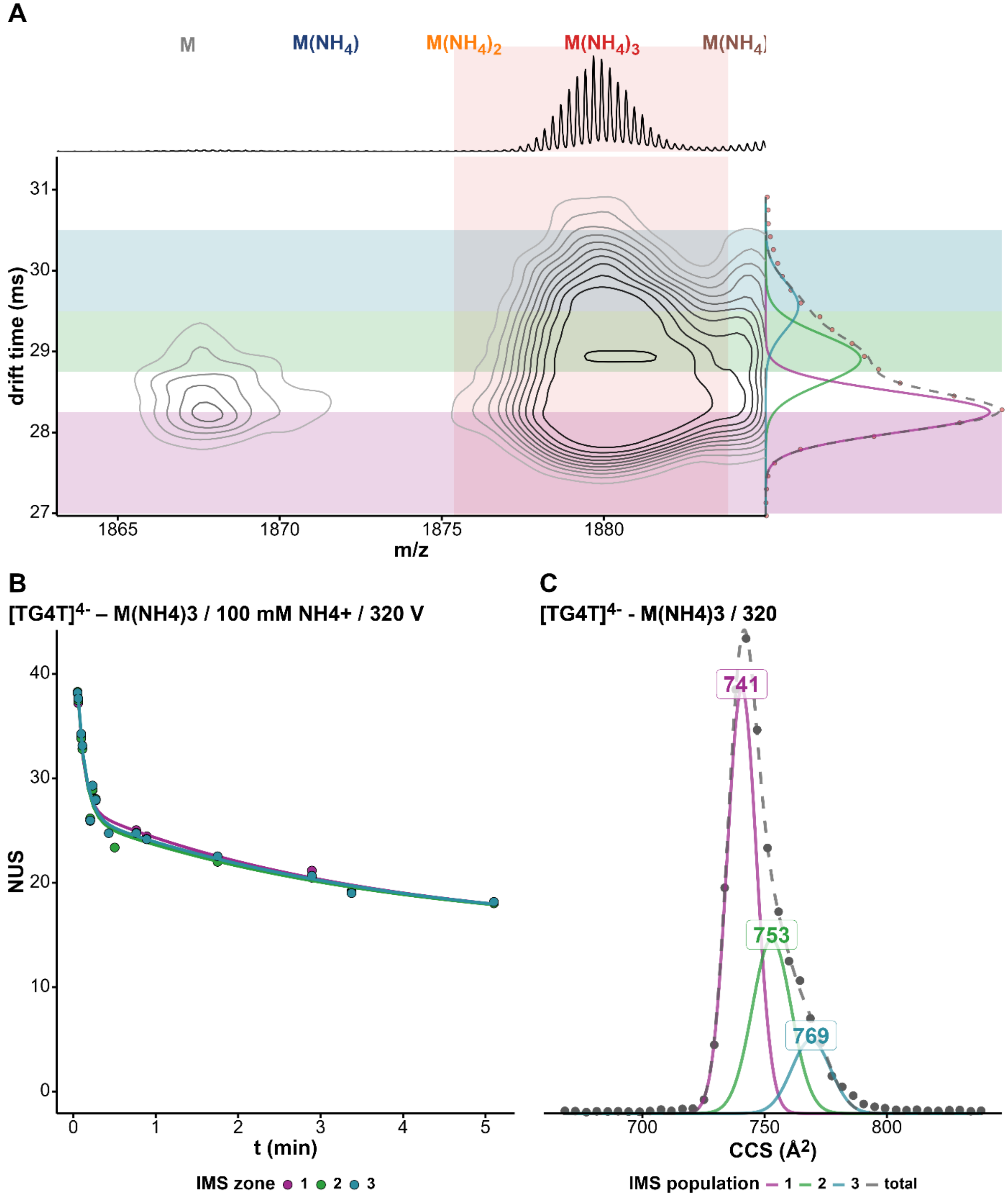


*Figure S136. (A) Density map of [TG4T]$_4$ in 100 mM $NH_4OAc$ (z=4-). The x-axis represents the mass spectrum and the y-axis the arrival-time distribution (ATD) of the 3$NH_4^+$ adduct with the Gaussian fits and the different zones selected for HDX/IMS. (B) HDX/IMS kinetics with colors matching the IMS zones of the density map for the same $NH_4^+$ adduct. (C) CCS distribution of the 3$NH_4^+$ adduct of TG4T adduct with the CCS values of each population.*

*TG4T / M($NH_4^+$)$_3$ / z = 5- / 1 mM KCl / 320 V*

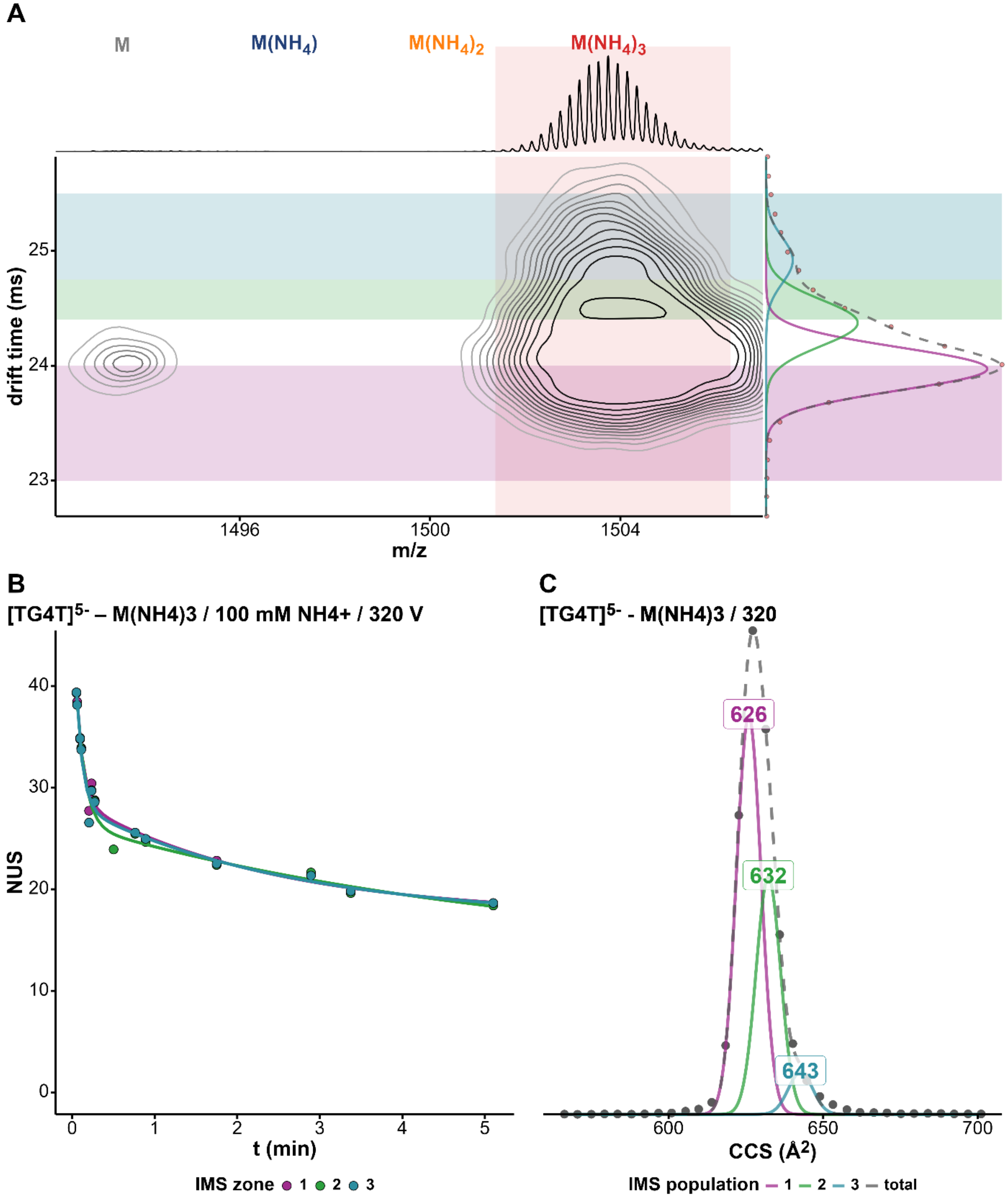


*Figure S137. (A) Density map of [TG4T]$_4$ in 100 mM $NH_4OAc$ (z=5-). The x-axis represents the mass spectrum and the y-axis the arrival-time distribution (ATD) of the 3$NH_4^+$ adduct with the Gaussian fits and the different zones selected for HDX/IMS. (B) HDX/IMS kinetics with colors matching the IMS zones of the density map for the same $NH_4^+$ adduct. (C) CCS distribution of the 3$NH_4^+$adduct of TG4T adduct with the CCS values of each population.*

### Isotopic distribution deconvolutions

*23TAG / $MK_2$ / 1 mM KCl / 370 V / zone 1*

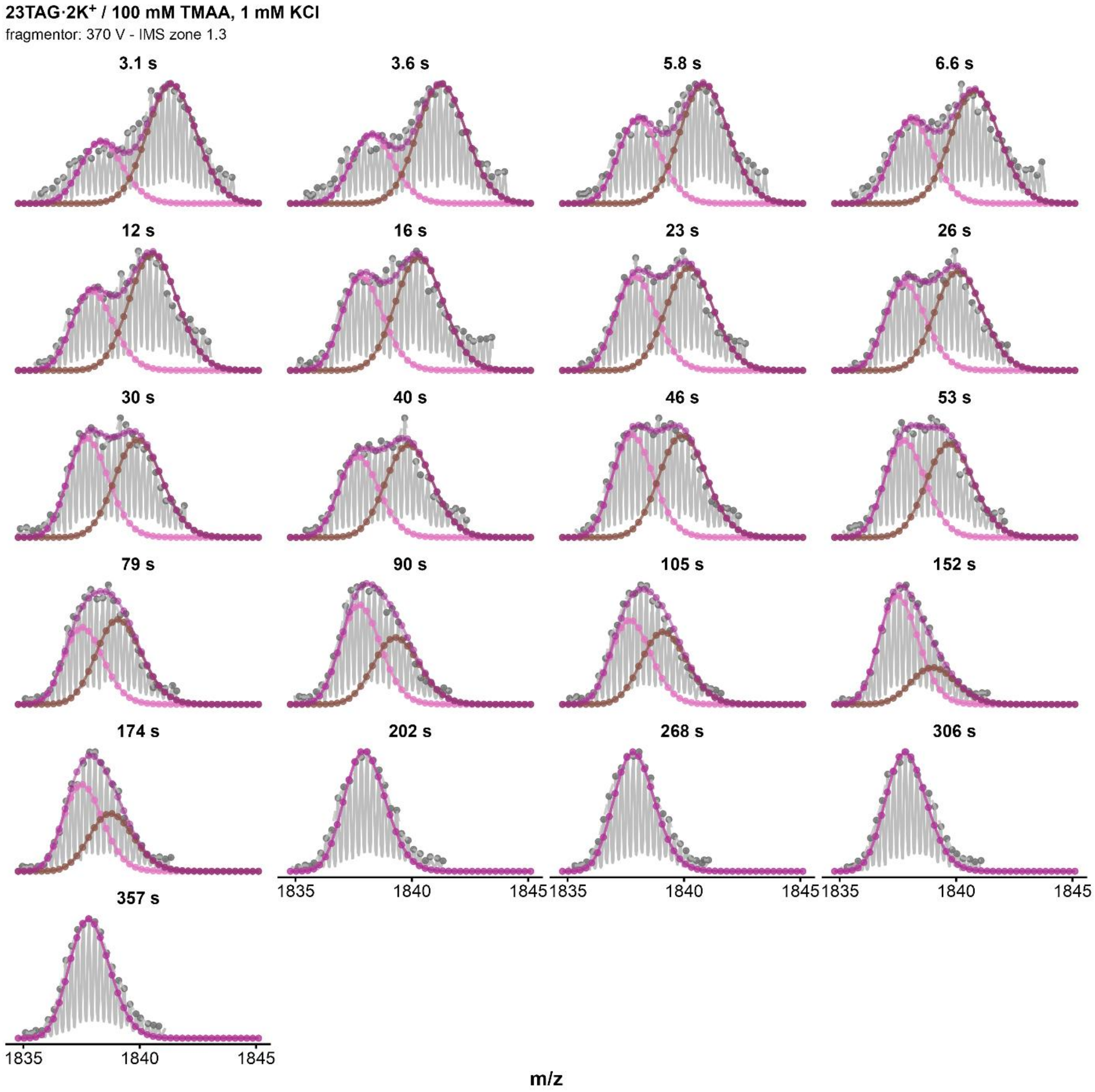


*Figure S138. Isotopic MS deconvolutions of 23TAG·2K⁺ zone 1 (fragmentor = 370V, z=4-) for every HDX timepoints. Purple curve correspond to the fit of the whole distribution, the pink curve to the highly exchanged population, and the brown one to the low-exchanged population. The distribution is considered monomodal after 202s.*

### *23TAG / $MK_2$ / 1 mM KCl / 370 V / zone 2*

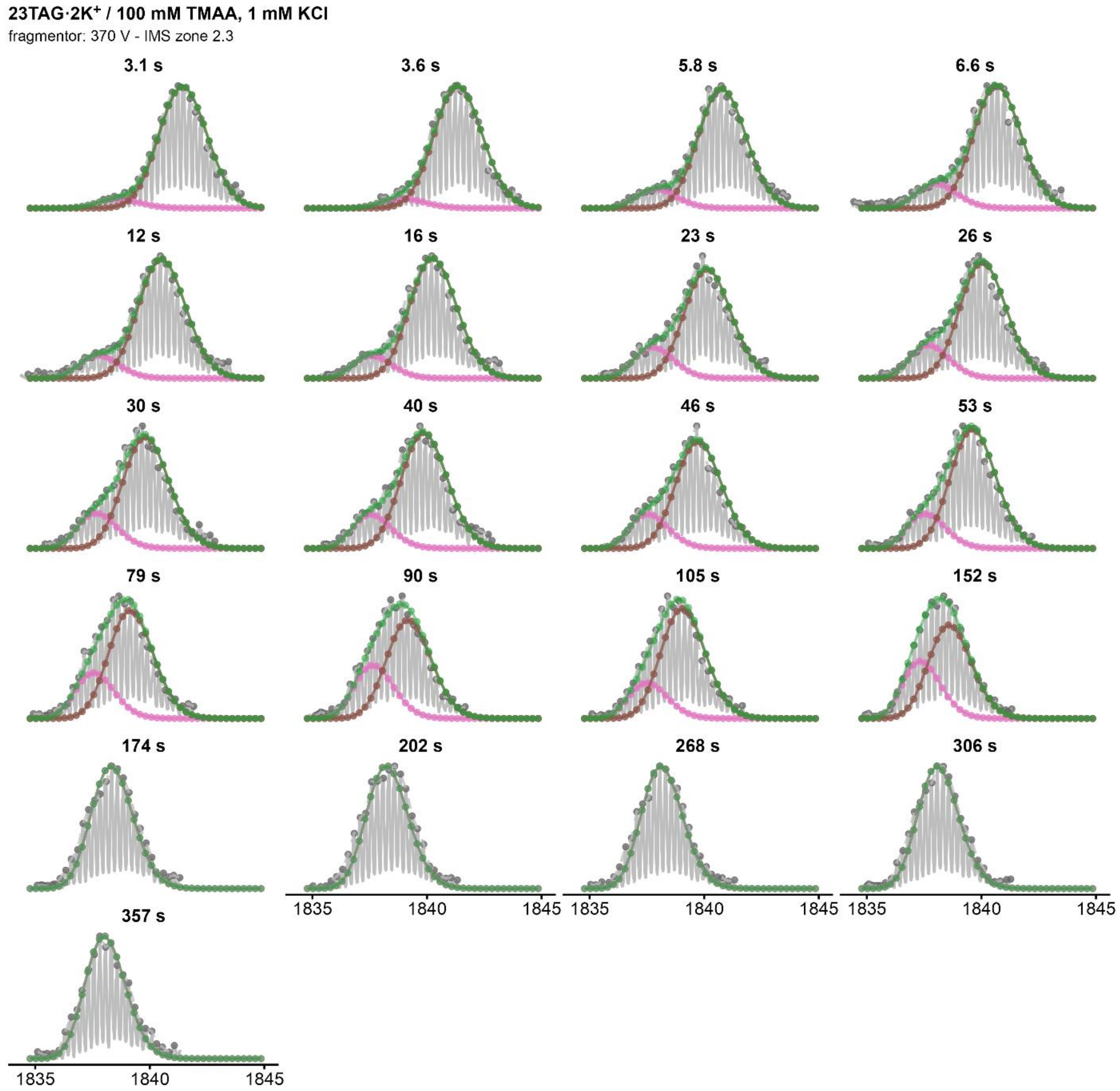


*Figure S139. Isotopic MS deconvolutions of 23TAG·$2K^+$ zone 2 (fragmentor = 370V, z=4-) for every HDX timepoints. Green curve correspond to the fit of the whole distribution, the pink curve to the highly exchanged population, and the brown one to the low-exchanged population. The distribution is considered monomodal after 174s.*

### *23TAG / $MK_2$ / 1 mM KCl / 320 V / zone 1*

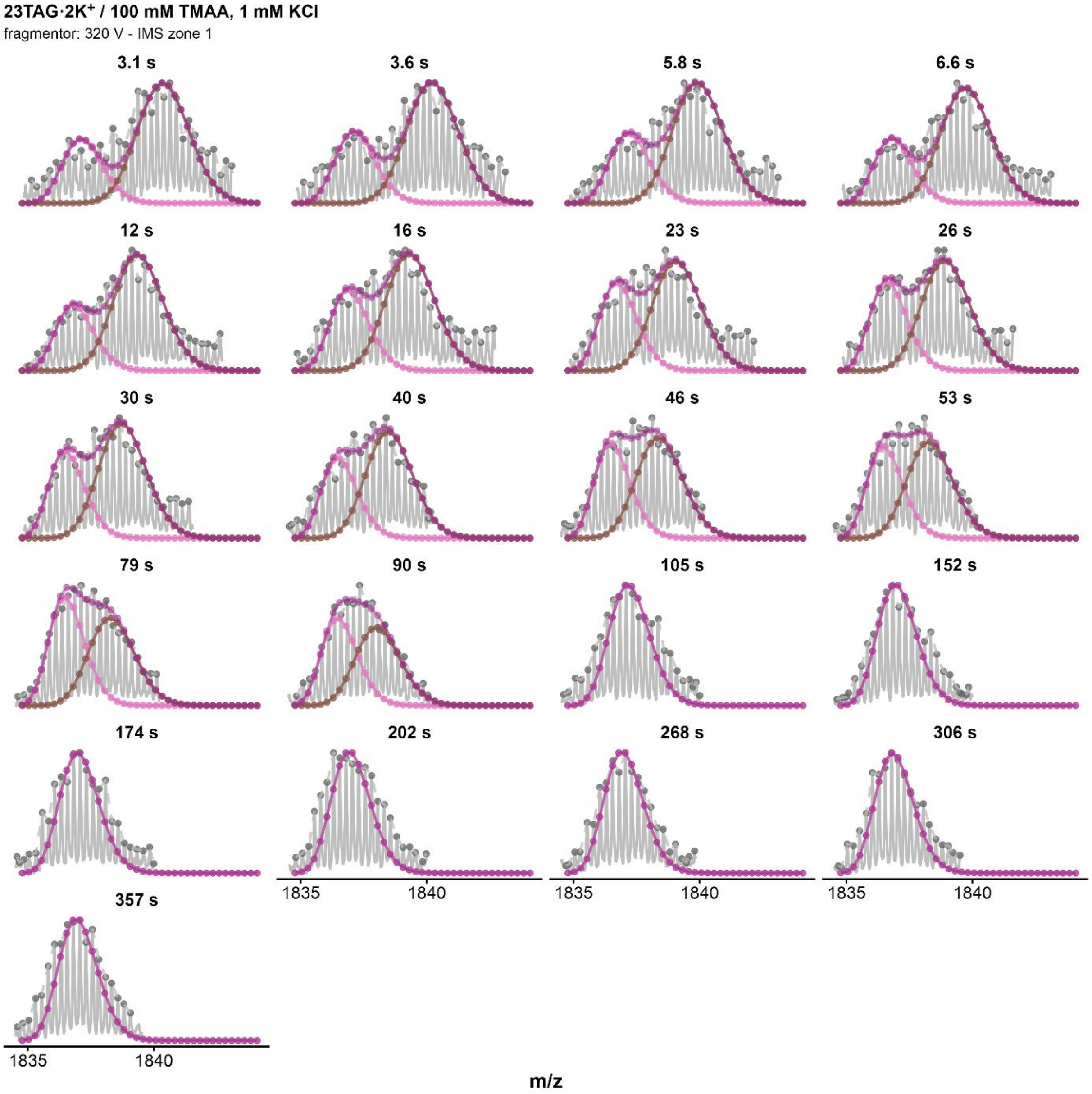


*Figure S140. Isotopic MS deconvolutions of 23TAG·$2K^+$ zone 1 (fragmentor = 320V, z=4-) for every HDX timepoints. Purple curve correspond to the fit of the whole distribution, the pink curve to the highly exchanged population, and the brown one to the low-exchanged population. The distribution is considered monomodal after 105s.*

### *23TAG / $MK_2$ / 1 mM KCl / 320 V / zone 2*

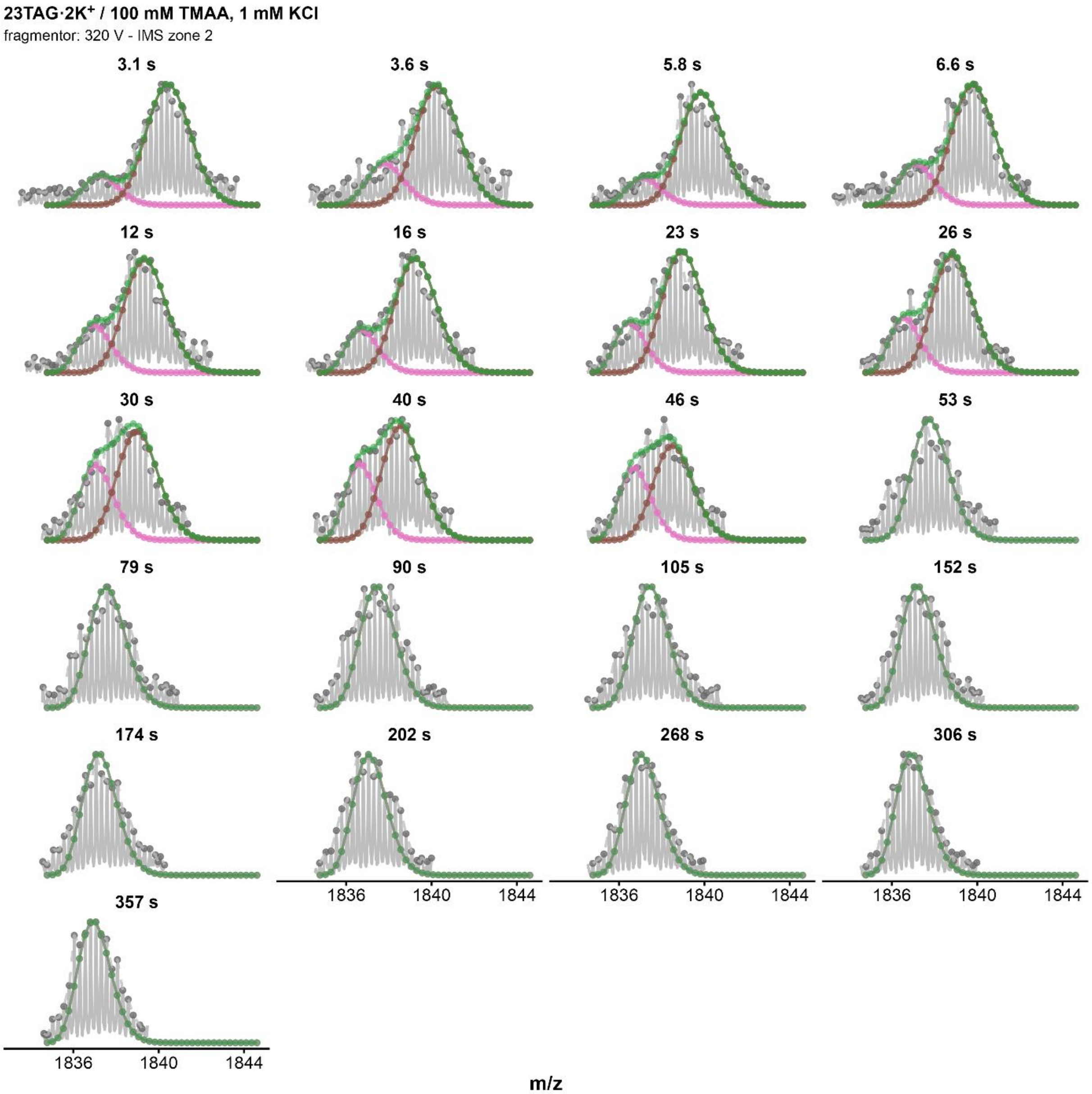


*Figure S141. Isotopic MS deconvolutions of 23TAG·$2K^+$ zone 2 (fragmentor = 320V, z=4-) for every HDX timepoints. Green curve correspond to the fit of the whole distribution, the pink curve to the highly exchanged population, and the brown one to the low-exchanged population. The distribution is considered monomodal after 174s.*

### *23TAG / $MK_2$ / 1 mM KCl / 350 V / zone 1*

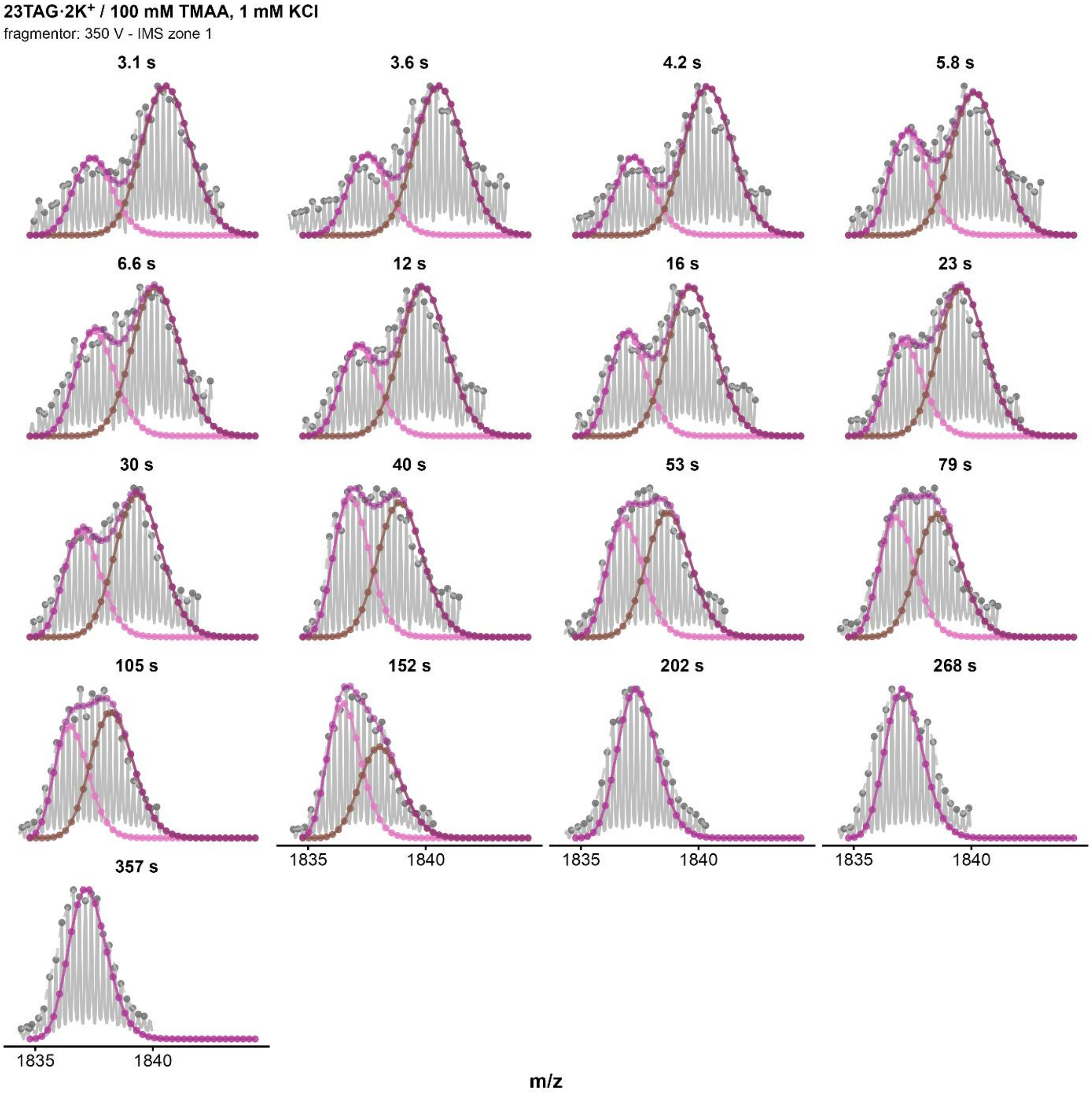


*Figure S142. Isotopic MS deconvolutions of 23TAG·$2K^+$ zone 1 (fragmentor = 350V, z=4-) for every HDX timepoints. Purple curve correspond to the fit of the whole distribution, the pink curve to the highly exchanged population, and the brown one to the low-exchanged population. The distribution is considered monomodal after 152s.*

### *23TAG / $MK_2$ / 1 mM KCl / 350 V / zone 2*

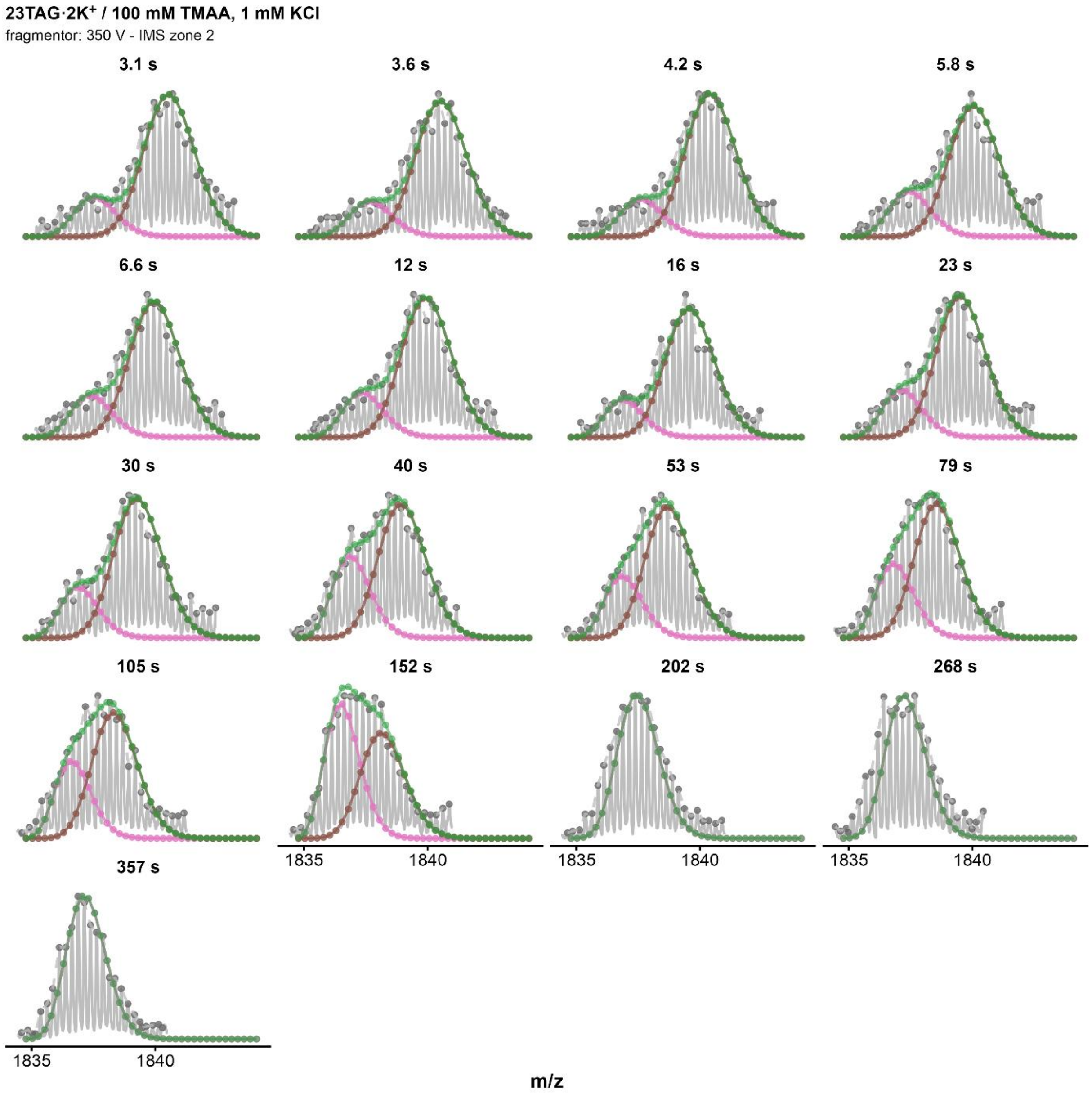


*Figure S143. Isotopic MS deconvolutions of 23TAG·2K⁺ zone 2 (fragmentor = 350V, z=4-) for every HDX timepoints. Green curve correspond to the fit of the whole distribution, the pink curve to the highly exchanged population, and the brown one to the low-exchanged population. The distribution is considered monomodal after 152s.*

### *23TAG / MK / 1 mM KCl / 370 V / zone 1*

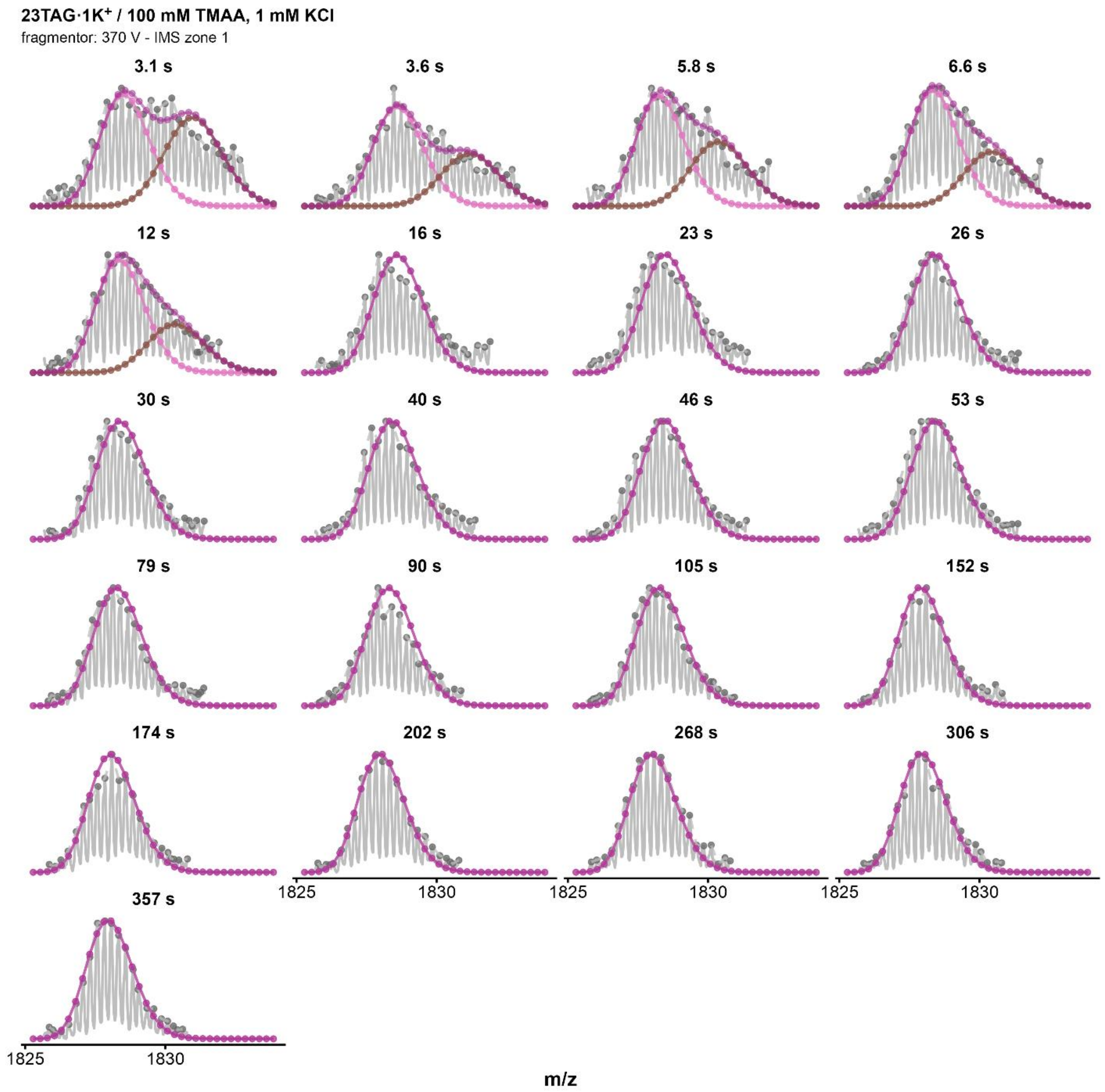


*Figure S144. Isotopic MS deconvolutions of 23TAG·1K+ zone 1 (fragmentor = 370V, z=4-) for every HDX timepoints. Purple curve correspond to the fit of the whole distribution, the pink curve to the highly exchanged population, and the brown one to the low-exchanged population. The distribution is considered monomodal after 12s.*

### *23TAG / MK / 1 mM KCl / 370 V / zone 2*

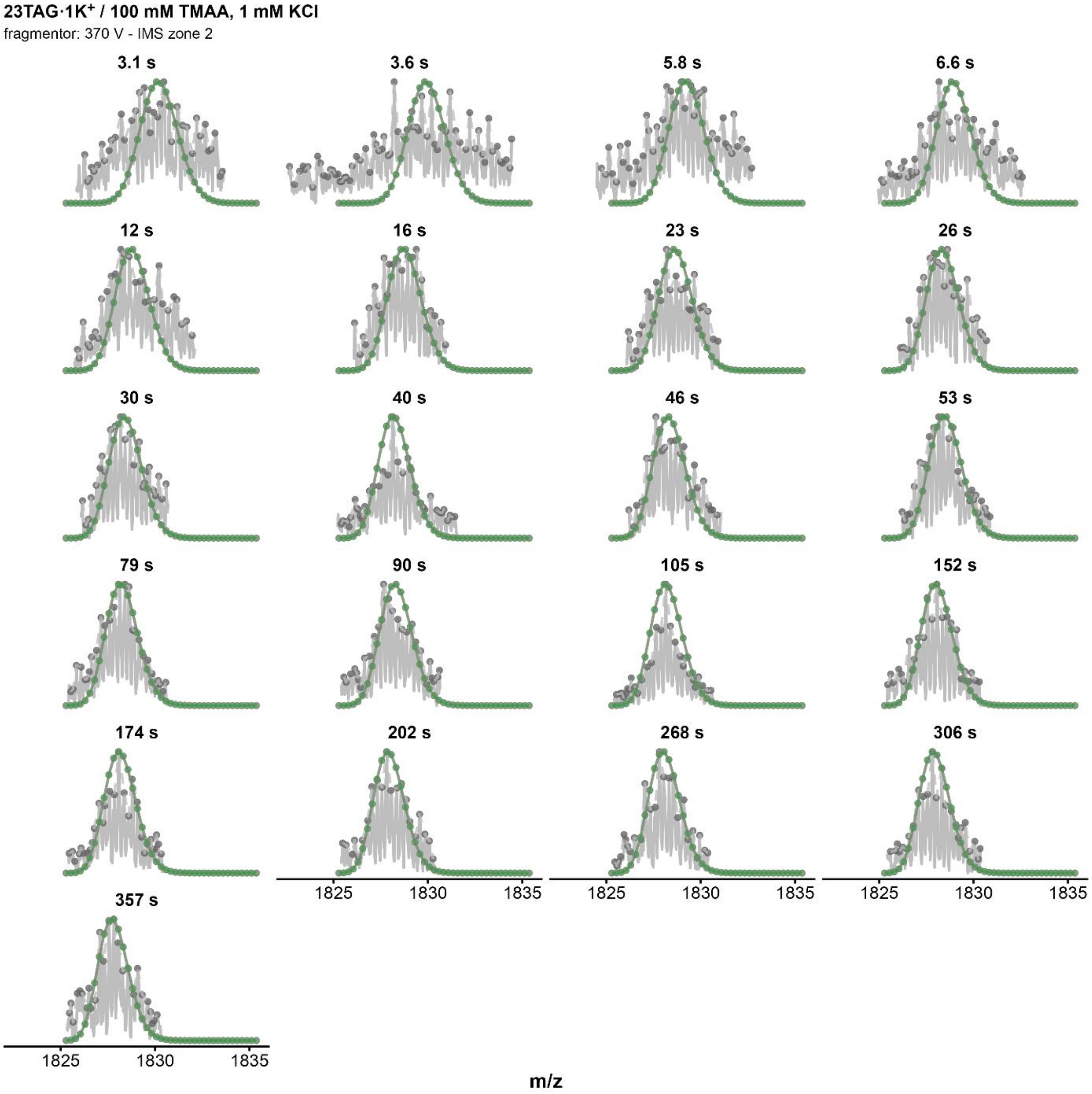


*Figure S145. Isotopic MS deconvolutions of 23TAG·2K⁺ zone 2 (fragmentor = 370V, z=4-) for every HDX timepoints. The data here was extremely noisy and could not allow to process to a decent deconvolution.*

### *5YEY / $MK_2$ / 1 mM KCl / 320 V / zone 1*

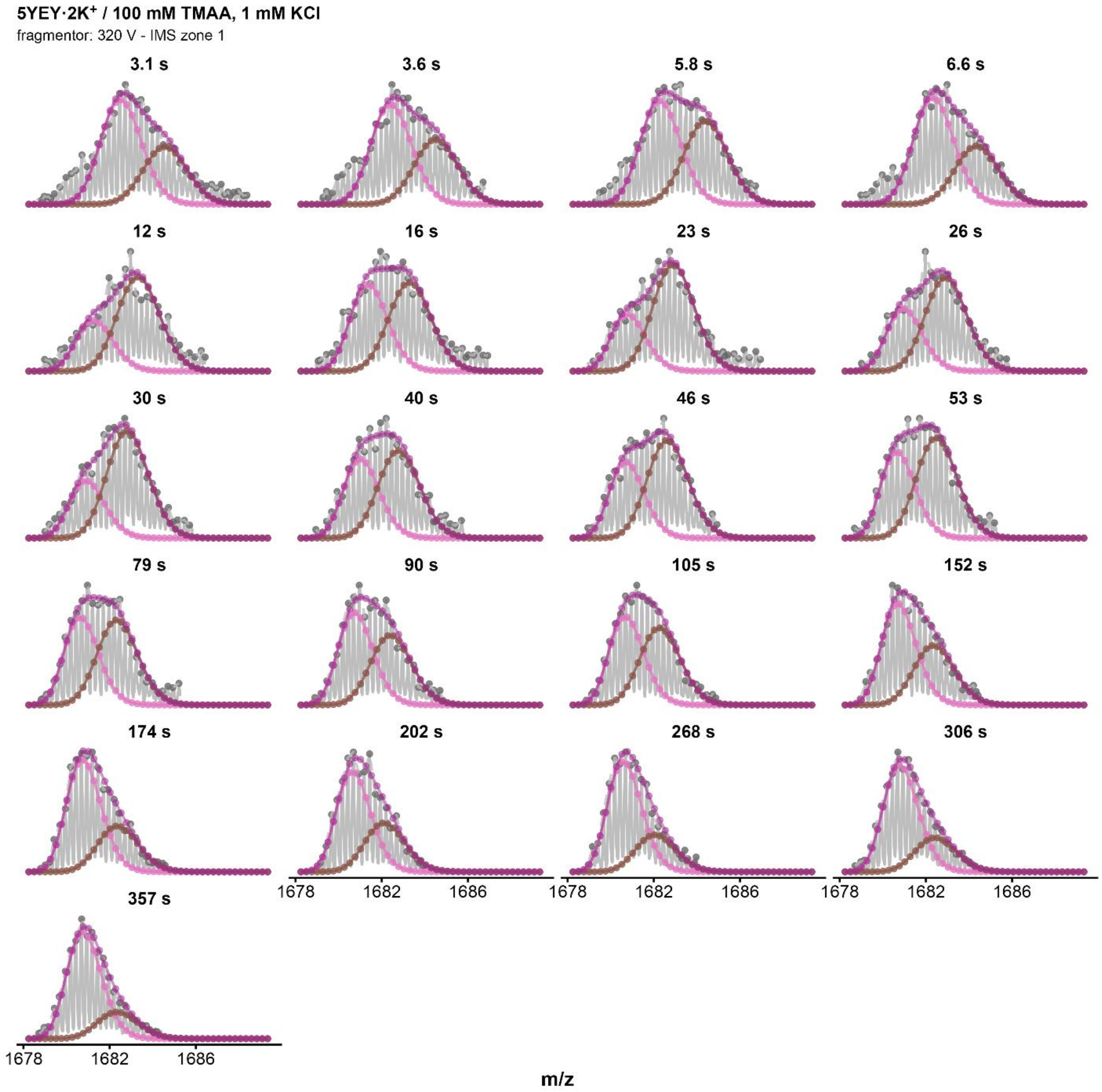


*Figure S146. Isotopic MS deconvolutions of 5YEY·2K⁺ zone 1 (fragmentor = 320V, z=4-) for every HDX timepoints. Purple curve correspond to the fit of the whole distribution, the pink curve to the highly exchanged population, and the brown one to the low-exchanged population.*

### *5YEY / $MK_2$ / 1 mM KCl / 320 V / zone 2*

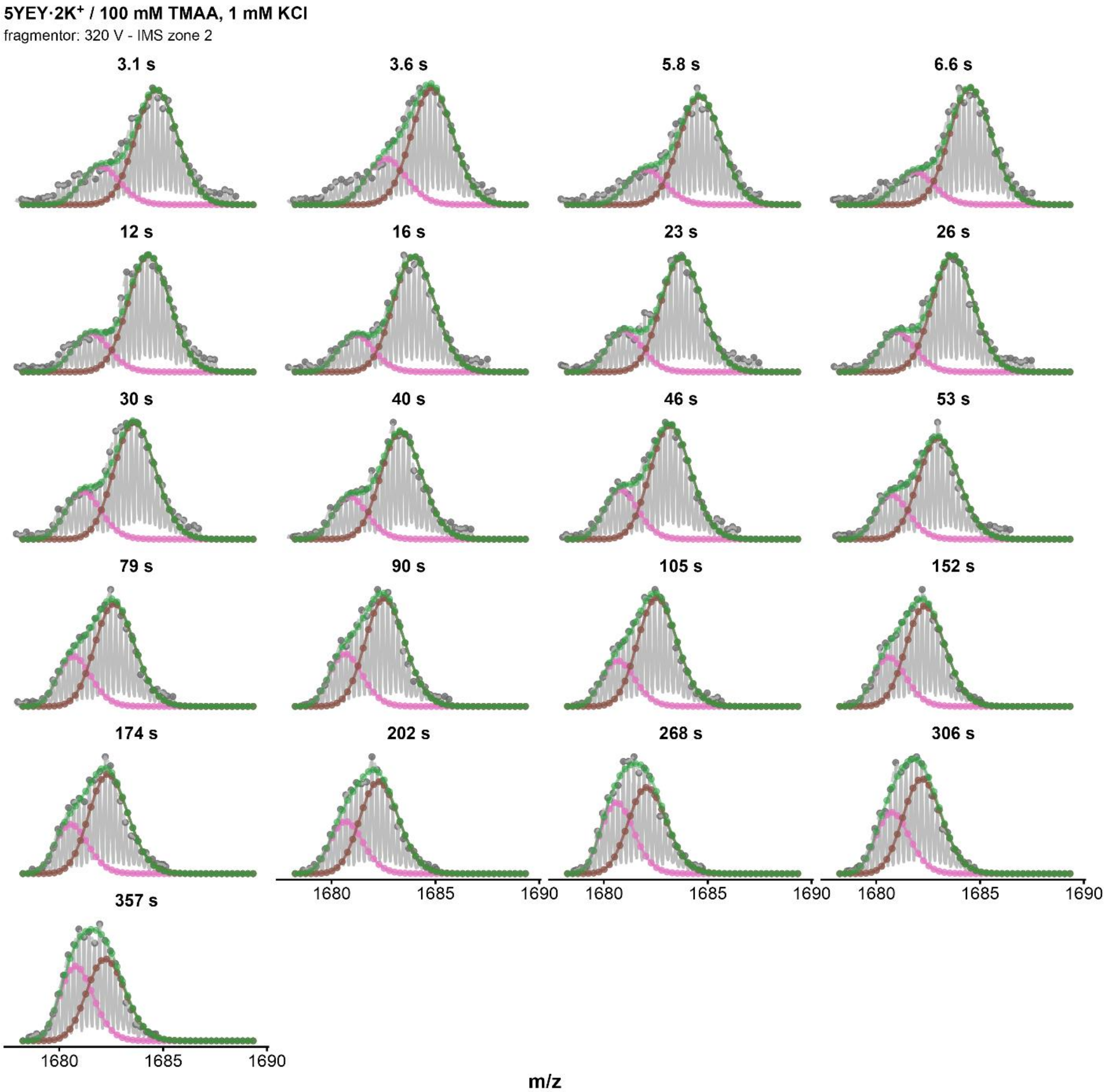


*Figure S147. Isotopic MS deconvolutions of 5YEY·2K$^+$ zone 2 (fragmentor = 320V, z=4-) for every HDX timepoints. Green curve correspond to the fit of the whole distribution, the pink curve to the highly exchanged population, and the brown one to the low-exchanged population.*

### *5YEY / MK₂ / 1 mM KCl / 320 V / zone 3*

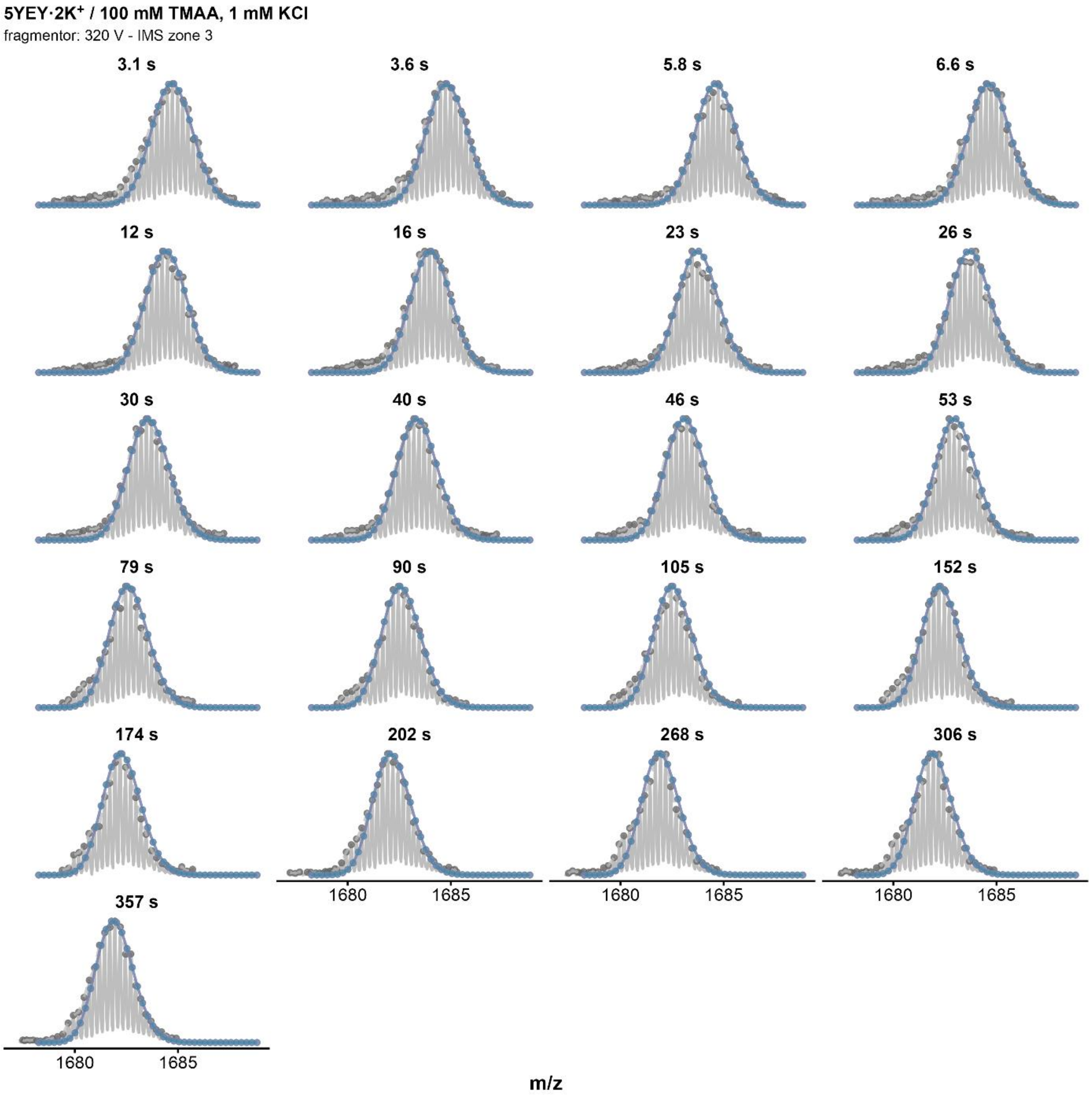


*Figure S148. Isotopic MS deconvolutions of 5YEY·2K+ zone 3 (fragmentor = 320V, z=4-) for every HDX timepoints. blue curve correspond to the fit of the whole distribution.*

### *5YEY / MK / 1 mM KCl / 320 V / zone 1*

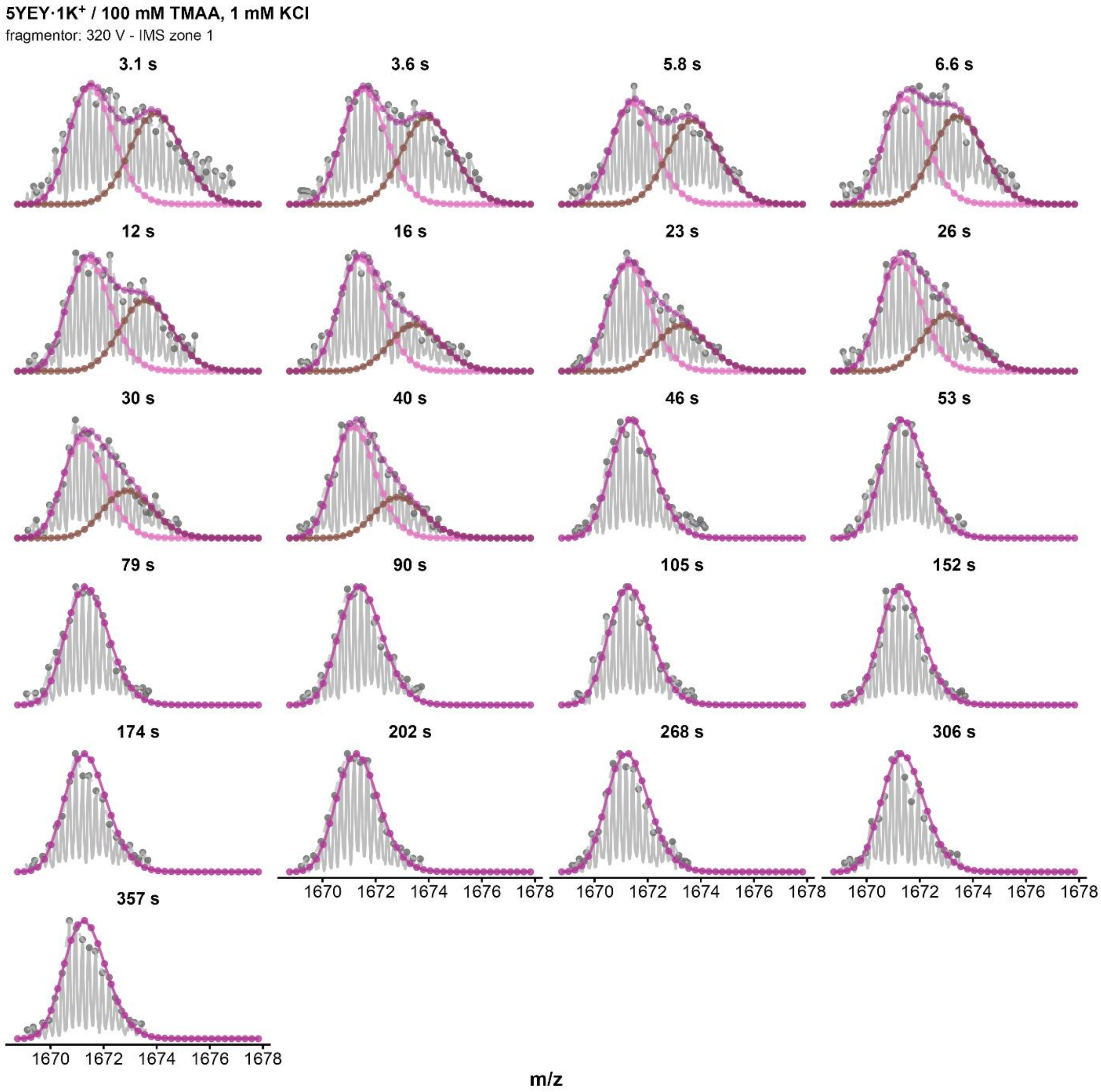


*Figure S149. Isotopic MS deconvolutions of 5YEY·1K⁺ zone 1 (fragmentor = 320V, z=4-) for every HDX timepoints. Purple curve correspond to the fit of the whole distribution, the pink curve to the highly exchanged population, and the brown one to the low-exchanged population. The distribution is considered monomodal after 40s.*

*5YEY / MK / 1 mM KCl / 320 V / zone 2*

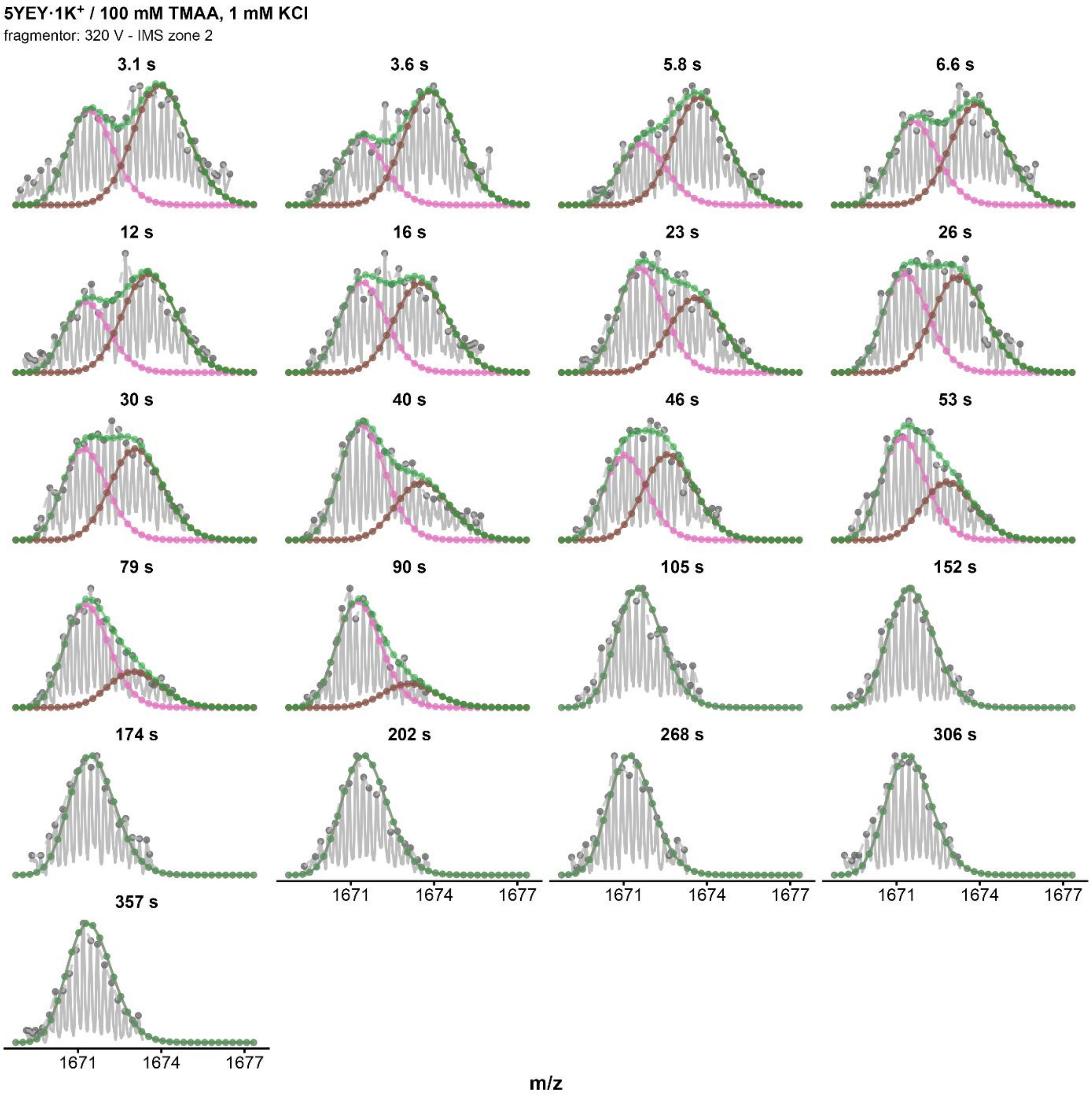


*Figure S150. Isotopic MS deconvolutions of 5YEY·1K⁺ zone 2 (fragmentor = 320V, z=4-) for every HDX timepoints. Green curve correspond to the fit of the whole distribution, the pink curve to the highly exchanged population, and the brown one to the low-exchanged population. The distribution is considered monomodal after 90s.*

### *22GT_18T / $MK_2$ / 1 mM KCl / 320 V / zone 1*

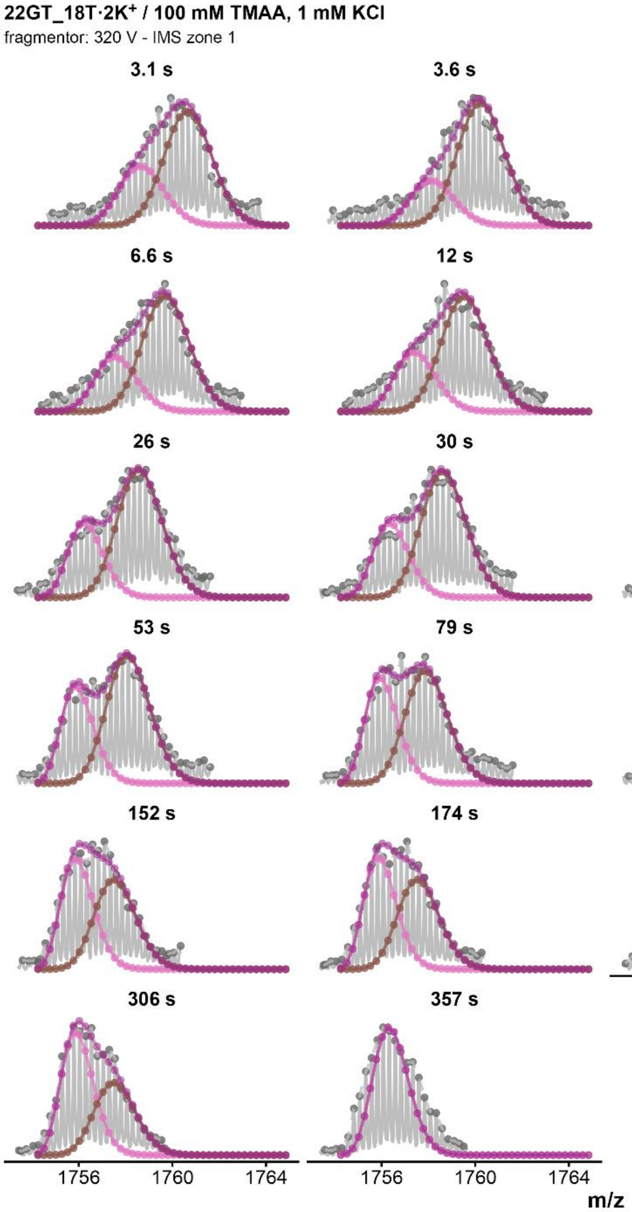


*Figure S151. Isotopic MS deconvolutions of 22GT_18T·$2K^+$ zone 1 (fragmentor = 320V, z=4-) for every HDX timepoints. Purple curve correspond to the fit of the whole distribution, the pink curve to the highly exchanged population, and the brown one to the low-exchanged population. The distribution is considered monomodal after 306s.*

### *22GT_18T / $MK_2$ / 1 mM KCl / 320 V / zone 2*

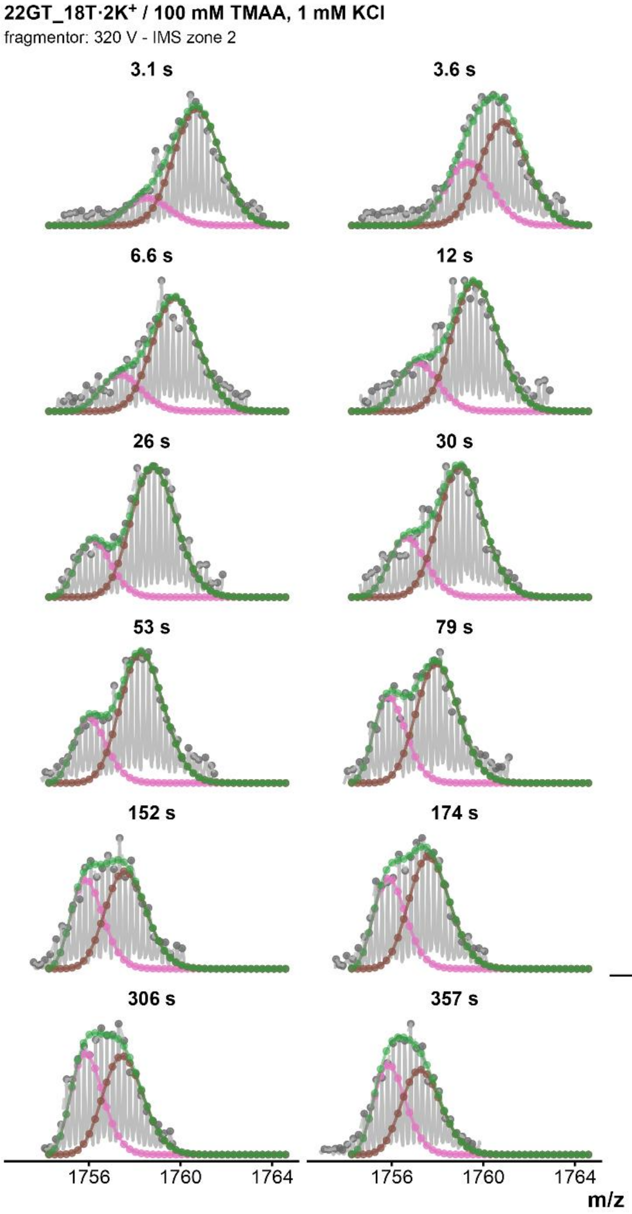


*Figure S152. Isotopic MS deconvolutions of 22GT_18T·2K⁺ zone 2 (fragmentor = 320V, z=4-) for every HDX timepoints. Green curve correspond to the fit of the whole distribution, the pink curve to the highly exchanged population, and the brown one to the low-exchanged population.*

### *24TTG / $MK_2$ / 1 mM KCl / 320 V / zone 1*

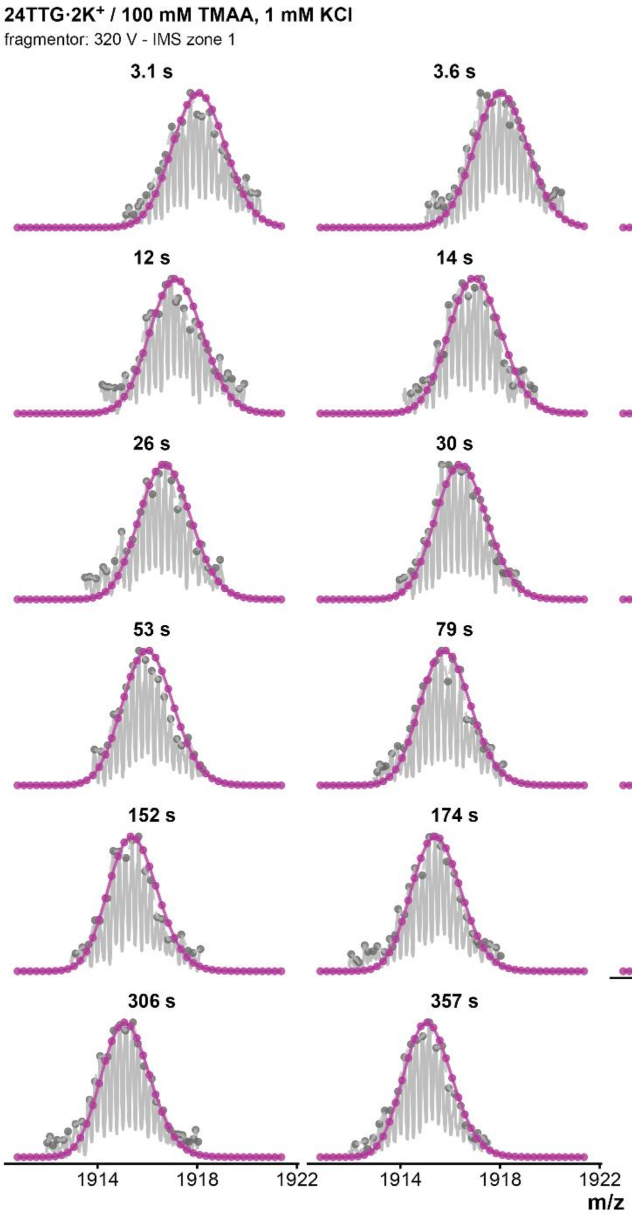


*Figure S153. Isotopic MS deconvolutions of 24TTG · 2K+ zone 2 (fragmentor = 320V, z=4-) in 1 mM KCl for every HDX timepoints. Purple curve correspond to the fit of the whole distribution.*

### *24TTG / $MK_2$ / 1 mM KCl / 320 V / zone 2*

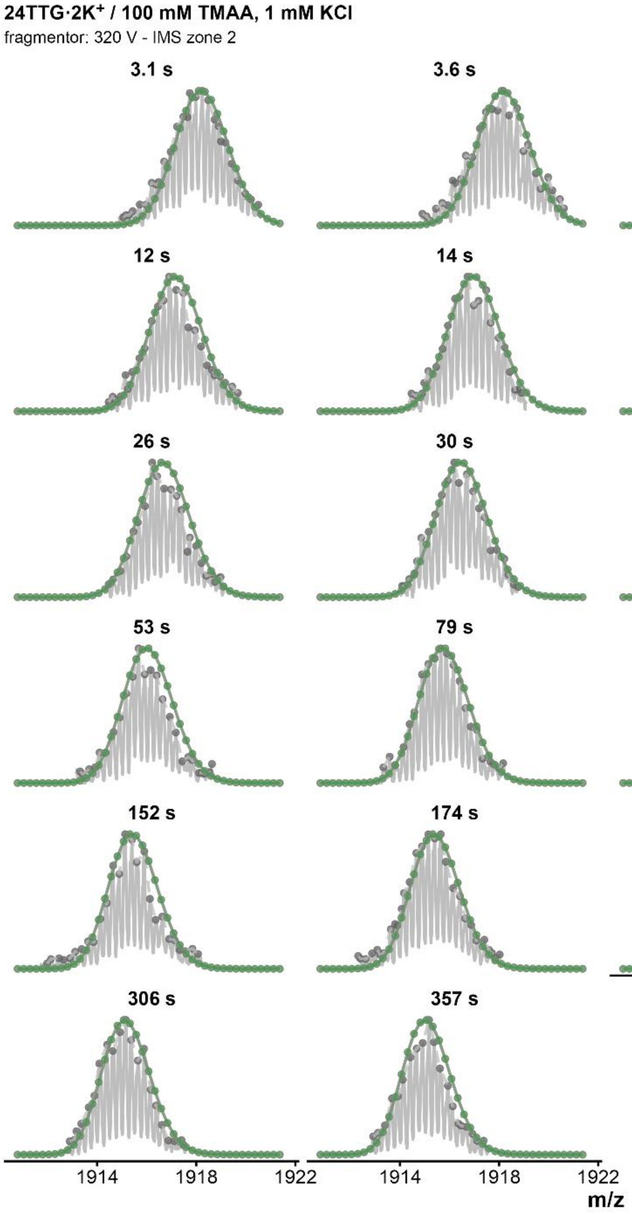


*Figure S154. Isotopic MS deconvolutions of 24TTG · 2K⁺ zone 2 (fragmentor = 320V, z=4-) in 1 mM KCl for every HDX timepoints. Green curve correspond to the fit of the whole distribution.*

### *24TTG / $MK_2$ / 0.5 mM KCl / 320 V / zone 1*

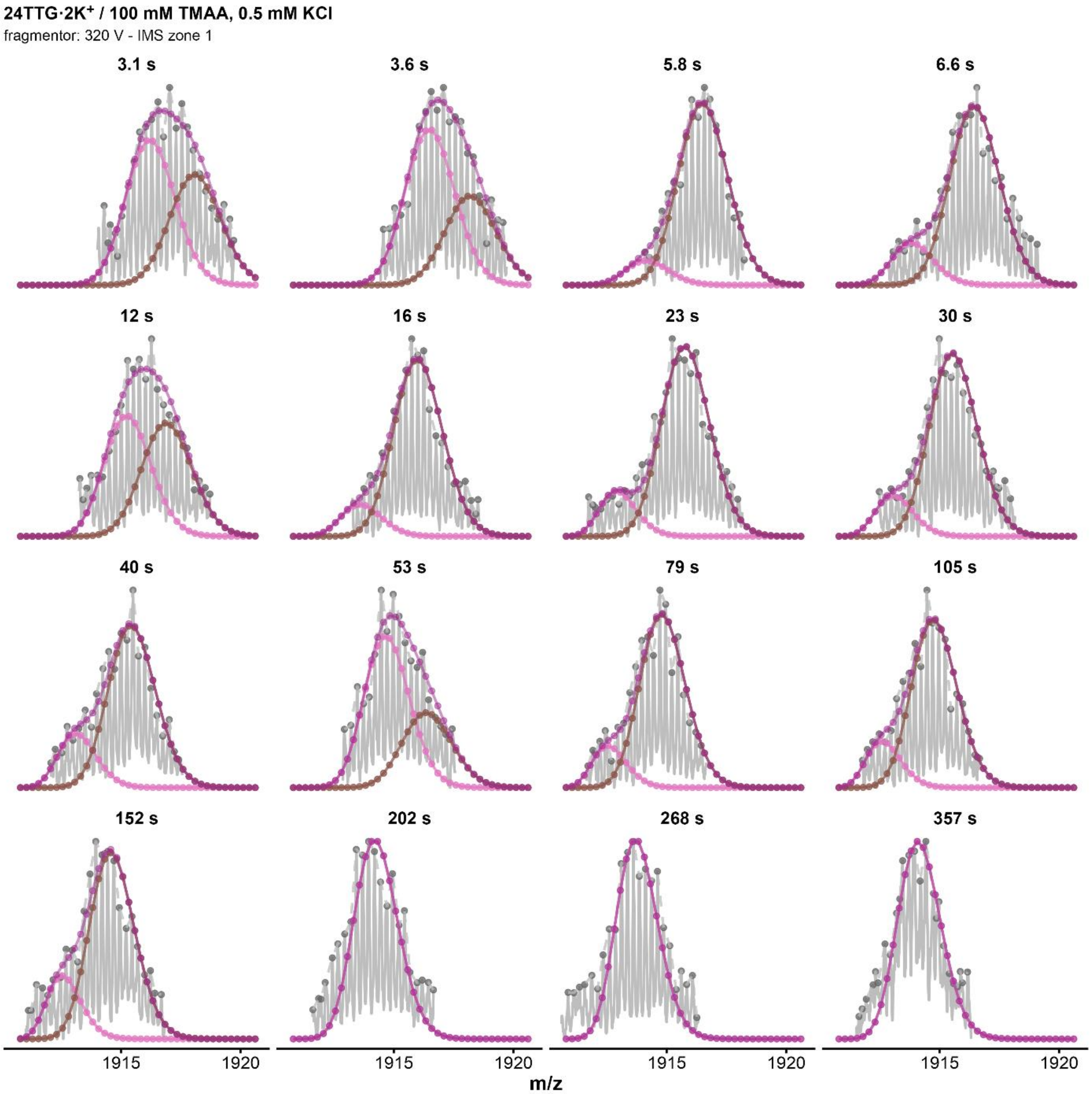


*Figure S155. Isotopic MS deconvolutions of 24TTG · $2K^+$ zone 1 (fragmentor = 320V, z=4-) in 0.5 mM KCl for every HDX timepoints. Purple curve correspond to the fit of the whole distribution distribution, the pink curve to the highly exchanged population, and the brown one to the low-exchanged population. The distribution is considered monomodal after 152s.*

### *24TTG / $MK_2$ / 0.5 mM KCl / 320 V / zone 2*

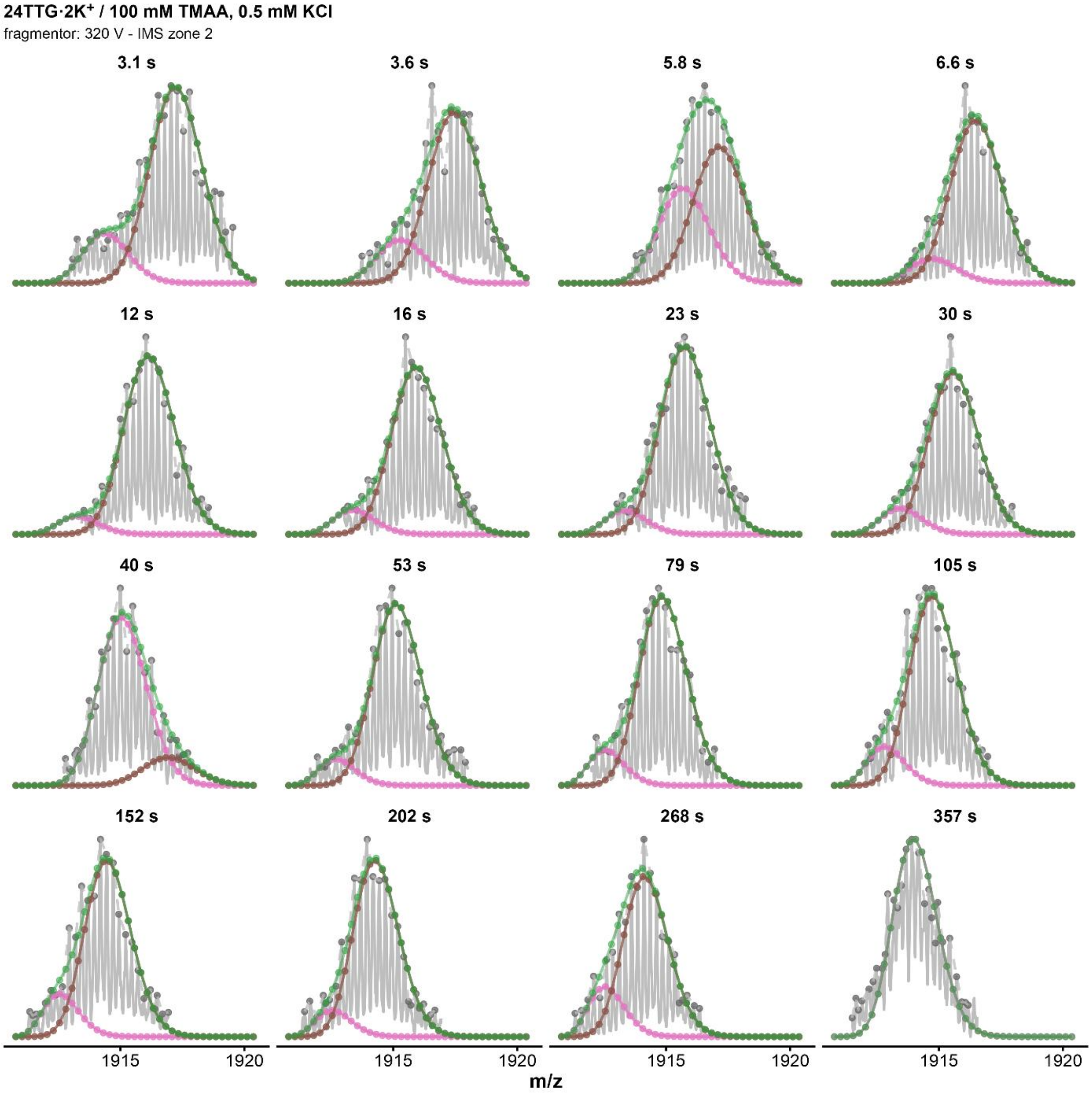


*Figure S156. Isotopic MS deconvolutions of 24TTG · $2K^+$ zone 2 (fragmentor = 320V, z=4-) in 0.5 mM KCl for every HDX timepoints. Green curve correspond to the fit of the whole distribution, the pink curve to the highly exchanged population, and the brown one to the low-exchanged population. The distribution is considered monomodal after 268s.*

### *24TTG / $MK_2$ / 0.1 mM KCl / 320 V / zone 1*

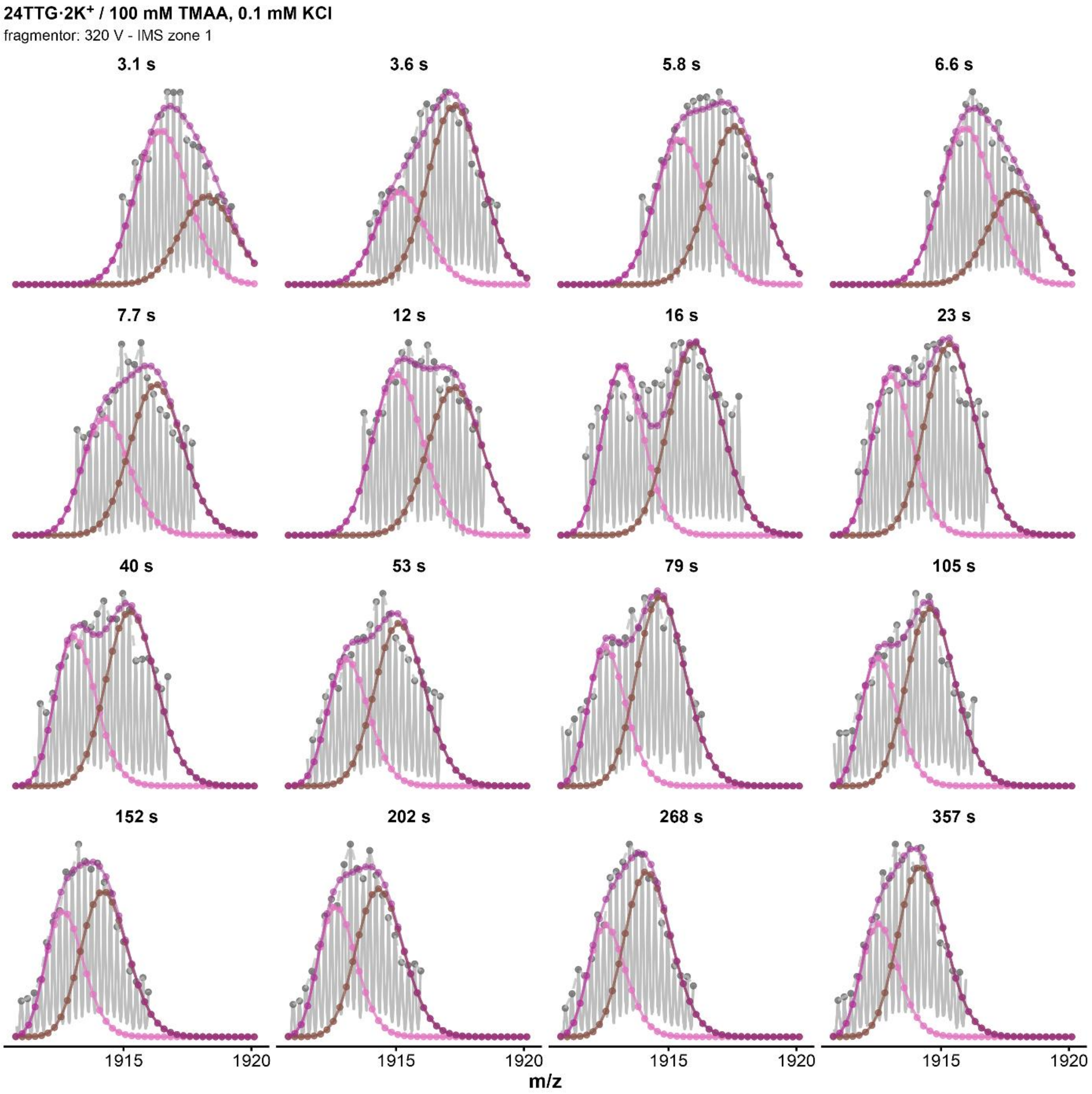


*Figure S157. Isotopic MS deconvolutions of 24TTG · 2K⁺ zone 1 (fragmentor = 320V, z=4-) in 0.1 mM KCl for every HDX timepoints. Purple curve correspond to the fit of the whole distribution.*

### *24TTG / MK$_2$ / 0.1 mM KCl / 320 V / zone 2*

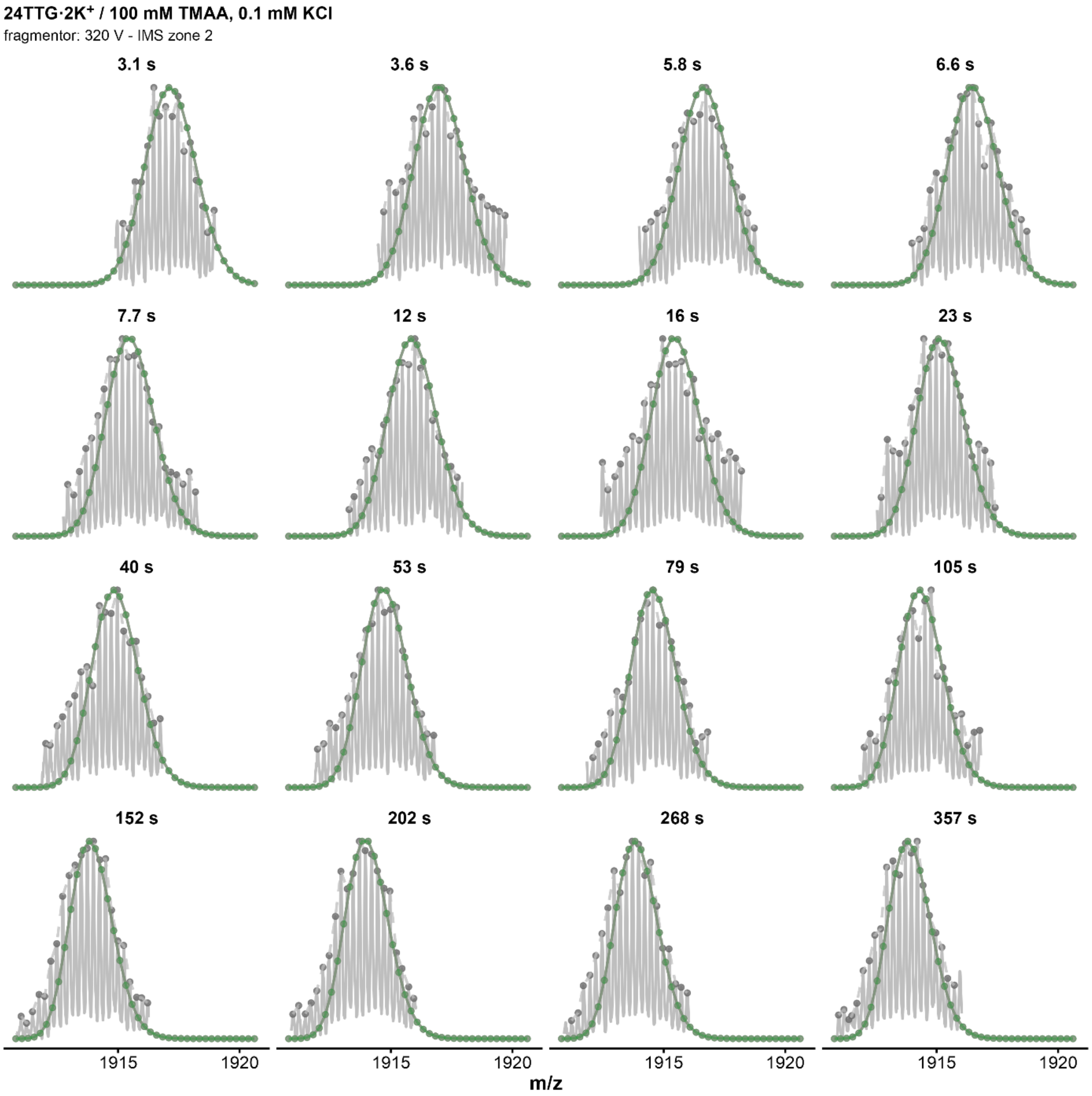


*Figure S158. Isotopic MS deconvolutions of 24TTG · 2K$^+$ zone 2 (fragmentor = 320V, z=4-) in 0.1 mM KCl for every HDX timepoints. Green curve correspond to the fit of the whole distribution*

### *222T / MK / 0.1 mM KCl / 320 V / zone 1*

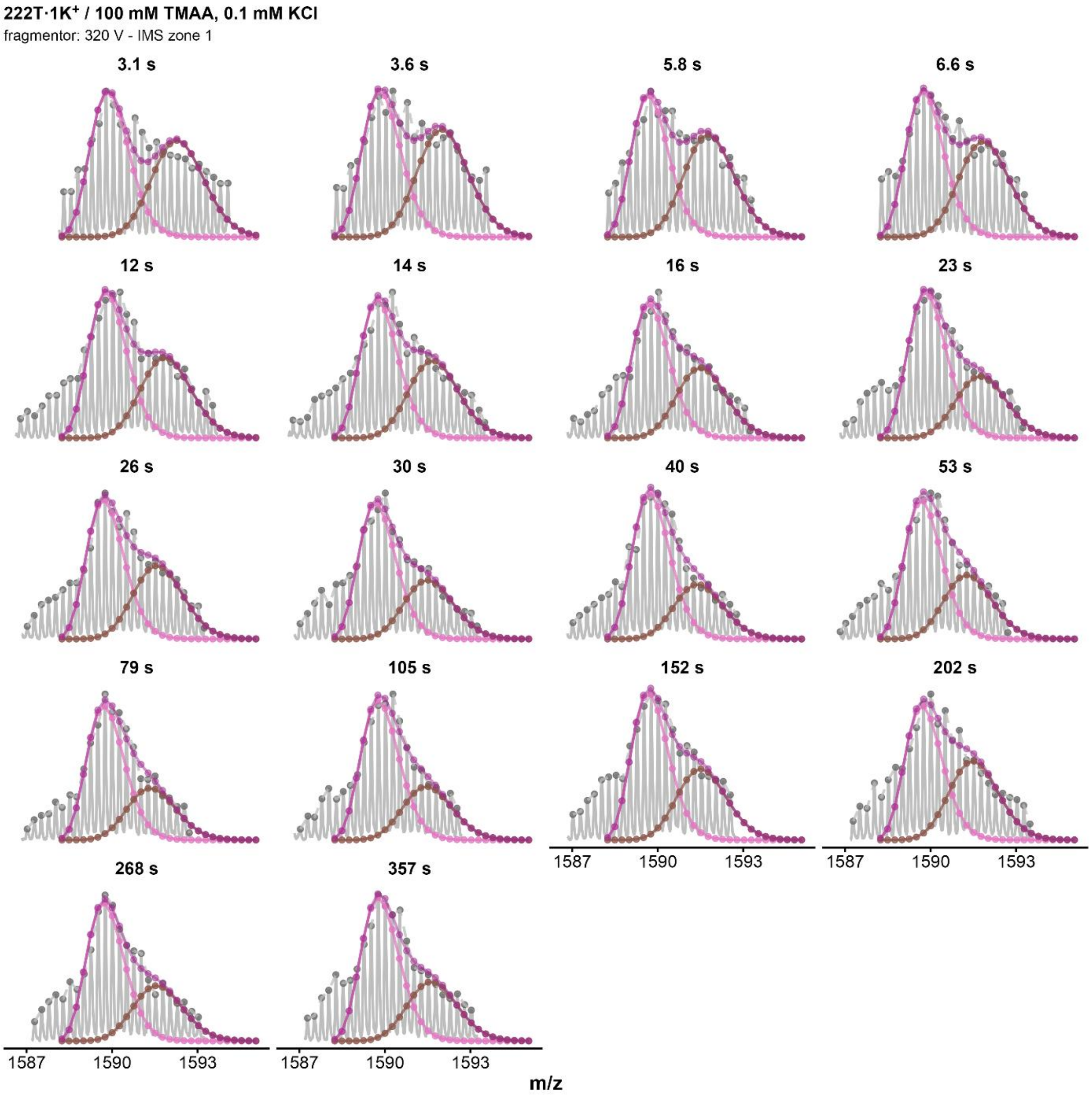


*Figure S159. Isotopic MS deconvolutions of 222T · 1K⁺ zone 1 (fragmentor = 320V, z=4-) in 0.1 mM KCl for every HDX timepoints. Purple curve correspond to the fit of the whole distribution, the pink curve to the highly exchanged population, and the brown one to the low-exchanged population.*

### *222T / MK / 0.1 mM KCl / 320 V / zone 2*

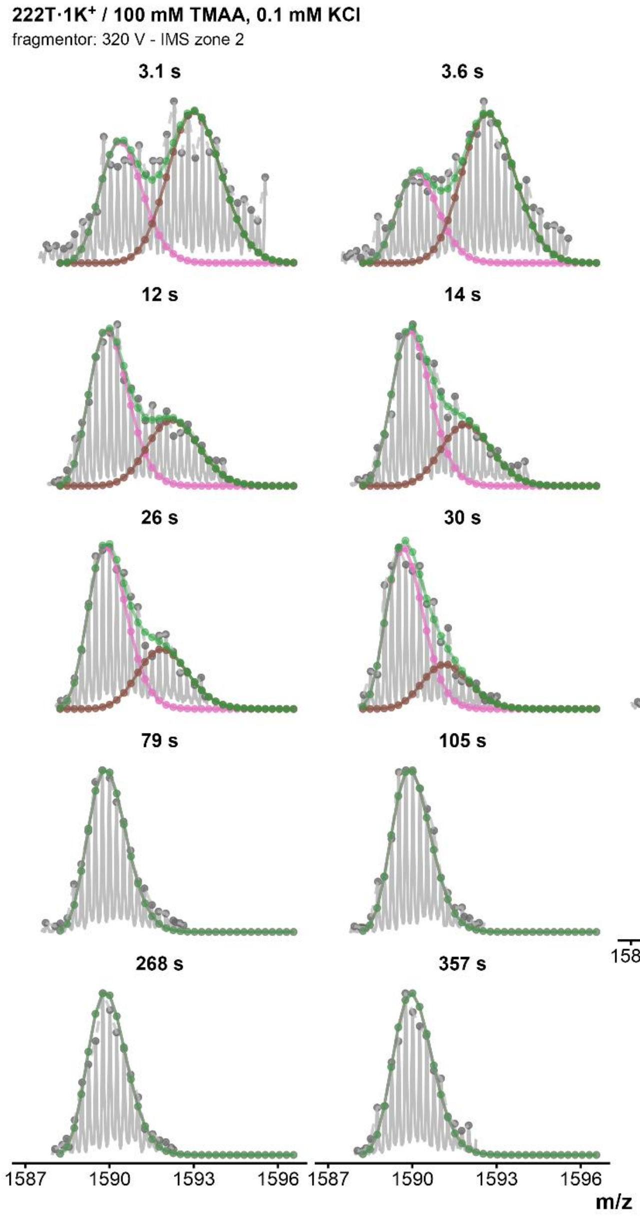


*Figure S160. Isotopic MS deconvolutions of 222T · 1K⁺ zone 2 (fragmentor = 320V, z=4-) in 0.1 mM KCl for every HDX timepoints. Green curve correspond to the fit of the whole distribution, the pink curve to the highly exchanged population, and the brown one to the low-exchanged population. The distribution is considered monomodal after 40s.*

*222T / $MK_2$ / 0.1 mM KCl / 320 V / zone 1*

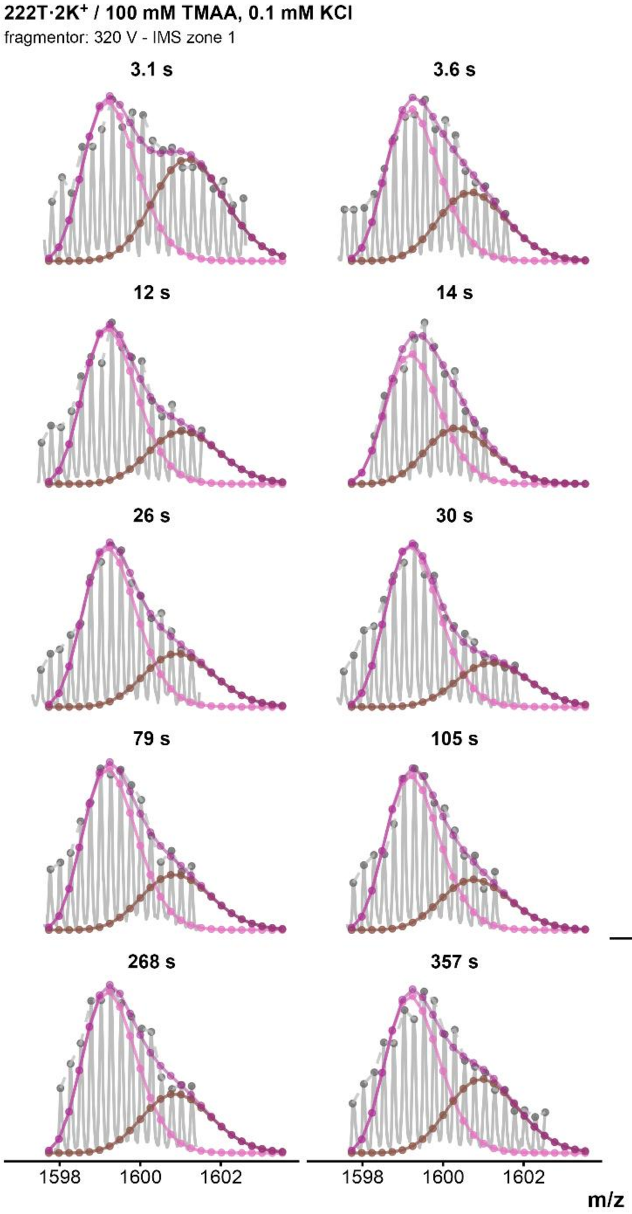


*Figure S161. Isotopic MS deconvolutions of 222T·$2K^+$ zone 1 (fragmentor = 320V, z=4-) in 0.1 mM KCl for every HDX timepoints. Purple curve correspond to the fit of the whole distribution, the pink curve to the highly exchanged population, and the brown one to the low-exchanged population.*

*222T / MK₂ / 0.1 mM KCl / 320 V / zone 2*

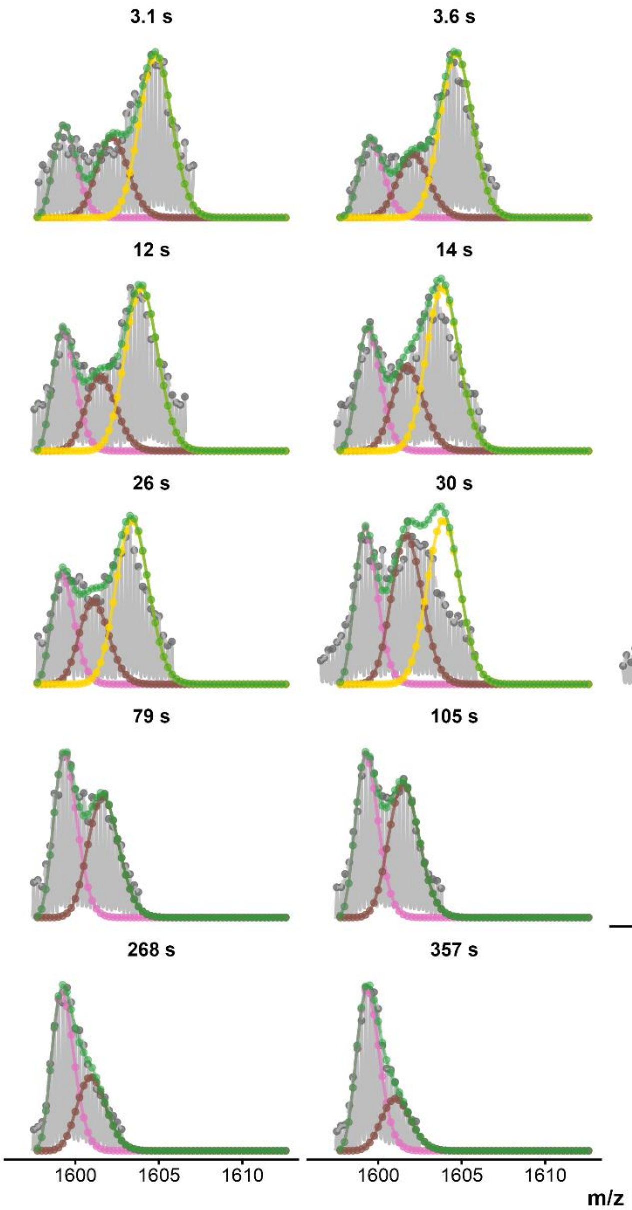


*Figure S162. Trimodal MS deconvolutions of 222T · 2K⁺ zone 2 (fragmentor = 320V, z=4-) in 0.1 mM KCl for every HDX timepoints. Green curve correspond to the fit of the whole distribution, the pink curve to the contaminated species, the brown curve to the highly exchanged population and the yellow one to the low-exchanged population.*

### *222T / $MK_2$ / 0.1 mM KCl / 320 V / zone 3*

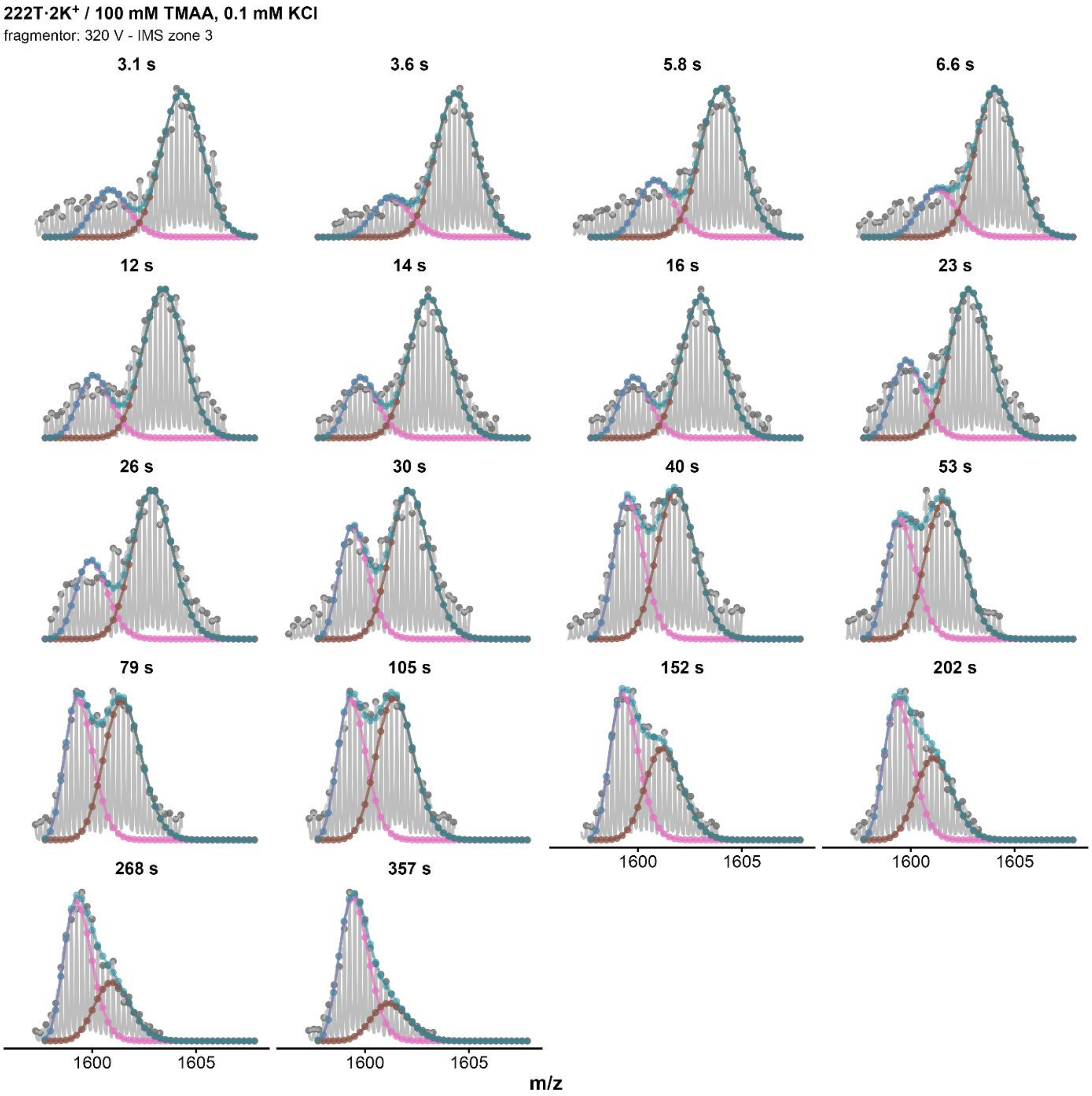


*Figure S163. Isotopic MS deconvolutions of 222T·$2K^+$ zone 3 (fragmentor = 320V, z=4-) in 0.1 mM KCl for every HDX timepoints. Blue curve correspond to the fit of the whole distribution, the pink curve to the highly exchanged population, and the brown one to the low-exchanged population.*

### *222T / $MK_2$ / 1 mM KCl / 320 V / zone 2*

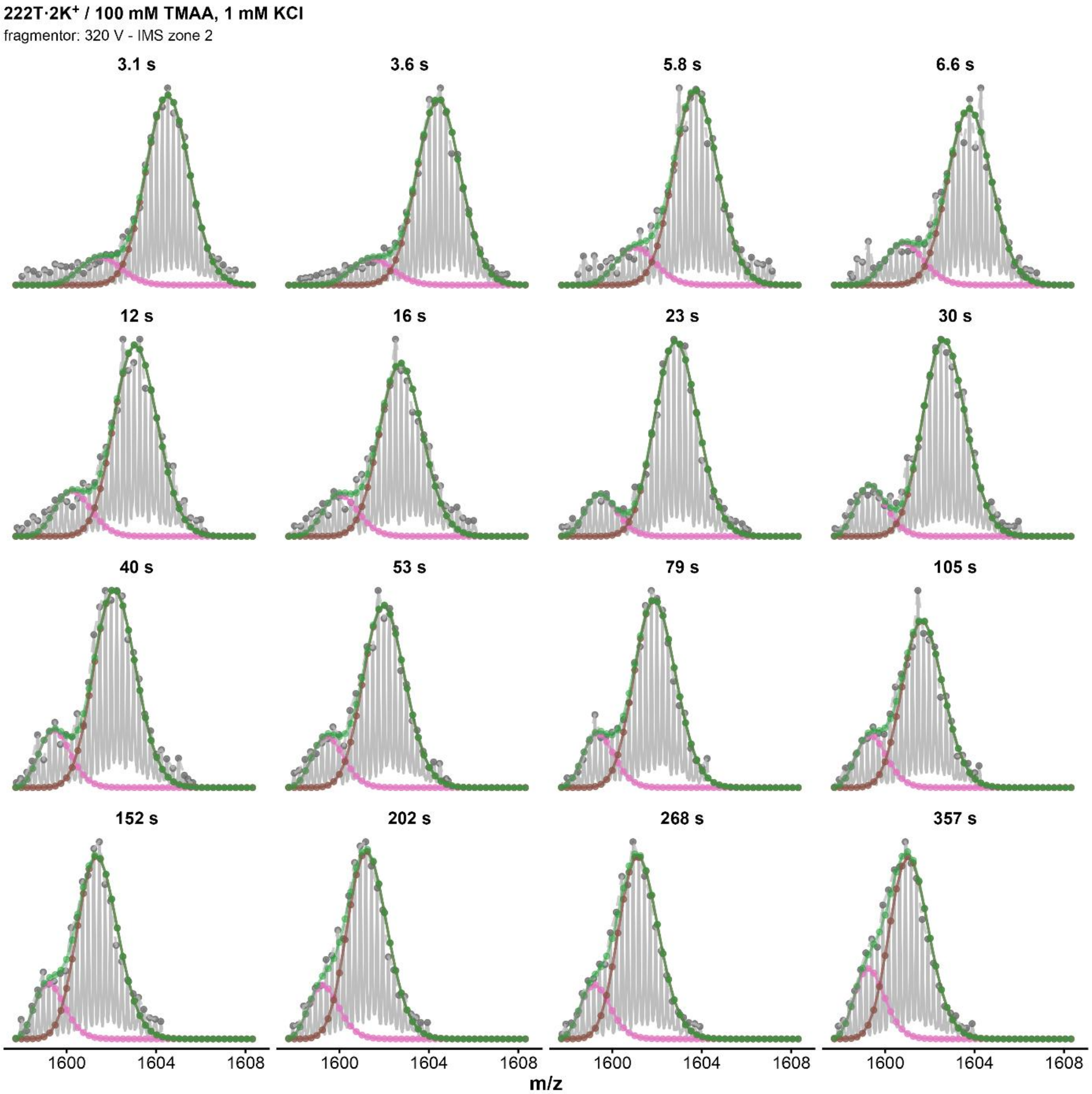


*Figure S164. Isotopic MS deconvolutions of 222T·$2K^+$ zone 2 (fragmentor = 320V, z=4-) in 1 mM KCl for every HDX timepoints. Green curve correspond to the fit of the whole distribution, the pink curve to the highly exchanged population, and the brown one to the low-exchanged population.*

### *222T / $MK_2$ / 1 mM KCl / 320 V / zone 3*

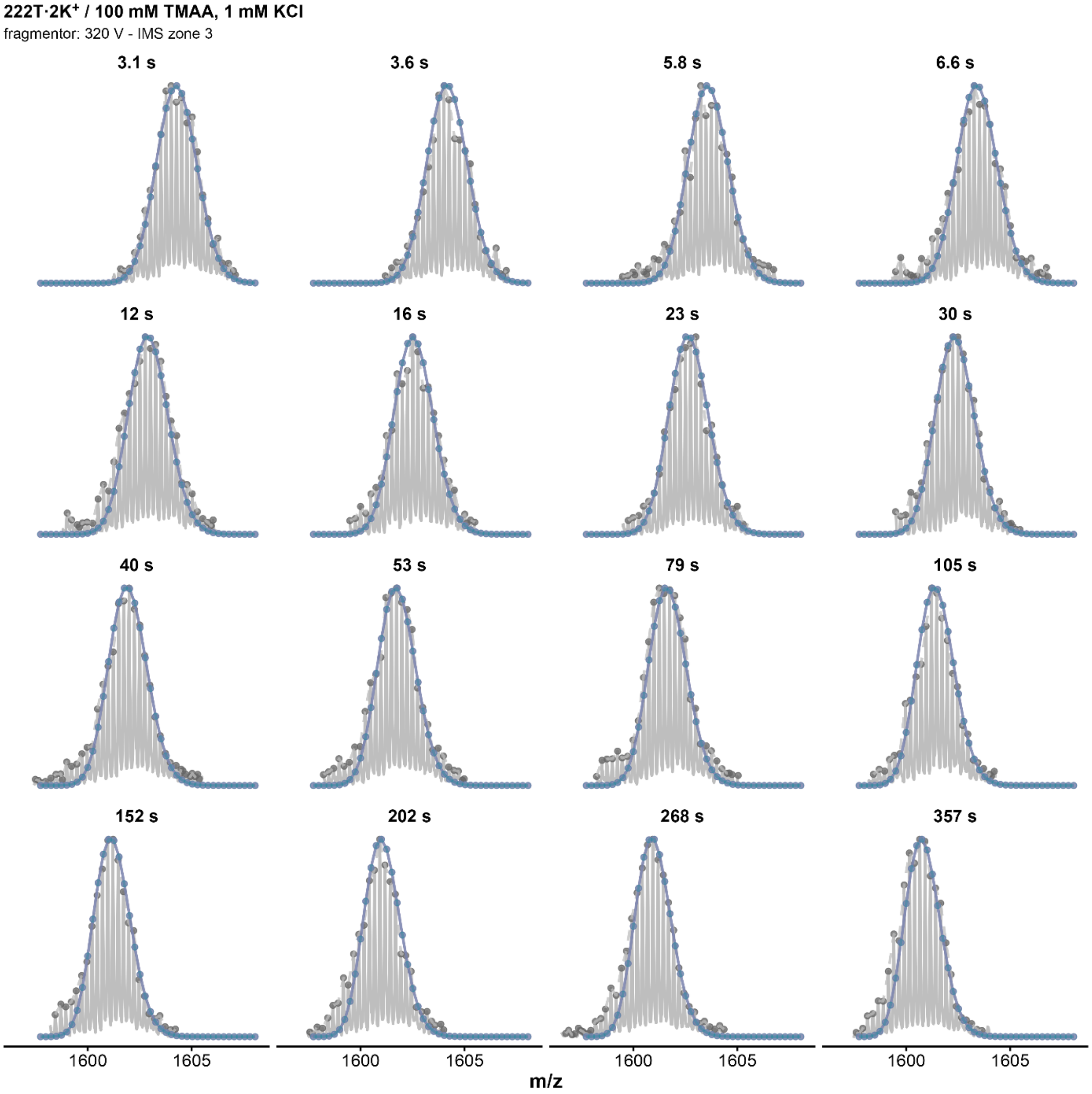


*Figure S165. Isotopic MS deconvolutions of 222T·2K⁺ zone 3 (fragmentor = 320V, z=4-) in 1 mM KCl for every HDX timepoints. Blue curve correspond to the fit of the whole distribution.*

### *Ckit87up / $MK_2$ / 1 mM KCl / 320 V / zone 1*

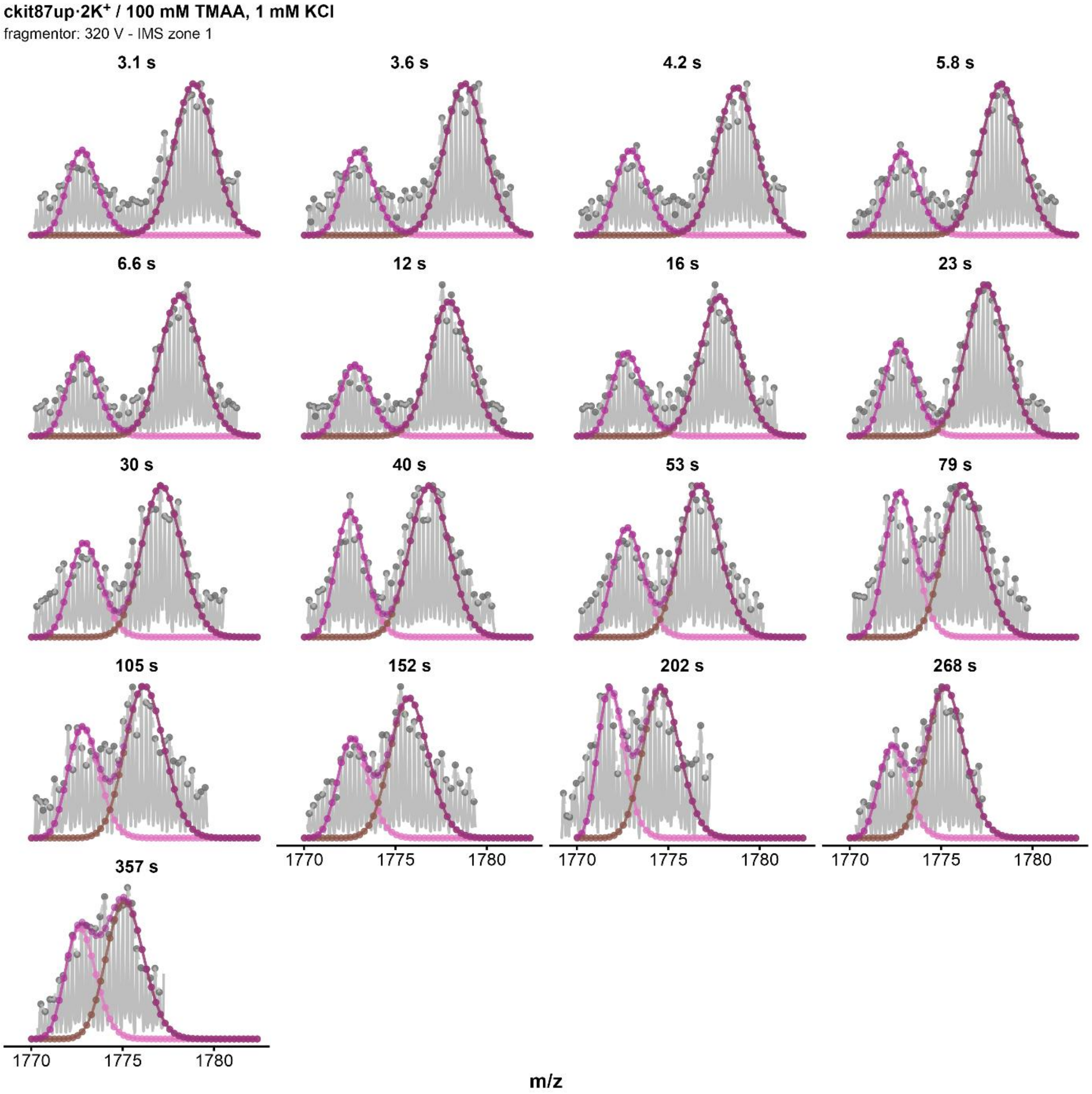


*Figure S166. Isotopic MS deconvolutions of ckit87up · 2K⁺ zone 1 (fragmentor = 320V, z=4-) in 1 mM KCl for every HDX timepoints. Purple curve correspond to the fit of the whole distribution, the pink curve to the highly exchanged population, and the brown one to the low-exchanged population.*

### *Ckit87up / $MK_2$ / 1 mM KCl / 320 V / zone 2*

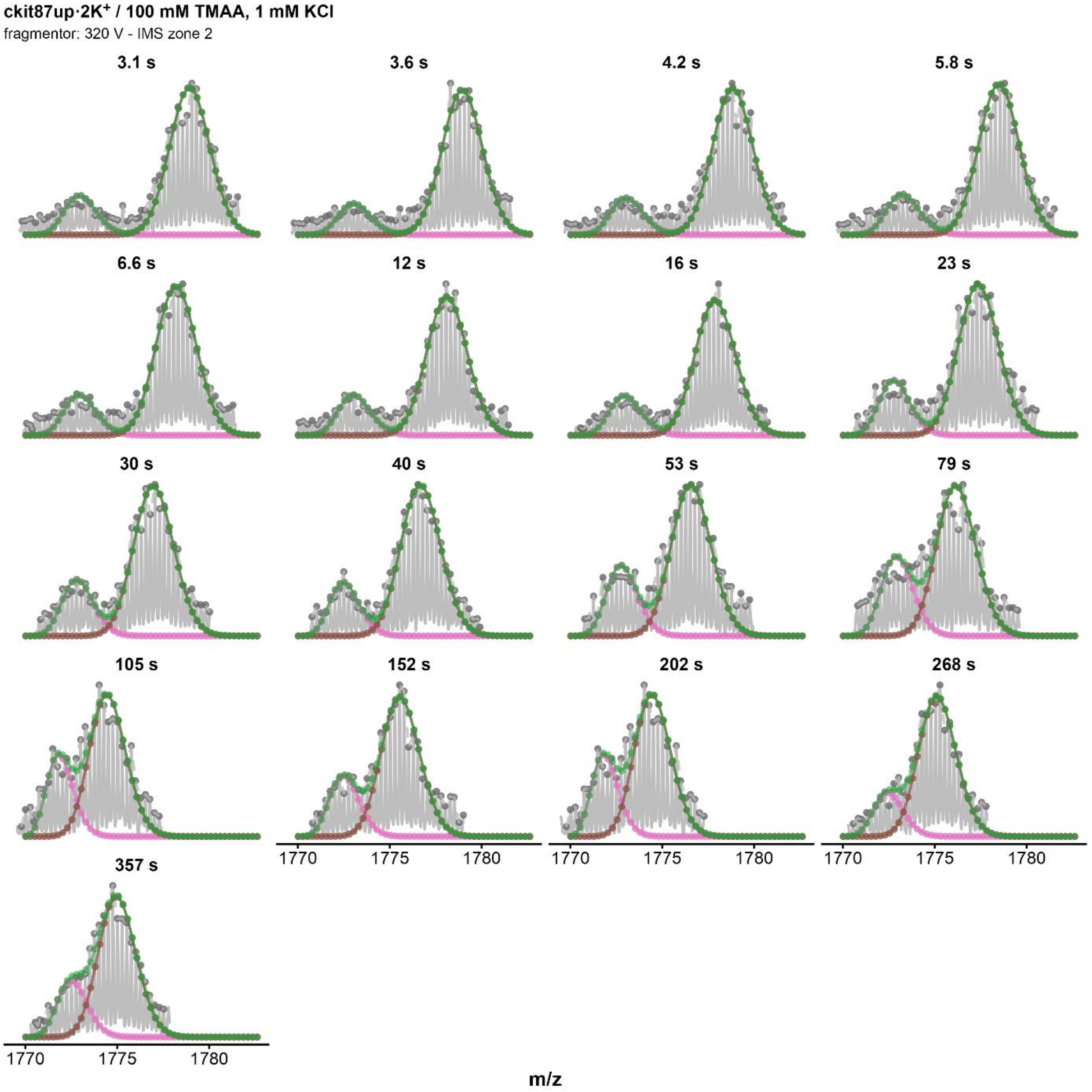


*Figure S167. Isotopic MS deconvolutions of ckit87up · $2K^+$ zone 2 (fragmentor = 320V, z=4-) in 1 mM KCl for every HDX timepoints. Green curve correspond to the fit of the whole distribution, the pink curve to the highly exchanged population, and the brown one to the low-exchanged population.*

### *Ckit87up / MK$_2$ / 1 mM KCl / 320 V / zone 3*

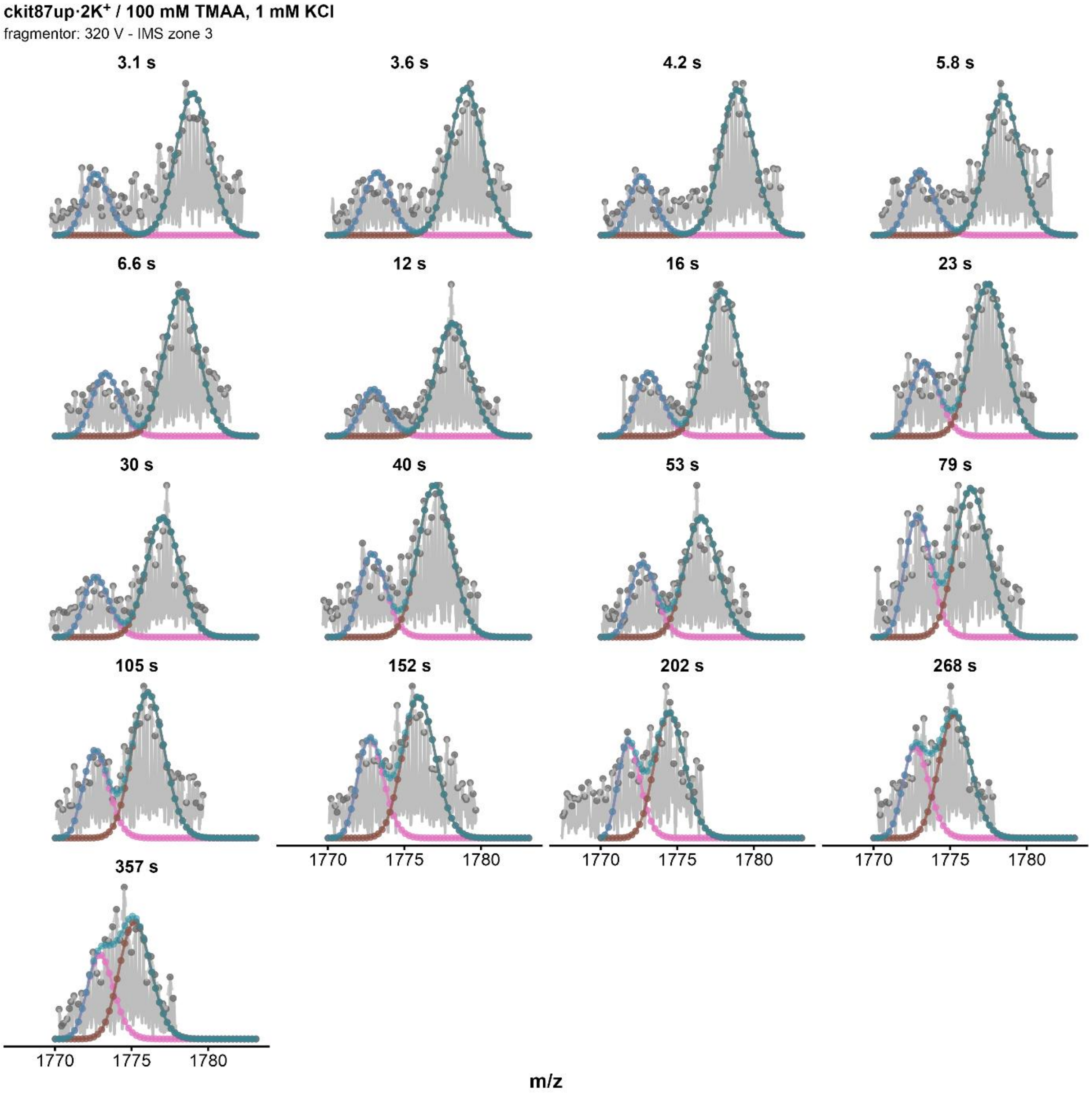


*Figure S168. Isotopic MS deconvolutions of ckit87up · 2K$^+$ zone 3 (fragmentor = 320V, z=4-) in 1 mM KCl for every HDX timepoints. Blue curve correspond to the fit of the whole distribution, the pink curve to the highly exchanged population, and the brown one to the low-exchanged population.*

### *Ckit87up / $MK_2$ / 1 mM KCl / 320 V / zone 1 – trimodal deconvolution*

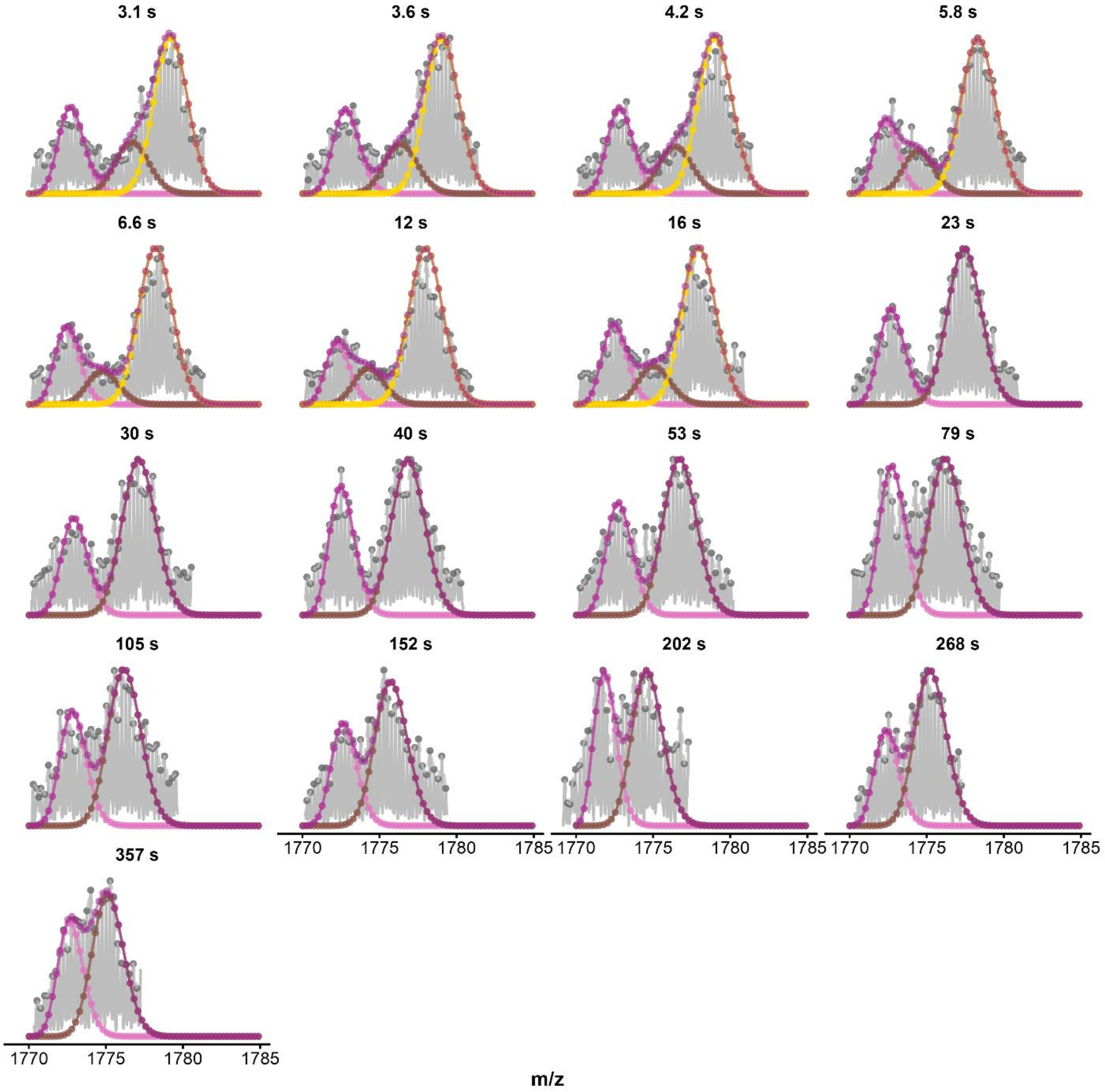


*Figure S169. Trimodal MS deconvolutions of ckit87up·2K⁺ zone 1 (fragmentor = 320V, z=4-) in 1 mM KCl for every HDX timepoints. Purple curve correspond to the fit of the whole distribution, the pink curve to the contaminated species, the brown curve to the highly exchanged population and the yellow one to the low-exchanged population.*

### *Ckit87up / MK$_2$ / 1 mM KCl / 320 V / zone 2 – trimodal deconvolution*

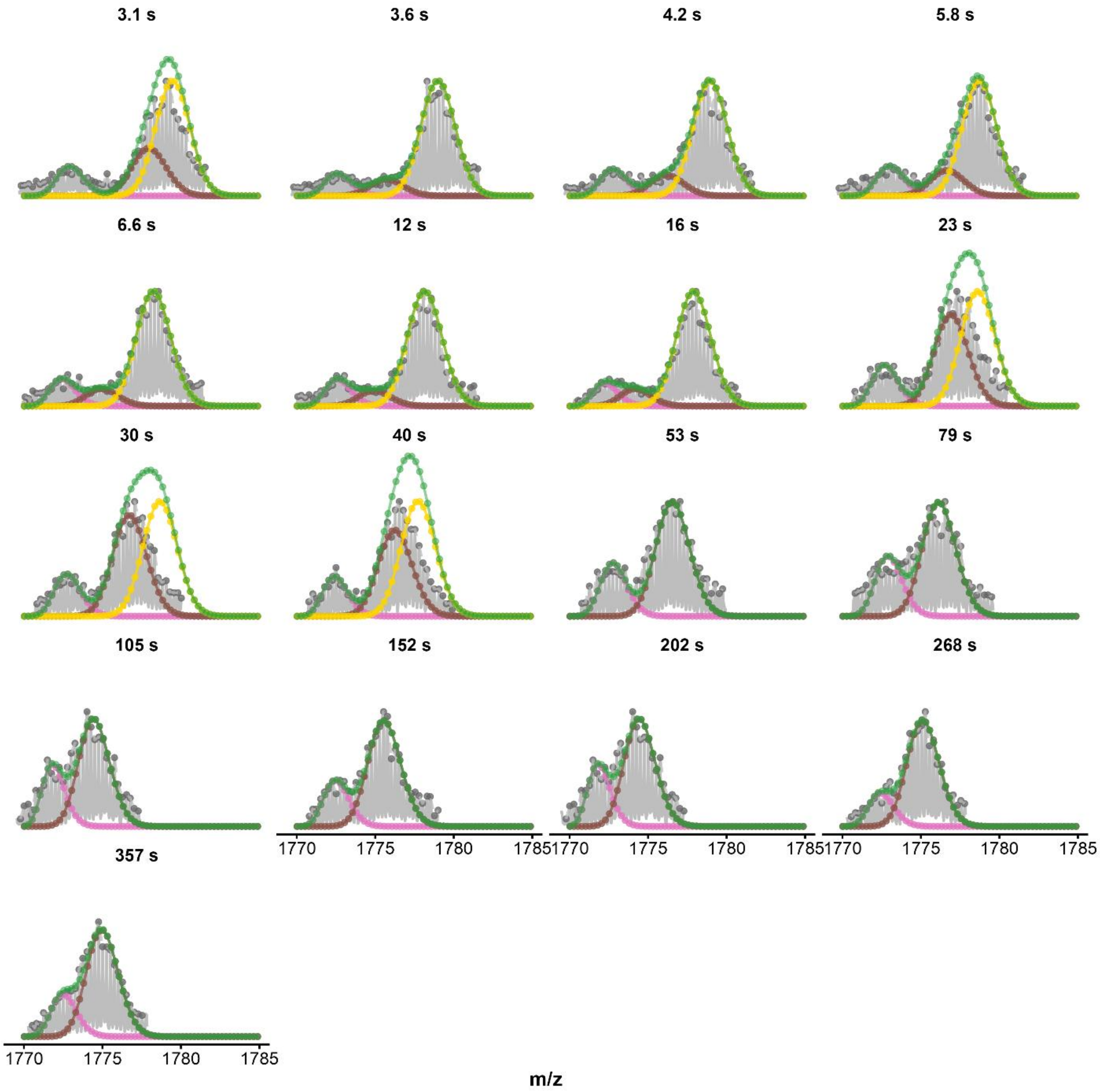


*Figure S170. Trimodal MS deconvolutions of ckit87up·2K$^+$ zone 2 (fragmentor = 320V, z=4-) in 1 mM KCl for every HDX timepoints. Green curve correspond to the fit of the whole distribution, the pink curve to the contaminated species, the brown curve to the highly exchanged population and the yellow one to the low-exchanged population.*

### *Ckit87up / $MK_2$ / 1 mM KCl / 320 V / zone 3 – trimodal deconvolution*

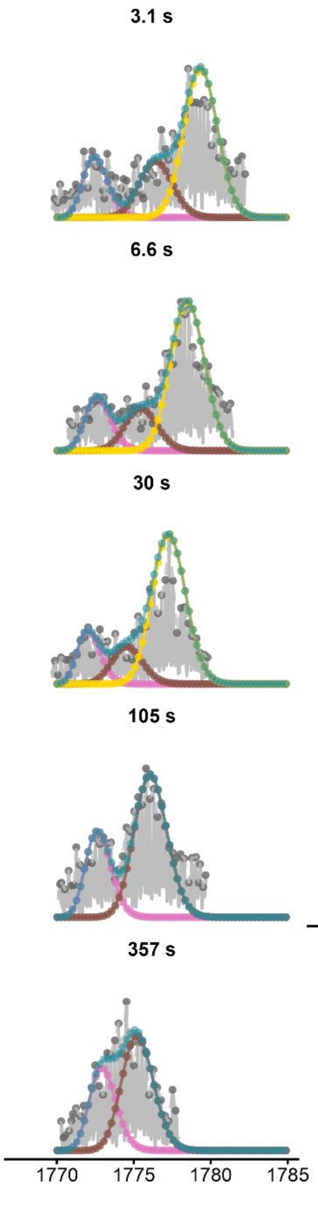


*Figure S171. Trimodal MS deconvolutions of ckit87up·$2K^+$ zone 3 (fragmentor = 320V, z = 4-) in 1 mM KCl for every HDX timepoints. Blue curve correspond to the fit of the whole distribution, the pink curve to the contaminated species, the brown curve to the highly exchanged population and the yellow one to the low-exchanged population.*

### *BCl2 / MK$_2$ / 1 mM KCl / 320 V / zone 1*

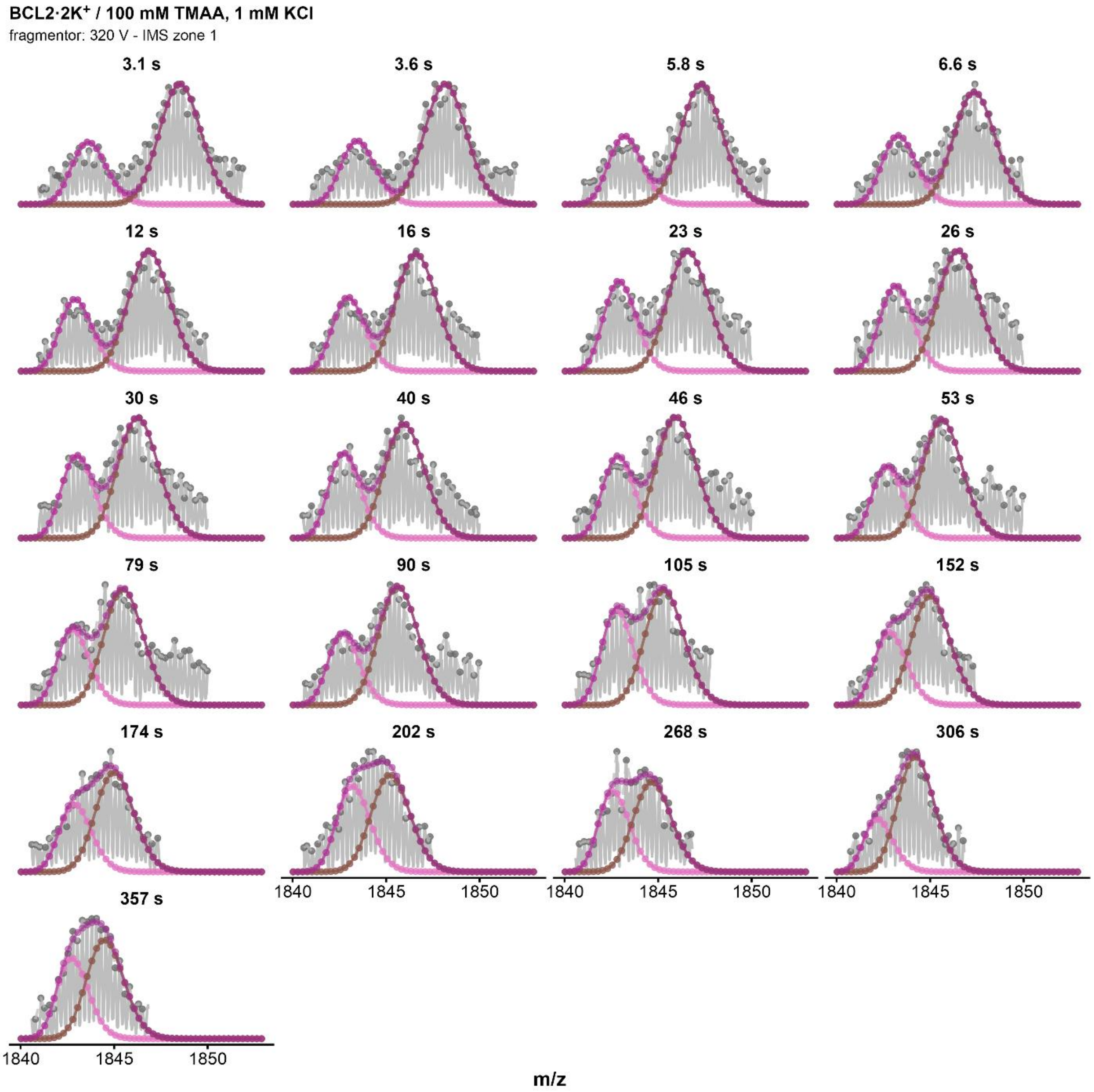


*Figure S172. Isotopic MS deconvolutions of BCL2·2K$^+$ zone 1 (fragmentor = 320V, z=4-) in 1 mM KCl for every HDX timepoints. Purple curve correspond to the fit of the whole distribution, the pink curve to the highly exchanged population, and the brown one to the low-exchanged population.*

### *BCl2 / MK₂ / 1 mM KCl / 320 V / zone 2*

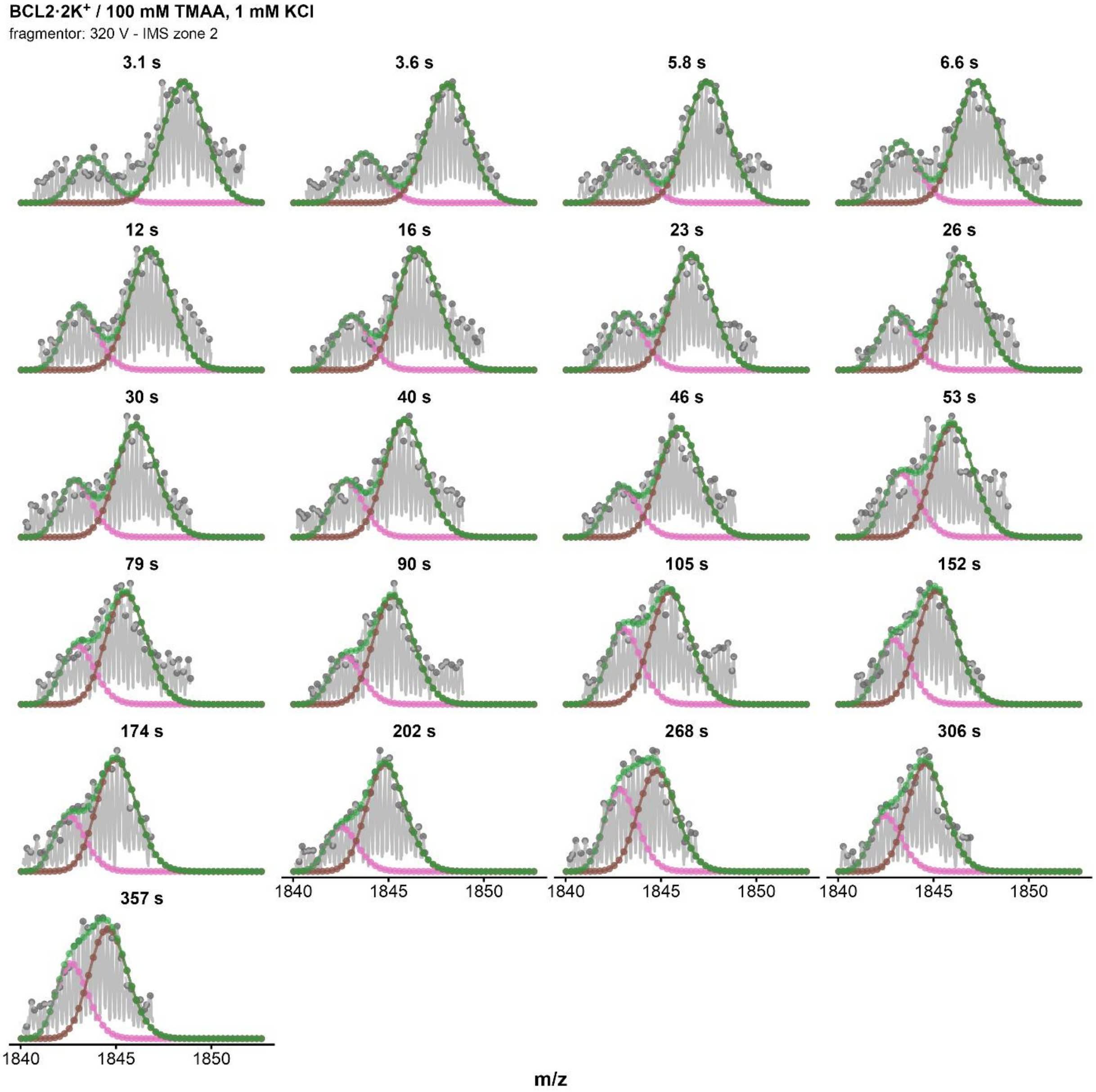


*Figure S173. Isotopic MS deconvolutions of BCL2·2K⁺ zone 2 (fragmentor = 320V, z=4-) in 1 mM KCl for every HDX timepoints. Green curve correspond to the fit of the whole distribution, the pink curve to the highly exchanged population, and the brown one to the low-exchanged population.*

### EX2 exchange kinetics

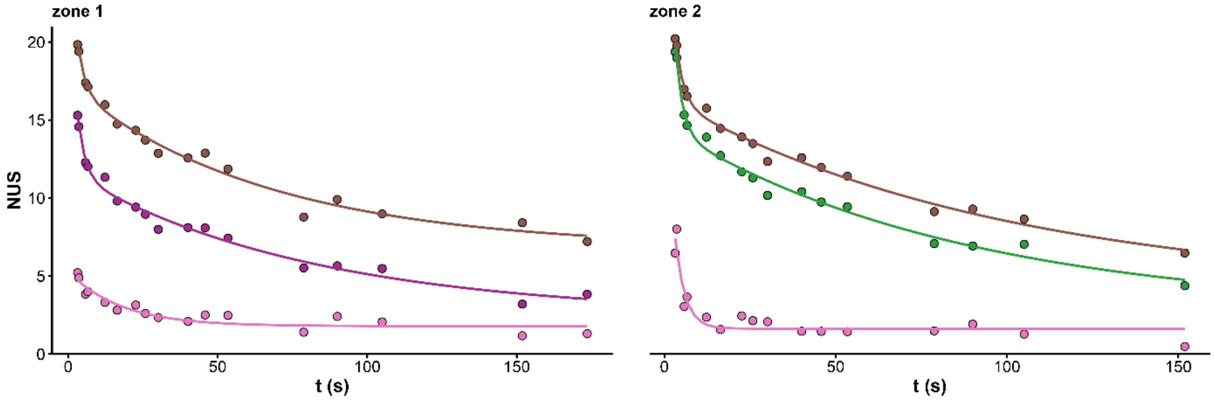


*Figure S174. Exchange kinetics of the overall (purple for zone 1, green for zone 2) and deconvoluted distributions (pink and brown) of 23TAG·2K⁺ (fragmentor = 370V, z=4-) in 1 mM KCl.*

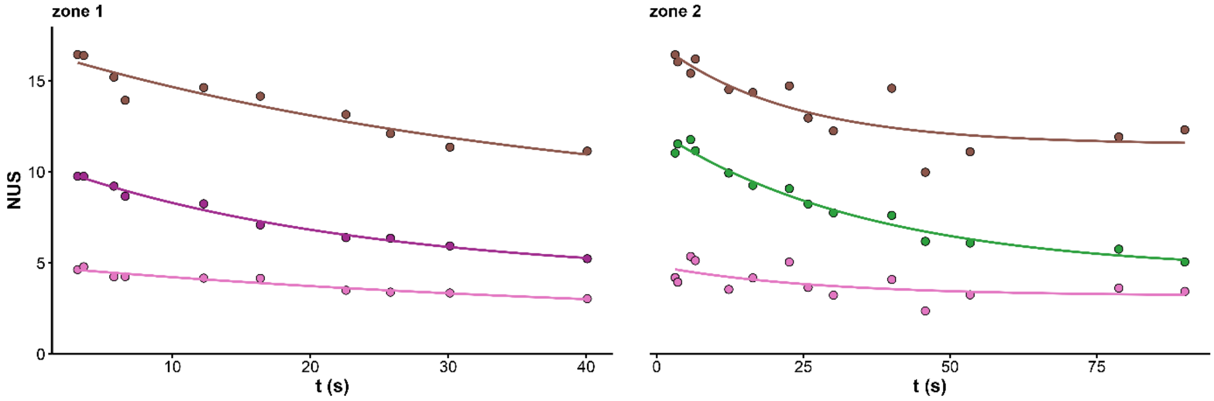


*Figure S175. Exchange kinetics of the overall (purple for zone 1, green for zone 2, blue for zone 3) and deconvoluted distributions (pink and brown) of 5YEY·1K⁺ (fragmentor = 320V, z=4-) in 1 mM KCl.*

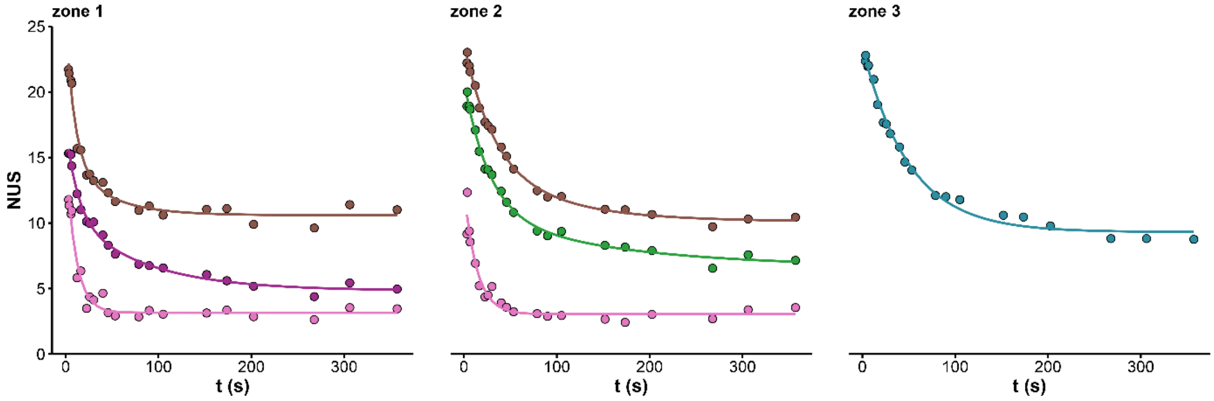


*Figure S176. Exchange kinetics of the overall (purple for zone 1, green for zone 2, blue for zone 3) and deconvoluted distributions (pink and brown) of 5YEY·2K⁺ (fragmentor = 320V, z=4-) in 1 mM KCl.*

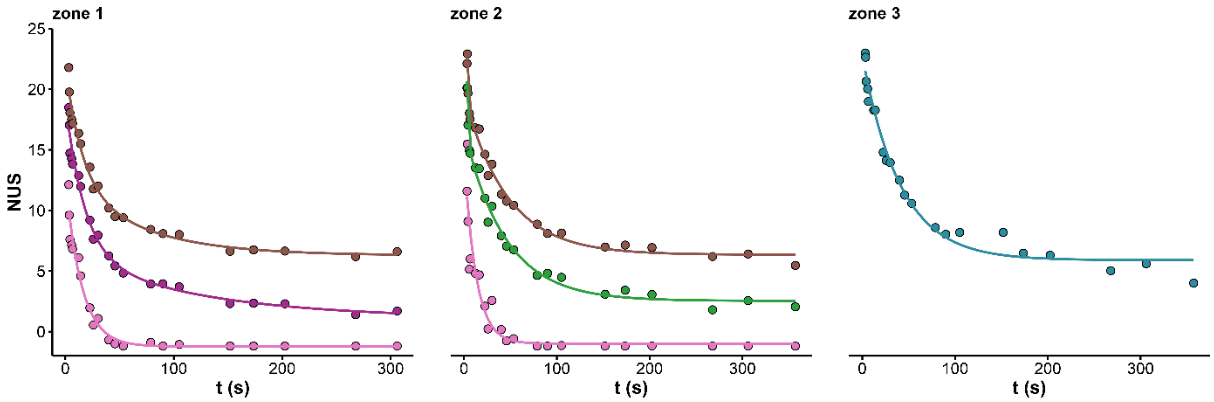


*Figure S177. Exchange kinetics of the overall (purple for zone 1, green for zone 2, blue for zone 3) and deconvoluted distributions (pink and brown) of 22GT_18T·2K⁺ (fragmentor = 320V, z=4-) in 1 mM KCl.*

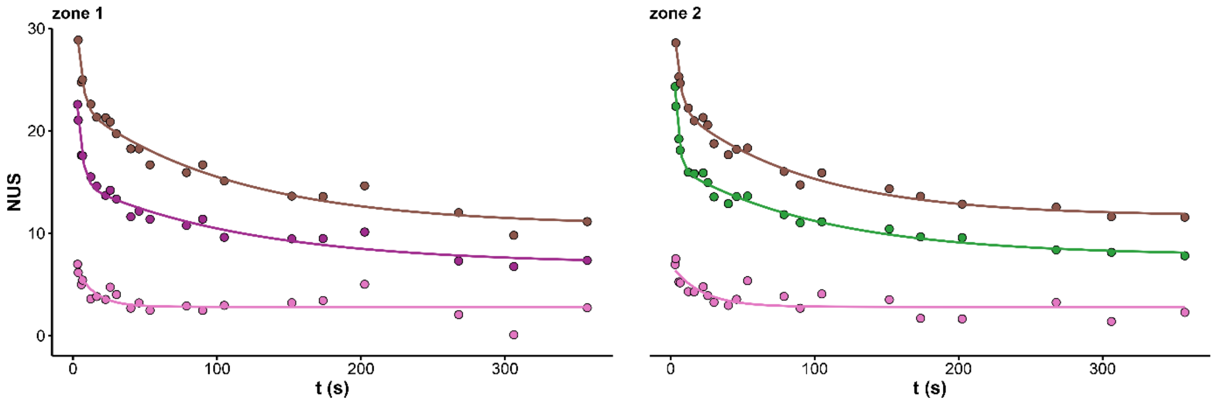


*Figure S178. Exchange kinetics of the overall (purple for zone 1, green for zone 2) and deconvoluted distributions (pink and brown) of BCL2·2K⁺ (fragmentor = 320V, z=4-) in 1 mM KCl.*

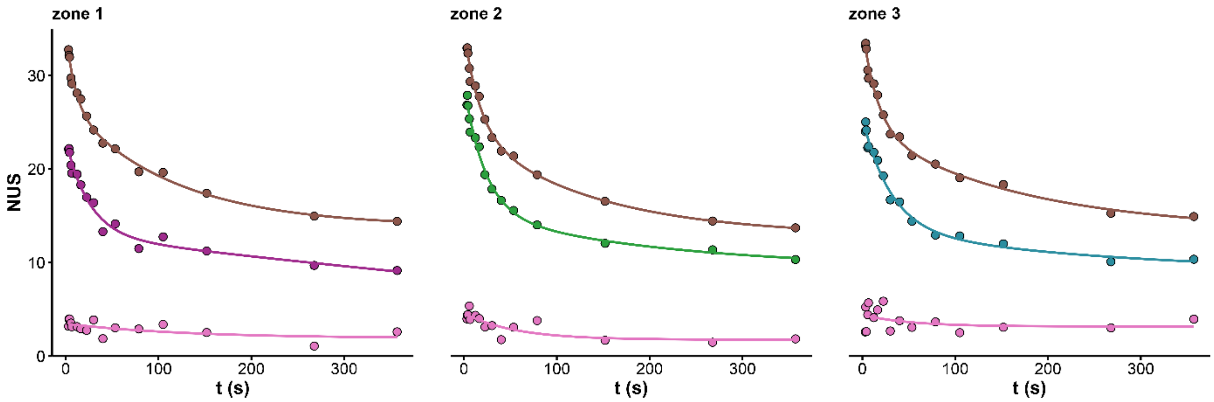


*Figure S179. Exchange kinetics of the overall (purple for zone 1, green for zone 2, blue for zone 3) and deconvoluted distributions (pink and brown) of ckit87up·2K⁺ (fragmentor = 320V, z=4-) in 1 mM KCl.*

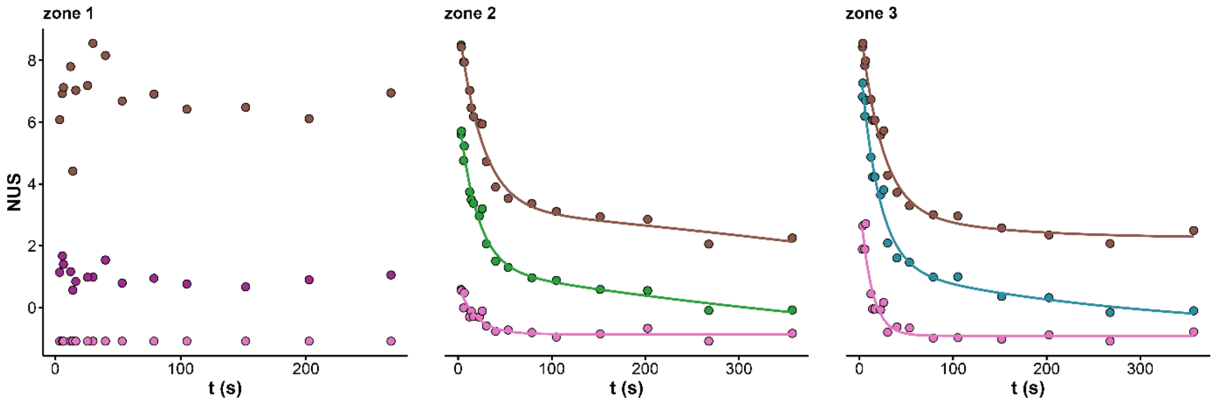


*Figure S180. Exchange kinetics of the overall (purple for zone 1, green for zone 2, blue for zone 3) and deconvoluted distributions (pink and brown) of 222T·2K⁺ (fragmentor = 320V, z=4-) in 0.1 mM KCl.*

### EX1 exchange kinetics

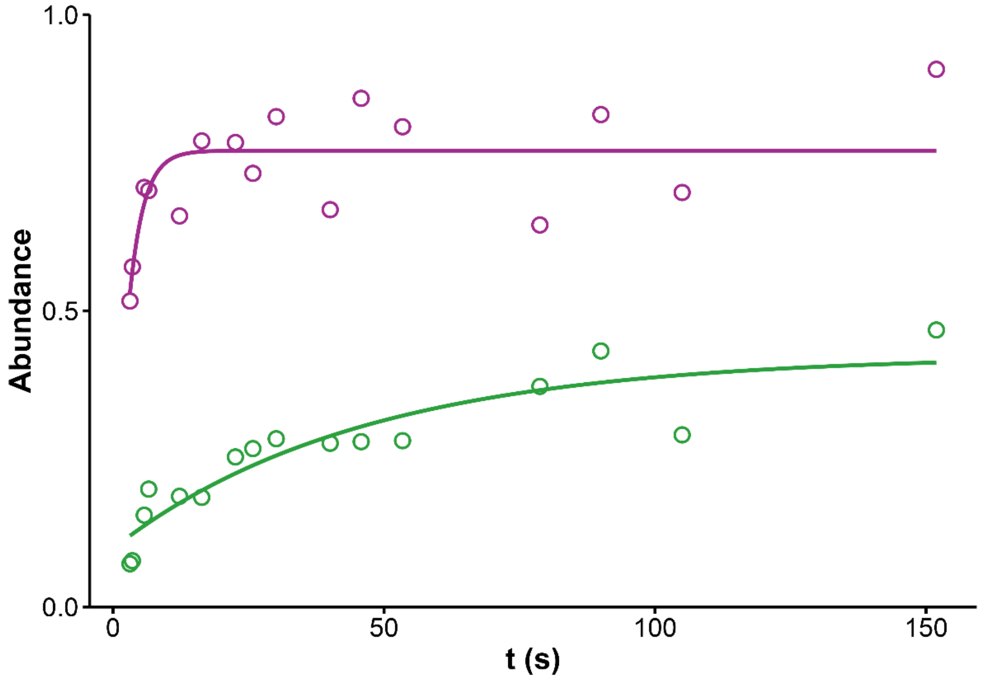


*Figure S181. EX1 dynamics of 23TAG·2K⁺ (TMAA+1mM KCl, fragmentor = 370V, z=4-). The abundance of the highly exchanged species is plotted against the time for each IMS zone (zone 1 in purple, zone 2 in green).*

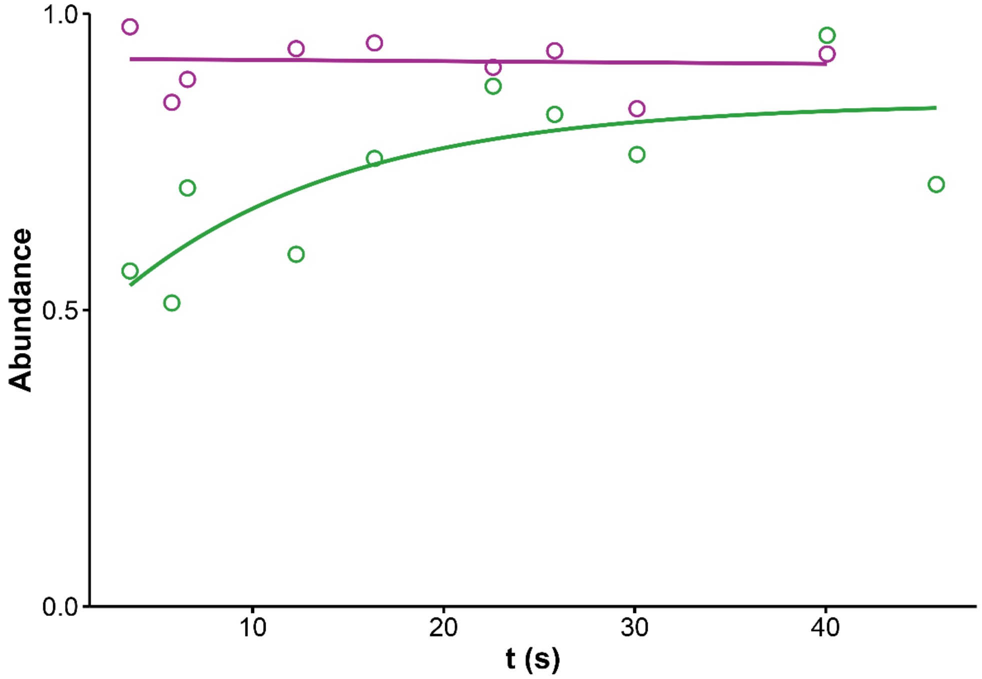


*Figure S182. EX1 dynamics of 5YEY·1K⁺ (TMAA+1mM KCl, fragmentor = 320V, z=4-). The abundance of the highly exchanged species is plotted against the time for each IMS zone (zone 1 in purple, zone 2 in green).*

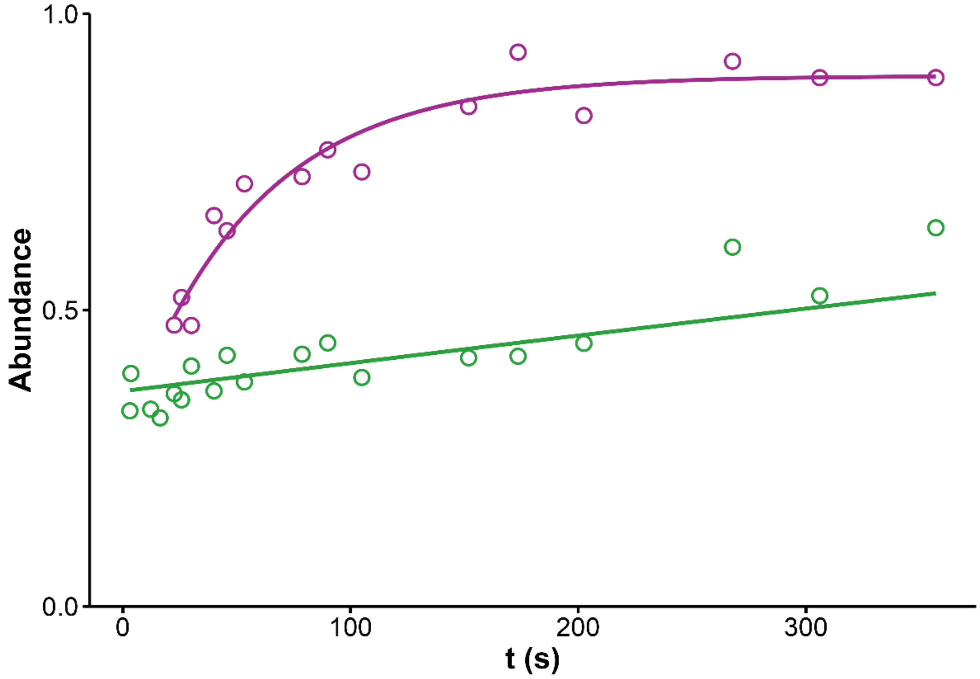


*Figure S183. EX1 dynamics of 5YEY·2K⁺ (TMAA+1mM KCl, fragmentor = 320V, z=4-). The abundance of the highly exchanged species is plotted against the time for each IMS zone (zone 1 in purple, zone 2 in green).*

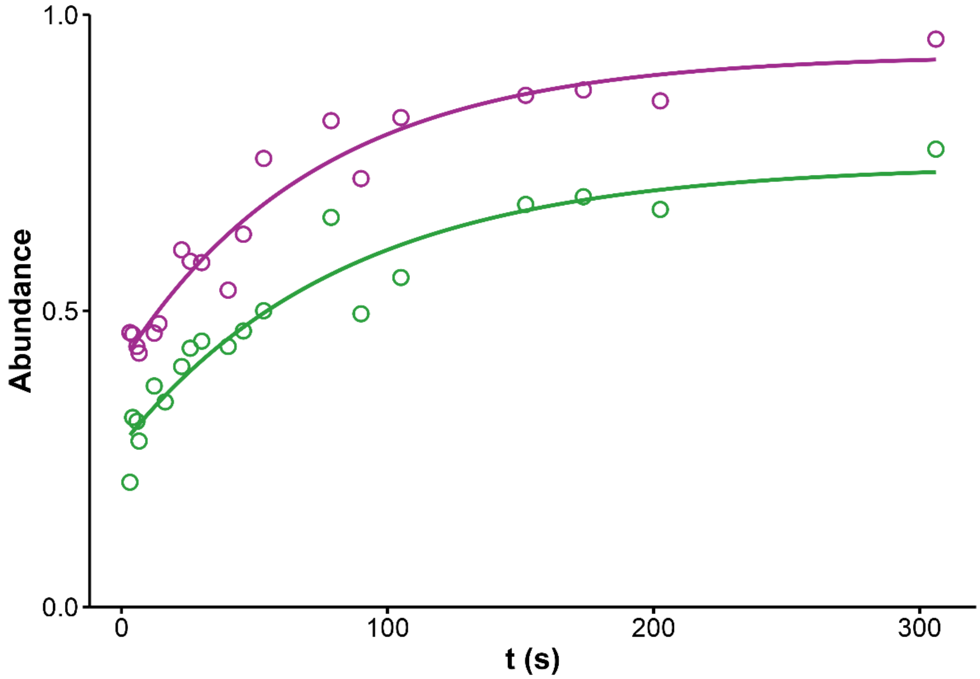


*Figure S184. EX1 dynamics of 22GT_18T·2K⁺ (TMAA+1mM KCl, fragmentor = 320V, z=4-).. The abundance of the highly exchanged species is plotted against the time for each IMS zone (zone 1 in purple, zone 2 in green).*

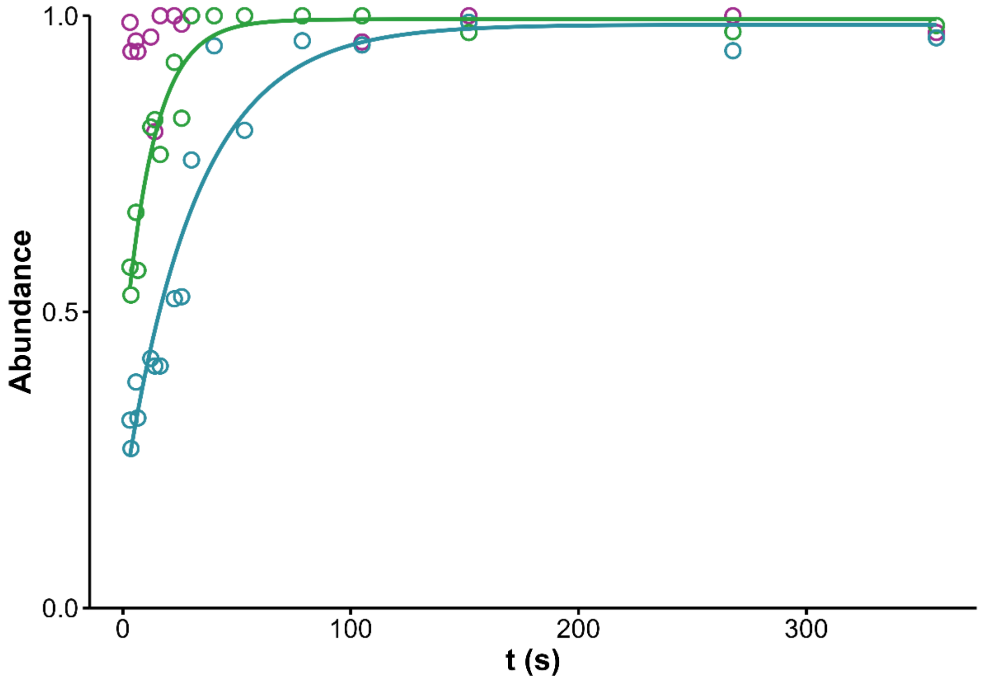


*Figure S185. EX1 dynamics of 222T·2K⁺ (TMAA+0.1mM KCl, fragmentor = 320V, z=4-). The abundance of the highly exchanged species is plotted against the time for each IMS zone (zone 1 in purple, zone 2 in green, zone 3 in blue).*

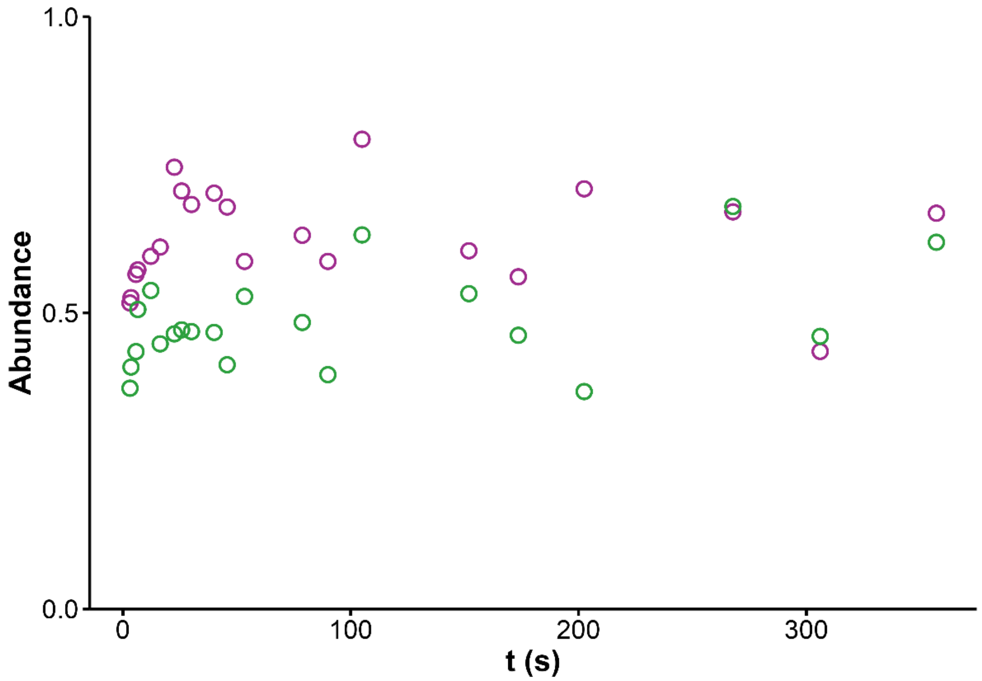


*Figure S186. EX1 dynamics of BCL2·2K⁺ (TMAA+1mM KCl, fragmentor = 320V, z=4-). The abundance of the highly exchanged species is plotted against the time for each IMS zone (zone 1 in purple, zone 2 in green).*

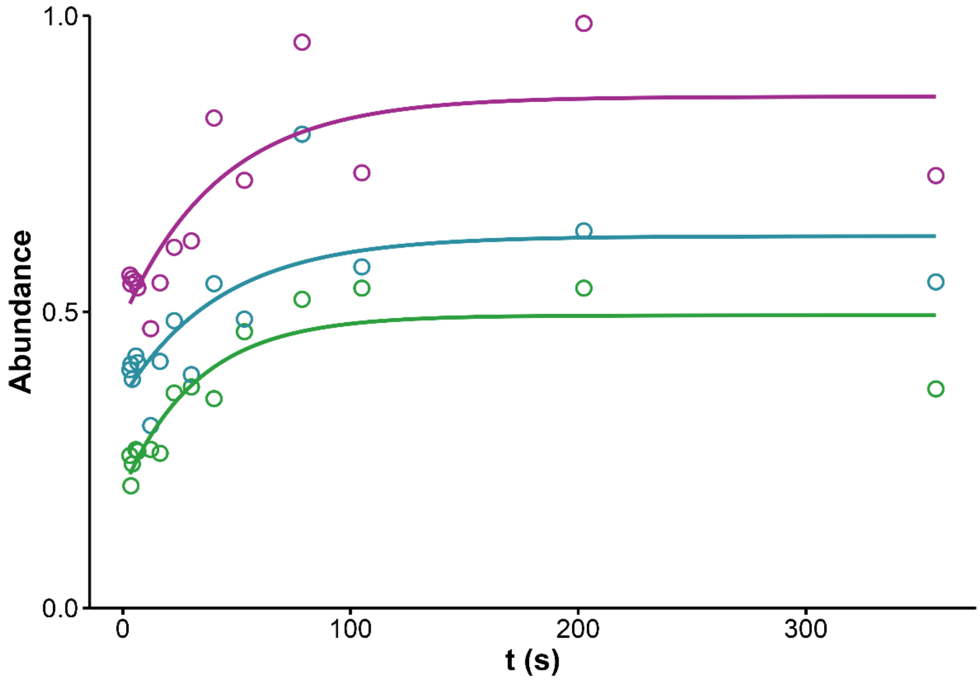


*Figure S187. EX1 dynamics of ckit87up · 2K$^+$ (TMAA+1mM KCl, fragmentor = 320V, z=4-). The abundance of the highly exchanged species is plotted against the time for each IMS zone (zone 1 in purple, zone 2 in green, zone 3 in blue).*

### Zone selections: influence of fragmentor voltage and robustness

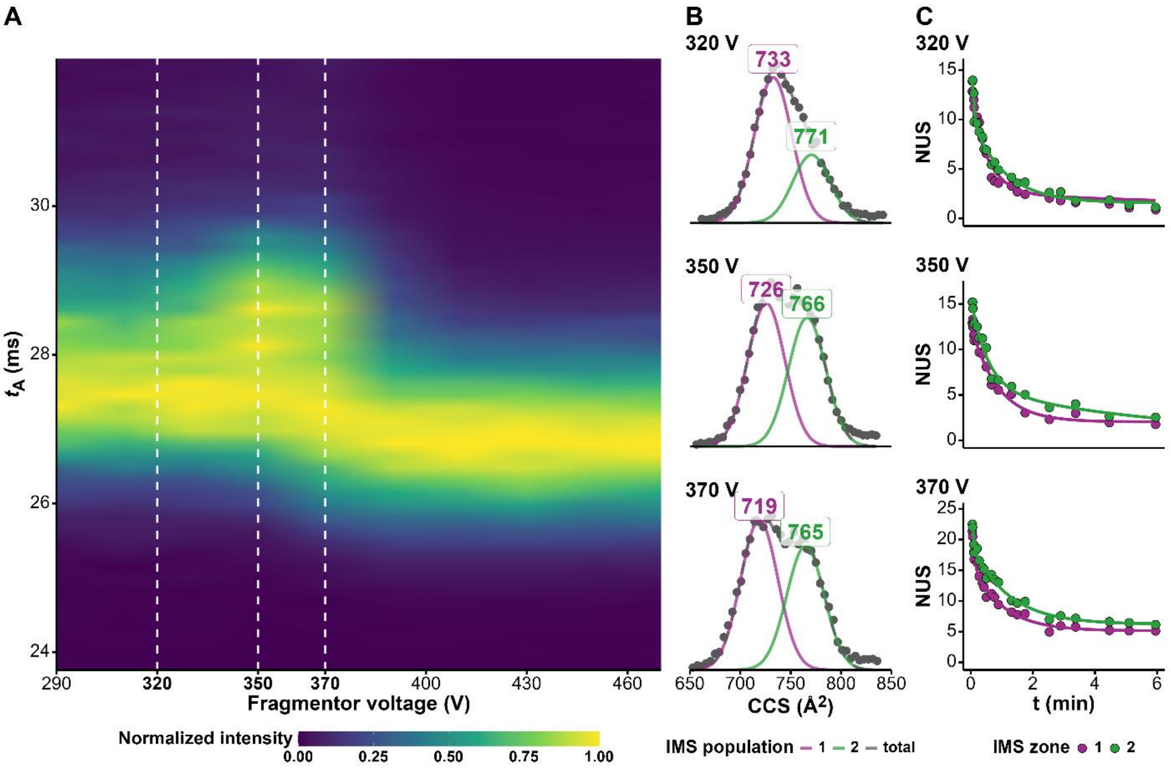


*Figure S188. Tuning the fragmentor voltage to improve the discrimination of two conformers of 23TAG·2K+: heatmap of the CIU experiment (A), corresponding CCS distributions at 320, 350 and 370 V, showing the compaction of one of the two populations, and IMS-separated HDX kinetics with improved discrimination at higher fragmentor voltage.*

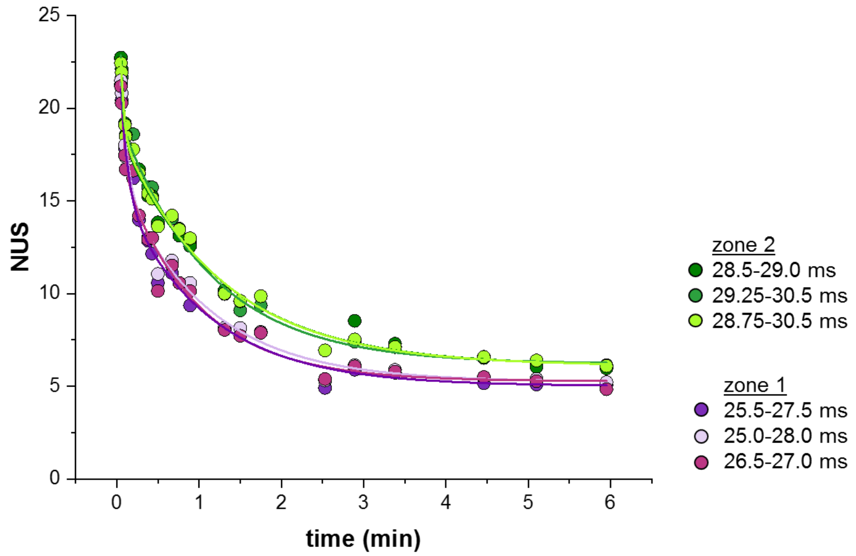


*Figure S189. Comparison of the selection of IMS zones on 23TAG·2K$^+$ in 100 mM TMAA + 1 mM KCl (fragmentor = 370V). The width of the IMS zones chosen do not significantly affect the HDX/MS kinetics as long as it is sufficiently restrictive and do not overlap on another population.*

### Probing the specificity of $K^+$ adducts on complex conformational mixes

22GT, 22CTA and 23AG_I14 are human telomeric sequences (the latter two being mutated) that have four repeats of three guanines, making them able to form 3-tetrad G4s. Despite this, 22GT mainly folds (about 60%) into a two-tetrad antiparallel basket-type G4 covered by a G-triad (Figure S190A),[17] and so does the mutated sequence 23AG_I14.[18] 22CTA adopts a 2-tetrad chair-type conformation capped by a GCGC tetrad.[19] The sequences 21G, 22AG, 23AG, whose structures have not been solved in potassium solution, are also most likely mixtures of conformers.[20] Mutating the sequence may be needed to displace the equilibrium towards the structure to be solved, e.g. dG to 7-$^{Br}$dG for 22GT and dG to 14-dI for 23AG.

In native MS experiments, the set of telomeric oligonucleotides feature both 1 and $2K^+$ peaks (Figure S190B): both 2- and 3-tetrad conformers can thus form. However, it is also possible for two tetrad conformers to bind an additional cation non-specifically.[21] CD spectra are consistent with the formation of antiparallel topologies expected of 2-tetrad conformers, but may not allow to detect minor 3-tetrad conformers (Figure S5).[6]

The conformer mixtures of both 1- and $2K^+$ species are characterized by broad ATD profiles, for which the exchange kinetics of three zones were extracted (Figure S190C et Figure S100 à Figure S117). Two salient observations can be made.

First, in each case, the IMS-resolved species generally exchange at the same rate (Figure S190D and Figure S100 to Figure S117). As expected, the quick exchange of $1K^+$ species (completed in less than 90s) is characteristic of two-tetrad conformers.[5]The greater than expected protection at early time points ($NUS_{12s} > 10$) is probably due to a base triad capping one of the G-tetrads, providing additional protection from exchange. Such secondary structures are known to be formed by 22GT and 23AG_I14.

Second, the IMS-resolved $2K^+$ species of 21G, 22AG and 22GT exchange at rates similar to their $1K^+$ counterparts (Figure S190D). In the case of $22CTA \cdot 2K^+$, however, two of the three IMS-resolved species have slower exchange kinetics. Similarly, both $23AG \cdot 2K^+$ species are more protected than the $1K^+$ ones.

Taken together, these results suggest that 21G and 22AG predominantly fold into 2-tetrad conformers capped by a G-triad, similar to 22GT and 23AG_I14. In $2K^+$-species, the second potassium is either coordinated between a tetrad and a G-triad or electrostatically bound to the phosphate backbone.[22] Conversely, 23AG and 22CTA adopt two significantly more protected conformations that are more likely to contain three tetrads.

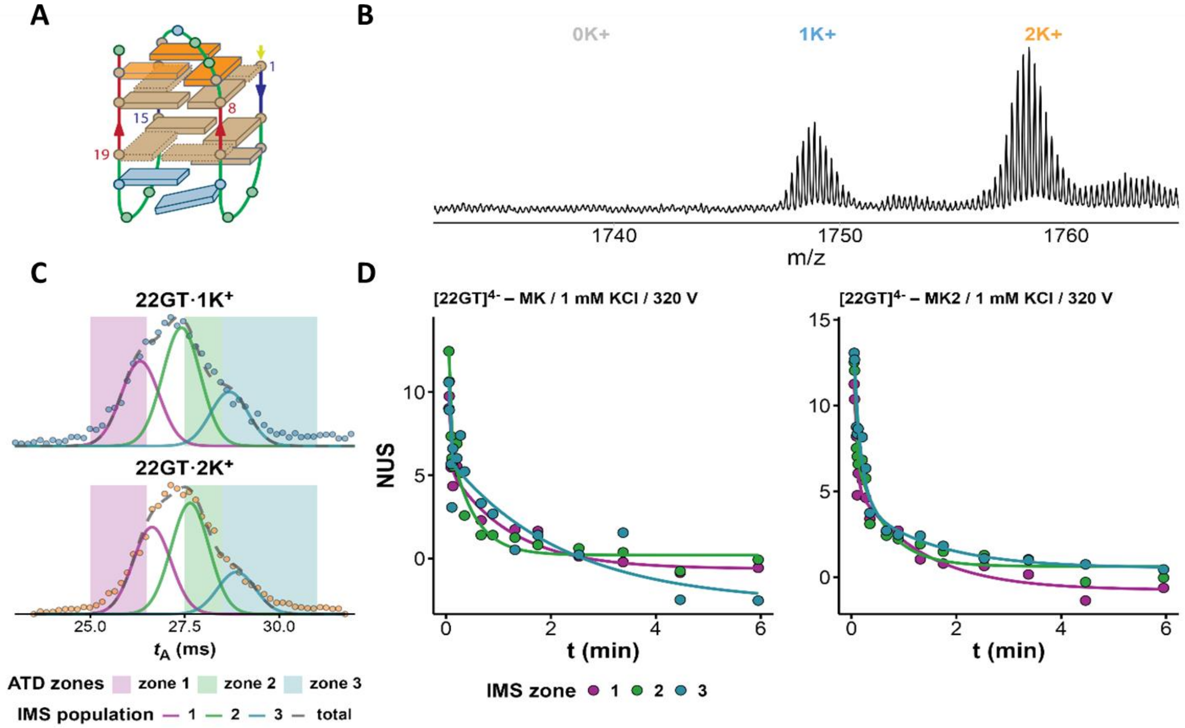


*Figure S190. HDX/IMS results on 22GT. (A) structure of the two-tetrad basket-type species formed by 22GT. The capping G-triad is indicated in orange. (B) Mass spectrum of 22GT showing the presence of both 1K⁺ and 2K⁺ species. (C) arrival-time distributions for the two K⁺ adducts. The similar shape led to the deconvolution of three zones. (D) HDX kinetics of the IMS-resolved species of the 1K⁺ and the 2K⁺ species.*
